# Modulation of the human cathelicidin dimerization interface separates antimicrobial activity from mammalian membrane disruption

**DOI:** 10.64898/2026.02.13.705689

**Authors:** John S. Albin, Dimuthu A. Vithanage, Wayne Vuong, Corey Johnson, Bradley L. Pentelute

## Abstract

The human *CAMP* gene product LL-37 is a prototypical cationic host defense peptide with potent activity against gram-negative bacteria. The development of LL-37 and other membrane-active host defense peptides as antibiotics, however, is limited in part by their activity against mammalian membranes. To better understand the structural features underlying LL-37 functions, we reconceived of LL-37 as a small protein and systematically ablated the sidechain content of entire surfaces while preserving helical character. This approach revealed that selected modifications within surface m4 retain wild-type levels of activity against gram-negative bacteria while disrupting activity against mammalian membranes by >100-fold. Separation of antimicrobial and mammalian membrane activities mapped primarily to residues I24 and L28, and was attributable to a combination of disrupted oligomerization and decreased hydrophobic content within the dimerization interface. Modulation of the LL-37 dimerization interface thus constitutes a rational pathway by which to engineer derivatives with improved therapeutic potential.

## Introduction

Most cationic host defense peptides (HDPs) are thought to exert their direct antimicrobial activities against gram-negative bacteria through the disruption of bacterial membranes(1–3). A classic example is the human cathelicidin, LL-37(4, 5). Initially expressed as a 16 kDa proprotein, this *CAMP* gene product undergoes proteolytic processing following neutrophil degranulation to release an antimicrobial 37-residue helical peptide starting with two Leu residues, hence the name LL-37(6, 7). Many prior studies have characterized the detailed mechanisms by which LL-37 and other membranolytic HDPs disrupt bacterial membranes(8–11), and this activity is ultimately thought to be protective against certain types of bacterial infections *in vivo* (*e.g.*, (12–14)).

Despite the potent activity of LL-37 against gram-negative bacteria, the development of antimicrobials based on this and other HDP templates has been challenging, in part due to incomplete selectivity for bacterial over mammalian membranes(15–20). Antibacterial activity is thought to depend on helical structure(10, 11, 21, 22), and multiple prior studies have described the assembly of LL-37 into oligomers, which may drive pore formation or other mechanisms of membrane disruption(14, 23–27). Although these structural factors are generally thought to correlate with membranolysis, it is unclear whether there may be a structural means by which to decouple antimicrobial activity from activity against mammalian cells.

To better understand the basis for the antimicrobial and hemolytic activities of full-length LL-37, we reconceived of LL-37 as a small protein with distinct surfaces imparting distinct functions. We then systematically ablated the sidechain content of entire surface regions while preserving helical character. Subsequent functional analysis of these mutants revealed that alteration of surface m4 yields mutants that retain antimicrobial activity comparable to wild-type LL-37 while demonstrating >100-fold decreases in hemolytic activity. Additional mutagenesis within the m4 surface identified the combination of residues I24 and L28 in mutant m15 as the primary drivers of this phenotypic separation.

Noting that changes in m4 overlap a previously described dimerization interface in LL-37(23, 24), we hypothesized that the disruption of oligomerization may contribute to the separability of antimicrobial and hemolytic functions in this classic helical amphipathic peptide. Further analysis by size exclusion chromatography (SEC) and biolayer interferometry (BLI) confirmed that the m4 and m15 mutants are deficient in their ability to form oligomeric structures, though reducing hemolytic activity depended also on reducing hydrophobic content within the dimerization interface. Rational modulation of residues within the LL-37 dimerization interface thus offers a pathway by which to engineer derivatives that separate antimicrobial activity from activity against mammalian membranes.

## Results

### LL-37 hemolytic and antimicrobial functions are separable

Prior investigations into LL-37 structure-activity relationships have focused primarily on the study of LL-37 fragments(28). Though commonly deployed and often useful, the study of such fragments risks the loss of structural context found in the intact peptide. We considered, however, that it may be possible to derive insight from the mutagenesis of multiple structurally clustered but linearly discontinuous residues in small but structured proteins like LL-37 and other HDPs. That is, in much the same way that larger proteins have distinct surfaces that impart distinct functions, we hypothesized that LL-37 might also consist of functionally distinguishable surfaces with associated functions that could be disrupted through clustered mutations.

To test this, we visually inspected a prior NMR structure of LL-37 and identified structurally contiguous residues that might cooperate in a given function(22). We then synthesized a series of LL-37 mutants in which every residue in each individual region was changed to Ala; a schematic summary of this strategy is shown in **Figure 1A**. Importantly, evaluation by circular dichroism (CD) spectroscopy showed that all of these mutants retain characteristic minima at 208 and 222 nm suggestive of retained helical structure; estimates of secondary structural content as calculated with BeStSel(29) indicated 100% helical content throughout. It is thus likely that these mutants retain helical structure comparable to wild-type LL-37, allowing for the isolation of sidechain effects through grouped Ala mutagenesis. **Supplementary Figure 1** shows the CD spectra for all **Figure 1** mutants as well as wild-type LL-37.

**Figure 1.**
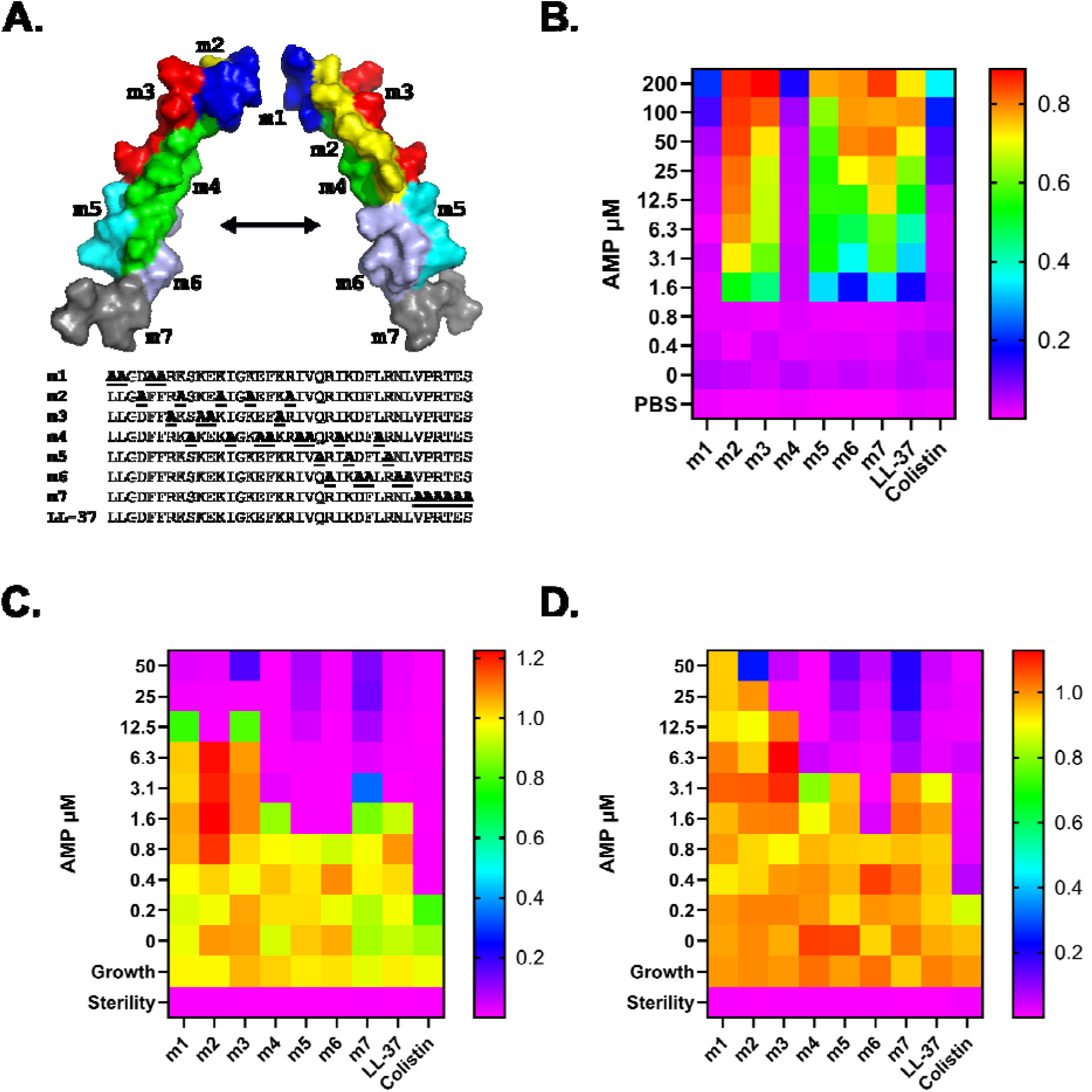
Mutations in region m4 separate LL-37 hemolytic and antimicrobial functions. **A.** Overview of mutagenesis strategy. Each mutant utilized has all residues within the putative structural surface changed to Ala, as shown in the associated sequences compared with wild-type LL-37. Regions are shown on a prior NMR structure 2K6O. **B.** Hemolysis assay quantifying the activity of each peptide relative to a 100% hemolysis control after 18 hours of incubation at 37 °C in the presence of different concentrations of each peptide. Warm colors indicate greater hemolysis as evidenced by absorbance at 414 nm for free hemoglobin in well supernatants; cool colors indicate low or no hemolysis. **C-D.** Antimicrobial susceptibility testing of *E. coli* (**C**) and *Pseudomonas* (**D**), which shows the OD600 after 18 hours of incubation at 37 °C in the presence of different concentrations of each peptide. Warm colors indicate growth; cool colors indicate no or low growth. Data shown are the mean of three independent experiments.

In characterizing these clustered surface mutants, we started with hemolytic activity against sheep blood as a surrogate for activity against mammalian cell membranes. As shown in **Figure 1B**, m1 and m4 among our initial set of seven clustered surface mutants demonstrated substantial reductions in hemolytic activity compared with wild-type LL-37, rendering these comparable in this measure of toxicity to the control antibiotic colistin. Further evaluation of the antimicrobial activity of these clustered surface mutants via broth dilution antimicrobial susceptibility testing, however, showed that only m4 retained activity comparable to wild-type LL-37, while m1 as well as m2 and m3 lost antimicrobial activity against both *Escherichia coli* and *Pseudomonas aeruginosa* (**Figure 1C-D**). We have expanded upon the contributions of N-terminal residues to LL-37 antimicrobial activity in separate manuscripts(30, 31). The current study focuses on this second observation: the separation of hemolytic and antimicrobial activity by mutant m4. Thus, clustered surface mutants yield new insights into the structural basis of LL-37 functional activities, and mutations within the m4 region ablate LL-37 hemolytic activity without affecting antimicrobial activity.

**Supplementary Figure 2** shows the hemolysis and antimicrobial susceptibility endpoint assays from **Figure 1** in histogram format with error bars. **Supplementary Figures 3-11** (hemolysis) and **12-20** (antimicrobial susceptibility) show the 18-hour kinetic curves associated with each endpoint measurement.

### Separation of hemolytic and antibacterial functions in LL-37 is mediated by amino acid changes at I24 and L28

Eight residues in mutant m4 are changed to Ala compared with wild-type LL-37. To better define those responsible for separation of hemolytic and antibacterial functions in LL-37, we made a series of overlapping mutants as shown in **Figure 2A** and subjected these to testing for hemolytic and antibacterial activities in a fashion similar to **Figure 1**. As shown in **Figure 2B**, disruption of hemolytic activity was initially localized to the four residues of the m10 region – I20, V21, I24, and L28, though mutation of any one of these four residues to Ala as in m11-m14 did not ablate hemolytic activity. In comparing mutants m8, m9, and m10, however, the only residues unique to m10 were I24A and L28A. We therefore prepared mutant m15 combining both I24A and L28A, which demonstrated hemolytic activity comparable to m4 (**Figure 2B**). Antimicrobial activity remained largely the same across mutants tested in this series of experiments as shown in **Figure 2C-D**. Thus, the separation of hemolytic and antimicrobial activity in LL-37 localizes primarily to residues I24 and L28.

**Figure 2.**
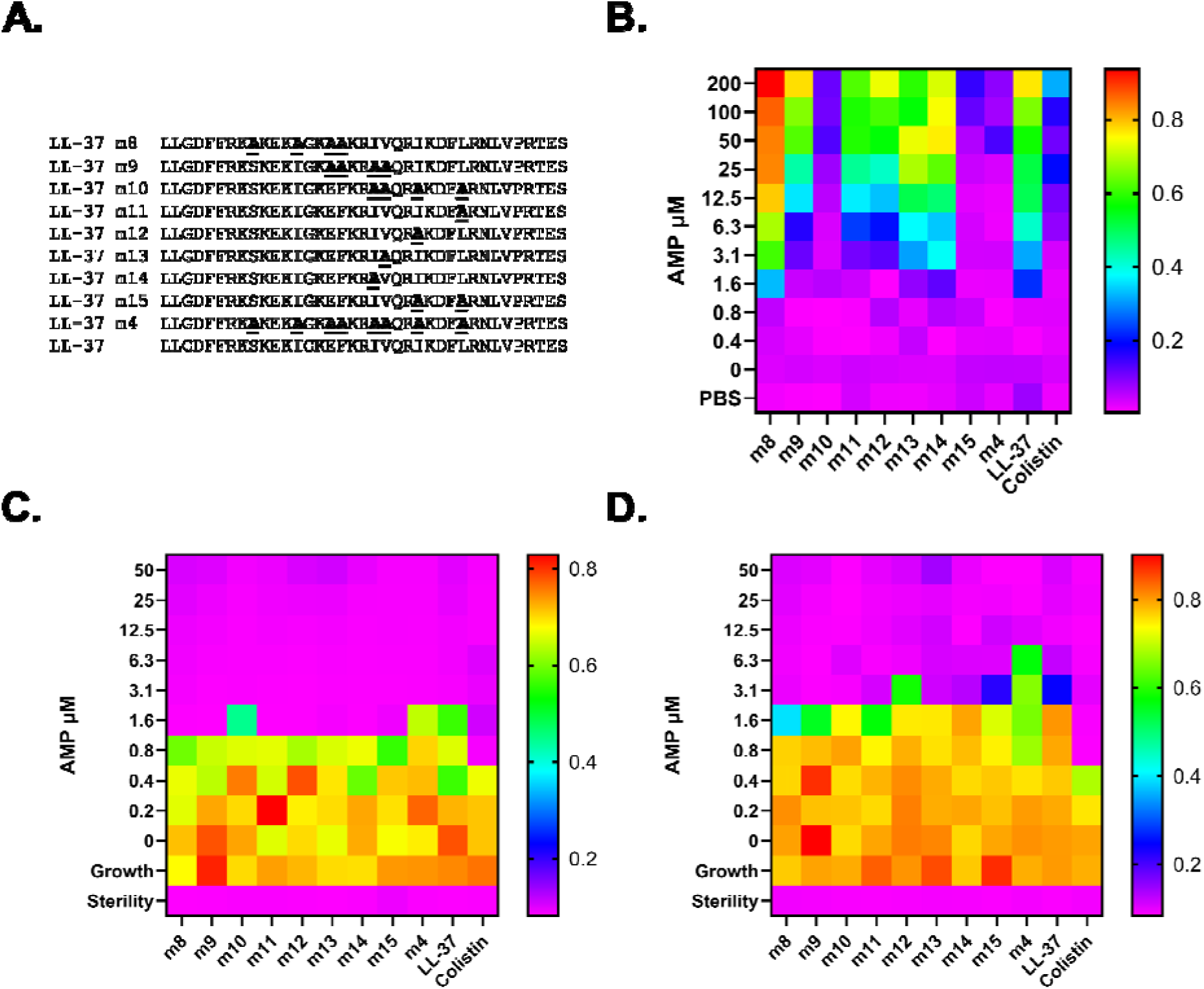
Separation of hemolytic and antimicrobial activities maps primarily to residues I24 and L28. **A.** Overlapping mutants were preparared as shown to localize the separation of function residues within LL-37 m4. **B.** Hemolysis assay quantifying the activity of each peptide relative to a freeze-thaw 100% hemolysis control. Warm colors indicate greater hemolysis as evidenced by absorbance at 414 nm for free hemoglobin in well supernatants; cool colors indicate low or no hemolysis. **C-D.** Antimicrobial susceptibility testing of *E. coli* (**C**) and *P. aeruginosa* (**D**), which shows the OD600 after 18 hours of incubation at 37 °C in the presence of different concentrations of each peptide. Warm colors indicate growth; cool colors indicate no or low growth.

**Supplementary Figure 21** shows CD spectra of mutants m8-m15, all of which demonstrated helical content comparable to wild-type LL-37. **Supplementary Figure 22** provides a histogram depiction of the data in **Figure 2** including error bars. 18-hour kinetic curves associated with each endpoint measurement are provided in **Supplementary Figures 23-33** (hemolysis) and **34-44** (antimicrobial susceptibility testing).

### Contributions of oligomerization and hydrophobic content to LL-37 hemolytic activity

On visualizing mutations associated with separation of hemolytic and antibacterial activities in prior crystal structures of LL-37, we noted that many of these were located in or adjacent to a core dimerization interface(23, 24). We therefore hypothesized that mutants with diminished hemolytic activity might also have a diminished propensity for oligomerization.

To test this, we first subjected LL-37 and m4 to SEC. As shown in **Supplementary Figure 45**, m4 showed greater retention and less tailing compared with LL-37. We next extended this analysis to mutants m1-m7. As shown in **Figure 3A**, migration patterns for most mutants were similar to that of wild-type LL-37. In contrast, m4 (and m6) had longer retention times and less tailing. This was further reinforced by evaluation of SEC traces among mutants m8-m15. As shown in **Figure 3B**, mutants with attenuated hemolytic activity such as m10 and m15 demonstrated peaks superimposable with that of m4. With the exceptions of the intermediate migration profiles observed in point mutants m13 and m14, mutants generally followed a clear dichotomy between LL-37-like species and m4-like species. On comparison with size controls, and accounting for the disproportionately large hydrodynamic radius per molecular weight of a helical peptide such as LL-37, this was most consistent with migration as dimers among LL-37-like species and as monomers among m4-like species (**Supplementary Table 1**).

**Figure 3.**
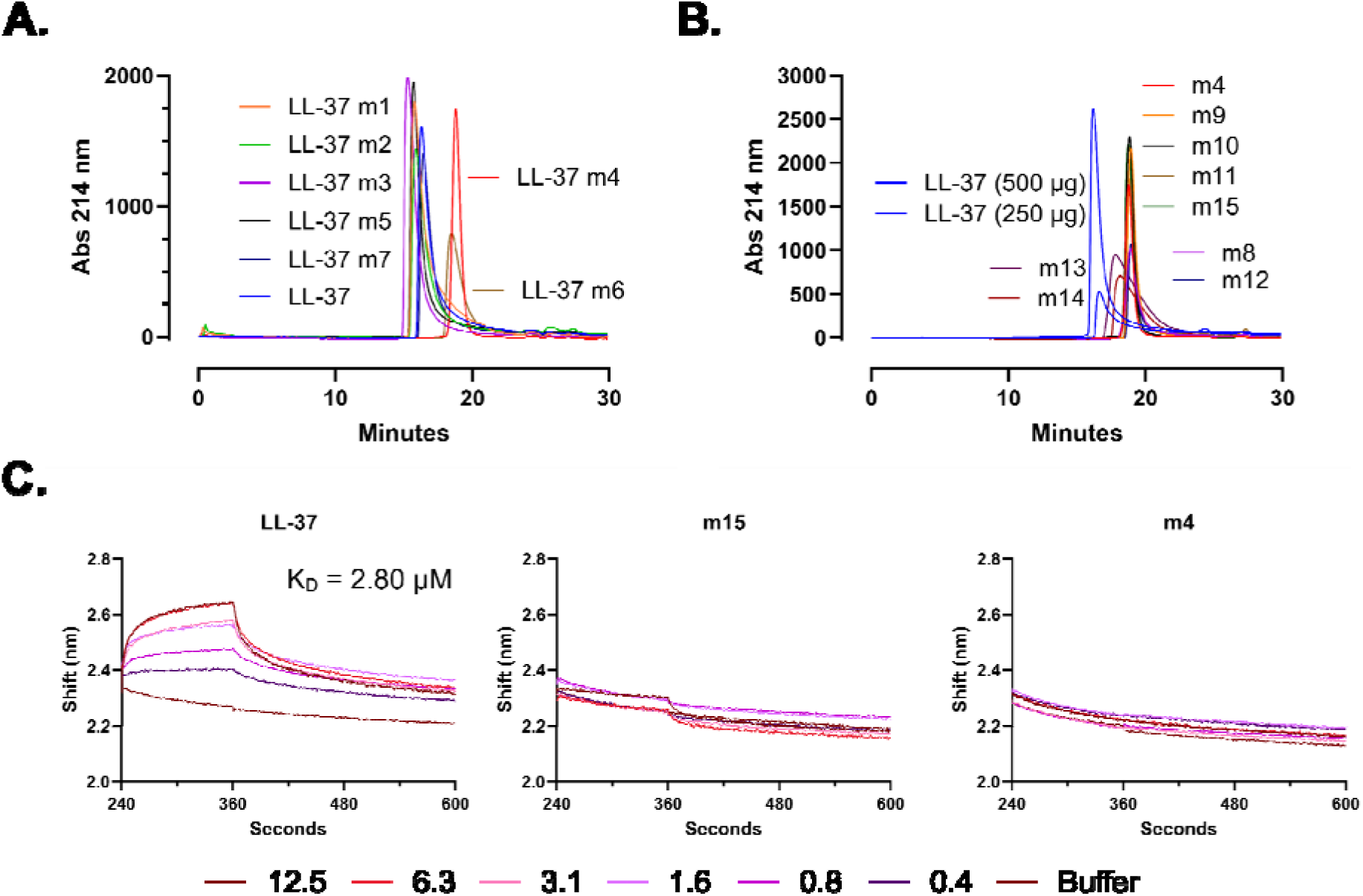
LL-37 mutants with decreased hemolytic activity have a decreased propensity for oligomerization. **A.** Size exclusion chromatography (SEC) of LL-37 surface mutants in PBS + 3.5% IPA as in Figure 1. **B.** SEC of overlapping mutants from Figure 2. **C.** Biolayer interferometry (BLI) showing the shifts associated with incubating the indicated analyte with a probe consisting of biotinylated LL-37.

On further analysis, it was evident that, while all mutants with diminished hemolytic activity were m4-like species, not all peptides migrating as m4 demonstrated diminished hemolytic activity – for example, m6 in **Figure 3A** and m8, m9, m11, and m12 in **Figure 3B**. This is consistent with an additional contribution of overall hydrophobic character in the dimerization interface to hemolytic activity. For instance, mutants m10 and m15 each change 2-4 hydrophobic residues at or immediately adjacent to I24 and L28 and have attenuated hemolytic activity. In contrast, mutants that migrate in a pattern similar to m4 but retain hemolytic activity each have either: i) Only one change in a hydrophobic sidechain at I24 (m12) or L28 (m11); or ii) 2-3 changes in hydrophobic residues in combinations that do not include I24A + L28A (m6, m8, m9). Thus, disruption of oligomerization alone is not sufficient to ablate hemolytic activity, and it is likely hydrophobic content in the dimerization interface also contributes to activity against mammalian membranes.

To better quantify the propensity for self-interaction among LL-37 mutants, we prepared an N-biotinylated derivative of LL-37 and used this as a probe in biolayer interferometry (BLI) experiments with LL-37, m15, or m4 as analytes. As shown in **Figure 3C**, LL-37 demonstrated a characteristic on-off curve with an average K_D_ across three independent experiments of 2.80 µM. By contrast, incubation of the LL-37 probe with m15 or m4 analytes yielded no measurable binding constant. Thus, hemolysis-deficient mutants that migrate as monomers by SEC do not efficiently form intermolecular interactions with LL-37, suggesting disruption of the interaction surface. **Supplementary Figure 46** shows the raw traces from three independent BLI experiments.

### Distinct mammalian membranes demonstrate differential susceptibility to oligomerization-deficient LL-37 derivatives

Within the standard model system that we use, the decreased activity of oligomerization-deficient LL-37 mutants toward sheep red blood cells (RBCs) was substantial. In **Figures 1** and **2**, for example, the ratio of the maximum peptide concentration with < 20% hemolysis for m4 relative to LL-37 is 125, slightly better than colistin at 62.5. To better define the activity spectrum of these mutants, we next carried out a series of hemolysis assays using a wider variety of sample types, including sheep and cow defibrinated blood, human whole blood from each of two donors, and isolated sheep and human RBCs.

As shown in **Figure 4A**, the extent of membrane disruption differed by sample type, though fold-improvement relative to wild-type LL-37 generally followed the same trend of colistin > m4 > m15 > LL-37. As discussed above, this may reflect additional contributions of the relative hydrophobicity of amino acid content at the LL-37 dimerization interface. Overall effect sizes are quantified in **Supplementary Table 2**, where the average fold-change in the highest peptide concentration yielding < 20% hemolysis within each blood type relative to LL-37 was 51-fold for m15, 135-fold for m4, and 208-fold for colistin, albeit with substantial deviation due to differences among sample types. In particular, LL-37 derivatives showed less fold-improvement in human cell types relative to other species. The basis for this diminished fold-improvement relative to other species is unclear, though some instances such as Human 1 suggest sample-specific fragility. Regardless, the relative trends in membranolytic activity among LL-37 derivatives as stated above hold true across conditions. **Supplementary Figure 47** provides the data from **Figure 4A** in histogram format. Kinetic hemolysis curves for each condition from triplicate experiments are provided in **Supplementary Figures 48-71**.

**Figure 4.**
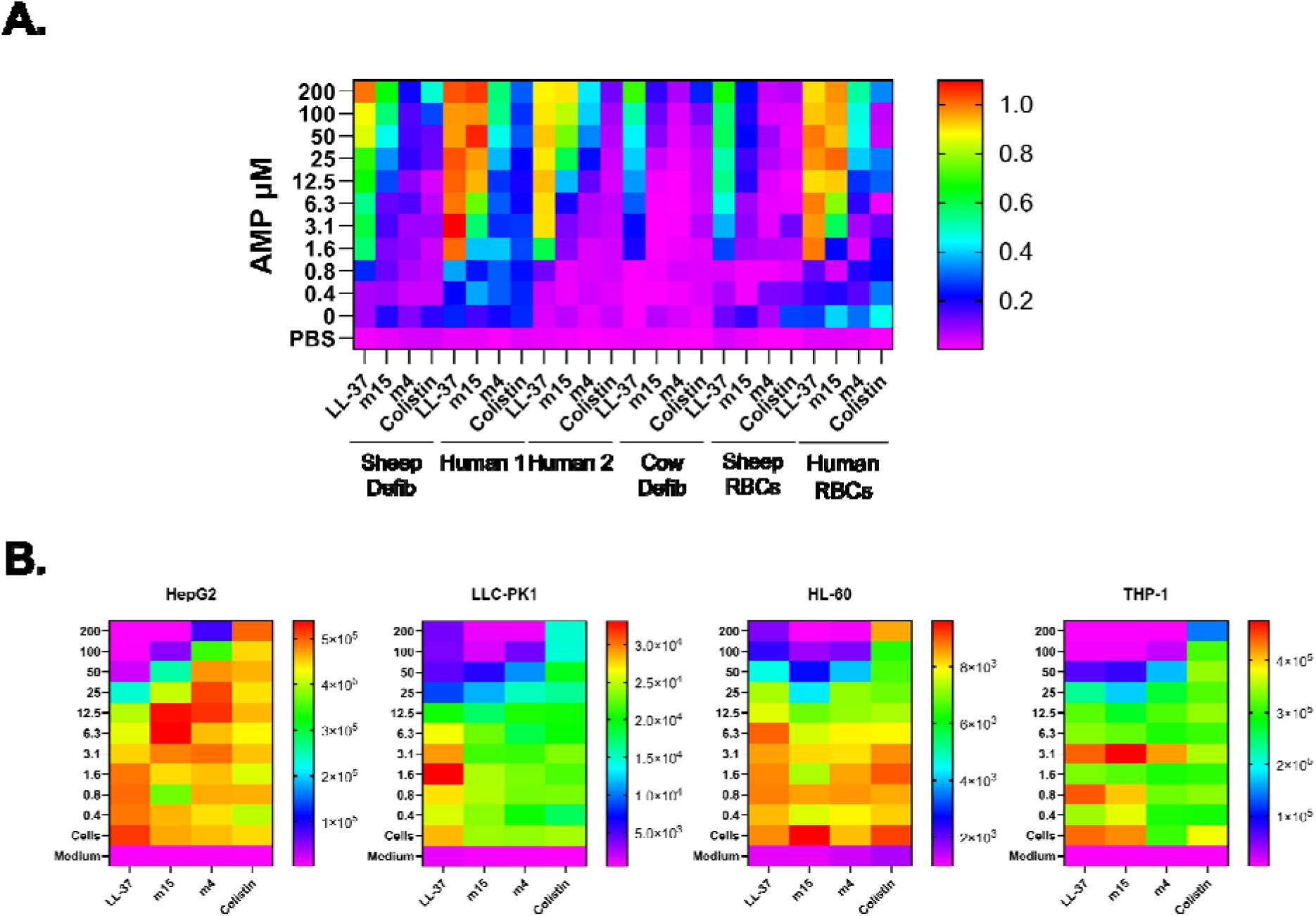
Effect of oligomerization deficient mutants on different cell types. **A.** Extension of hemolysis assays to the sample types shown. Sheep and cow blood are defibrinated. Human blood is purchased whole blood from each of two donors. Sheep and human RBCs indicate isolated RBCs. **B.** Luminescence toxicity assay with each of the indicated cell types.

To further assess the effects of the m15 and m4 mutants on metabolically active cells relative to LL-37, we carried out RealTime Glo (Promega) toxicity assays using HepG2, LLC-PK1, HL-60, and THP-1 cells. While some cell types such as HepG2 demonstrated a gradual decrease in toxicity by roughly one dilution each from LL-37 to m15 to m4 to colistin, the differences between oligomerization-deficient mutants and wild-type LL-37 were smaller in this context due to overall lower baseline toxicity of LL-37 (**Figure 4B**). Thus, the relatively low baseline toxicity of LL-37 itself and of its derivatives limits the dynamic range available for interpretation in metabolically active cells. Residual toxicity of LL-37 may be mediated, in part, by interactions other than those occurring directly with membranes, consistent with the reported interactions of LL-37 with protein receptors (*e.g.*, (32–36)). Histogram depictions of the data in **Figure 4B** are provided in **Supplementary Figure 72**. Kinetic curves leading into the endpoint measures in **Figure 4B** are shown in **Supplementary Figures 73-88**.

### Limited effect of divalent cation concentrations on the relative hemolytic activity of LL-37 derivatives

Divalent cation concentrations have been found previously to influence the antimicrobial activity of LL-37 and other cationic antimicrobials(37–39). To determine whether divalent cation concentrations would influence the relative activity of distinct mutants against mammalian membranes, we carried out hemolysis assays with defibrinated sheep blood in a fashion similar to **Figures 1-2**. In addition to PBS without added divalent cations, however, we supplemented calcium and magnesium content to Low, Medium, or High levels as defined under Methods. As shown in **Figure 5**, the relative hemolytic activity among LL-37 and derivatives thereof remained largely unchanged across most divalent cation concentrations. In this series, for example, LL-37 hemolytic activity was approximately 2-3 dilutions greater than that of m15, which in turn was 2-3 dilutions more active than m4. Results under High divalent cation concentrations demonstrated substantially more variability (**Supplementary Figure 89**), though again with diminished activity of the m15 and m4 mutants relative to wild-type. Thus, there was little overall effect of the divalent cation concentrations tested here on hemolysis levels among LL-37 and the derivatives tested. **Supplementary Figures 90-105** show the kinetic hemolysis curves associated with the endpoint results shown in **Figure 5**.

**Figure 5.**
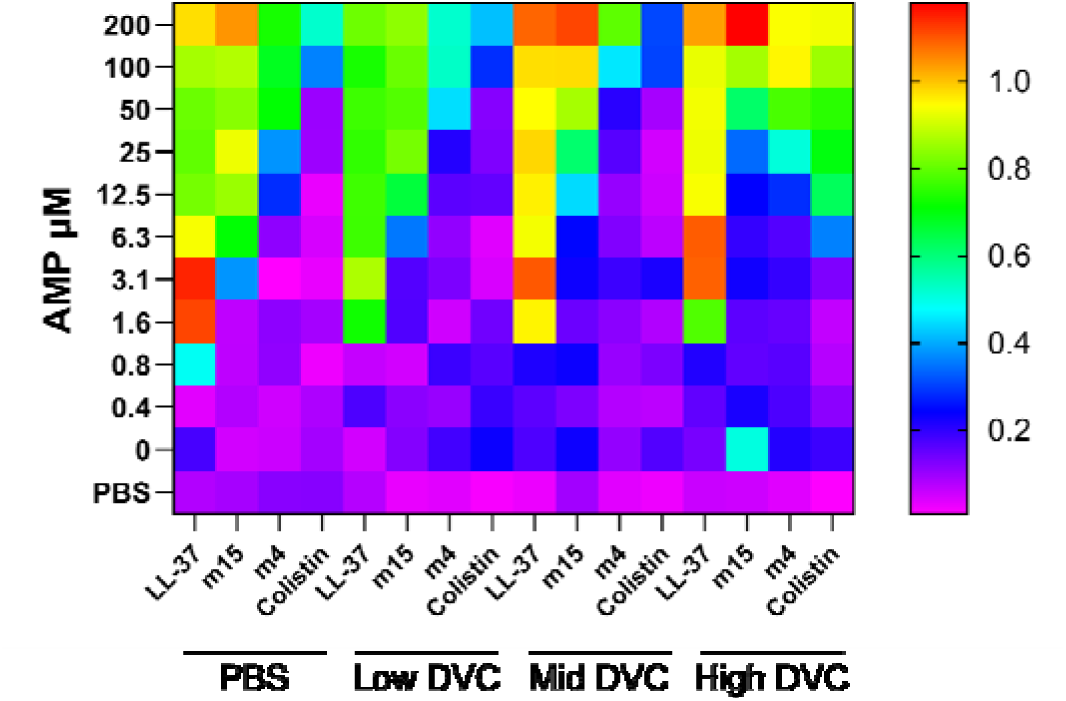
Effect of divalent cation concentrations on hemolytic activity. Hemolysis assays were carried out as in Figures 1-2 using defibrinated sheep blood in PBS (No DVCs) and PBS supplemented with each of three concentrations of Ca^2+^ / Mg^2+^ (Low, Mid, High).

## Discussion

The engineering of peptides with antimicrobial properties from any source often focuses on the modulation of biophysical properties such as length, charge, and hydrophobicity. In this work, we propose an alternative approach for naturally-occurring host-derived sequences that regards HDPs as small proteins. Starting from the hypothesis that the classic HDP LL-37 has distinct surfaces that impart distinct functions, we demonstrate the viability of separating these functions by systematically disrupting entire surfaces while preserving overall helical content. In addition to validating the importance of selected N-terminal residues for antimicrobial activity(30, 31), this approach leads us to the central observation of this paper: the separability of antimicrobial and hemolytic functions in LL-37 oligomerization-deficient mutants with variable degrees of hydrophobic blunting in the dimerization interface.

In our studies, all mutants with diminished hemolytic activity also display diminished capacity for oligomerization. The converse, however, is not necessarily true – we do observe mutants with diminished capacity for oligomerization that have intact hemolytic activity. This likely represents an additional contribution of hydrophobic content within the dimerization interface to activity against mammalian membranes. For example, although m15 itself displays substantially less activity against mammalian membranes than LL-37, m4 has an additional 2-3 dilutions less activity than m15. The most prominent difference between these two, neither of which displays detectable interaction with LL-37 by BLI, is the replacement of an additional four hydrophobic residues in the m4 dimerization interface with Ala (I13, F17, I20, V21) in addition to the core changes at I24 and L28 in m15.

Despite the likely contribution of hydrophobic content in the dimerization interface to the functional loss of hemolytic activity, there is also positional dependence to these phenotypes that would argue against hydrophobic content alone as an explanation for decreased hemolytic activity. Specifically, m8 and m9 both behave as monomers by SEC and lack 2-3 hydrophobic residues relative to LL-37, yet remain strongly hemolytic. By contrast, the loss of the two specific hydrophobic residues at I24 and L28 in m15 confers the majority of the diminished hemolytic activity observed in m4. It is unclear whether it may be possible to diminish hydrophobic content at I24 and L28 while preserving oligomerization, or conversely to preserve equivalent hydrophobic content while disrupting oligomerization.

For the purpose of engineering improved derivatives of LL-37 for therapeutic applications, the path forward is the same regardless of the relative contributions of oligomerization and hydrophobic content to the activity of LL-37 and derivatives thereof against mammalian membranes. The overall degree of diminished mammalian membrane toxicity in the mutants described here approaches that of the approved control antibiotic colistin, which though still toxic(40–42) may be of use as an *in vitro* benchmark for when something may be approaching utility (**Supplementary Table 2**). To meet and exceed this minimum bar to entry, we anticipate that additional modifications at hydrophobic residues within the dimerization interface will be of greatest utility. In doing so, it may be necessary to test for emergent effects among large numbers of pooled peptides rather than individual candidates(43, 44).

We report here that the antimicrobial activity of oligomerization-deficient mutants against gram-negative bacteria remains intact. What we have not reported is how these mutations impact the ability of LL-37 derivatives to permeabilize gram-negative membranes. This is the subject of a separate manuscript derived from this same series of studies(45).

## Methods

### Peptide synthesis and purification

Methods for peptide synthesis and purification are generally as previously described (*e.g.*, (46–48)). Peptides used in this study were made on a third-generation automated flow synthesizer following preloading of 0.45 mmol/g 4-(4-Hydroxymethyl-3-methoxyphenoxy)butyric acid (HMPB) resin (ChemMatrix) with the relevant C-terminal amino acid using overnight reaction with N,N’-Diisopropylcarbodiimide (DIC) and 4-dimethylamino pyridine (DMAP).

Following flow synthesis, resin was cleaved by treatment with Reagent K (82.5% trifluoroacetic acid (TFA), 5% water, 5% phenol, 2.5% 1,2-ethanedithiol) for 2 hours at room temperature followed by peptide precipitation with ether chilled on dry ice. Precipitated peptide was then resuspended in 50% water / 50% acetonitrile with 0.1% TFA, flash frozen, and lyophilized. Crude peptide was purified on reverse phase columns using a Biotage Selekt flash chromatography system. Analytical data associated with peptides used in the course of these studies are provided in **Supplementary Figures 106-126**.

### Circular dichroism spectroscopy

Samples were prepared for circular dichroism spectroscopy by dissolving peptides at 0.5 mg/mL in water and then diluting to 20% v/v trifluoroethanol. Data were collected on an Aviv 420 CD Spectrometer at the indicated temperatures in 0.1 cm pathlength cuvettes. Calculation of secondary structural content was completed using BeStSel(29).

### Size exclusion chromatography

Size exclusion chromatography was performed on an Agilent 1260 Infinity II Bio-Inert liquid chromatography system using a Superdex Increase 75 100/300 GL column (Cytiva) and PBS buffer supplemented with 3.5% isopropanol with a flow rate of 0.8 mL / minute over 30 minutes and monitoring at 214 nm.

### Biolayer interferometry

Biolayer interferometry was performed on a GatorPlus bio-layer interferometry system (GatorBio) using streptavidin probes. Following an initial baseline measure for 30 seconds, probes were loaded with 800 nM N-biotinylated LL-37 in PBS for 180 seconds. Following another 30 second baseline read, association was measured for 120 seconds followed by dissociation for 240 seconds. Analyte concentration series of LL-37, m4, or m15 were 12.5 µM, 6.3 µM, 3.1 µM, 1.6 µM, or 0.8 µM with no analyte (buffer) and no probe (analyte only at 12.5 µM) controls. Calculation of interaction kinetics was completed in GatorOne software associated with the instrument.

### Antimicrobial susceptibility testing

Testing for antimicrobial activity of peptides of interest was completed as per Clinical and Laboratory Standards Institute methods with previously described modifications for the use of cationic peptides, specifically the use of polypropylene plates (Greiner) and unadjusted Mueller Hinton Broth (Difco) (39). These methods were scaled down to either 35 or 50 µL per well in 384-well plates. In general, peptide was added initially as a 10x concentrate in water followed by the addition of a concentrated broth and cell mix to a final 1x concentration of all reagents. Following plating, growth was monitored by OD600 on a Tecan Spark plate reader over the course of 18 hours at 37 °C, during which time the plate was incubated in a Spark Large Humidity Cassette to minimize evaporation. The bacterial strains used in these studies were *E. coli* ATCC 25922 and *Pseudomonas* ATCC 27853, which are quality control strains for antimicrobial susceptibility testing(49).

### Hemolysis Assays

Hemolysis assays were carried out by treating blood samples with a peptide of interest over a range of concentrations. In general, blood was spun for 10 minutes at 800 rcf and supernatant decanted to clearance, after which cells were brought up in PBS at 1% v/v for final dilution to 0.5% with a 2x concentration of peptide dissolved in PBS. Final well volumes were 50 µL each in 384-well polypropylene plates (Greiner). Reactions were then monitored over 18 hours by OD600 as a kinetic marker of hemolysis. In addition to this kinetic marker, endpoint measures of hemolysis were made by adding 25 µL of PBS to each well at the end of incubation, spinning plates for 10 minutes at 800 rcf, and then removing 25 µL of supernatant for evaluation of free hemoglobin levels by absorbance at 414 nm. Types of blood used in these studies included defibrinated sheep and cow blood (Hardy Diagnostics), purchased human whole blood from two separate donors (ZenBio), sheep RBCs (Innovative Research), and human RBCs (Rockland Immunochemicals).

Quantification of hemolysis was completed by comparison to a thrice freeze-thawed control of the same blood used in plating set to 100%. Thus, percentages are not a literal percentage of cells, but rather a bulk percentage of signal. The choice of the percentage cutoff for determining samples with substantial hemolysis (20% for most blood types with the exception of two samples set to 40%) was based on the percentage at which there was a clear cutoff between signal and noise in the colistin concentration gradient, the premise being that colistin can be used *in vivo* and thus provides a rough surrogate for approximating where a peptide is on the toxicity spectrum from LL-37 to colistin.

PBC without divalent cations was used in hemolysis assays described here with the excetion of those in **Figure 5**. Divalent cation concentrations used to supplement PBS in **Figure 5** were chosen to reflect levels commonly used in antimicrobial susceptibility testing. Low approximates the content of unadjusted MHB (4.9 mg/L Mg^2+^, 4.0 mg/L Ca^2+^), while Mid approximates the content of cation adjusted MHB (12.2 mg/L Mg^2+^, 20.0 mg/L Ca^2+^(39, 50). The High condition approximates the divalent cation levels found in human blood (19.4 mg/L Mg^2+^, 48.1 mg/L Ca^2+^(51).

### Luminescence toxicity assays

Luminescence toxicity assays were completed using RealTime Glo reagents per the manufacturer’s instructions. Following initial pilot studies, HepG2 cells were plated at a density of 5,000 cells per well, while LLC-PK1, HL-60, and THP-1 were plated at 20,000 cells/well in 25 µL per well of white clear-bottom plates. These were then allowed to settle for approximately 12 hours, after which 25 µL of RealTime Glo reagents in the appropriate medium and 5 µL of a 10x concentration of peptide in water were added. Wells were then incubated over 18 hours in a Tecan Spark Plate reader with Small Humidity Cassette to minimize evaporation. HepG2 cells were maintained in EMEM medium (ATCC) with 10% FBS. LLC-PK1 cells were maintained in Medium 199 (Sigma) with 5% FBS. THP-1 cells were maintained in RPMI medium (ATCC) with 10% FBS and supplemented with fresh β mercaptoethanol to 0.05 mM final concentration at the time of usage (Gibco). HL-60 cells were maintained in IMDM medium (ATCC) with 20% FBS. Adherent cells were split using Trypsin-EDTA (ATCC).

## Supporting information

Combined_Tables_S1-2_Figures_S1-126

## Acknowledgements

This work has been funded in by the National Institute of Allergy and Infectious Diseases (T32 AI007061 and K08 AI166345 to JSA; U19 AI142780 to BLP) and by the Cystic Fibrosis Foundation (ALBIN19F0, ALBIN21Q0, ALBIN22A0-KB to JSA).

## Author Contributions

JSA designed and conducted the experiments and wrote the paper. DAV, WV, and CJ performed the experiments. BLP guided the experiments and wrote the paper.

