## Supplementary material for "Modulation of the human cathelicidin dimerization interface separates antimicrobial activity from mammalian membrane disruption": Combined_Tables_S1-2_Figures_S1-126

**Table of Contents:**


| | |
|---|---|
| Table S1 Values for Figure 3B molecular weight estimation | S3-S4 |
| Table S2 Relative decrease in hemolytic activity across species | S5-S6 |
| Figure S1 CD spectroscopy of Figure 1 mutants | S7-S8 |
| Figure S2 Histogram depiction of data in Figure 1 | S9-S10 |
| Figures S3-S11 Kinetic hemolysis curves associated with Figure 1 | S11-S28 |
| Figures S12-S20 Kinetic microbial growth curves associated with Figure 1 | S29-S46 |
| Figure S21 CD spectroscopy of Figure 2 mutants | S47-S48 |
| Figure S22 Histogram depiction of data in Figure 2 | S49-S50 |
| Figures S23-S33 Kinetic hemolysis curves associated with Figure 2 | S51-S72 |
| Figures S34-S44 Kinetic microbial growth curves associated with Figure 2 | S73-S94 |
| Figure S45 SEC profiles of LL-37 and m4 | S95-S96 |
| Figure S46 Raw BLI traces for PEG4-LL-37 with LL-37, m15, or m4 | S97-S98 |
| Figure S47 Histogram depiction of data in Figure 4A | S99-S100 |
| Figures S48-S71 Kinetic hemolysis curves associated with Figure 4A | S101-S148 |
| Figure S72 Histogram depiction of data in Figure 4B | S149-S150 |
| Figures S73-S88 Kinetic luminescence toxicity assay associated with Figure 4B | S151-S182 |
| Figure S89 Histogram depiction of Figure 5 data | S183-S184 |
| Figures S90-S105 Kinetic hemolysis curves associated with Figure 5 | S185-S216 |
| Figures S106-S126 Analytical data associated with peptide synthesis | S217-S258 |

| Standards | MW | RT | EV | Kav | LogMW |
|---|---|---|---|---|---|
| Thyroglobulin | 670000 | 10.181 | 8.145 | | |
| γ-globulin | 158000 | 10.671 | 8.537 | 0.02543 | 5.19866 |
| Ovalbumin | 44000 | 13.194 | 10.555 | 0.15635 | 4.64345 |
| Myoglobin | 17000 | 16.575 | 13.260 | 0.33179 | 4.23045 |
| Vitamin B12 | 1350 | 23.177 | 18.542 | 0.67437 | 3.13033 |

$y = -0.3203x + 1.6744$ $R^2 = 0.9942$

| Peptide | MW | RT | EV | Kav | LogMW | $10^{(LogMW)}$/m4 |
|---|---|---|---|---|---|---|
| LL-37 (500 µg) | 4493.3 | 16.185 | 12.948 | 0.31155 | 4.25492 | 2.62 |
| LL-37 (250 µg) | 4493.3 | 16.626 | 13.301 | 0.33443 | 4.18348 | 2.23 |
| m4 | 4146.8 | 18.771 | 15.017 | 0.44574 | 3.83597 | 1.00 |
| m8 | 4301.1 | 18.900 | 15.120 | 0.45243 | 3.81508 | 0.95 |
| m9 | 4289.0 | 18.928 | 15.142 | 0.45388 | 3.81054 | 0.94 |
| m10 | 4339.0 | 18.840 | 15.072 | 0.44932 | 3.82480 | 0.97 |
| m11 | 4451.2 | 18.834 | 15.067 | 0.44901 | 3.82577 | 0.98 |
| m12 | 4451.2 | 18.945 | 15.156 | 0.45477 | 3.80779 | 0.94 |
| m13 | 4465.2 | 17.811 | 14.249 | 0.39592 | 3.99150 | 1.43 |
| m14 | 4451.2 | 18.153 | 14.522 | 0.41367 | 3.93609 | 1.26 |
| m15 | 4409.1 | 18.759 | 15.007 | 0.44511 | 3.83792 | 1.00 |

**Supplementary Table 1 – Values associated with molecular weight estimation for peptides in Figure 3B.** Observed retention times (RT) and elution volumes (EV) used in the calculation of Kav and conversion to molecular weight (MW) are shown for BioRad gel filtration standards and test peptides depicted in **Figure 3B**.

| Sheep Defib | < 20% Fold Δ | |
|---|---|---|
| LL-37 | 0.4 | |
| m15 | 6.3 | 15.8 |
| m4 | 100 | 250 |
| Colistin | 50 | 125 |

| Cow Defib | < 20% Fold Δ | |
|---|---|---|
| LL-37 | 0.8 | |
| m15 | 200 | 250 |
| m4 | 200 | 250 |
| Colistin | 100 | 125 |

| Human 1 | < 40% Fold Δ | |
|---|---|---|
| LL-37 | 0.8 | |
| m15 | 1.6 | 2 |
| m4 | 25 | 31.3 |
| Colistin | 200 | 250 |

| Sheep RBCs | < 20% Fold Δ | |
|---|---|---|
| LL-37 | 0.8 | |
| m15 | 25 | 31.3 |
| m4 | 200 | 250 |
| Colistin | 200 | 250 |

| Human 2 | < 20% Fold Δ | |
|---|---|---|
| LL-37 | 0.8 | |
| m15 | 3.1 | 3.9 |
| m4 | 12.5 | 15.6 |
| Colistin | 200 | 250 |

| Human RBCs | < 40% Fold Δ | |
|---|---|---|
| LL-37 | 0.8 | |
| m15 | 1.6 | 2 |
| m4 | 12.5 | 15.6 |
| Colistin | 200 | 250 |

| | Avg. | SD |
|---|---|---|
| m15 | 51 | 98 |
| m4 | 135 | 126 |
| Colistin | 208 | 65 |

**Supplementary Table 2 – Fold-improvement in hemolytic activity relative to LL-37 across species.** For each sample type, a number indicating the greatest peptide concentration in µM at which hemolysis levels are < 20% or < 40% is given. The choice of threshold is generally based on the overall propensity of a given sample for hemolysis in the presence of colistin, which despite itself being a relatively toxic compound *in vivo* serves as a reference point. A second column divides this concentration by the value obtained for LL-37 on that sample type to estimate a fold-improvement in hemolytic activity relative to LL-37.

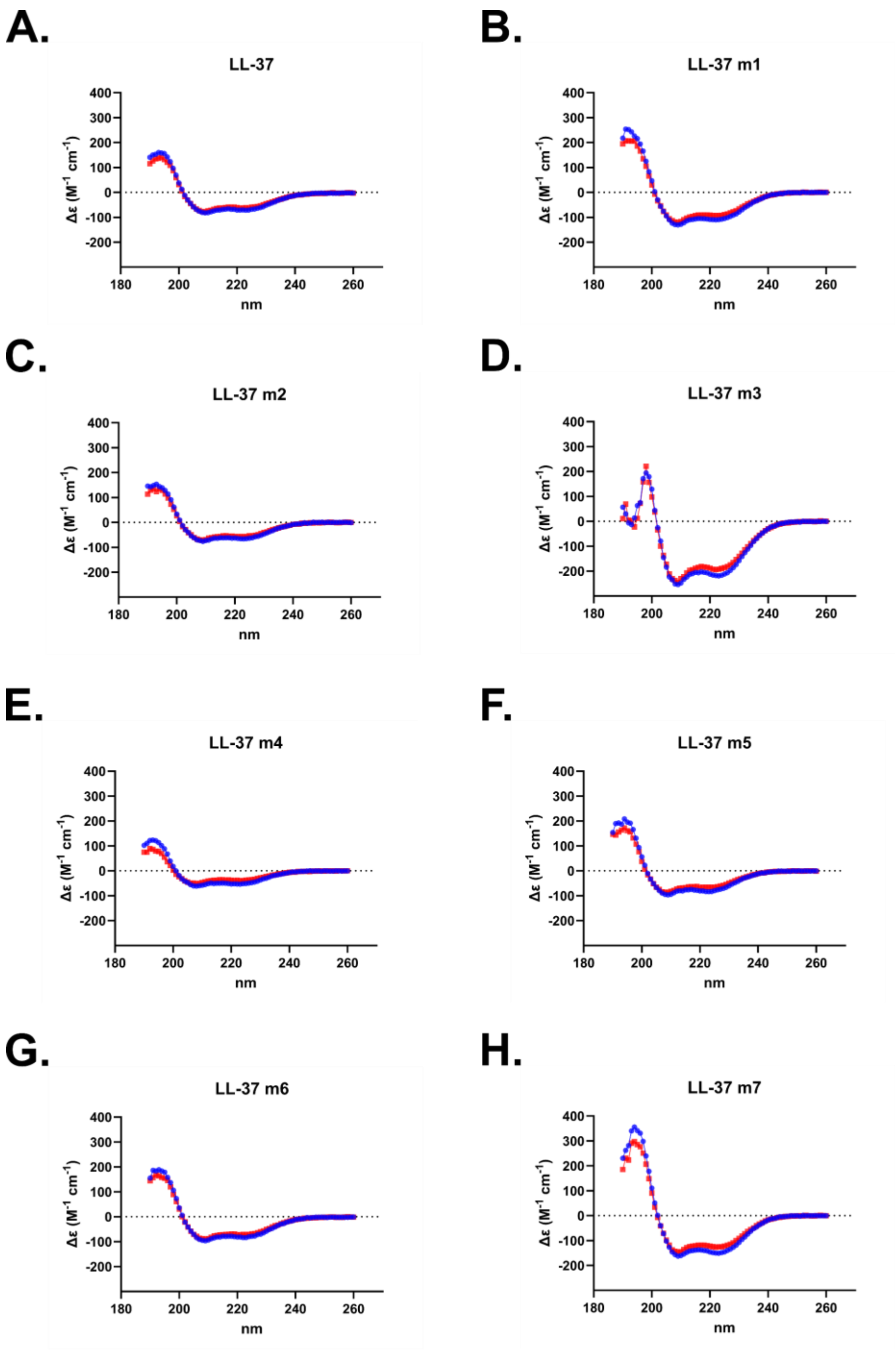


I.

| | % Helix | |
|---|---|---|
| | 20 °C | 37 °C |
| LL-37 | 100 | 100 |
| LL-37 m1 | 100 | 100 |
| LL-37 m2 | 100 | 100 |
| LL-37 m3 | 100 | 100 |
| LL-37 m4 | 100 | 100 |
| LL-37 m5 | 100 | 100 |
| LL-37 m6 | 100 | 100 |
| LL-37 m7 | 100 | 100 |

**Supplementary Figure 1 – Circular dichroism (CD) spectroscopy of Figure 1 mutants. A-H.** CD spectra are displayed for each mutant dissolved in water plus 20% trifluoroethanol, with spectra covering 190-260 nm at both 20 °C (Blue) and 37 °C (Red). **I.** Estimation of helical content as calculated with BeStSel for each mutant.

**A.**

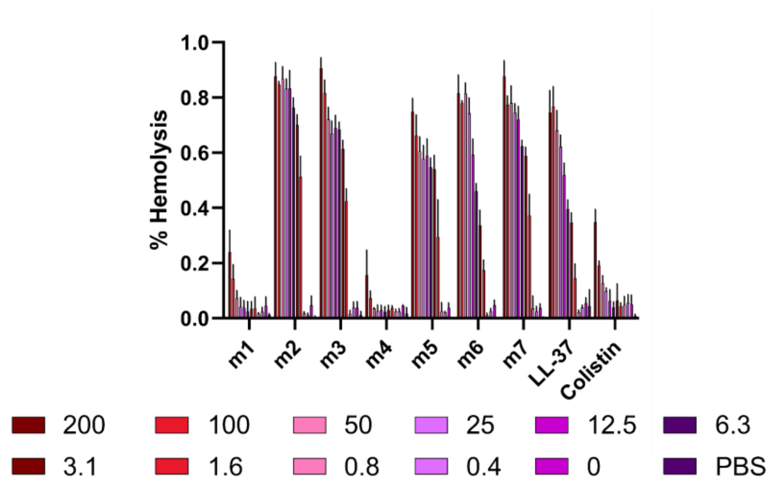
% Hemolysis
1.0
0.8
0.6
0.4
0.2
0.0
m1
m2
m3
m4
m5
m6
m7
LL-37
Colistin
200
100
50
25
12.5
6.3
3.1
1.6
0.8
0.4
0
PBS

**B.**

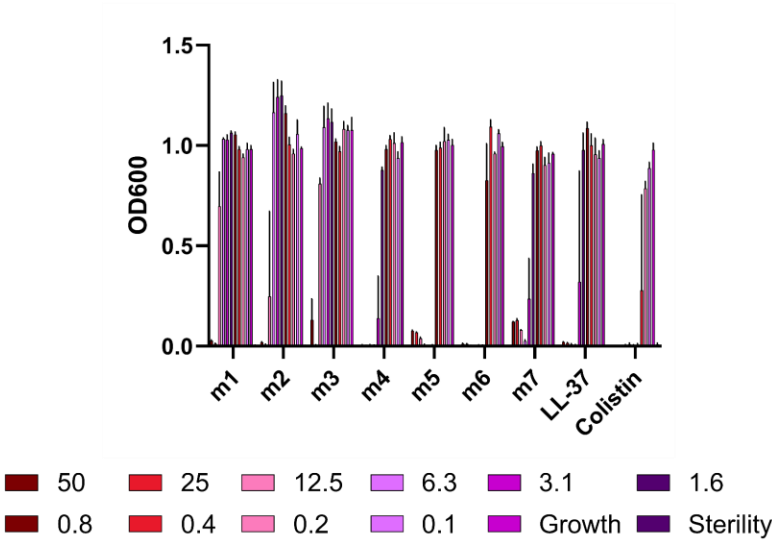
OD600
1.5
1.0
0.5
0.0
m1
m2
m3
m4
m5
m6
m7
LL-37
Colistin
50
25
12.5
6.3
3.1
1.6
0.8
0.4
0.2
0.1
Growth
Sterility

**C.**

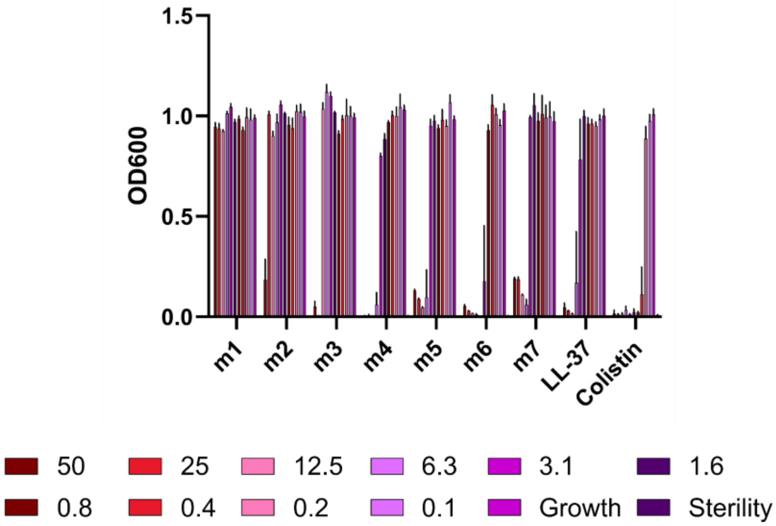
OD600
1.5
1.0
0.5
0.0
m1
m2
m3
m4
m5
m6
m7
LL-37
Colistin
50
25
12.5
6.3
3.1
1.6
0.8
0.4
0.2
0.1
Growth
Sterility

**Supplementary Figure 2 – Histogram depiction of Figure 1 data. A.** Endpoint hemolytic activity of each peptide against defibrinated sheep blood relative to a 100% hemolysis thrice freeze-thawed control after 18 hours at 37 °C. Raw data were obtained by absorbance at 414 nm in well supernatants for free hemoglobin. **B-C.** Endpoint activity of each peptide against *E. coli* (**B**) or *P. aeruginosa* (**C**) after 18 hours of incubation at 37 °C in the presence of the indicated concentrations of each peptide. Data are the raw OD600 after 18 hours at 37 °C for three independent experiments. Peptide concentrations are given in μM.

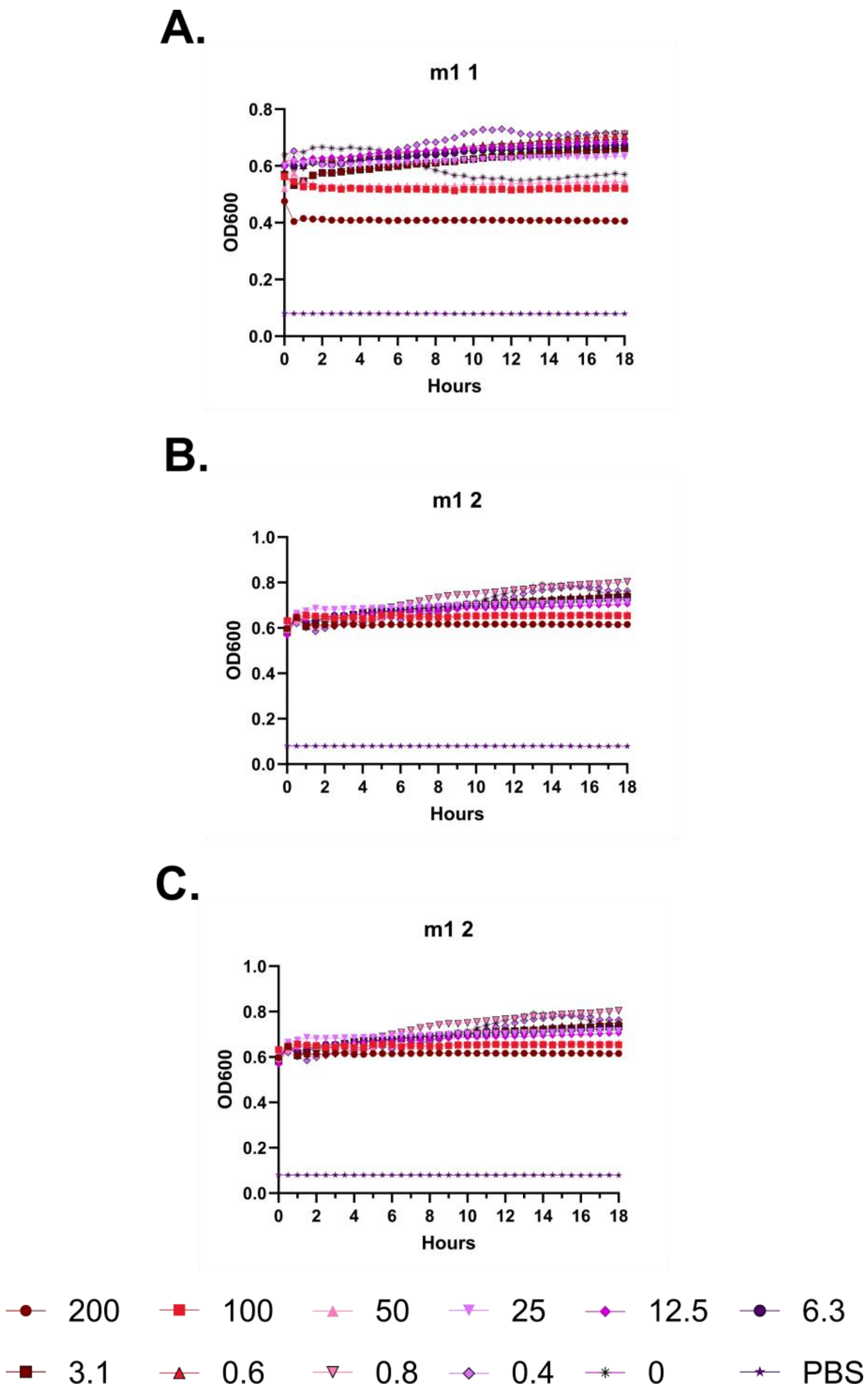
A.
m1 1
OD600
Hours
B.
m1 2
OD600
Hours
C.
m1 2
OD600
Hours
200
100
50
25
12.5
6.3
3.1
0.6
0.8
0.4
0
PBS

**Supplementary Figure 3 – Kinetic hemolysis assays for LL-37 m1. A.** Hemolysis was monitored by OD600 over 18 hours at 37 °C in the presence of the indicated peptide concentrations. Decreasing OD600 indicates increasing hemolysis. Peptide concentrations are given in μM. Each curve indicates an independent experiment.

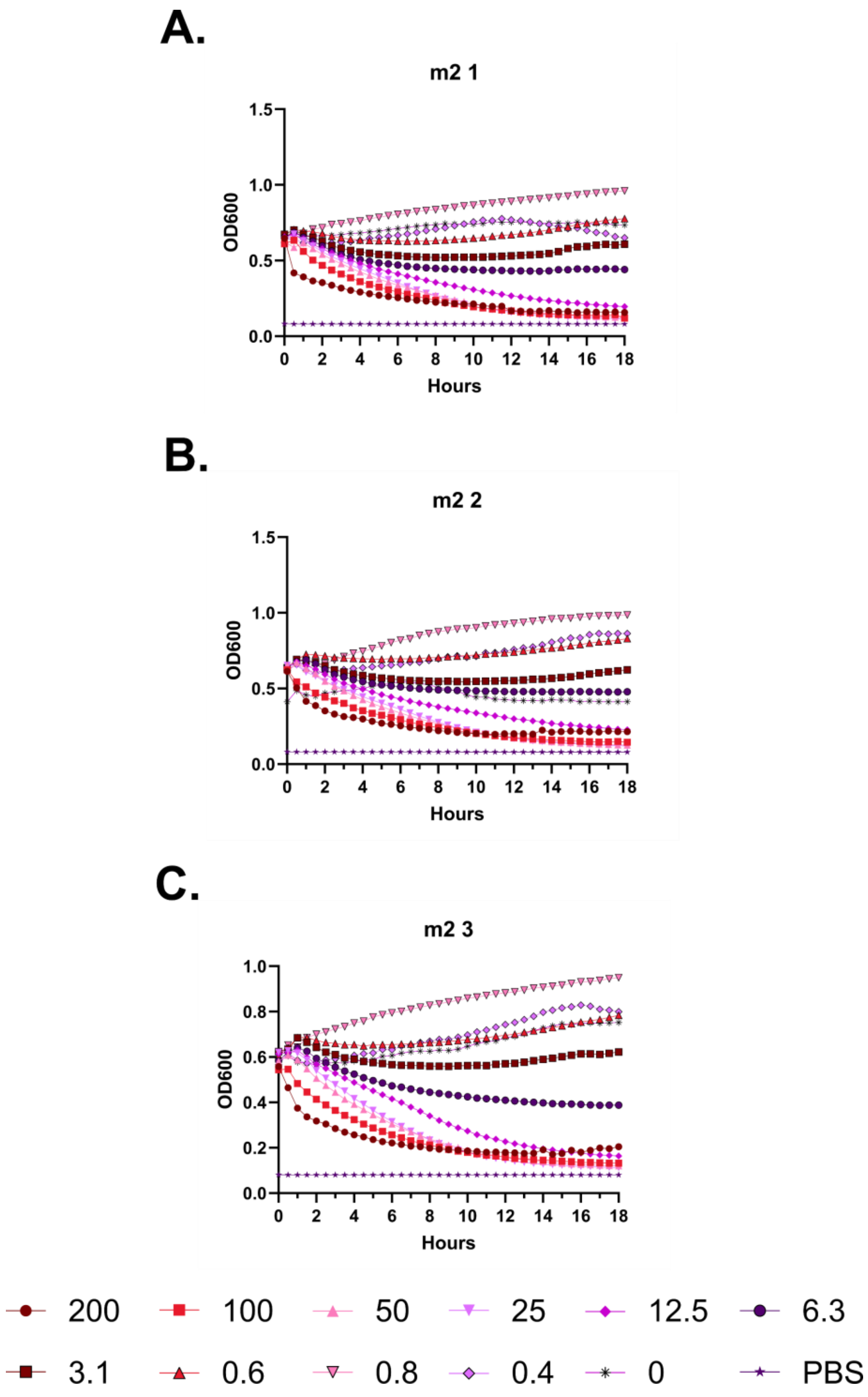
A.
m2 1
OD600
Hours
B.
m2 2
OD600
Hours
C.
m2 3
OD600
Hours
200
100
50
25
12.5
6.3
3.1
0.6
0.8
0.4
0
PBS

**Supplementary Figure 4 – Kinetic hemolysis assays for LL-37 m2. A-C.** Hemolysis was monitored by OD600 over 18 hours at 37 °C in the presence of the indicated peptide concentrations. Decreasing OD600 indicates increasing hemolysis. Peptide concentrations are given in μM. Each curve indicates an independent experiment.

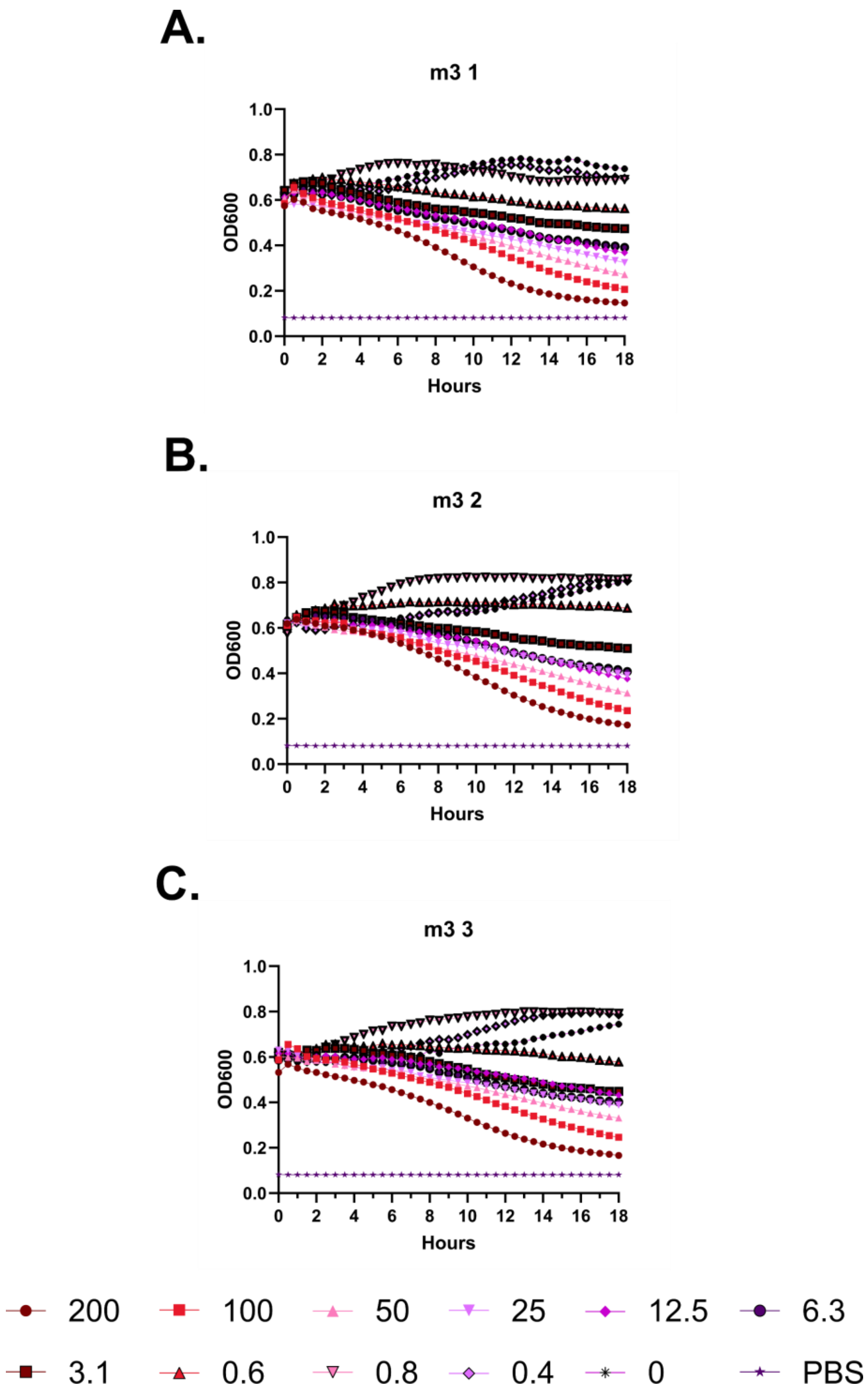
A.
m3 1
OD600
1.0
0.8
0.6
0.4
0.2
0.0
0 2 4 6 8 10 12 14 16 18
Hours
B.
m3 2
OD600
1.0
0.8
0.6
0.4
0.2
0.0
0 2 4 6 8 10 12 14 16 18
Hours
C.
m3 3
OD600
1.0
0.8
0.6
0.4
0.2
0.0
0 2 4 6 8 10 12 14 16 18
Hours
200
100
50
25
12.5
6.3
3.1
0.6
0.8
0.4
0
PBS

**Supplementary Figure 5 – Kinetic hemolysis assays for LL-37 m3. A-C.** Hemolysis was monitored by OD600 over 18 hours at 37 °C in the presence of the indicated peptide concentrations. Decreasing OD600 indicates increasing hemolysis. Peptide concentrations are given in μM. Each curve indicates an independent experiment.

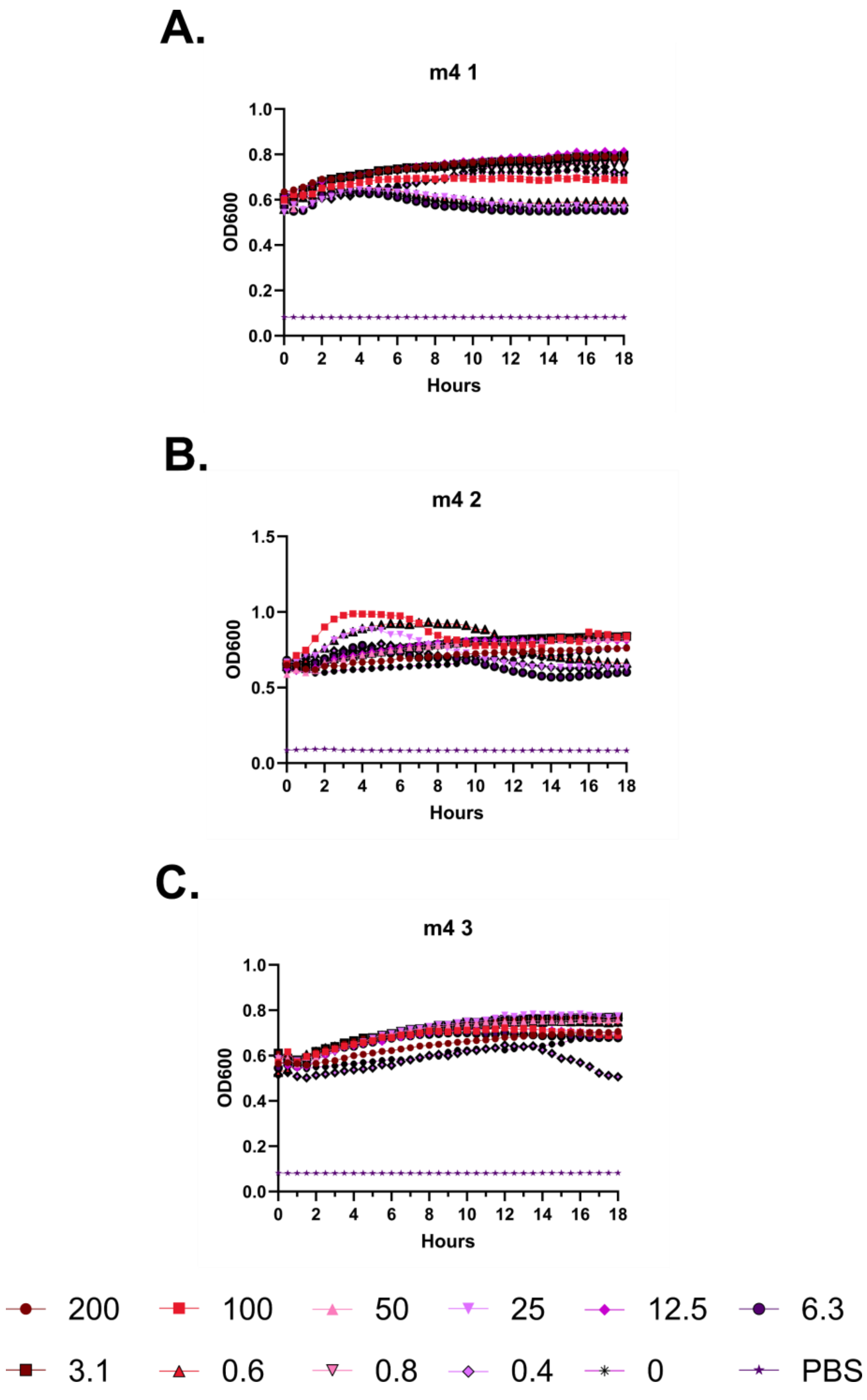
A.
m4 1
OD600
Hours
B.
m4 2
OD600
Hours
C.
m4 3
OD600
Hours
200
100
50
25
12.5
6.3
3.1
0.6
0.8
0.4
0
PBS

**Supplementary Figure 6 – Kinetic hemolysis assays for LL-37 m4. A-C.** Hemolysis was monitored by OD600 over 18 hours at 37 °C in the presence of the indicated peptide concentrations. Decreasing OD600 indicates increasing hemolysis. Peptide concentrations are given in μM. Each curve indicates an independent experiment.

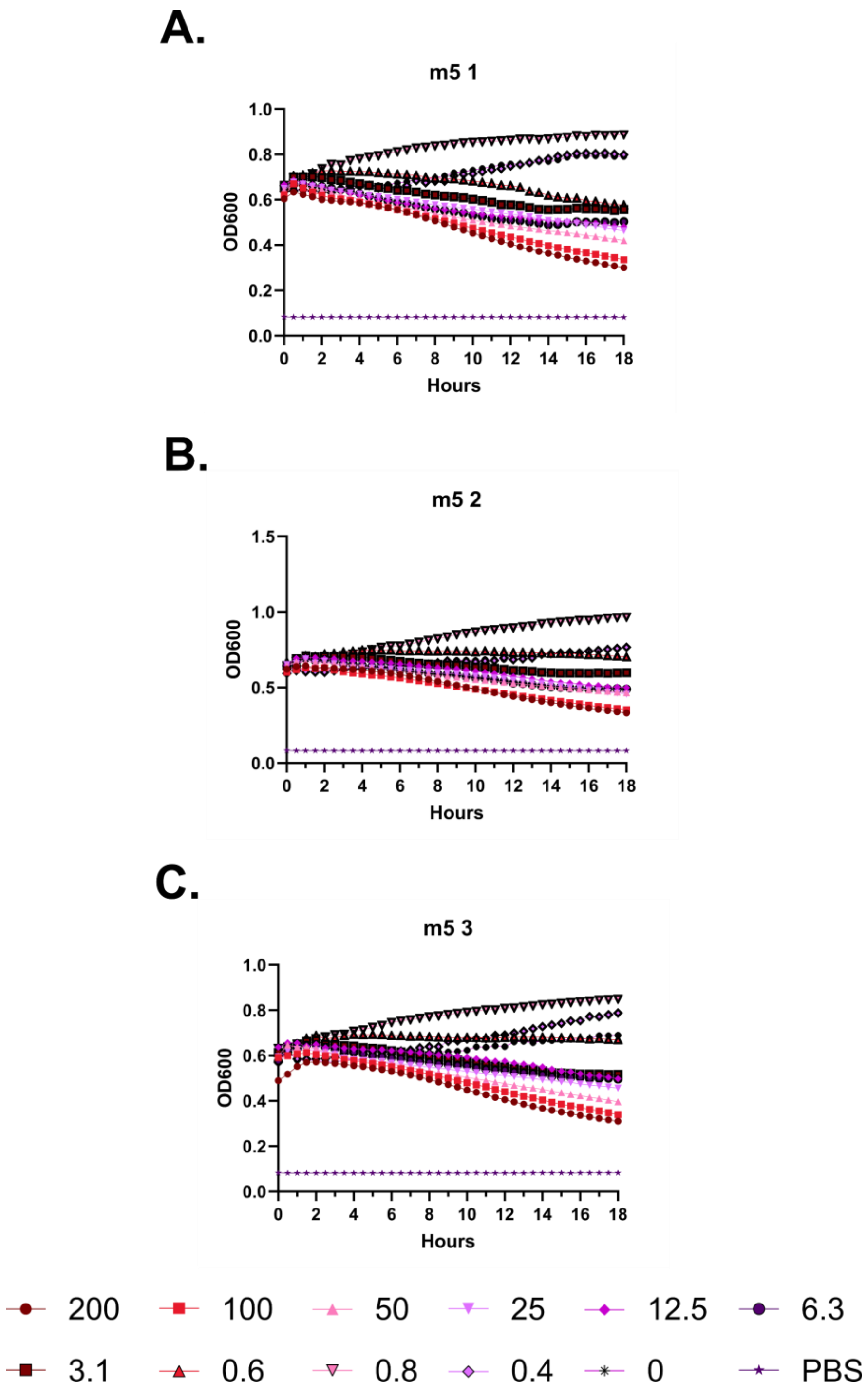
A.
m5 1
OD600
Hours
B.
m5 2
OD600
Hours
C.
m5 3
OD600
Hours
200
100
50
25
12.5
6.3
3.1
0.6
0.8
0.4
0
PBS

**Supplementary Figure 7 – Kinetic hemolysis assays for LL-37 m5. A-C.** Hemolysis was monitored by OD600 over 18 hours at 37 °C in the presence of the indicated peptide concentrations. Decreasing OD600 indicates increasing hemolysis. Peptide concentrations are given in μM. Each curve indicates an independent experiment.

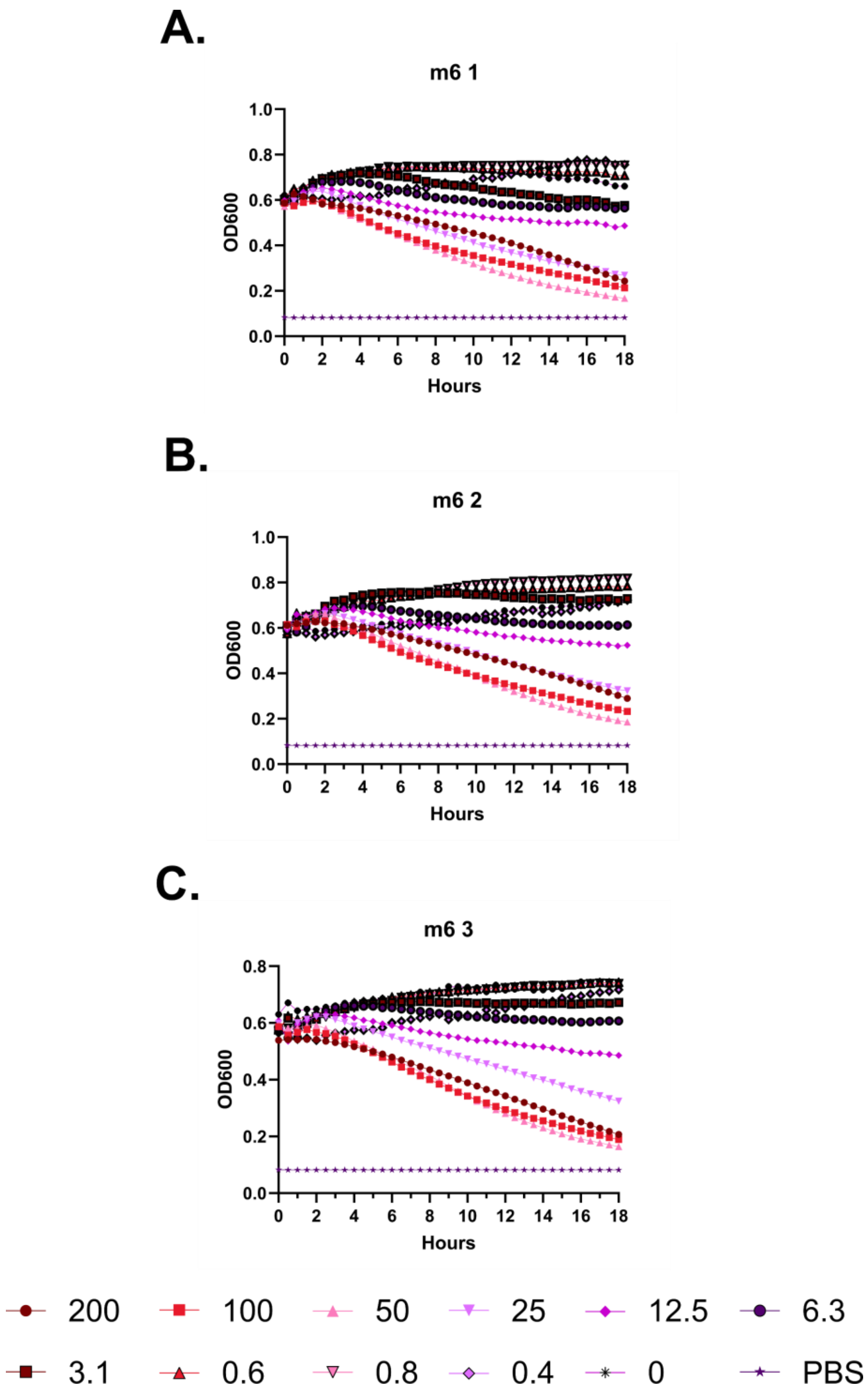
A.
m6 1
1.0
0.8
0.6
0.4
0.2
0.0
OD600
0
2
4
6
8
10
12
14
16
18
Hours
B.
m6 2
1.0
0.8
0.6
0.4
0.2
0.0
OD600
0
2
4
6
8
10
12
14
16
18
Hours
C.
m6 3
0.8
0.6
0.4
0.2
0.0
OD600
0
2
4
6
8
10
12
14
16
18
Hours
200
100
50
25
12.5
6.3
3.1
0.6
0.8
0.4
0
PBS

**Supplementary Figure 8 – Kinetic hemolysis assays for LL-37 m6. A-C.** Hemolysis was monitored by OD600 over 18 hours at 37 °C in the presence of the indicated peptide concentrations. Decreasing OD600 indicates increasing hemolysis. Peptide concentrations are given in µM. Each curve indicates an independent experiment.

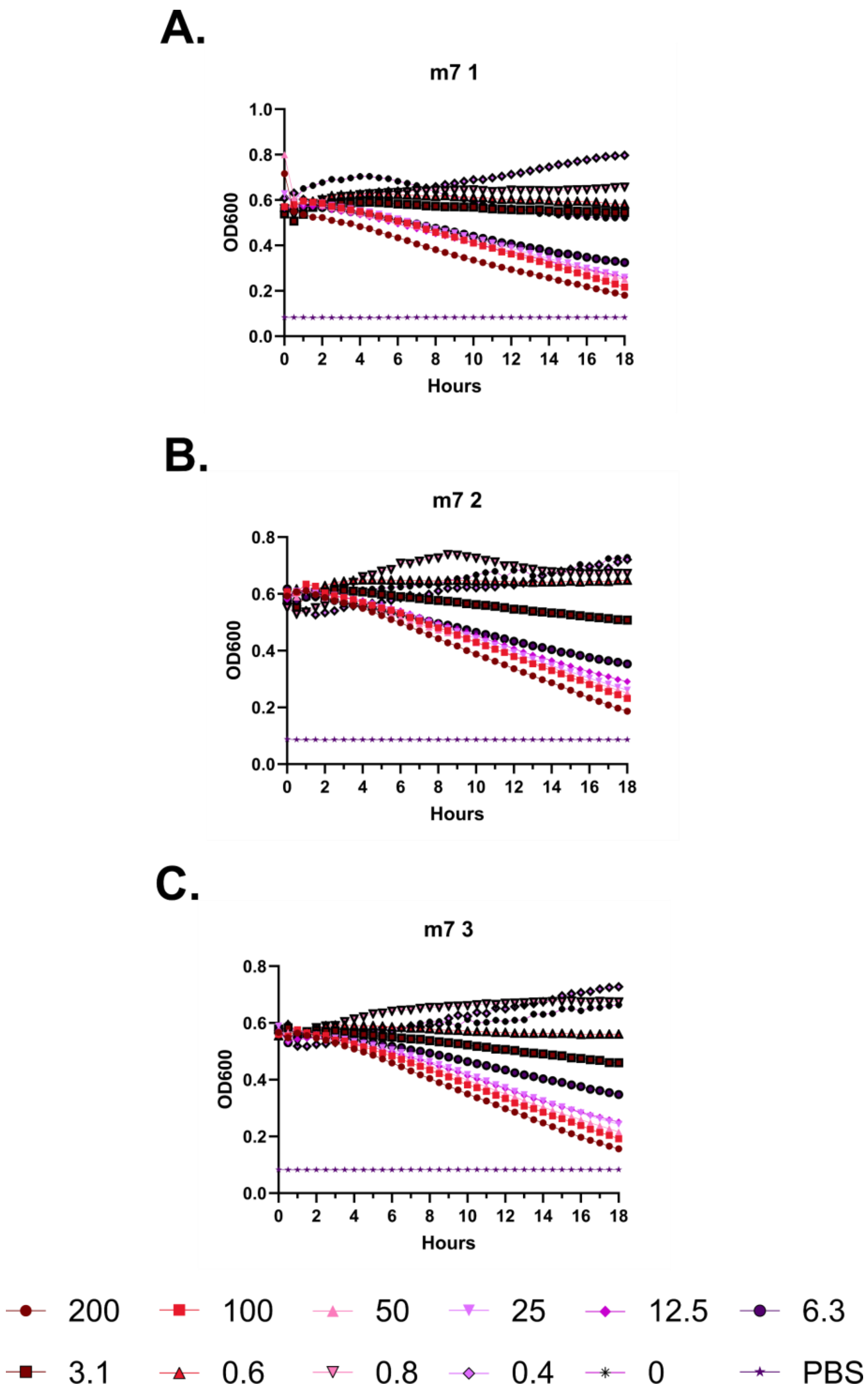

A.
m7 1
OD600
1.0
0.8
0.6
0.4
0.2
0.0
0
2
4
6
8
10
12
14
16
18
Hours
B.
m7 2
OD600
0.8
0.6
0.4
0.2
0.0
0
2
4
6
8
10
12
14
16
18
Hours
C.
m7 3
OD600
0.8
0.6
0.4
0.2
0.0
0
2
4
6
8
10
12
14
16
18
Hours
200
100
50
25
12.5
6.3
3.1
0.6
0.8
0.4
0
PBS

**Supplementary Figure 9 – Kinetic hemolysis assays for LL-37 m7. A-C.** Hemolysis was monitored by OD600 over 18 hours at 37 °C in the presence of the indicated peptide concentrations. Decreasing OD600 indicates increasing hemolysis. Peptide concentrations are given in μM. Each curve indicates an independent experiment.

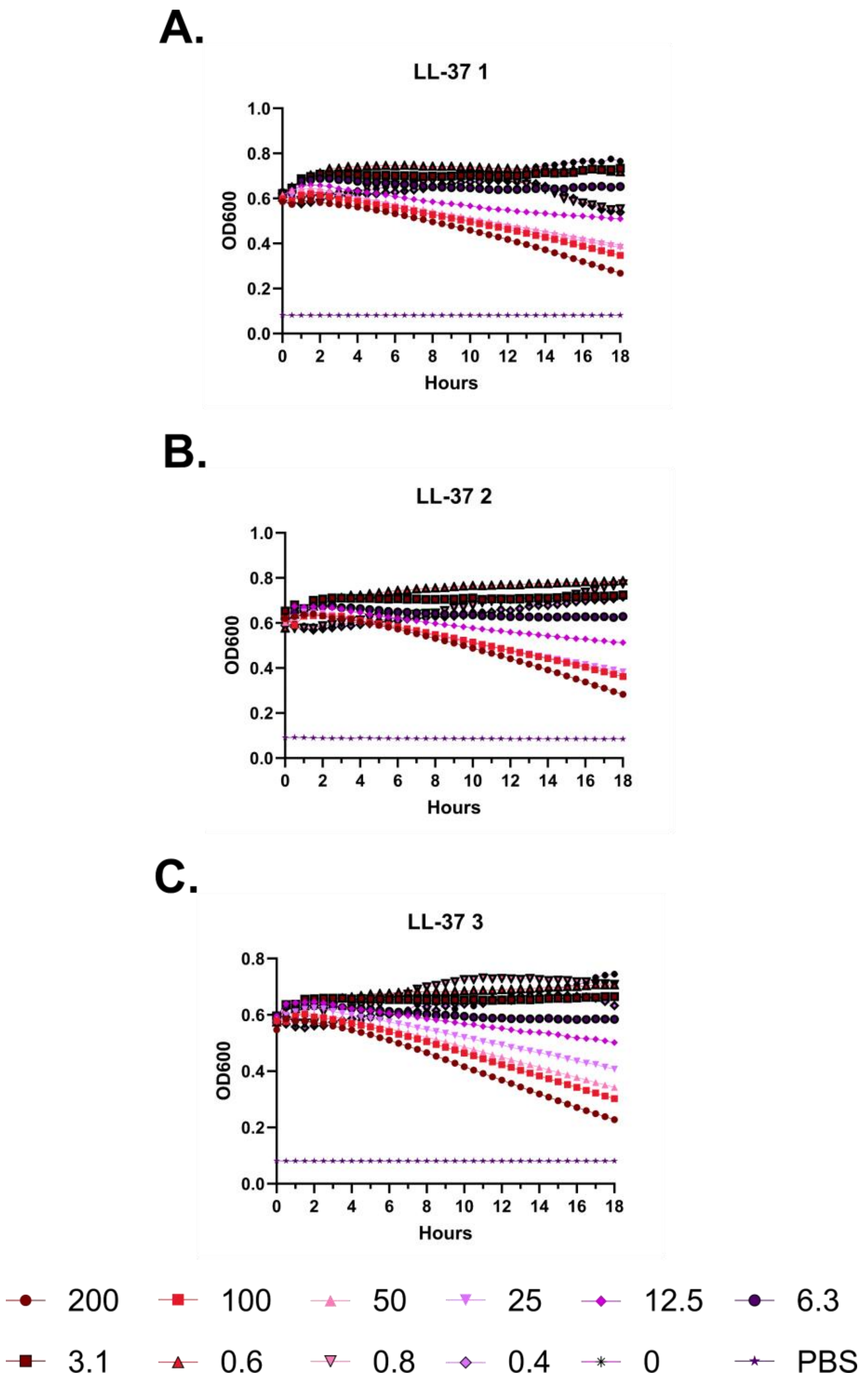
A.
LL-37 1
OD600
Hours
B.
LL-37 2
OD600
Hours
C.
LL-37 3
OD600
Hours
200
100
50
25
12.5
6.3
3.1
0.6
0.8
0.4
0
PBS

**Supplementary Figure 10 – Kinetic hemolysis assays for LL-37. A-C.** Hemolysis was monitored by OD600 over 18 hours at 37 °C in the presence of the indicated peptide concentrations. Decreasing OD600 indicates increasing hemolysis. Peptide concentrations are given in μM. Each curve indicates an independent experiment.

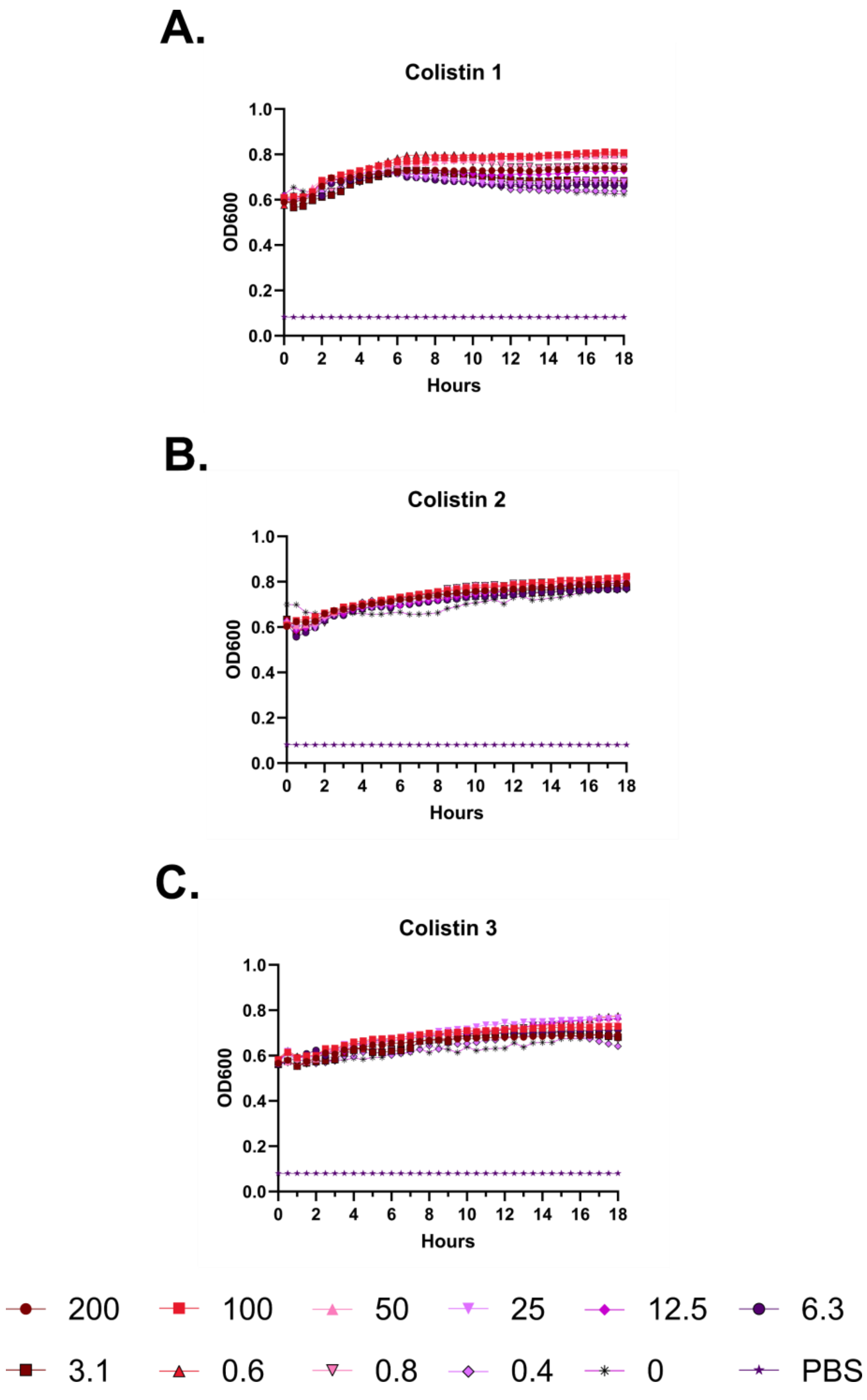
A.
Colistin 1
OD600
Hours
B.
Colistin 2
OD600
Hours
C.
Colistin 3
OD600
Hours
200
100
50
25
12.5
6.3
3.1
0.6
0.8
0.4
0
PBS

**Supplementary Figure 11 – Kinetic hemolysis assays for colistin. A-C.** Hemolysis was monitored by OD600 over 18 hours at 37 °C in the presence of the indicated peptide concentrations. Decreasing OD600 indicates increasing hemolysis. Peptide concentrations are given in μM. Each curve indicates an independent experiment.

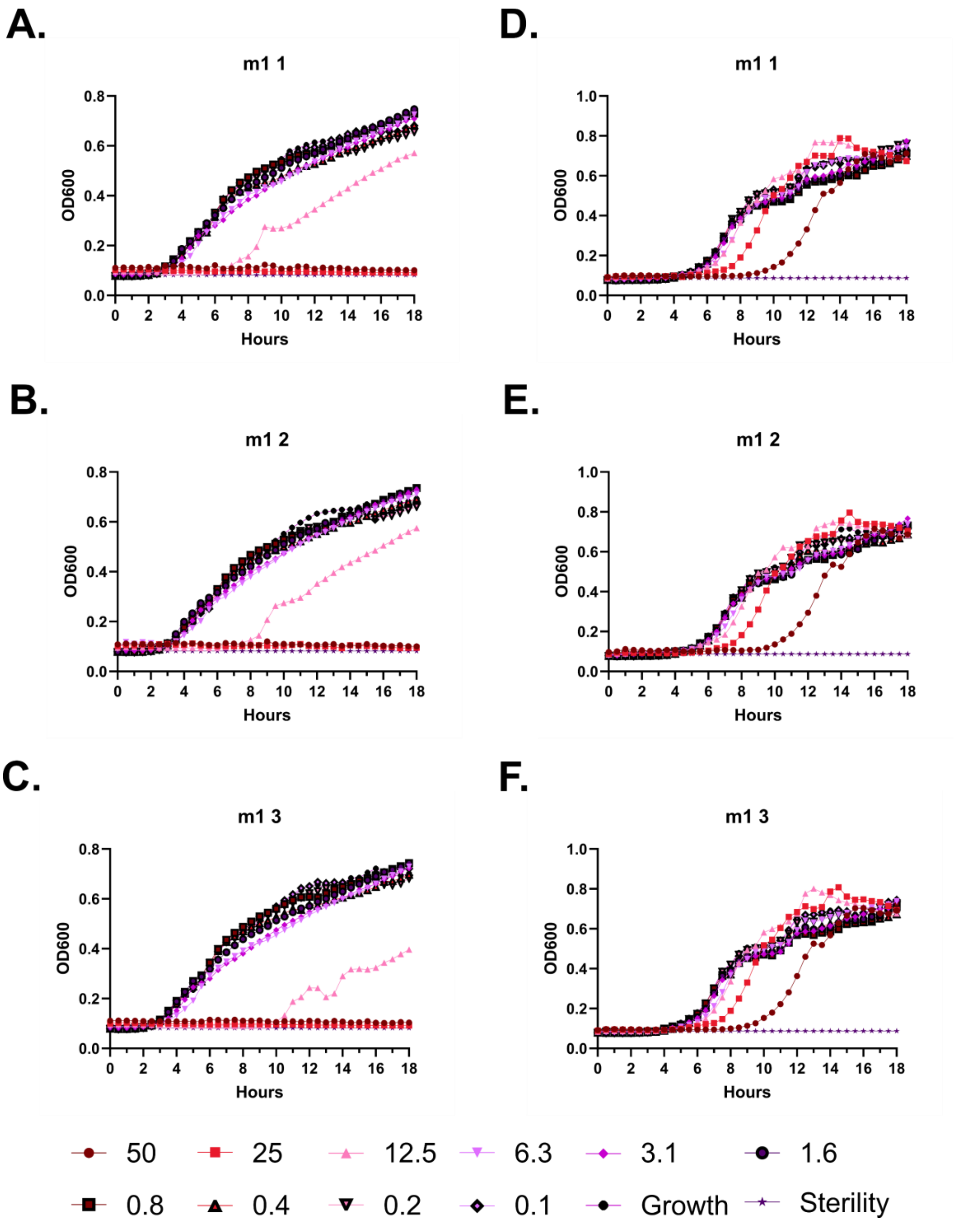
A.
m1 1
OD600
Hours
B.
m1 2
OD600
Hours
C.
m1 3
OD600
Hours
D.
m1 1
OD600
Hours
E.
m1 2
OD600
Hours
F.
m1 3
OD600
Hours
50
25
12.5
6.3
3.1
1.6
0.8
0.4
0.2
0.1
Growth
Sterility

**Supplementary Figure 12 – Kinetic antimicrobial susceptibility testing for LL-37 m1. A-C.** *E. coli*. **D-F.** *Pseudomonas.* Bacterial growth was monitored by OD600 over 18 hours at 37 °C in the presence of the indicated peptide concentrations. Increasing OD600 indicates increasing bacterial growth. Peptide concentrations are given in μM. Each curve indicates an independent experiment.

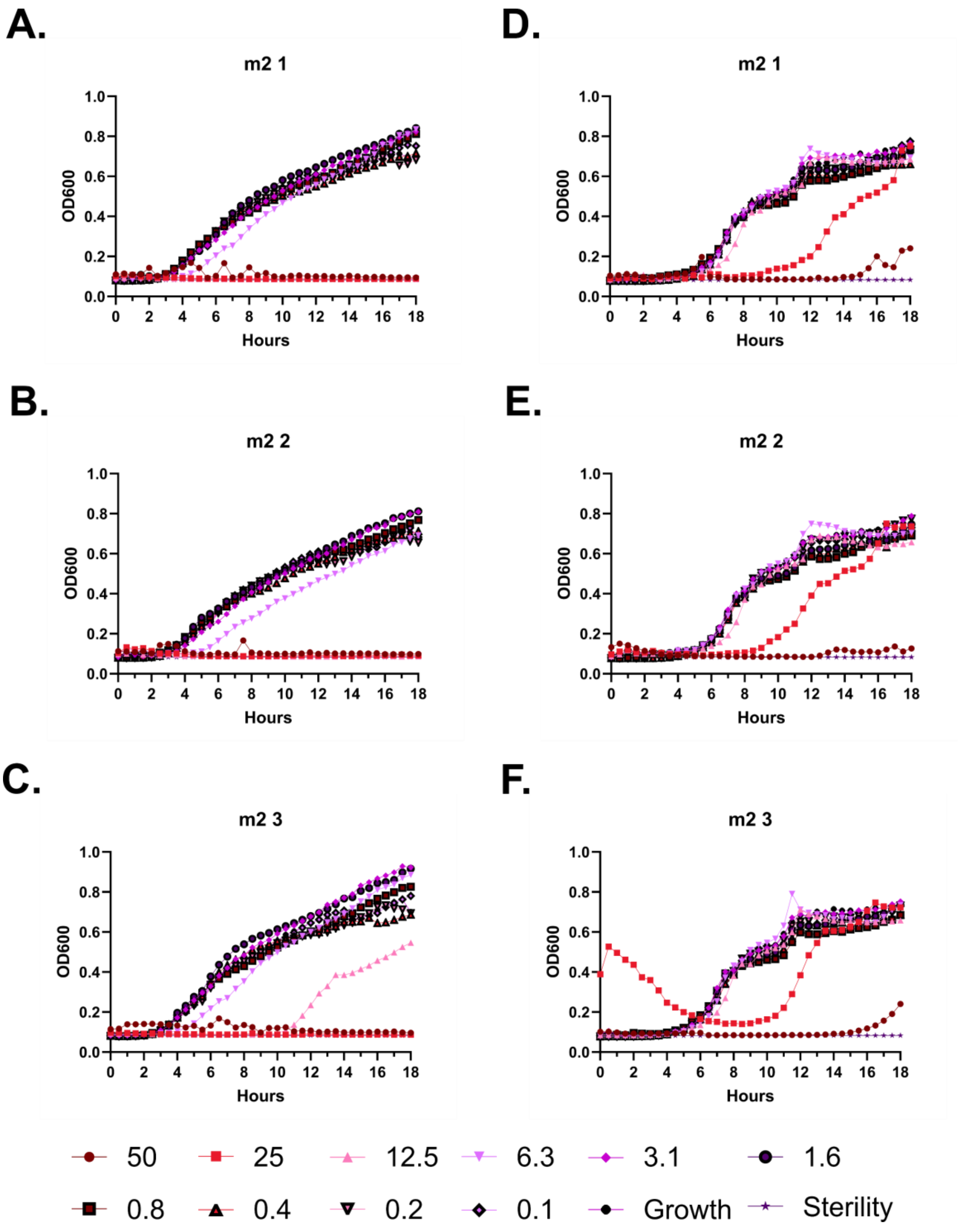
A.
m2 1
OD600
Hours
D.
m2 1
OD600
Hours
B.
m2 2
OD600
Hours
E.
m2 2
OD600
Hours
C.
m2 3
OD600
Hours
F.
m2 3
OD600
Hours
0.0
0.2
0.4
0.6
0.8
1.0
0 2 4 6 8 10 12 14 16 18
50
25
12.5
6.3
3.1
1.6
0.8
0.4
0.2
0.1
Growth
Sterility

**Supplementary Figure 13 – Kinetic antimicrobial susceptibility testing for LL-37 m2. A-C.** *E. coli*. **D-F.** *Pseudomonas.* Bacterial growth was monitored by OD600 over 18 hours at 37 °C in the presence of the indicated peptide concentrations. Increasing OD600 indicates increasing bacterial growth. Peptide concentrations are given in μM. Each curve indicates an independent experiment.

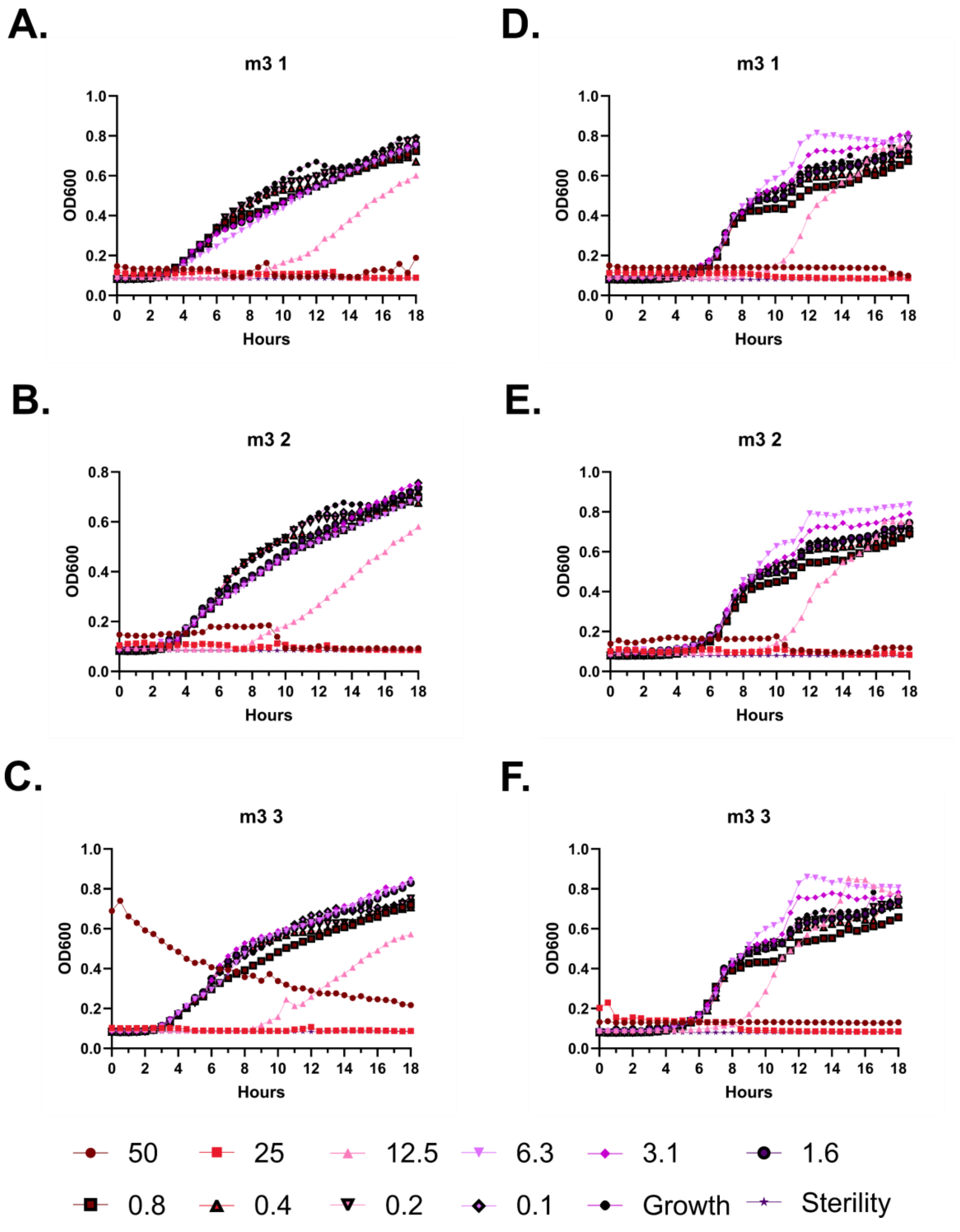
A.
m3 1
OD600
Hours
D.
m3 1
OD600
Hours
B.
m3 2
OD600
Hours
E.
m3 2
OD600
Hours
C.
m3 3
OD600
Hours
F.
m3 3
OD600
Hours
50
25
12.5
6.3
3.1
1.6
0.8
0.4
0.2
0.1
Growth
Sterility

**Supplementary Figure 14 – Kinetic antimicrobial susceptibility testing for LL-37 m3. A-C.** *E. coli*. **D-F.** *Pseudomonas.* Bacterial growth was monitored by OD600 over 18 hours at 37 °C in the presence of the indicated peptide concentrations. Increasing OD600 indicates increasing bacterial growth. Peptide concentrations are given in μM. Each curve indicates an independent experiment.

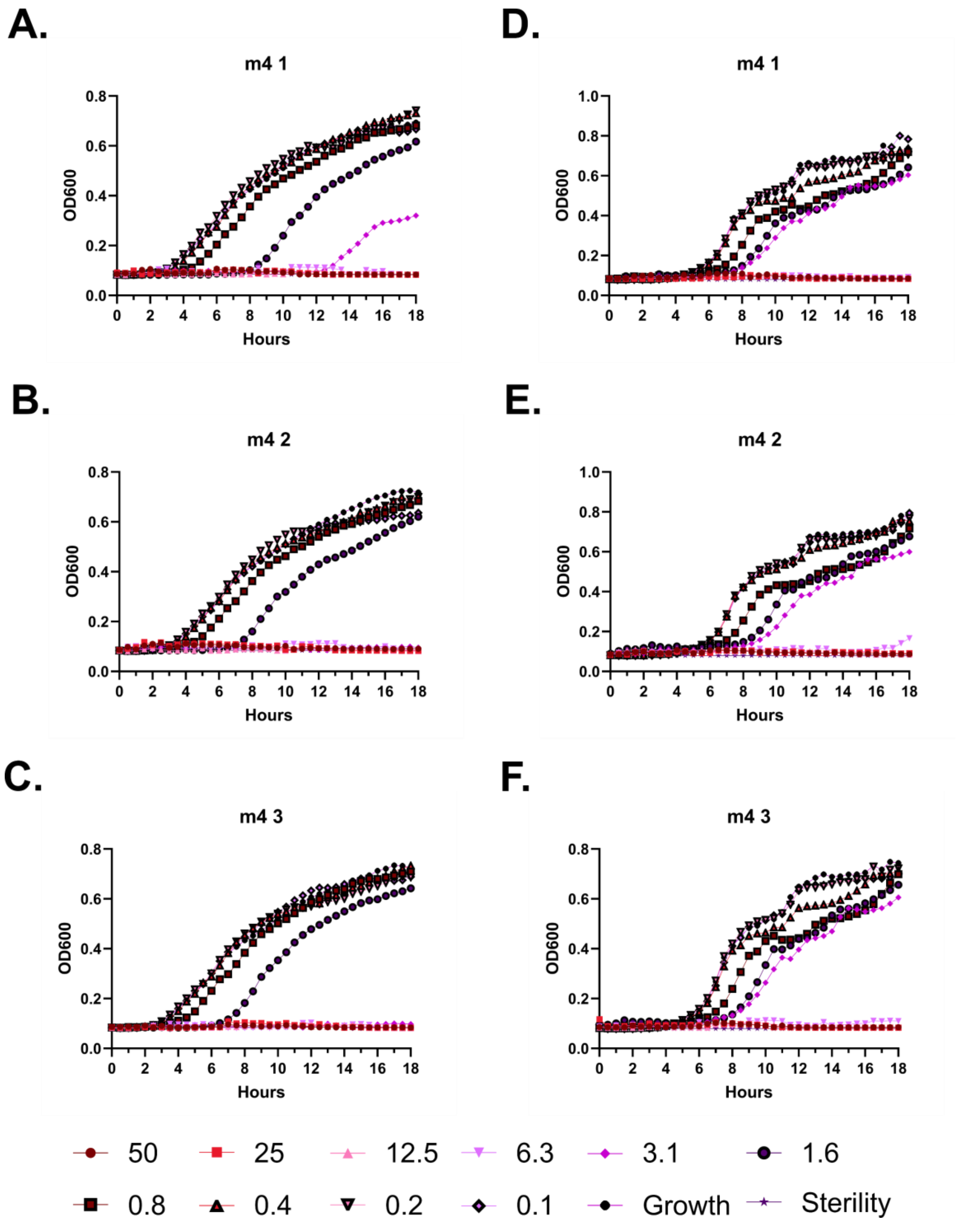
A.
m4 1
OD600
Hours
D.
m4 1
B.
m4 2
E.
m4 2
C.
m4 3
F.
m4 3
50
25
12.5
6.3
3.1
1.6
0.8
0.4
0.2
0.1
Growth
Sterility

**Supplementary Figure 15 – Kinetic antimicrobial susceptibility testing for LL-37 m4. A-C.** *E. coli*. **D-F.** *Pseudomonas.* Bacterial growth was monitored by OD600 over 18 hours at 37 °C in the presence of the indicated peptide concentrations. Increasing OD600 indicates increasing bacterial growth. Peptide concentrations are given in μM. Each curve indicates an independent experiment.

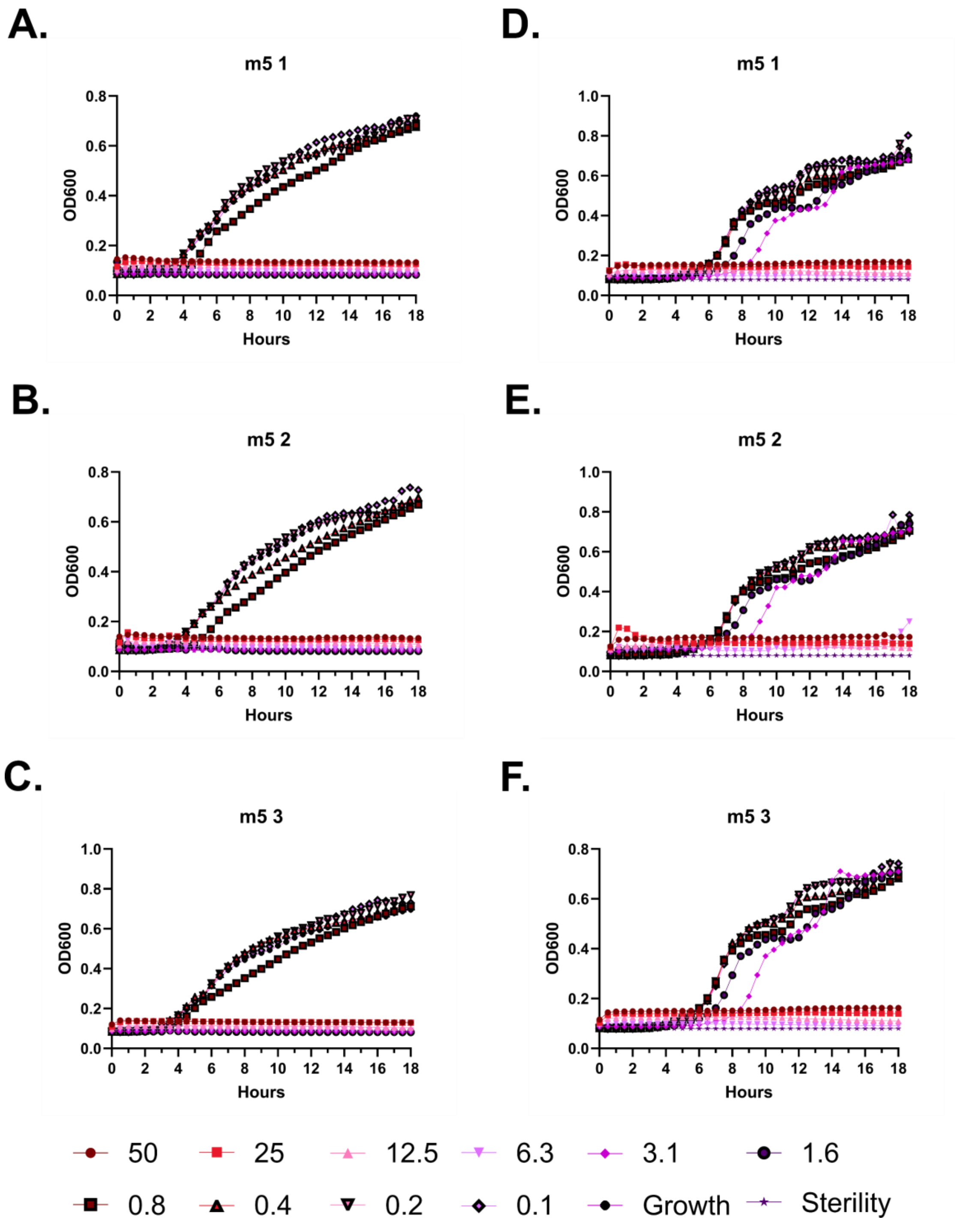
A.
m5 1
OD600
Hours
B.
m5 2
C.
m5 3
D.
m5 1
E.
m5 2
F.
m5 3
50
25
12.5
6.3
3.1
1.6
0.8
0.4
0.2
0.1
Growth
Sterility

**Supplementary Figure 16 – Kinetic antimicrobial susceptibility testing for LL-37 m5. A-C.** *E. coli*. **D-F.** *Pseudomonas.* Bacterial growth was monitored by OD600 over 18 hours at 37 °C in the presence of the indicated peptide concentrations. Increasing OD600 indicates increasing bacterial growth. Peptide concentrations are given in μM. Each curve indicates an independent experiment.

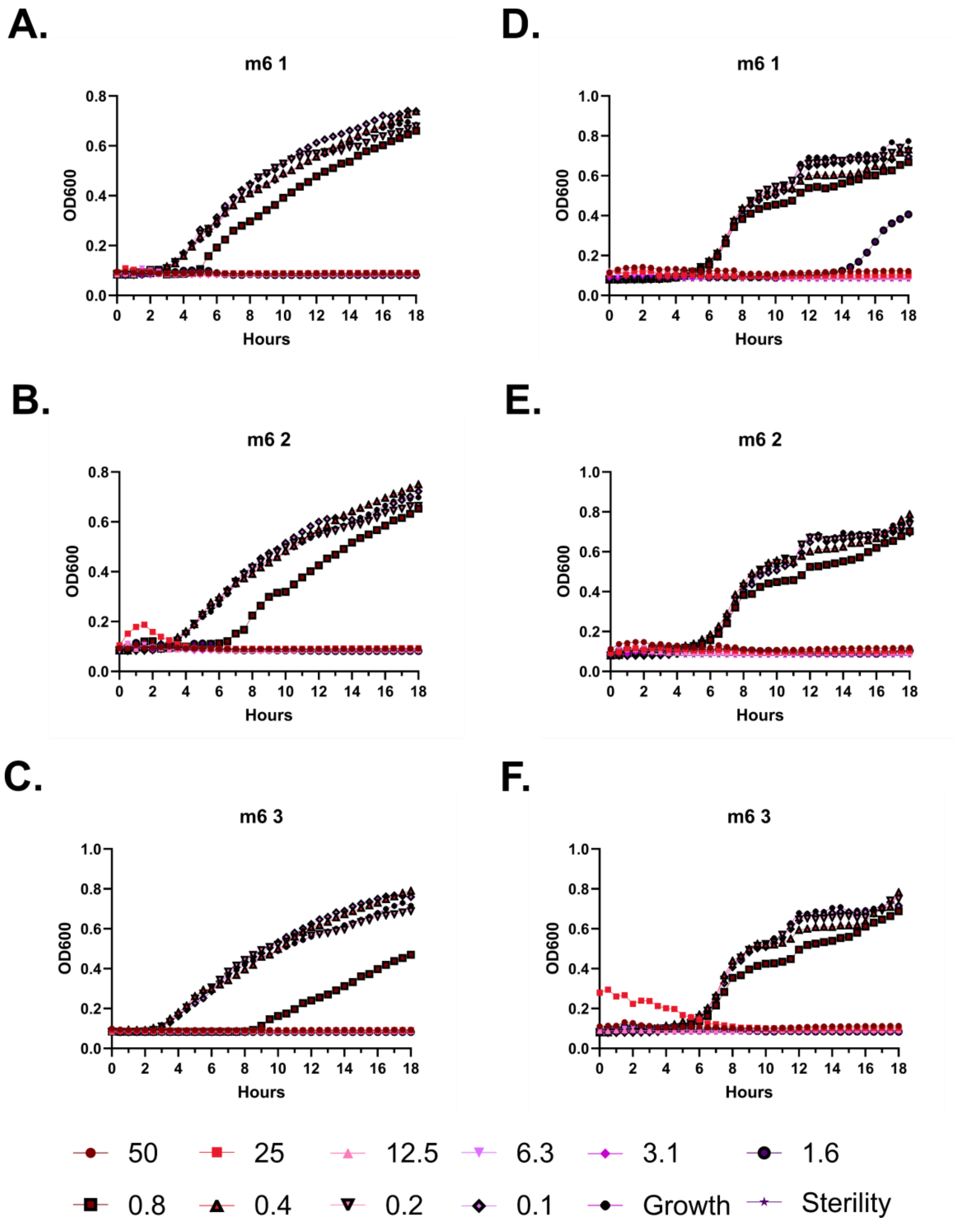
A.
m6 1
OD600
Hours
D.
m6 1
OD600
Hours
B.
m6 2
OD600
Hours
E.
m6 2
OD600
Hours
C.
m6 3
OD600
Hours
F.
m6 3
OD600
Hours
50
25
12.5
6.3
3.1
1.6
0.8
0.4
0.2
0.1
Growth
Sterility

**Supplementary Figure 17 – Kinetic antimicrobial susceptibility testing for LL-37 m6. A-C.** *E. coli*. **D-F.** *Pseudomonas.* Bacterial growth was monitored by OD600 over 18 hours at 37 °C in the presence of the indicated peptide concentrations. Increasing OD600 indicates increasing bacterial growth. Peptide concentrations are given in μM. Each curve indicates an independent experiment.

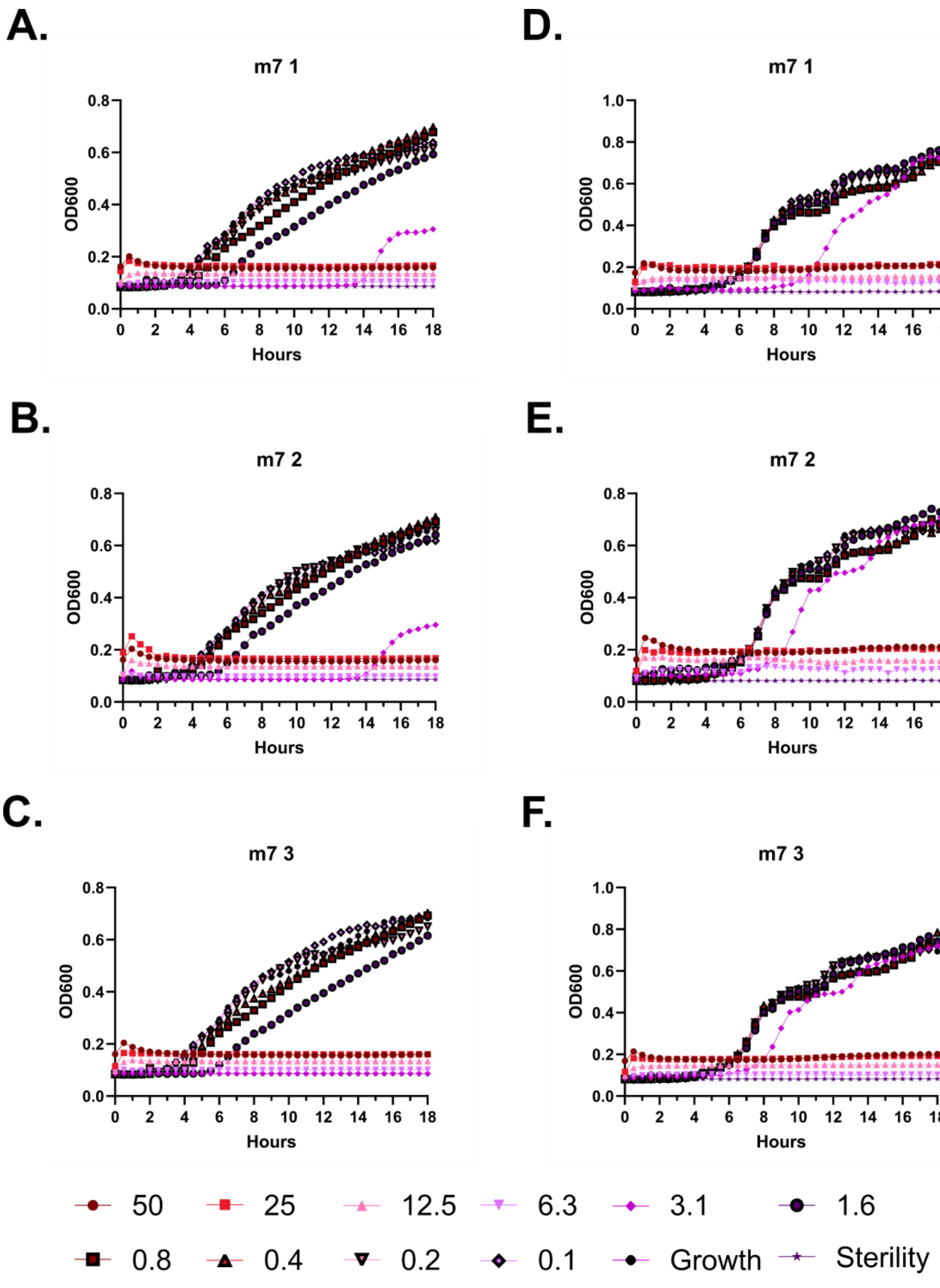
A.
m7 1
OD600
Hours
D.
m7 1
OD600
Hours
B.
m7 2
OD600
Hours
E.
m7 2
OD600
Hours
C.
m7 3
OD600
Hours
F.
m7 3
OD600
Hours
50
25
12.5
6.3
3.1
1.6
0.8
0.4
0.2
0.1
Growth
Sterility

**Supplementary Figure 18 – Kinetic antimicrobial susceptibility testing for LL-37 m7. A-C.** *E. coli*. **D-F.** *Pseudomonas.* Bacterial growth was monitored by OD600 over 18 hours at 37 °C in the presence of the indicated peptide concentrations. Increasing OD600 indicates increasing bacterial growth. Peptide concentrations are given in μM. Each curve indicates an independent experiment.

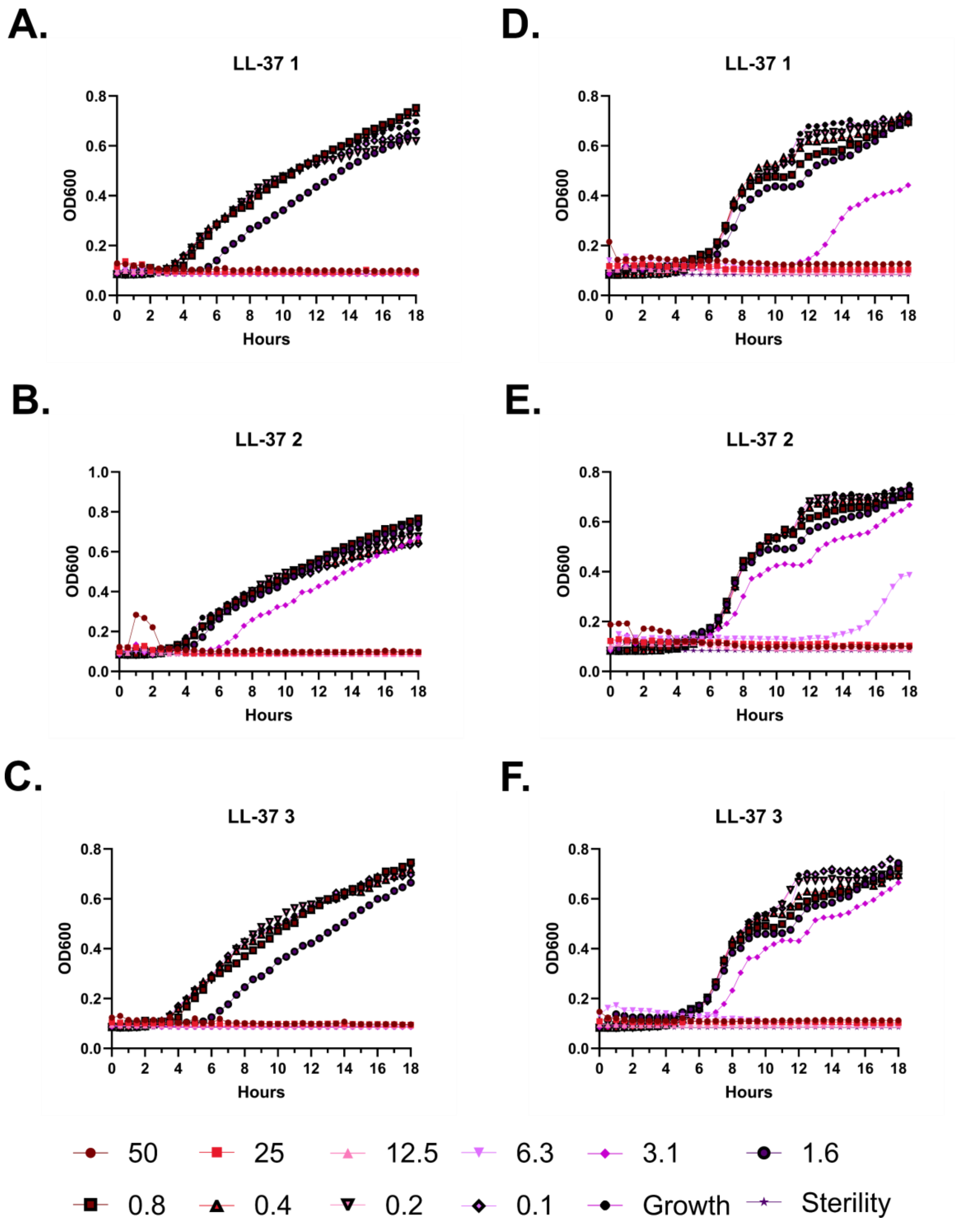
A.
LL-37 1
OD600
Hours
D.
LL-37 1
B.
LL-37 2
E.
LL-37 2
C.
LL-37 3
F.
LL-37 3
50
25
12.5
6.3
3.1
1.6
0.8
0.4
0.2
0.1
Growth
Sterility

**Supplementary Figure 19 – Kinetic antimicrobial susceptibility testing for LL-37. A-C.** *E. coli.* **D-F.** *Pseudomonas.* Bacterial growth was monitored by OD600 over 18 hours at 37 °C in the presence of the indicated peptide concentrations. Increasing OD600 indicates increasing bacterial growth. Peptide concentrations are given in μM. Each curve indicates an independent experiment.

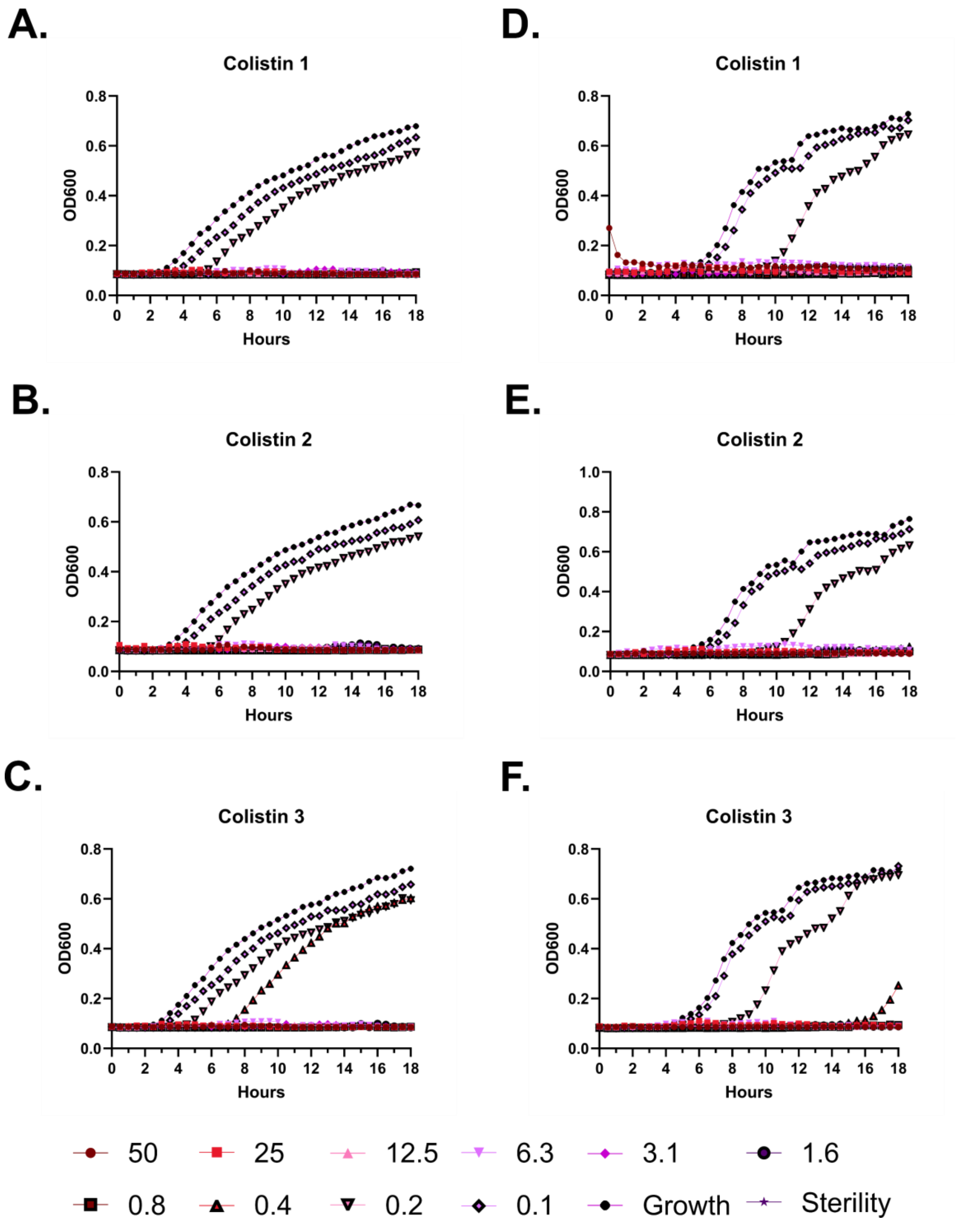
A.
Colistin 1
OD600
Hours
B.
Colistin 2
OD600
Hours
C.
Colistin 3
OD600
Hours
D.
Colistin 1
OD600
Hours
E.
Colistin 2
OD600
Hours
F.
Colistin 3
OD600
Hours
50
25
12.5
6.3
3.1
1.6
0.8
0.4
0.2
0.1
Growth
Sterility

**Supplementary Figure 20 – Kinetic antimicrobial susceptibility testing for colistin. A-C.** *E. coli*. **D-F.** *Pseudomonas.* Bacterial growth was monitored by OD600 over 18 hours at 37 °C in the presence of the indicated peptide concentrations. Increasing OD600 indicates increasing bacterial growth. Peptide concentrations are given in μM. Each curve indicates an independent experiment.

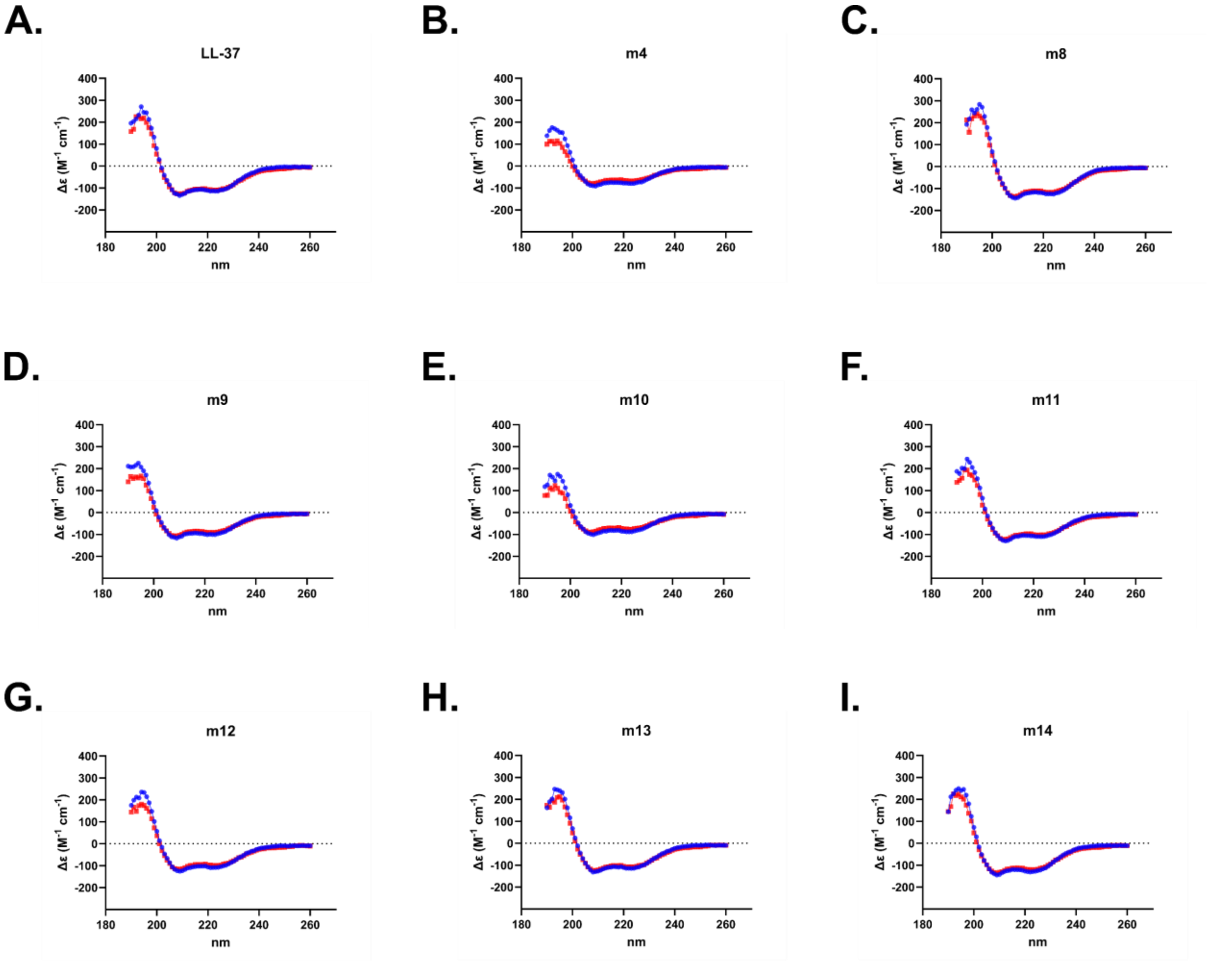


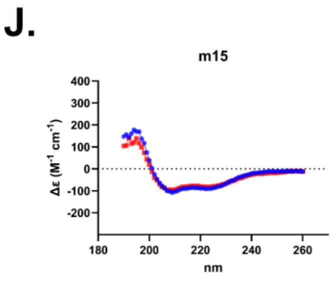


K.

| | % Helix | |
|---|---|---|
| | 20 °C | 37 °C |
| LL-37 | 100 | 100 |
| LL-37 m4 | 100 | 100 |
| LL-37 m8 | 100 | 100 |
| LL-37 m9 | 100 | 100 |
| LL-37 m10 | 100 | 100 |
| LL-37 m11 | 100 | 100 |
| LL-37 m12 | 100 | 100 |
| LL-37 m13 | 100 | 100 |
| LL-37 m14 | 100 | 100 |
| LL-37 m15 | 100 | 100 |

**Supplementary Figure 21 – Circular dichroism (CD) spectroscopy of Figure 2 mutants. A-J.** CD spectra are displayed for each mutant dissolved in water plus 20% trifluoroethanol. Data were collected from 190-260 nm at both 20 °C (Blue) and 37 °C (Red). **K.** Estimation of helical content as calculated with BeStSel for each mutant.

**A.**

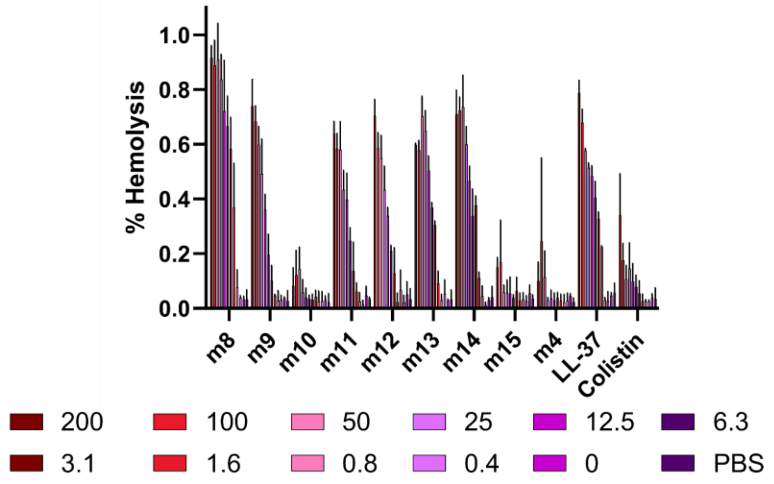
% Hemolysis
1.0
0.8
0.6
0.4
0.2
0.0
m8
m9
m10
m11
m12
m13
m14
m15
m4
LL-37
Colistin
200
100
50
25
12.5
6.3
3.1
1.6
0.8
0.4
0
PBS

**B.**

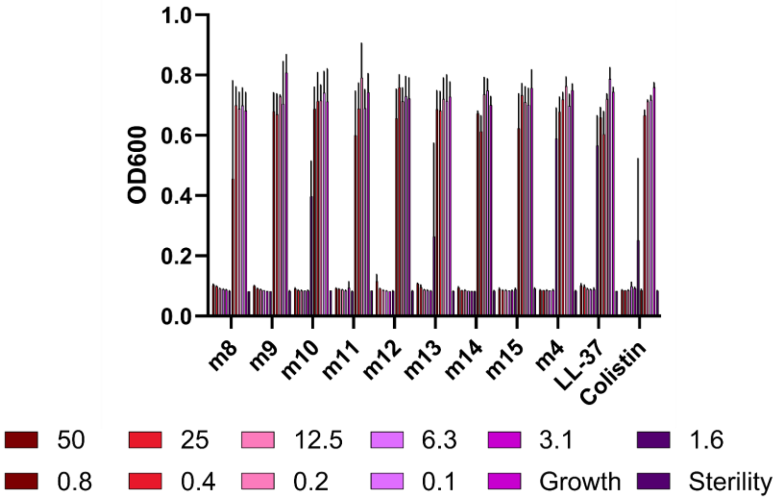
OD600
1.0
0.8
0.6
0.4
0.2
0.0
m8
m9
m10
m11
m12
m13
m14
m15
m4
LL-37
Colistin
50
25
12.5
6.3
3.1
1.6
0.8
0.4
0.2
0.1
Growth
Sterility

**C.**

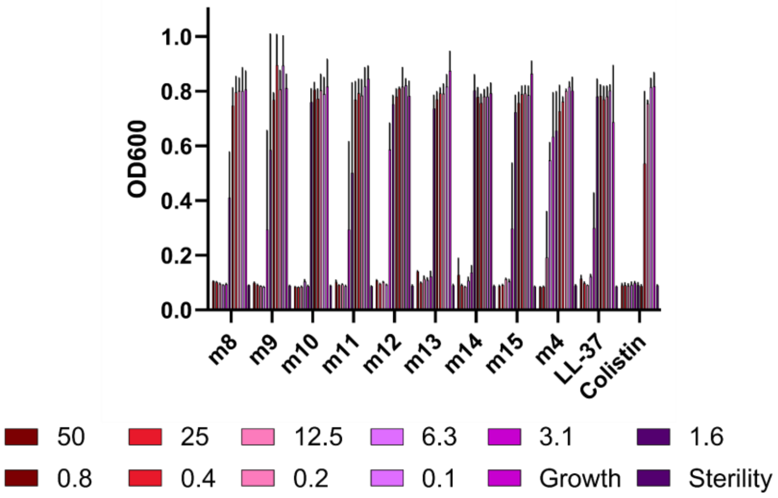
OD600
1.0
0.8
0.6
0.4
0.2
0.0
m8
m9
m10
m11
m12
m13
m14
m15
m4
LL-37
Colistin
50
25
12.5
6.3
3.1
1.6
0.8
0.4
0.2
0.1
Growth
Sterility

**Supplementary Figure 22 – Histogram depiction of Figure 2 data. A.** Endpoint hemolytic activity of each peptide relative to a 100% hemolysis freeze-thaw control after 18 hours at 37 °C. Raw data were obtained by absorbance at 414 nm in well supernatants for free hemoglobin signal. **B-C.** Endpoint activity of each peptide against *E. coli* (**B**) or *P. aeruginosa* (**C**). Data are the raw OD600 after 18 hours at 37 °C. Peptide concentrations are given in μM. Data throughout show the mean and standard deviation of three independent experiments.

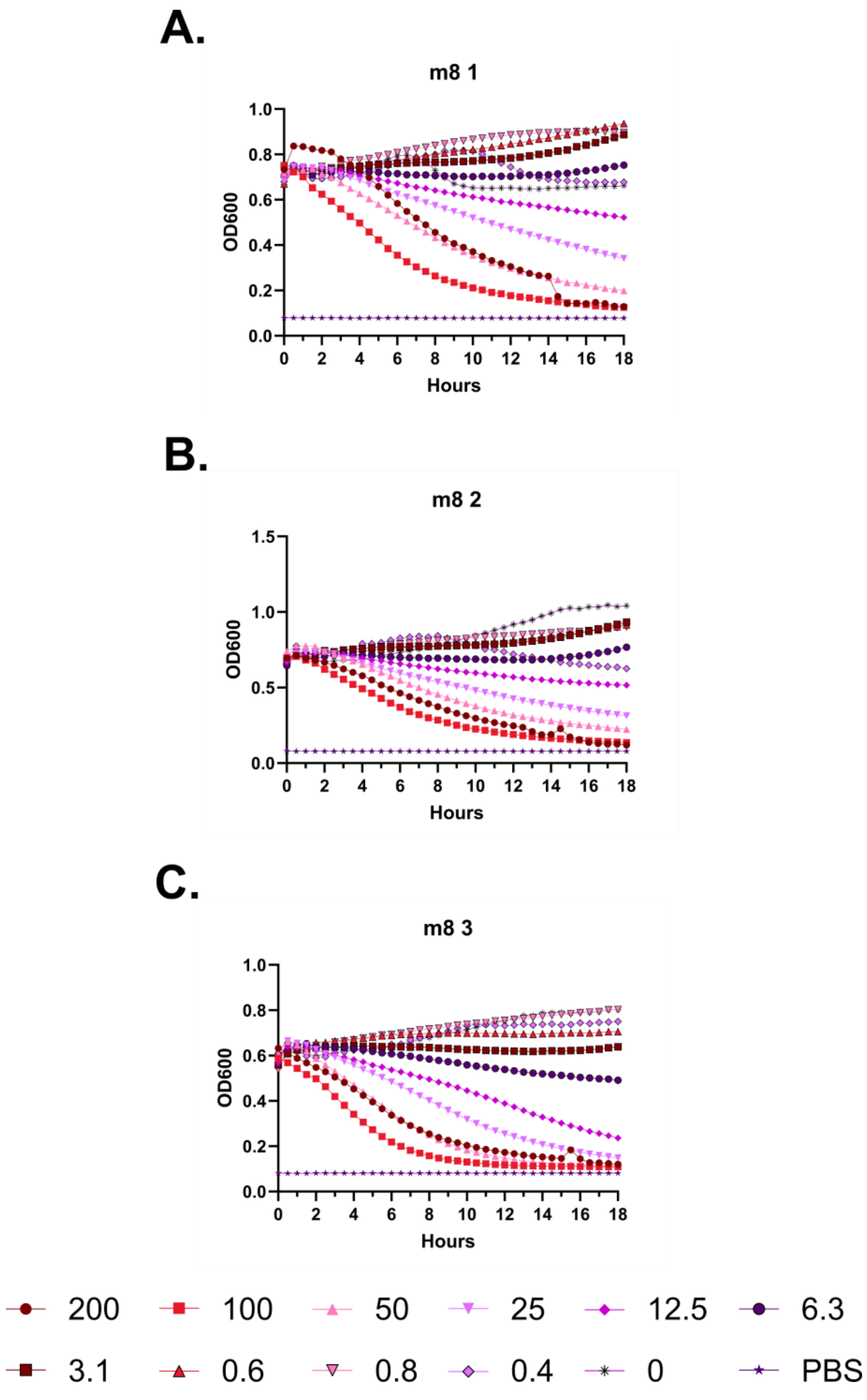
A.
m8 1
OD600
Hours
B.
m8 2
OD600
Hours
C.
m8 3
OD600
Hours
200
100
50
25
12.5
6.3
3.1
0.6
0.8
0.4
0
PBS

**Supplementary Figure 23 – Kinetic hemolysis assays for LL-37 m8. A-C.** Hemolysis was monitored by OD600 over 18 hours at 37 °C in the presence of the indicated peptide concentrations. Decreasing OD600 indicates increasing hemolysis. Peptide concentrations are given in μM. Each curve indicates an independent experiment.

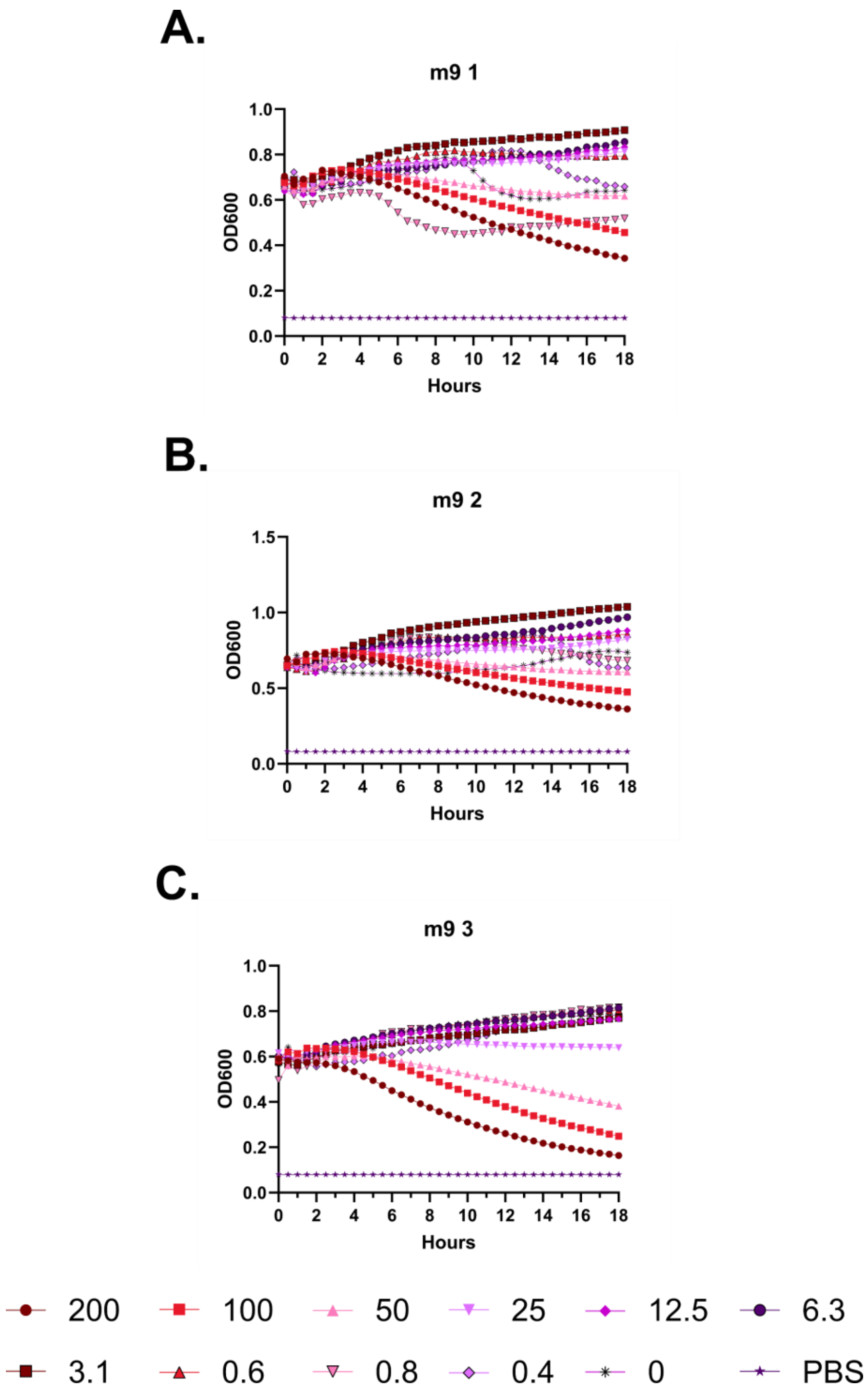
A.
m9 1
OD600
Hours
0.0
0.2
0.4
0.6
0.8
1.0
0 2 4 6 8 10 12 14 16 18
B.
m9 2
OD600
Hours
0.0
0.5
1.0
1.5
0 2 4 6 8 10 12 14 16 18
C.
m9 3
OD600
Hours
0.0
0.2
0.4
0.6
0.8
1.0
0 2 4 6 8 10 12 14 16 18
200
100
50
25
12.5
6.3
3.1
0.6
0.8
0.4
0
PBS

**Supplementary Figure 24 – Kinetic hemolysis assays for LL-37 m9. A-C.** Hemolysis was monitored by OD600 over 18 hours at 37 °C in the presence of the indicated peptide concentrations. Decreasing OD600 indicates increasing hemolysis. Peptide concentrations are given in μM. Each curve indicates an independent experiment.

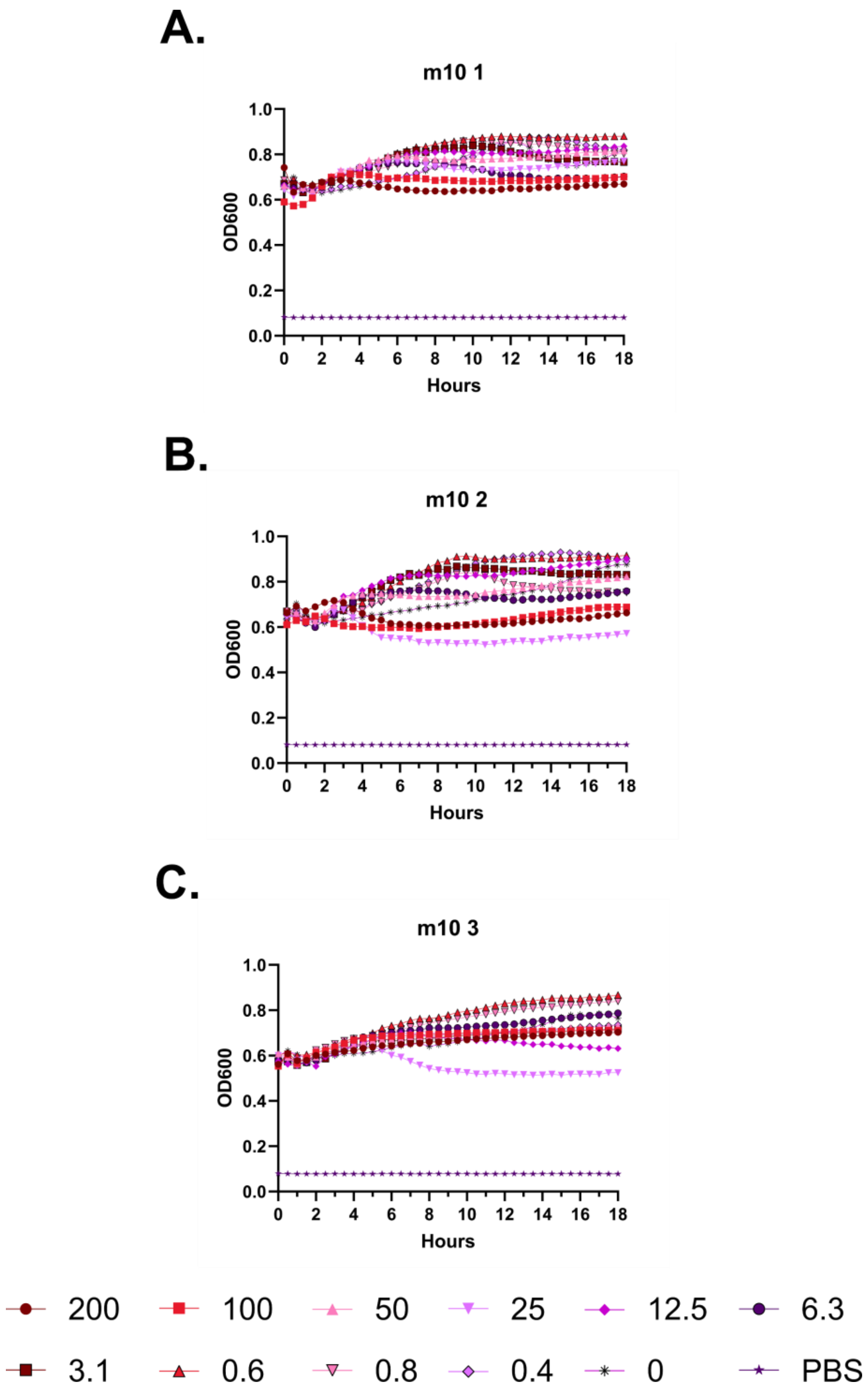
A.
m10 1
OD600
Hours
B.
m10 2
OD600
Hours
C.
m10 3
OD600
Hours
200
100
50
25
12.5
6.3
3.1
0.6
0.8
0.4
0
PBS

**Supplementary Figure 25 – Kinetic hemolysis assays for LL-37 m10. A-C.** Hemolysis was monitored by OD600 over 18 hours at 37 °C in the presence of the indicated peptide concentrations. Decreasing OD600 indicates increasing hemolysis. Peptide concentrations are given in μM. Each curve indicates an independent experiment.

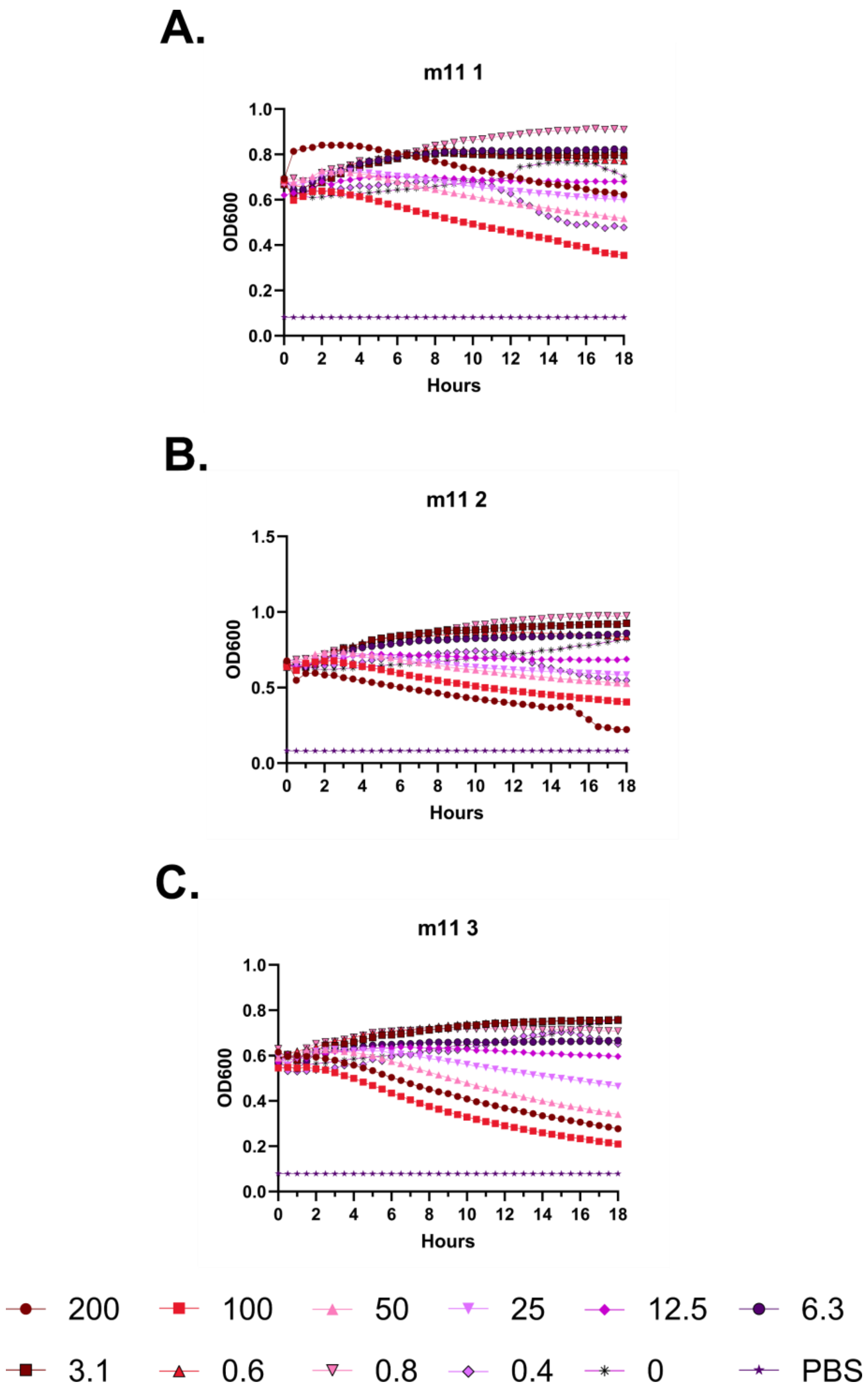
A.
m11 1
OD600
1.0
0.8
0.6
0.4
0.2
0.0
0 2 4 6 8 10 12 14 16 18
Hours
B.
m11 2
OD600
1.5
1.0
0.5
0.0
0 2 4 6 8 10 12 14 16 18
Hours
C.
m11 3
OD600
1.0
0.8
0.6
0.4
0.2
0.0
0 2 4 6 8 10 12 14 16 18
Hours
200
100
50
25
12.5
6.3
3.1
0.6
0.8
0.4
0
PBS

**Supplementary Figure 26 – Kinetic hemolysis assays for LL-37 m11. A-C.** Hemolysis was monitored by OD600 over 18 hours at 37 °C in the presence of the indicated peptide concentrations. Decreasing OD600 indicates increasing hemolysis. Peptide concentrations are given in μM. Each curve indicates an independent experiment.

**A.**

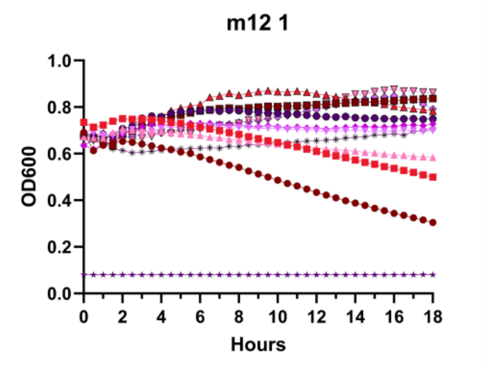
m12 1
OD600
Hours
1.0
0.8
0.6
0.4
0.2
0.0
0
2
4
6
8
10
12
14
16
18

**B.**

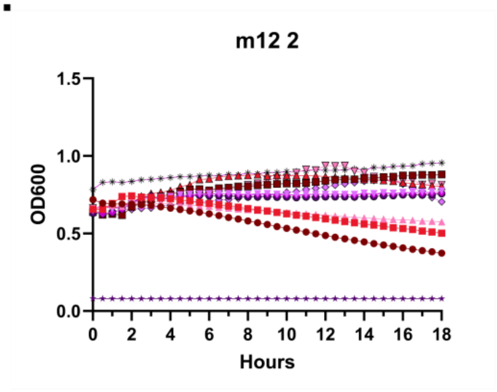
m12 2
OD600
Hours
1.5
1.0
0.5
0.0
0
2
4
6
8
10
12
14
16
18

**C.**

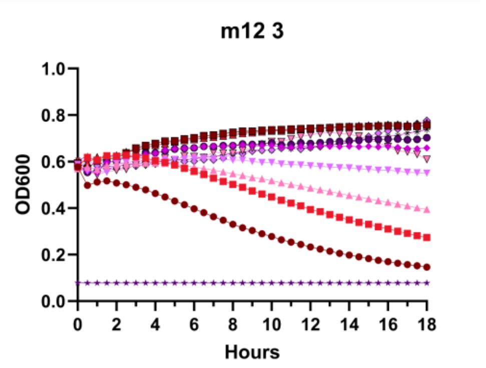
m12 3
OD600
Hours
1.0
0.8
0.6
0.4
0.2
0.0
0
2
4
6
8
10
12
14
16
18

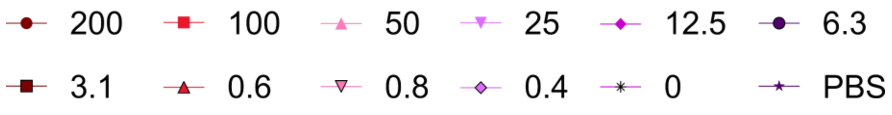
200
100
50
25
12.5
6.3
3.1
0.6
0.8
0.4
0
PBS

**Supplementary Figure 27 – Kinetic hemolysis assays for LL-37 m12. A-C.** Hemolysis was monitored by OD600 over 18 hours at 37 °C in the presence of the indicated peptide concentrations. Decreasing OD600 indicates increasing hemolysis. Peptide concentrations are given in μM. Each curve indicates an independent experiment.

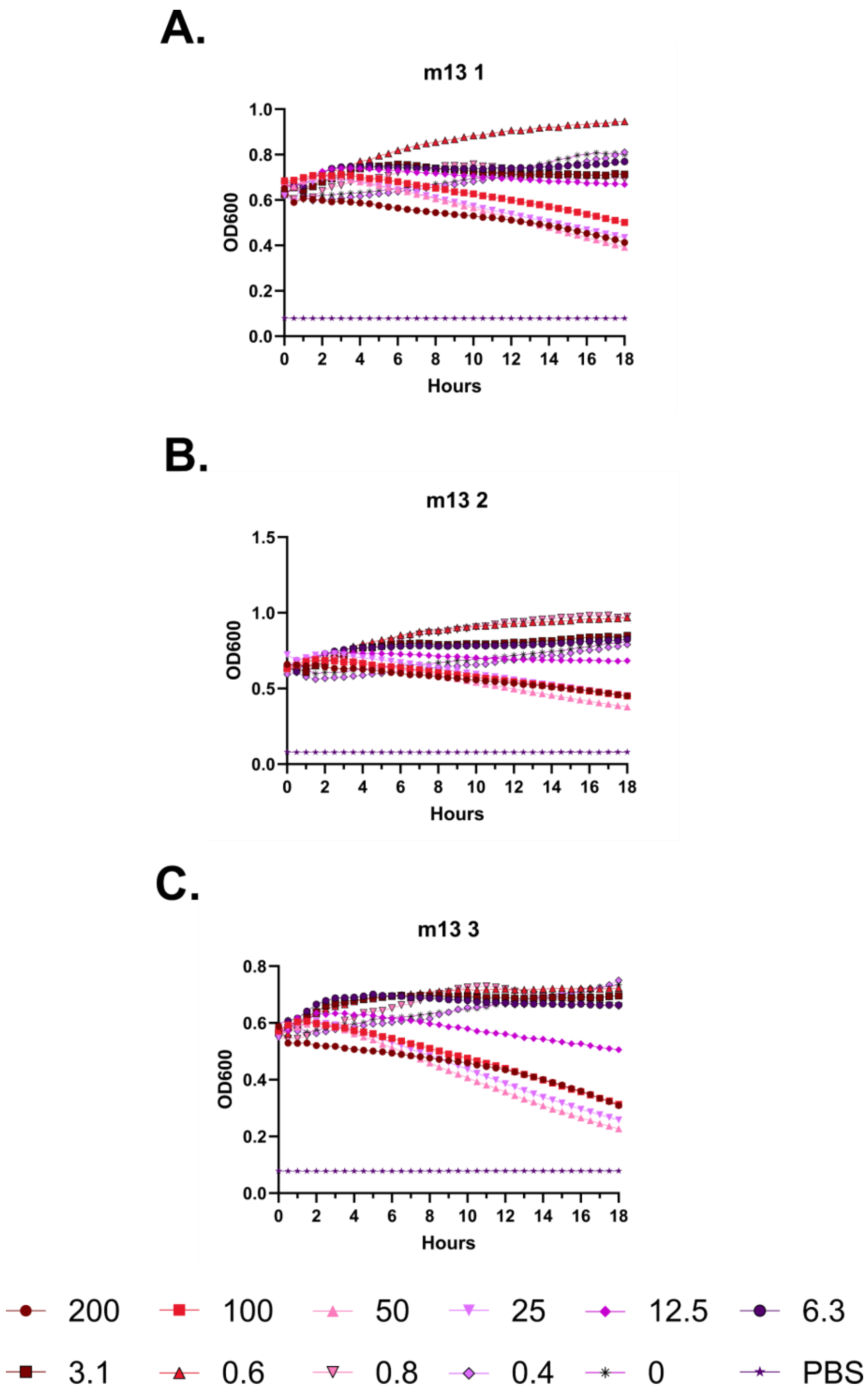
A.
m13 1
OD600
Hours
B.
m13 2
OD600
Hours
C.
m13 3
OD600
Hours
200
100
50
25
12.5
6.3
3.1
0.6
0.8
0.4
0
PBS

**Supplementary Figure 28 – Kinetic hemolysis assays for LL-37 m13. A-C.** Hemolysis was monitored by OD600 over 18 hours at 37 °C in the presence of the indicated peptide concentrations. Decreasing OD600 indicates increasing hemolysis. Peptide concentrations are given in μM. Each curve indicates an independent experiment.

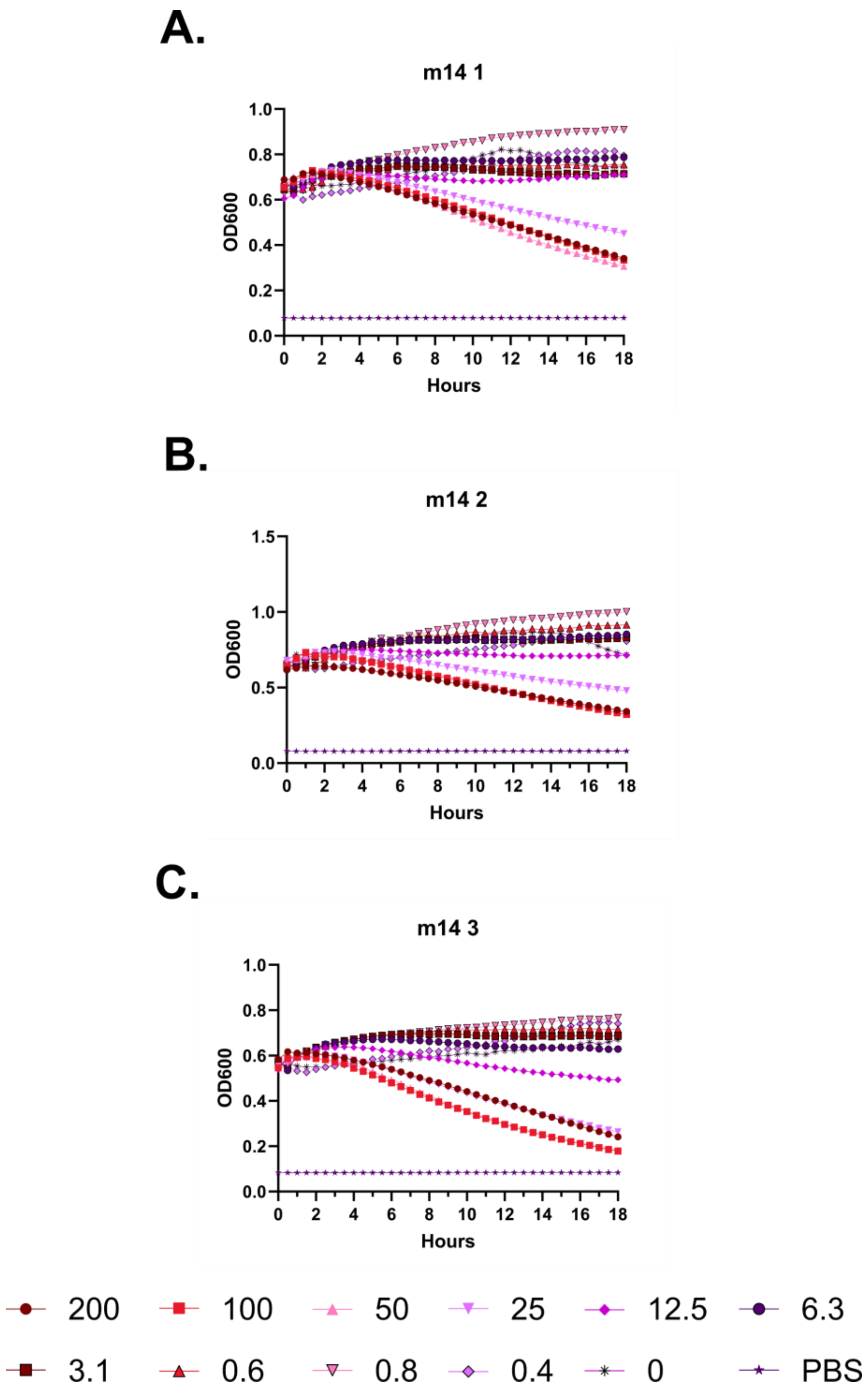
A.
m14 1
OD600
Hours
B.
m14 2
OD600
Hours
C.
m14 3
OD600
Hours
200
100
50
25
12.5
6.3
3.1
0.6
0.8
0.4
0
PBS

**Supplementary Figure 29 – Kinetic hemolysis assays for LL-37 m14. A-C.** Hemolysis was monitored by OD600 over 18 hours at 37 °C in the presence of the indicated peptide concentrations. Decreasing OD600 indicates increasing hemolysis. Peptide concentrations are given in μM. Each curve indicates an independent experiment.

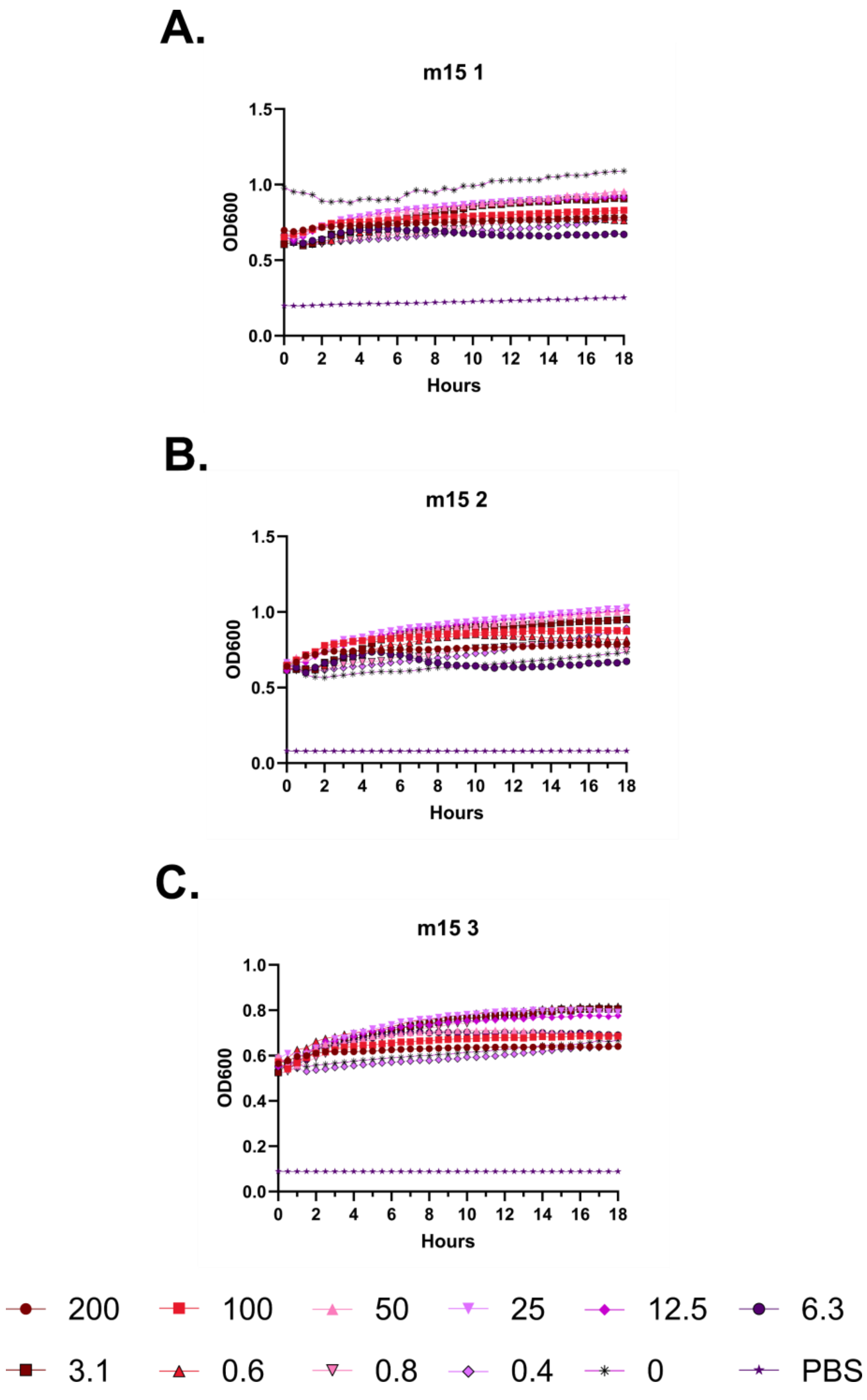
A.
m15 1
OD600
1.5
1.0
0.5
0.0
0 2 4 6 8 10 12 14 16 18
Hours
B.
m15 2
OD600
1.5
1.0
0.5
0.0
0 2 4 6 8 10 12 14 16 18
Hours
C.
m15 3
OD600
1.0
0.8
0.6
0.4
0.2
0.0
0 2 4 6 8 10 12 14 16 18
Hours
200
100
50
25
12.5
6.3
3.1
0.6
0.8
0.4
0
PBS

**Supplementary Figure 30 – Kinetic hemolysis assays for LL-37 m15. A-C.** Hemolysis was monitored by OD600 over 18 hours at 37 °C in the presence of the indicated peptide concentrations. Decreasing OD600 indicates increasing hemolysis. Peptide concentrations are given in µM. Each curve indicates an independent experiment.

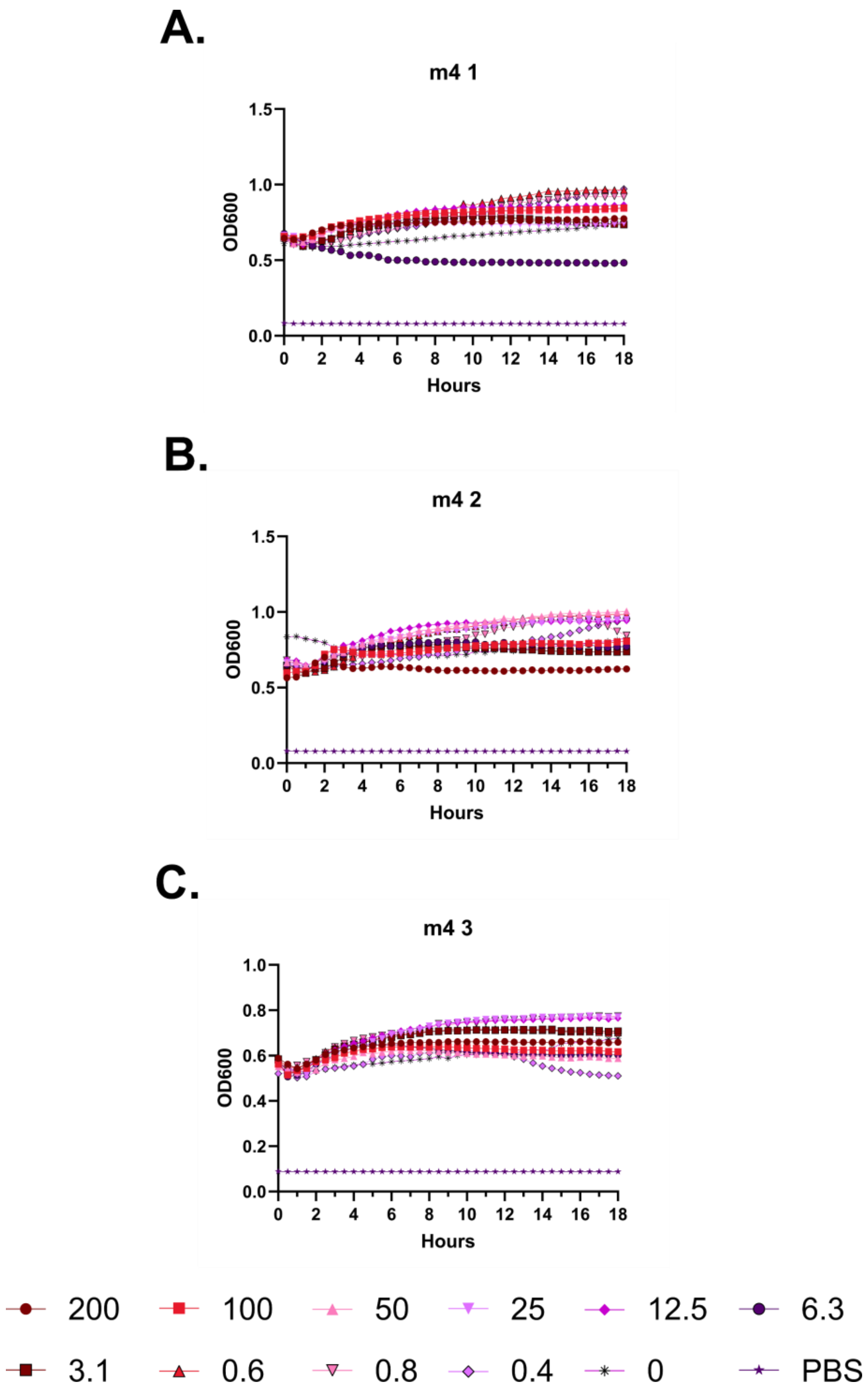
A.
m4 1
OD600
Hours
B.
m4 2
OD600
Hours
C.
m4 3
OD600
Hours
200
100
50
25
12.5
6.3
3.1
0.6
0.8
0.4
0
PBS

**Supplementary Figure 31 – Kinetic hemolysis assays for LL-37 m4. A-C.** Hemolysis was monitored by OD600 over 18 hours at 37 °C in the presence of the indicated peptide concentrations. Decreasing OD600 indicates increasing hemolysis. Peptide concentrations are given in μM. Each curve indicates an independent experiment.

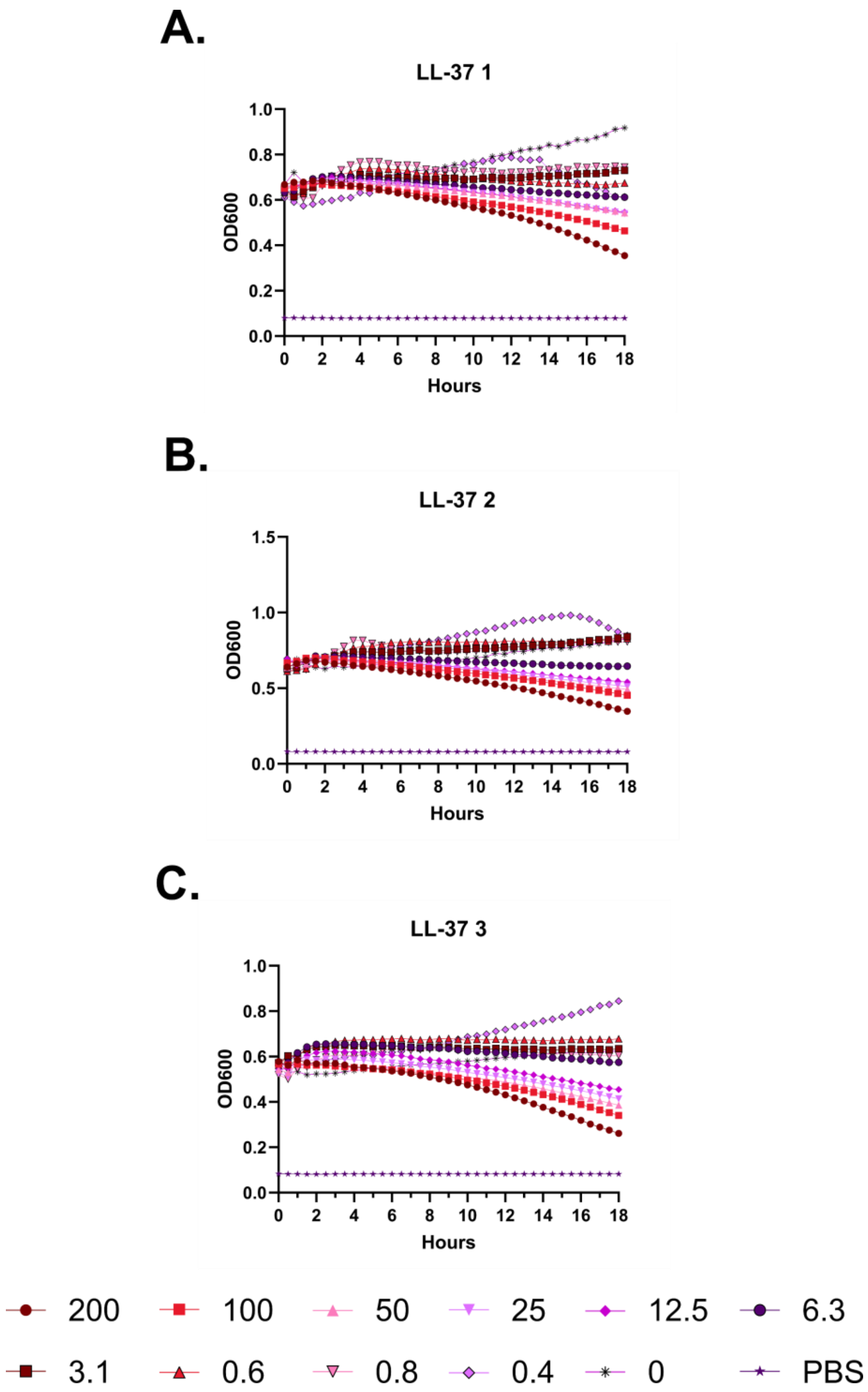
A.
LL-37 1
OD600
Hours
B.
LL-37 2
OD600
Hours
C.
LL-37 3
OD600
Hours
200
100
50
25
12.5
6.3
3.1
0.6
0.8
0.4
0
PBS

**Supplementary Figure 32 – Kinetic hemolysis assays for LL-37. A-C.** Hemolysis was monitored by OD600 over 18 hours at 37 °C in the presence of the indicated peptide concentrations. Decreasing OD600 indicates increasing hemolysis. Peptide concentrations are given in μM. Each curve indicates an independent experiment.

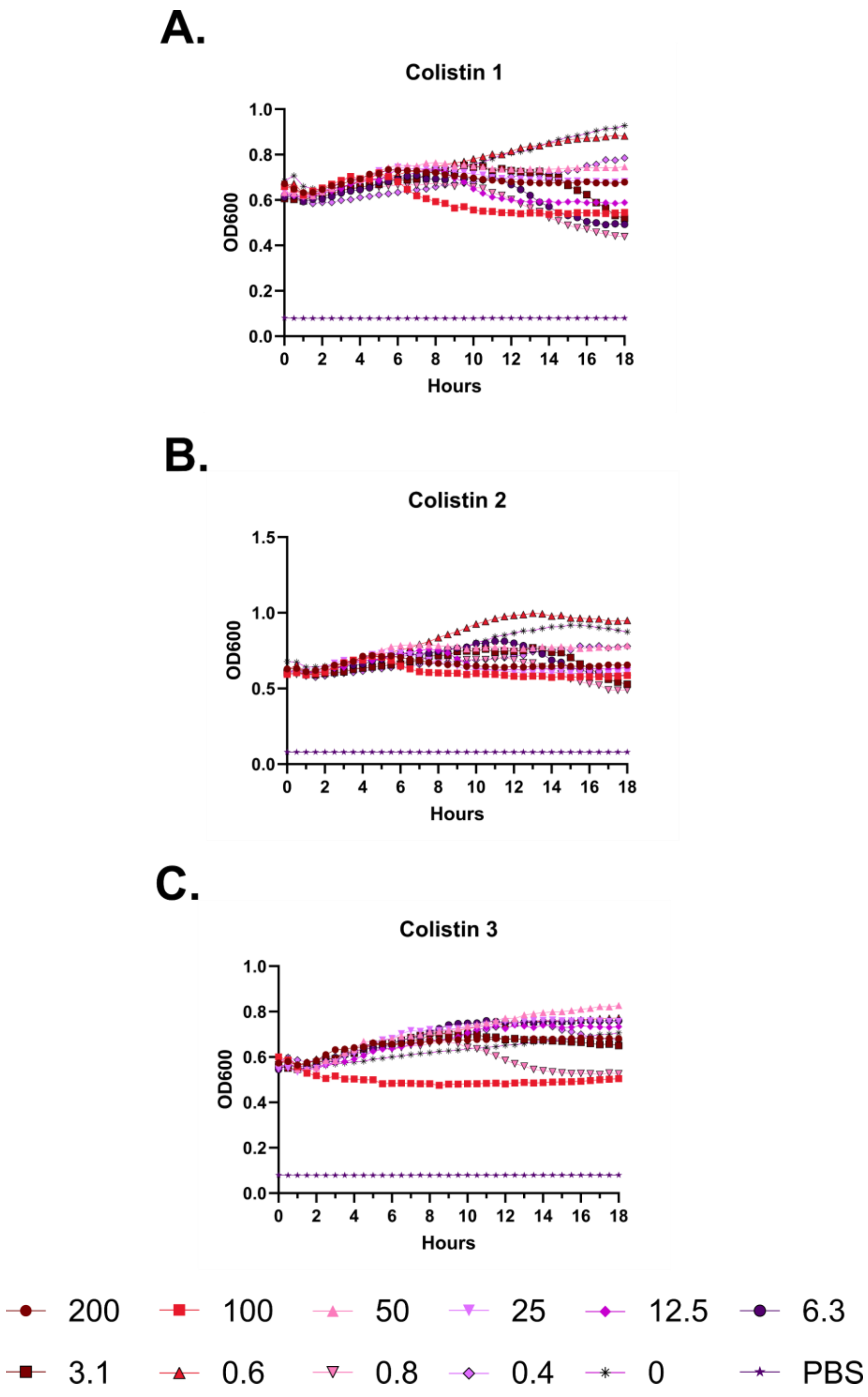
A.
Colistin 1
OD600
Hours
B.
Colistin 2
OD600
Hours
C.
Colistin 3
OD600
Hours
200
100
50
25
12.5
6.3
3.1
0.6
0.8
0.4
0
PBS

**Supplementary Figure 33 – Kinetic hemolysis assays for colistin. A-C.** Hemolysis was monitored by OD600 over 18 hours at 37 °C in the presence of the indicated peptide concentrations. Decreasing OD600 indicates increasing hemolysis. Peptide concentrations are given in μM. Each curve indicates an independent experiment.

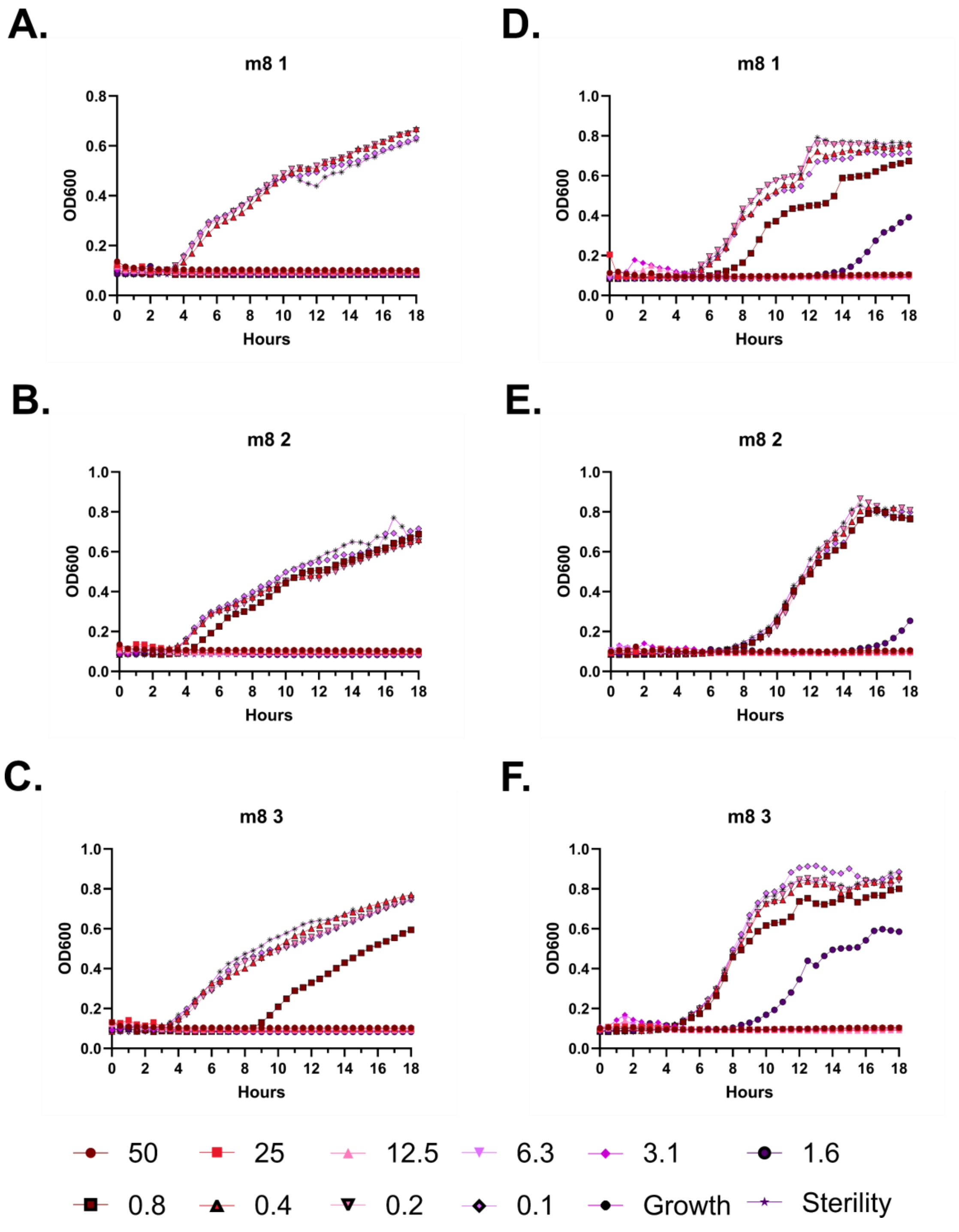
A.
m8 1
OD600
Hours
B.
m8 2
OD600
Hours
C.
m8 3
OD600
Hours
D.
m8 1
OD600
Hours
E.
m8 2
OD600
Hours
F.
m8 3
OD600
Hours
50
25
12.5
6.3
3.1
1.6
0.8
0.4
0.2
0.1
Growth
Sterility

**Supplementary Figure 34 – Kinetic antimicrobial susceptibility testing for LL-37 m8. A-C.** *E. coli*. **D-F.** *P. aeruginosa.* Bacterial growth was monitored by OD600 over 18 hours at 37 °C in the presence of the indicated peptide concentrations. Increasing OD600 indicates increasing bacterial growth. Peptide concentrations are given in μM. Each curve indicates an independent experiment.

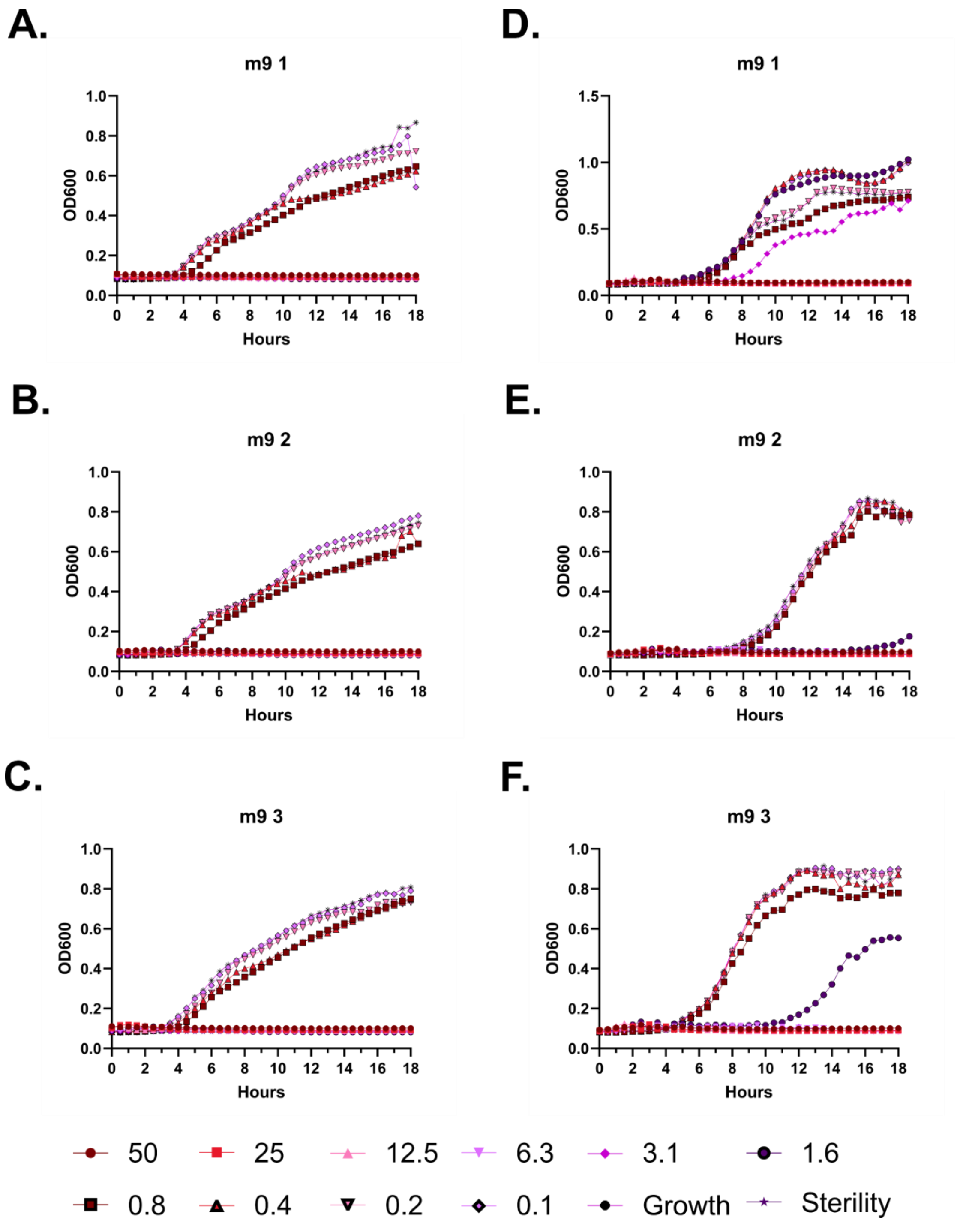
A.
m9 1
OD600
Hours
D.
m9 1
OD600
Hours
B.
m9 2
OD600
Hours
E.
m9 2
OD600
Hours
C.
m9 3
OD600
Hours
F.
m9 3
OD600
Hours
50
25
12.5
6.3
3.1
1.6
0.8
0.4
0.2
0.1
Growth
Sterility

**Supplementary Figure 35 – Kinetic antimicrobial susceptibility testing for LL-37 m9. A-C.** *E. coli*. **D-F.** *P. aeruginosa.* Bacterial growth was monitored by OD600 over 18 hours at 37 °C in the presence of the indicated peptide concentrations. Increasing OD600 indicates increasing bacterial growth. Peptide concentrations are given in μM. Each curve indicates an independent experiment.

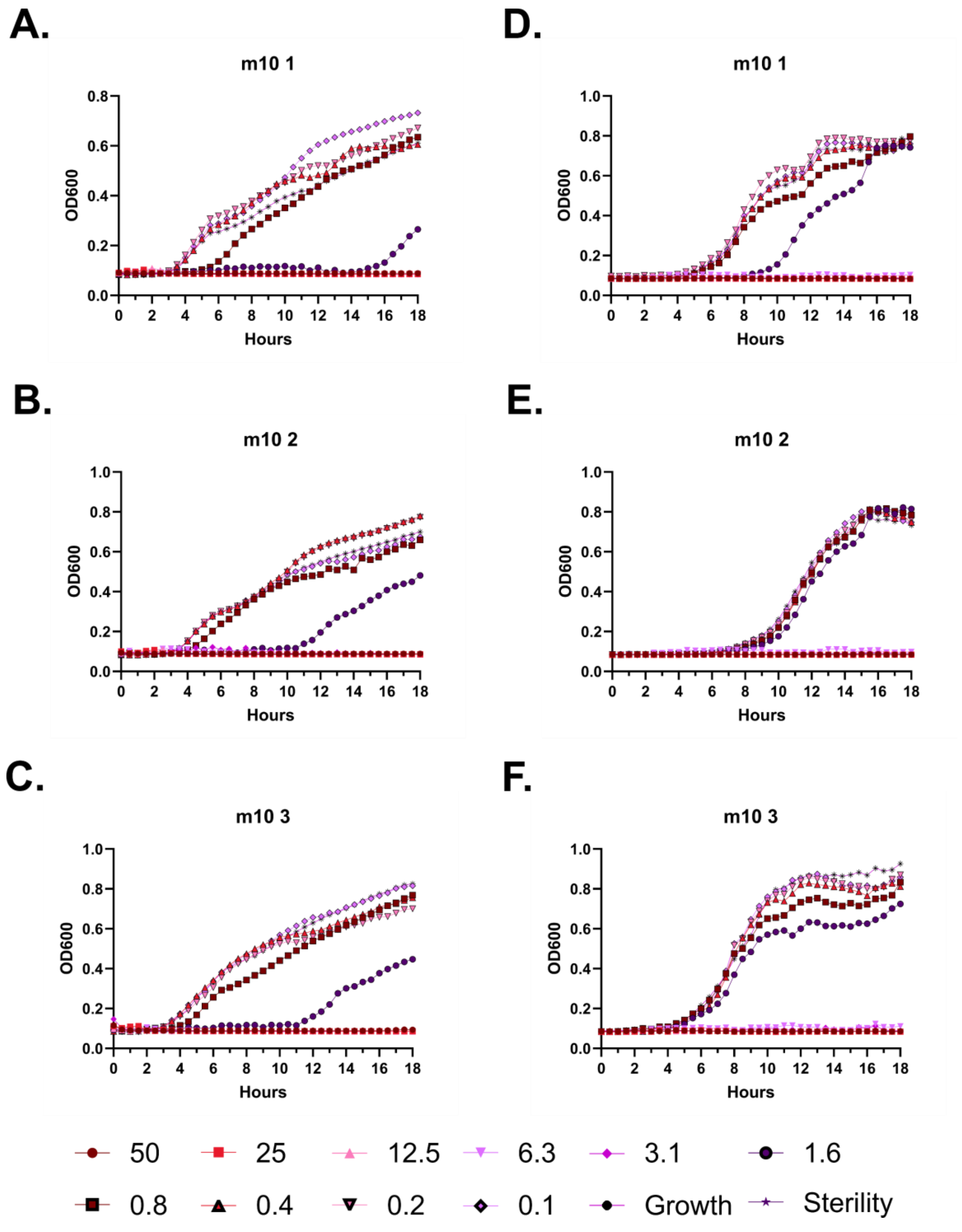
A.
m10 1
OD600
Hours
B.
m10 2
C.
m10 3
D.
m10 1
E.
m10 2
F.
m10 3
50
25
12.5
6.3
3.1
1.6
0.8
0.4
0.2
0.1
Growth
Sterility

**Supplementary Figure 36 – Kinetic antimicrobial susceptibility testing for LL-37 m10. A-C.** *E. coli*. **D-F.** *P. aeruginosa.* Bacterial growth was monitored by OD600 over 18 hours at 37 °C in the presence of the indicated peptide concentrations. Increasing OD600 indicates increasing bacterial growth. Peptide concentrations are given in μM. Each curve indicates an independent experiment.

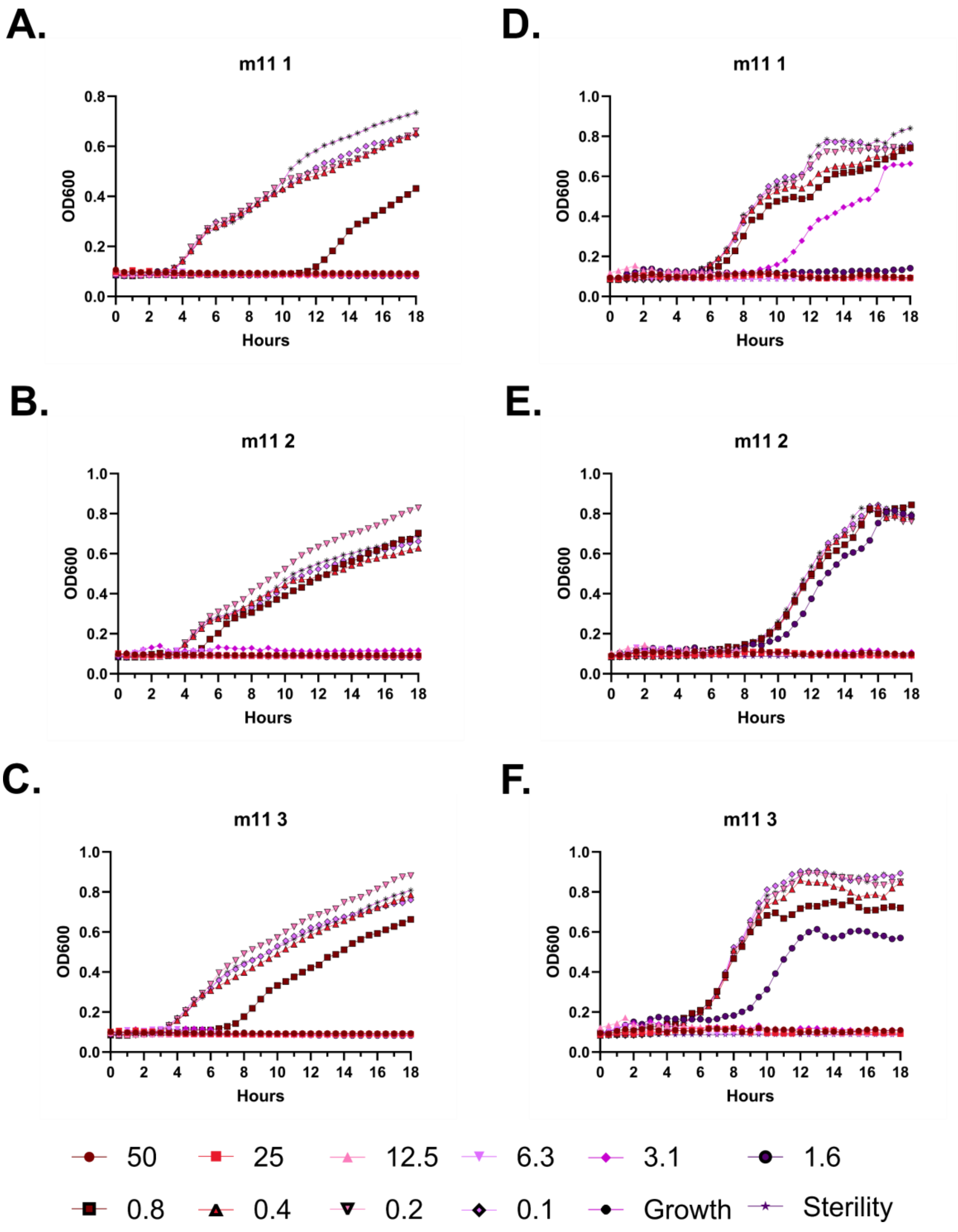
A.
m11 1
OD600
Hours
B.
m11 2
OD600
Hours
C.
m11 3
OD600
Hours
D.
m11 1
OD600
Hours
E.
m11 2
OD600
Hours
F.
m11 3
OD600
Hours
50
25
12.5
6.3
3.1
1.6
0.8
0.4
0.2
0.1
Growth
Sterility

**Supplementary Figure 37 – Kinetic antimicrobial susceptibility testing for LL-37 m11. A-C.** *E. coli*. **D-F.** *P. aeruginosa.* Bacterial growth was monitored by OD600 over 18 hours at 37 °C in the presence of the indicated peptide concentrations. Increasing OD600 indicates increasing bacterial growth. Peptide concentrations are given in μM. Each curve indicates an independent experiment.

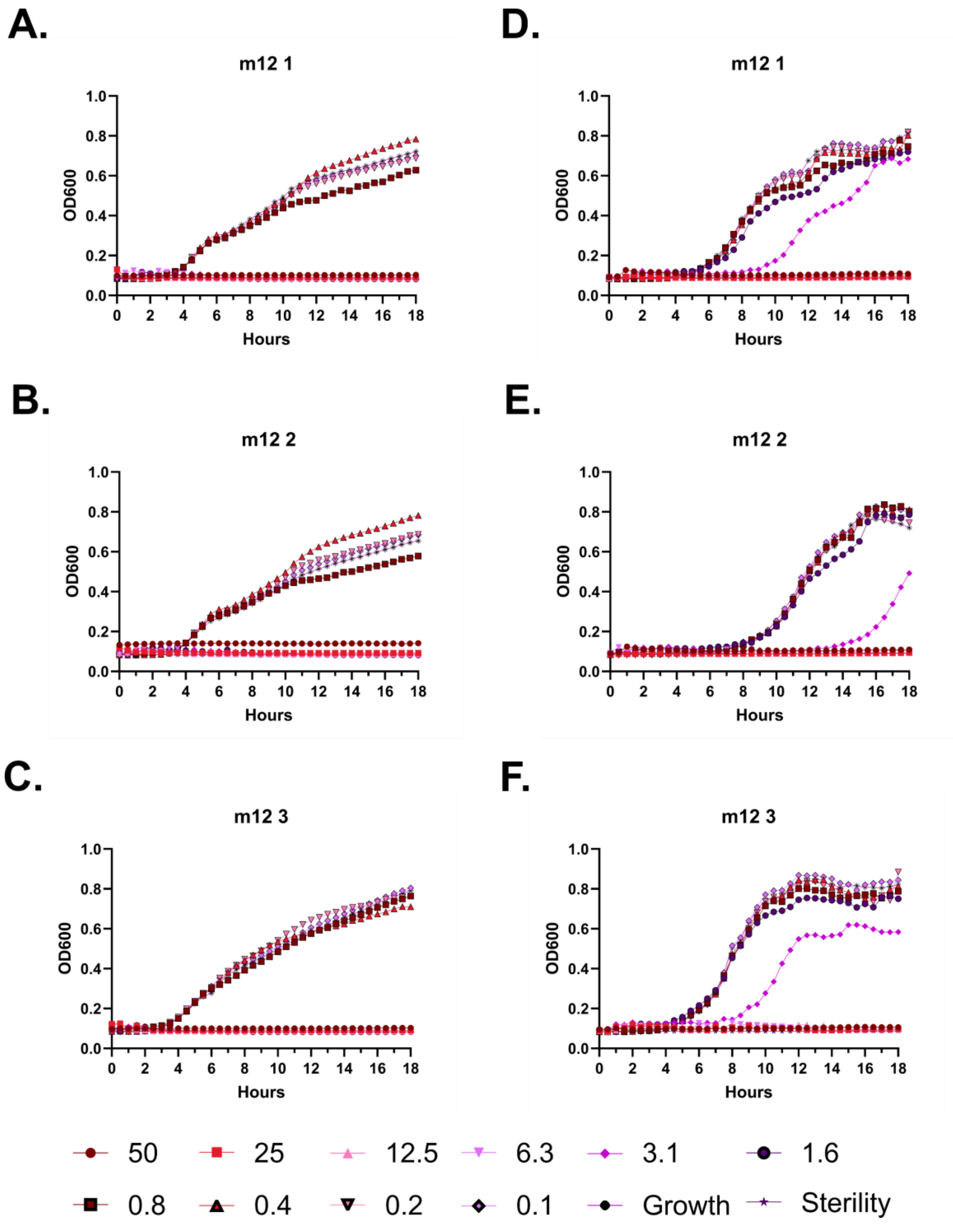
A.
m12 1
D.
m12 1
B.
m12 2
E.
m12 2
C.
m12 3
F.
m12 3
OD600
Hours
50
25
12.5
6.3
3.1
1.6
0.8
0.4
0.2
0.1
Growth
Sterility

**Supplementary Figure 38 – Kinetic antimicrobial susceptibility testing for LL-37 m12. A-C.** *E. coli*. **D-F.** *P. aeruginosa.* Bacterial growth was monitored by OD600 over 18 hours at 37 °C in the presence of the indicated peptide concentrations. Increasing OD600 indicates increasing bacterial growth. Peptide concentrations are given in µM. Each curve indicates an independent experiment.

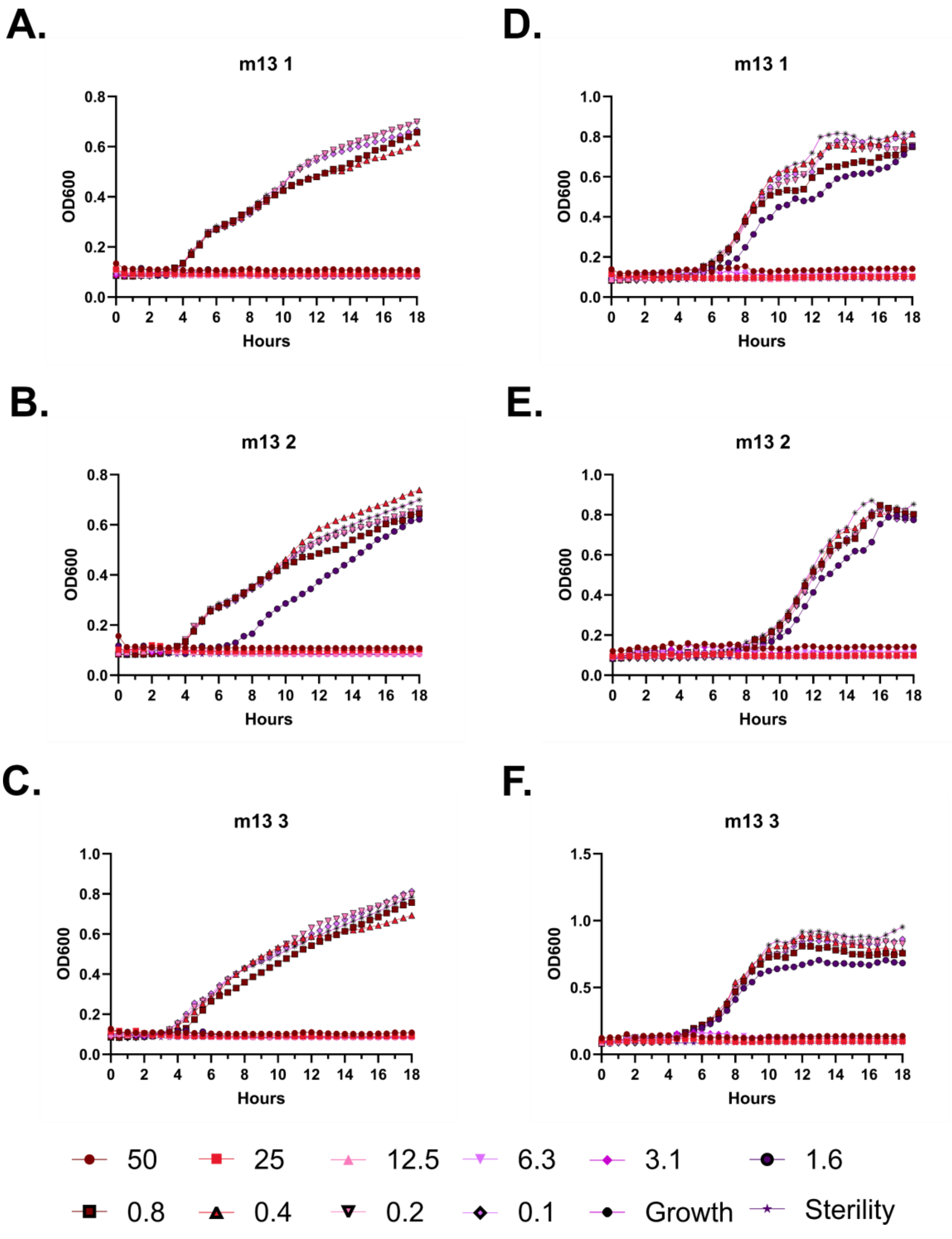
A.
m13 1
OD600
Hours
D.
m13 1
OD600
Hours
B.
m13 2
OD600
Hours
E.
m13 2
OD600
Hours
C.
m13 3
OD600
Hours
F.
m13 3
OD600
Hours
50
25
12.5
6.3
3.1
1.6
0.8
0.4
0.2
0.1
Growth
Sterility

**Supplementary Figure 39 – Kinetic antimicrobial susceptibility testing for LL-37 m13. A-C.** *E. coli*. **D-F.** *P. aeruginosa.* Bacterial growth was monitored by OD600 over 18 hours at 37 °C in the presence of the indicated peptide concentrations. Increasing OD600 indicates increasing bacterial growth. Peptide concentrations are given in μM. Each curve indicates an independent experiment.

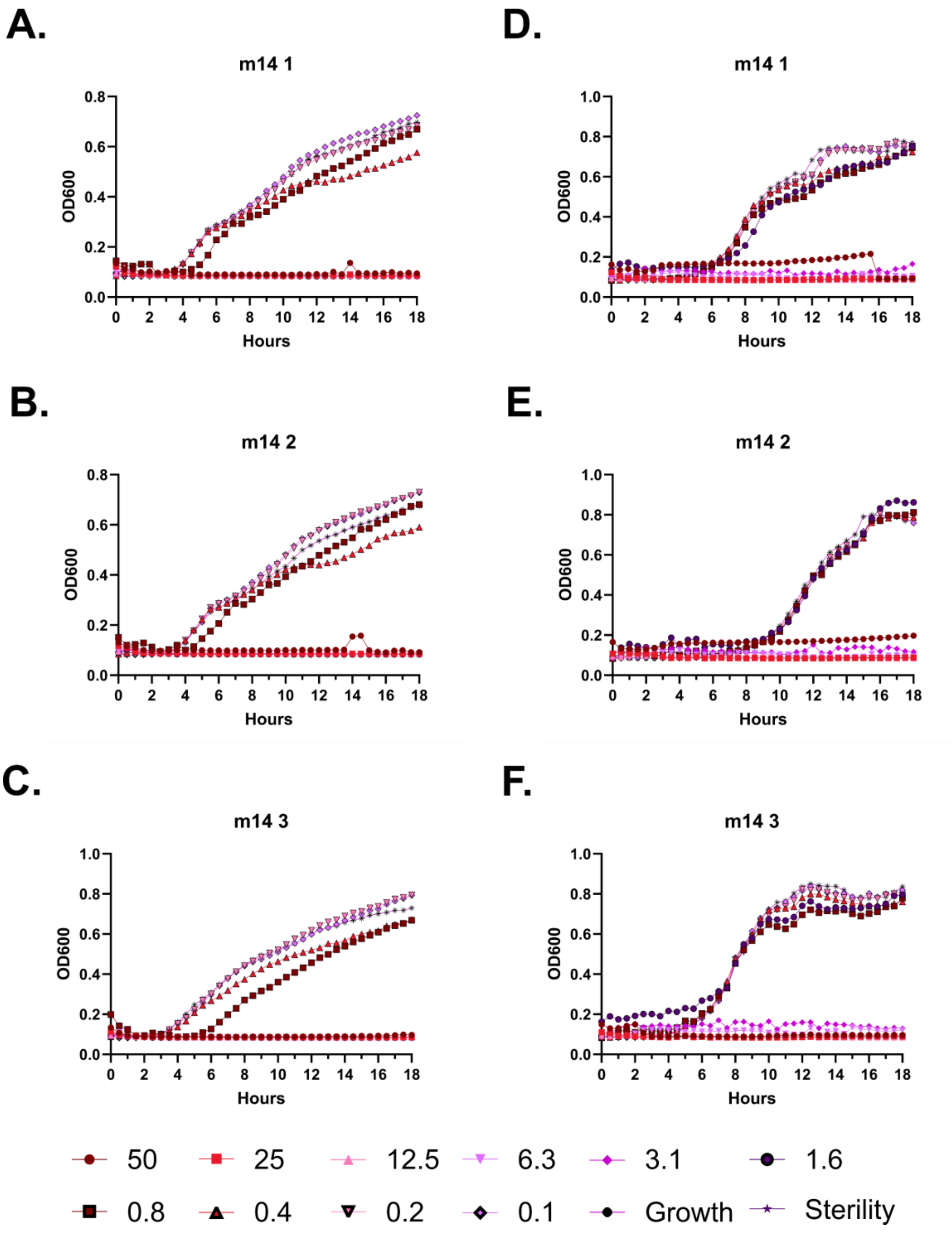
A.
m14 1
OD600
Hours
D.
m14 1
OD600
Hours
B.
m14 2
OD600
Hours
E.
m14 2
OD600
Hours
C.
m14 3
OD600
Hours
F.
m14 3
OD600
Hours
50
25
12.5
6.3
3.1
1.6
0.8
0.4
0.2
0.1
Growth
Sterility

**Supplementary Figure 40 – Kinetic antimicrobial susceptibility testing for LL-37 m14. A-C.** *E. coli*. **D-F.** *P. aeruginosa.* Bacterial growth was monitored by OD600 over 18 hours at 37 °C in the presence of the indicated peptide concentrations. Increasing OD600 indicates increasing bacterial growth. Peptide concentrations are given in μM. Each curve indicates an independent experiment.

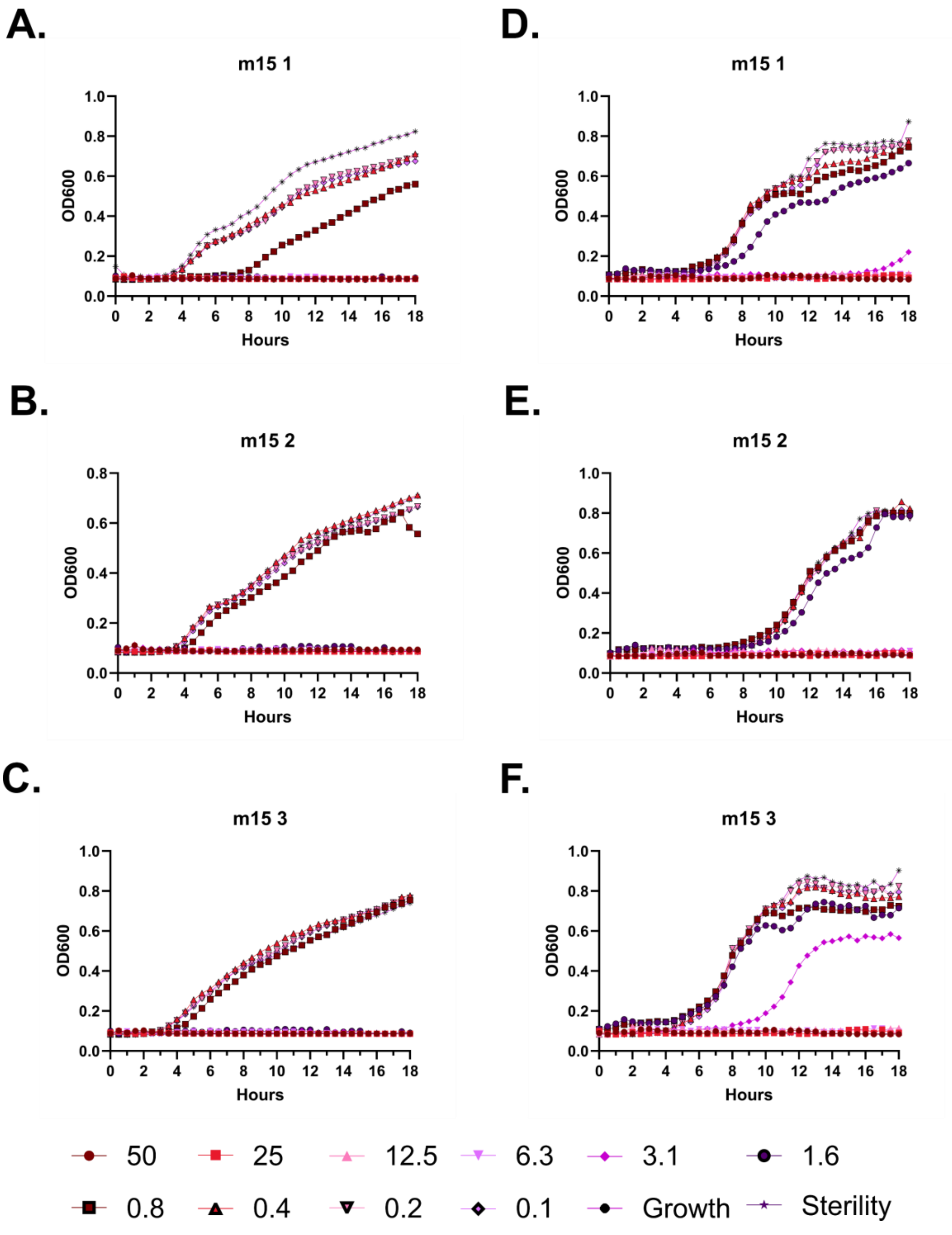
A.
m15 1
OD600
Hours
D.
m15 1
OD600
Hours
B.
m15 2
OD600
Hours
E.
m15 2
OD600
Hours
C.
m15 3
OD600
Hours
F.
m15 3
OD600
Hours
50
25
12.5
6.3
3.1
1.6
0.8
0.4
0.2
0.1
Growth
Sterility

**Supplementary Figure 41 – Kinetic antimicrobial susceptibility testing for LL-37 m15. A-C.** *E. coli*. **D-F.** *P. aeruginosa.* Bacterial growth was monitored by OD600 over 18 hours at 37 °C in the presence of the indicated peptide concentrations. Increasing OD600 indicates increasing bacterial growth. Peptide concentrations are given in μM. Each curve indicates an independent experiment.

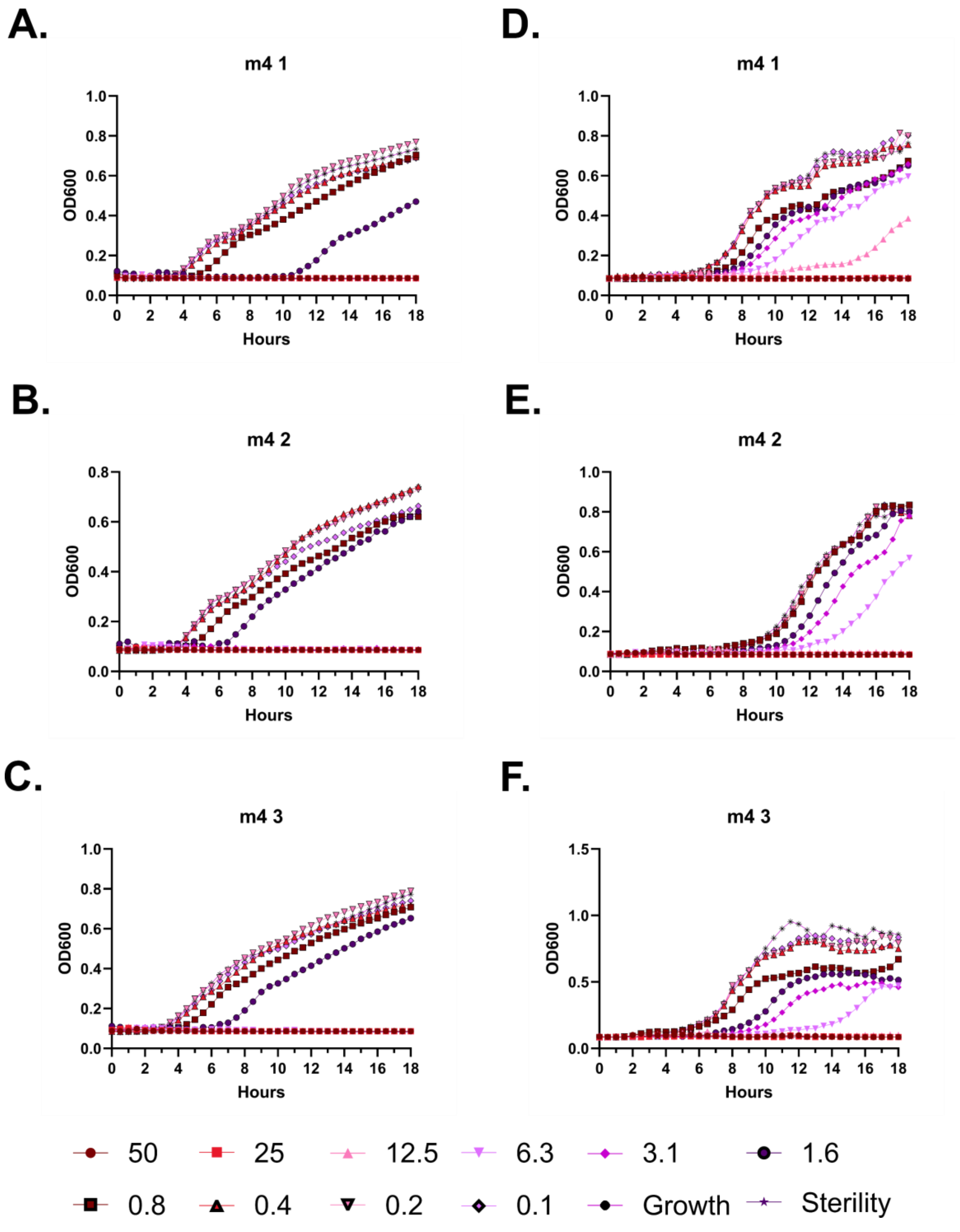
A.
m4 1
B.
m4 2
C.
m4 3
D.
m4 1
E.
m4 2
F.
m4 3
OD600
Hours
50
25
12.5
6.3
3.1
1.6
0.8
0.4
0.2
0.1
Growth
Sterility

**Supplementary Figure 42 – Kinetic antimicrobial susceptibility testing for LL-37 m4. A-C.** *E. coli*. **D-F.** *P. aeruginosa.* Bacterial growth was monitored by OD600 over 18 hours at 37 °C in the presence of the indicated peptide concentrations. Increasing OD600 indicates increasing bacterial growth. Peptide concentrations are given in μM. Each curve indicates an independent experiment.

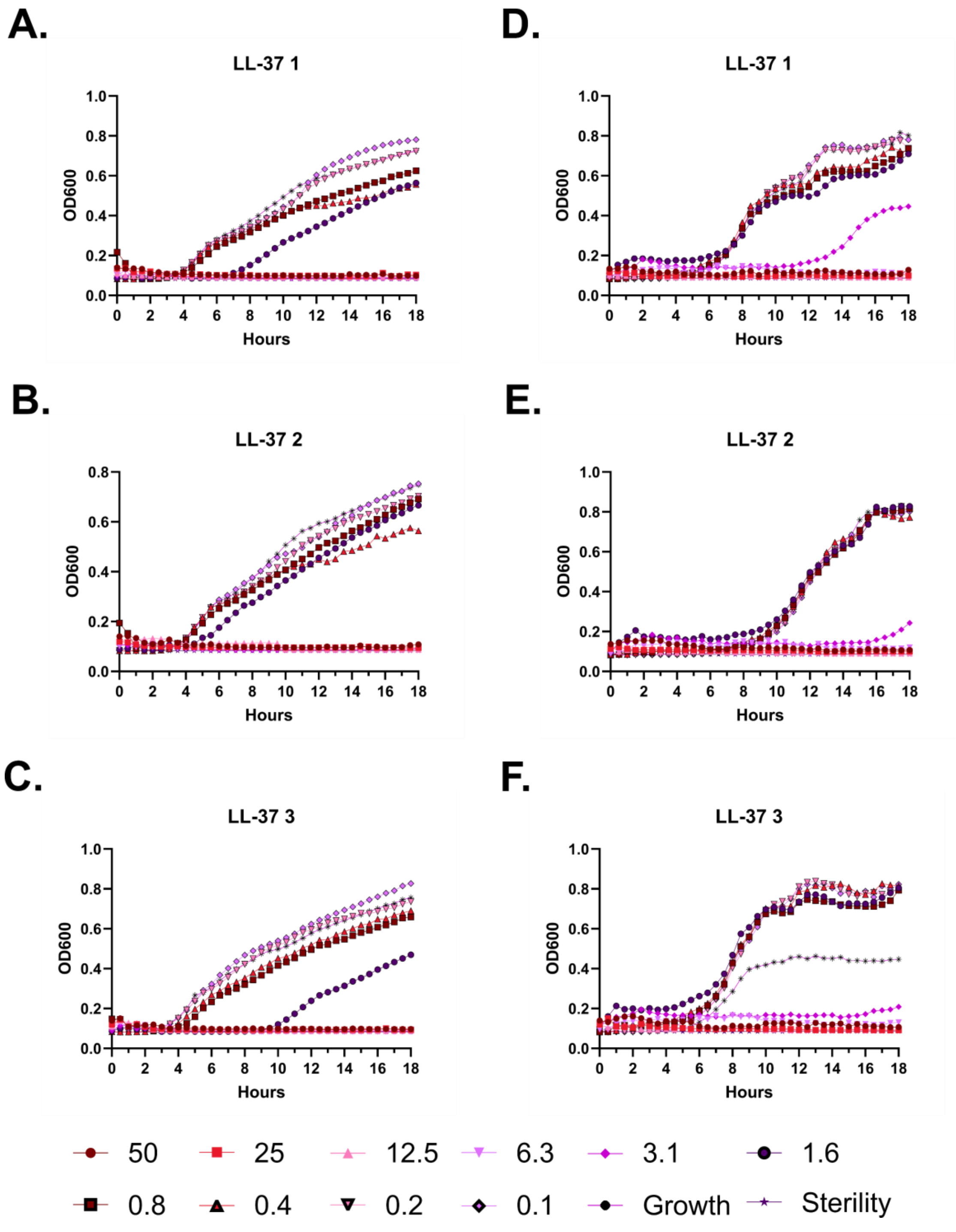
A.
LL-37 1
B.
LL-37 2
C.
LL-37 3
D.
LL-37 1
E.
LL-37 2
F.
LL-37 3
OD600
Hours
50
25
12.5
6.3
3.1
1.6
0.8
0.4
0.2
0.1
Growth
Sterility

**Supplementary Figure 43 – Kinetic antimicrobial susceptibility testing for LL-37. A-C.** *E. coli.* **D-F.** *P. aeruginosa.* Bacterial growth was monitored by OD600 over 18 hours at 37 °C in the presence of the indicated peptide concentrations. Increasing OD600 indicates increasing bacterial growth. Peptide concentrations are given in μM. Each curve indicates an independent experiment.

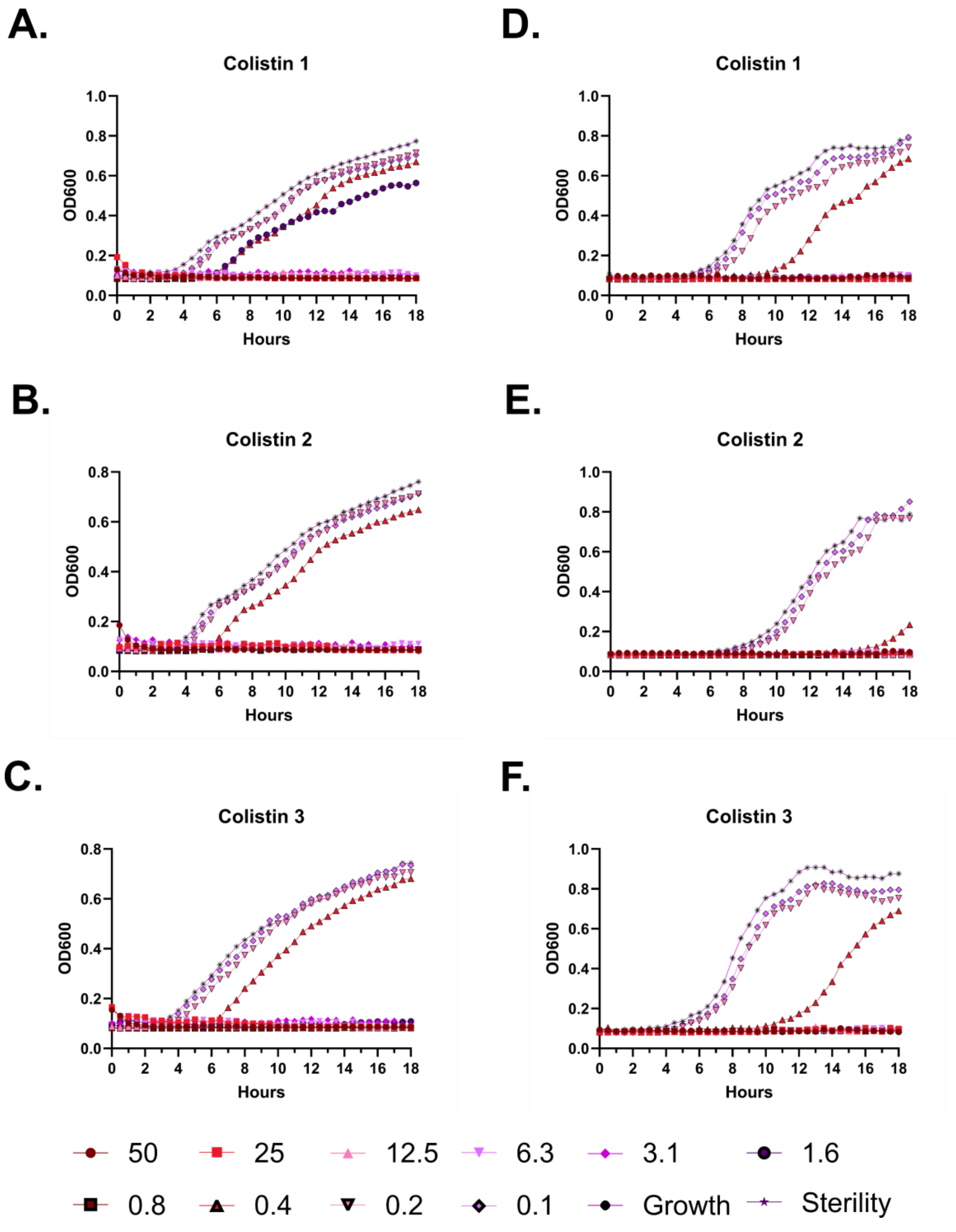

A.
Colistin 1
D.
Colistin 1
B.
Colistin 2
E.
Colistin 2
C.
Colistin 3
F.
Colistin 3
OD600
Hours
50
25
12.5
6.3
3.1
1.6
0.8
0.4
0.2
0.1
Growth
Sterility

**Supplementary Figure 44 – Kinetic antimicrobial susceptibility testing for colistin. A-C.** *E. coli*. **D-F.** *P. aeruginosa.* Bacterial growth was monitored by OD600 over 18 hours at 37 °C in the presence of the indicated peptide concentrations. Increasing OD600 indicates increasing bacterial growth. Peptide concentrations are given in μM. Each curve indicates an independent experiment.

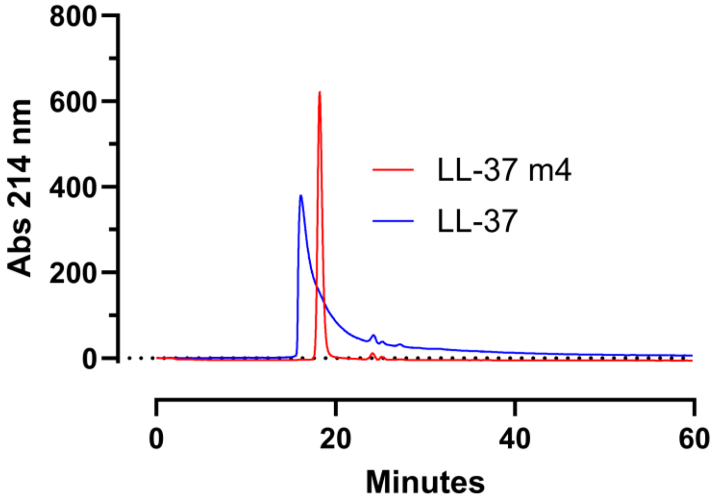

800
600
400
200
0
Abs 214 nm
LL-37 m4
LL-37
0
20
40
60
Minutes

**Figure 45 – Comparison of SEC profiles of LL-37 and m4.** SEC profiles of LL-37 and m4, where the latter demonstrates longer retention and sharper peaks with less tailing. Additional comparisons among mutants are shown in **Figure 3** and in **Supplementary Table 1**.

A.

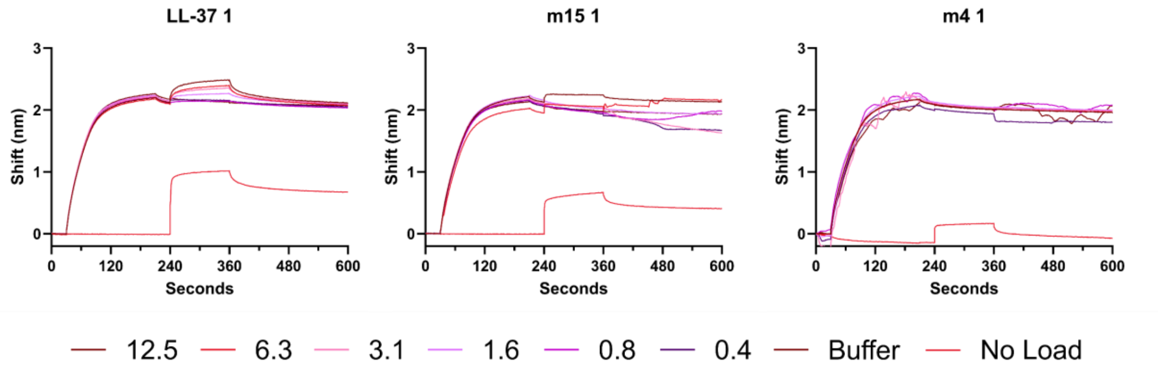
LL-37 1
m15 1
m4 1
Shift (nm)
Seconds
12.5
6.3
3.1
1.6
0.8
0.4
Buffer
No Load

B.

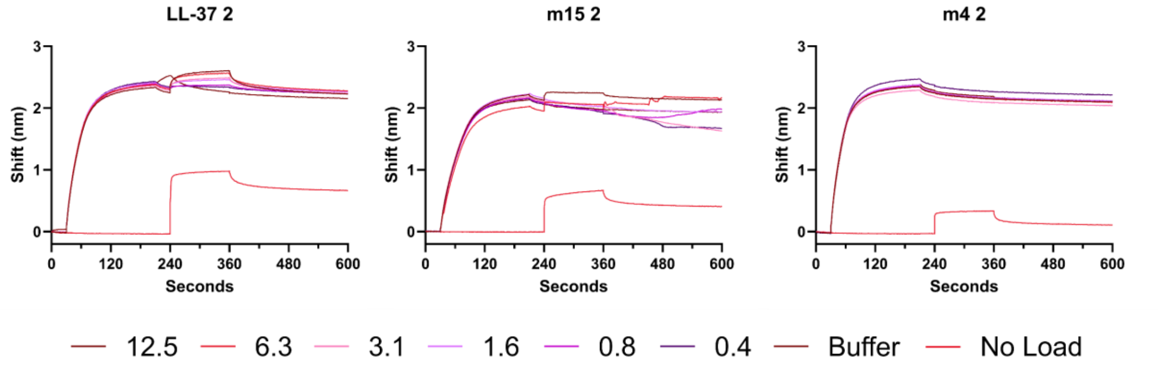
LL-37 2
m15 2
m4 2
Shift (nm)
Seconds
12.5
6.3
3.1
1.6
0.8
0.4
Buffer
No Load

C.

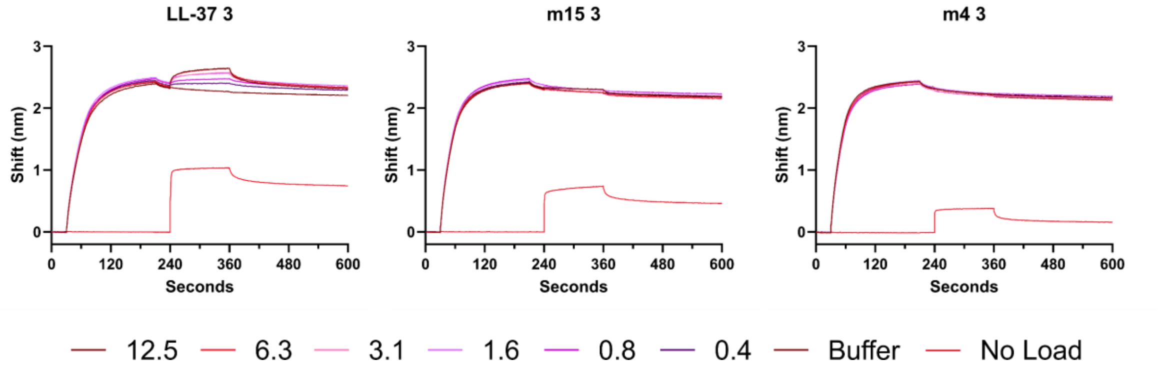
LL-37 3
m15 3
m4 3
Shift (nm)
Seconds
12.5
6.3
3.1
1.6
0.8
0.4
Buffer
No Load

**Supplementary Figure 45 – Raw biolayer interferometry (BLI) traces for interactions between LL-37 probe and LL-37, m15, or m4 analytes. A.** BLI traces for Experiment 1. Buffer indicates a tip loaded with biotinylated LL-37 and then treated with buffer analyte (PBS). No load indicates a probe without any biotinylated LL-37 that is subsequently incubated with the corresponding peptide analyte at 12.5 μM. **B.** BLI traces for Experiment 2. **C.** BLI traces for Experiment 3. Peptide concentrations are given in μM.

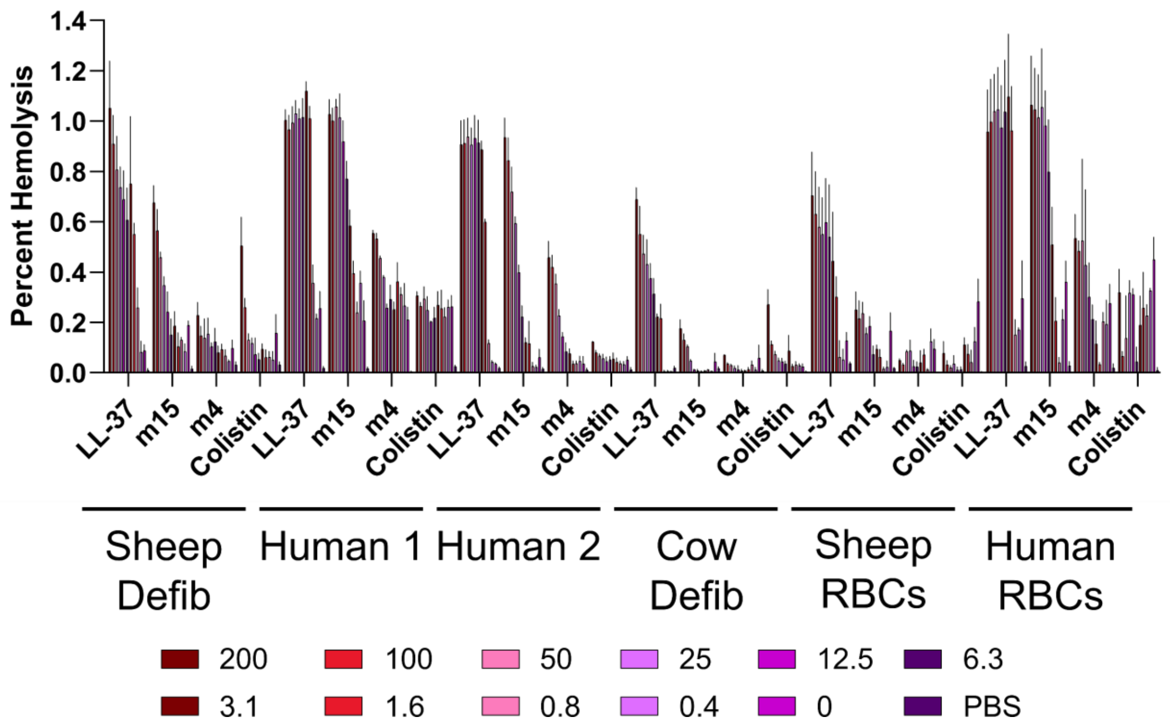
Percent Hemolysis
1.4
1.2
1.0
0.8
0.6
0.4
0.2
0.0
LL-37
m15
m4
Colistin
Sheep Defib
Human 1
Human 2
Cow Defib
Sheep RBCs
Human RBCs
200
100
50
25
12.5
6.3
3.1
1.6
0.8
0.4
0
PBS

**Supplementary Figure 47 – Histogram depiction of Figure 4A data. A.** Endpoint hemolytic activity of each peptide relative to a 100% hemolysis freeze-thaw control after 18 hours at 37 °C. Free hemoglobin levels in cell supernatants were assessed by measuring absorbance at 414 nm.

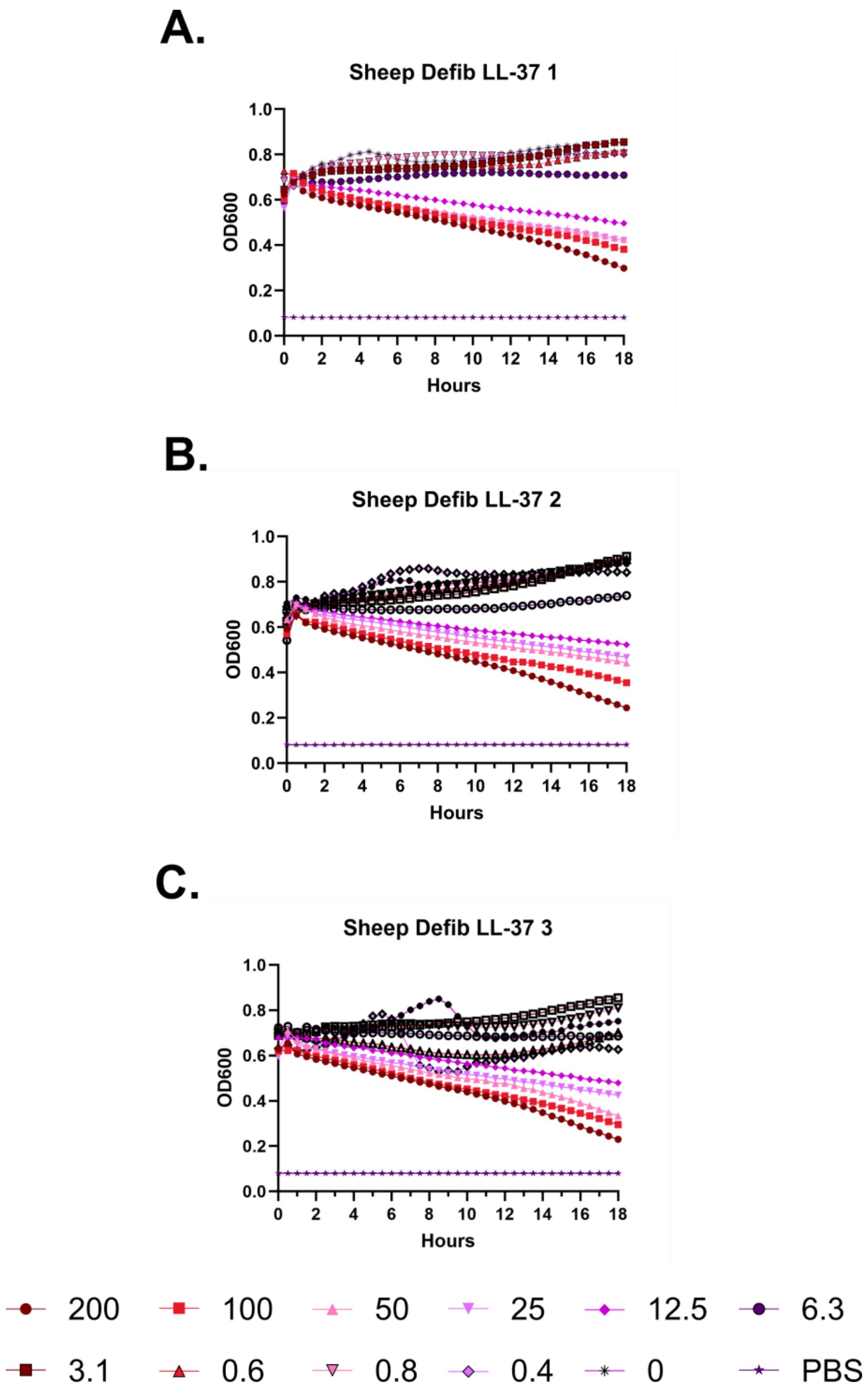
A.
Sheep Defib LL-37 1
OD600
1.0
0.8
0.6
0.4
0.2
0.0
0 2 4 6 8 10 12 14 16 18
Hours
B.
Sheep Defib LL-37 2
OD600
1.0
0.8
0.6
0.4
0.2
0.0
0 2 4 6 8 10 12 14 16 18
Hours
C.
Sheep Defib LL-37 3
OD600
1.0
0.8
0.6
0.4
0.2
0.0
0 2 4 6 8 10 12 14 16 18
Hours
200
100
50
25
12.5
6.3
3.1
0.6
0.8
0.4
0
PBS

**Supplementary Figure 48 – Kinetic hemolysis assays for LL-37 in defibrinated sheep blood. A-C.** Hemolysis was monitored by OD600 over 18 hours at 37 °C in the presence of the indicated peptide concentrations. Decreasing OD600 indicates increasing hemolysis. Peptide concentrations are given in μM. Each curve indicates an independent experiment.

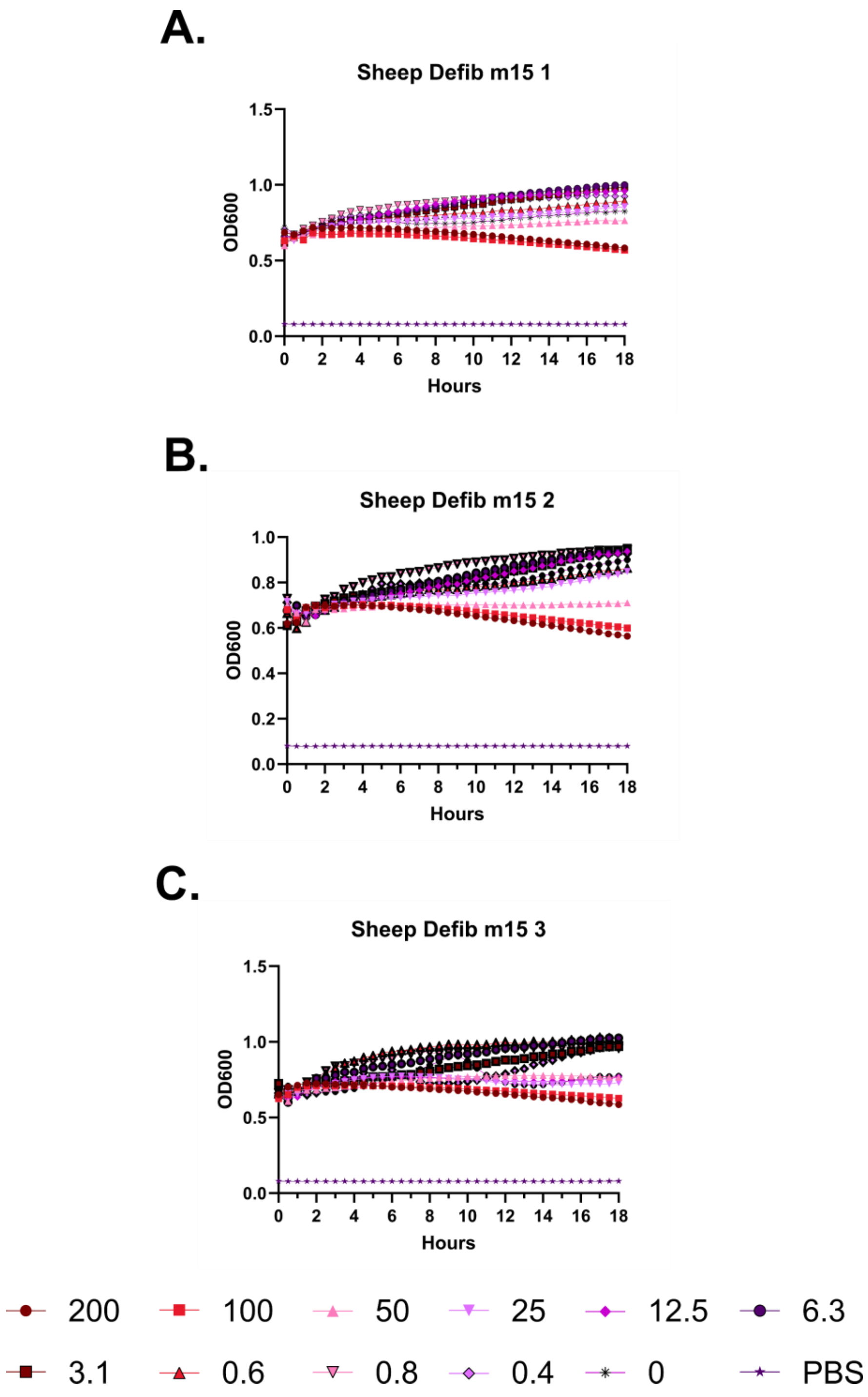
A.
Sheep Defib m15 1
OD600
Hours
B.
Sheep Defib m15 2
OD600
Hours
C.
Sheep Defib m15 3
OD600
Hours
200
100
50
25
12.5
6.3
3.1
0.6
0.8
0.4
0
PBS

**Supplementary Figure 49 – Kinetic hemolysis assays for m15 in defibrinated sheep blood. A-C.** Hemolysis was monitored by OD600 over 18 hours at 37 °C in the presence of the indicated peptide concentrations. Decreasing OD600 indicates increasing hemolysis. Peptide concentrations are given in μM. Each curve indicates an independent experiment.

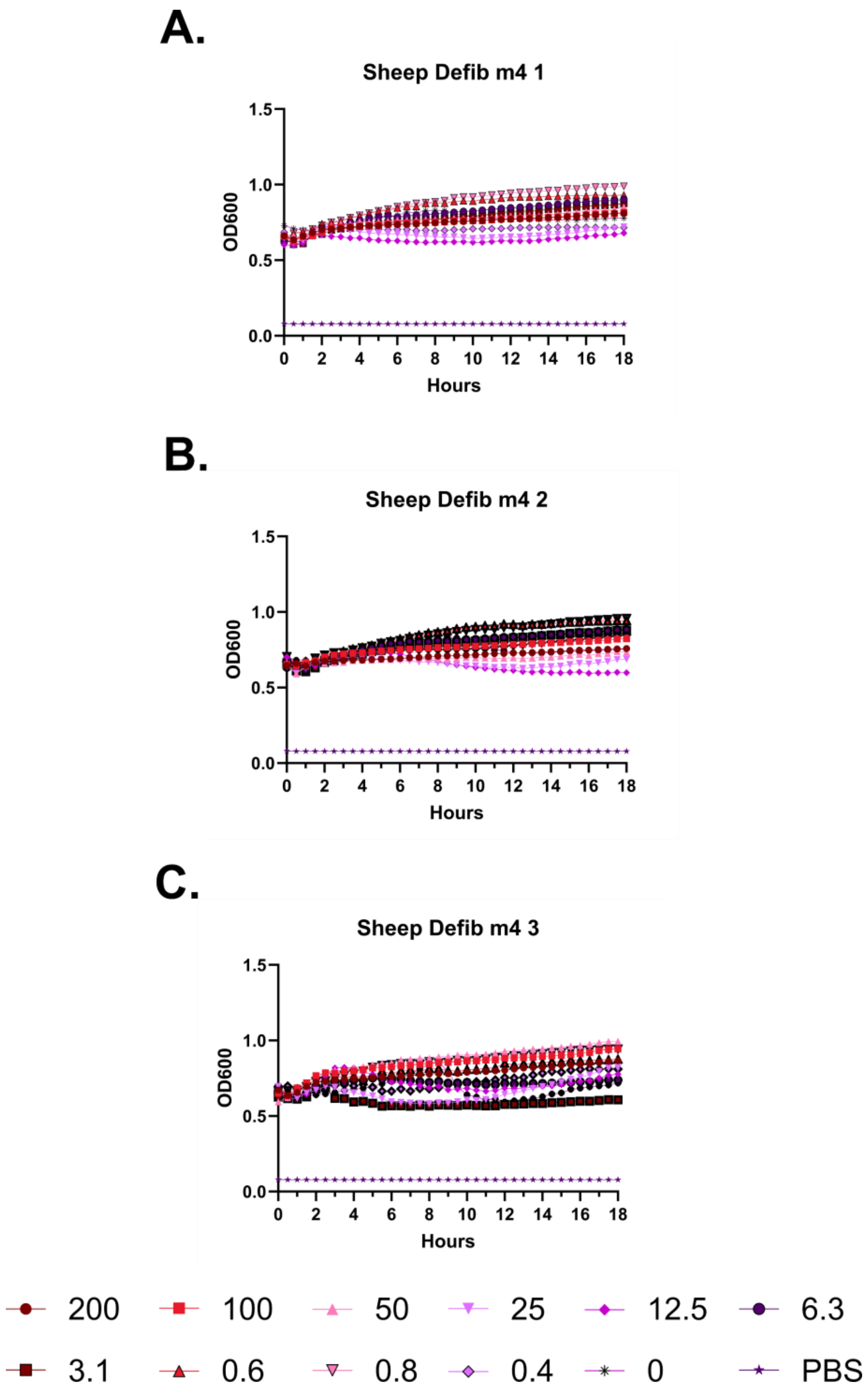
A.
Sheep Defib m4 1
OD600
Hours
B.
Sheep Defib m4 2
OD600
Hours
C.
Sheep Defib m4 3
OD600
Hours
200
100
50
25
12.5
6.3
3.1
0.6
0.8
0.4
0
PBS

**Supplementary Figure 50 – Kinetic hemolysis assays for m4 in defibrinated sheep blood. A-C.** Hemolysis was monitored by OD600 over 18 hours at 37 °C in the presence of the indicated peptide concentrations. Decreasing OD600 indicates increasing hemolysis. Peptide concentrations are given in μM. Each curve indicates an independent experiment.

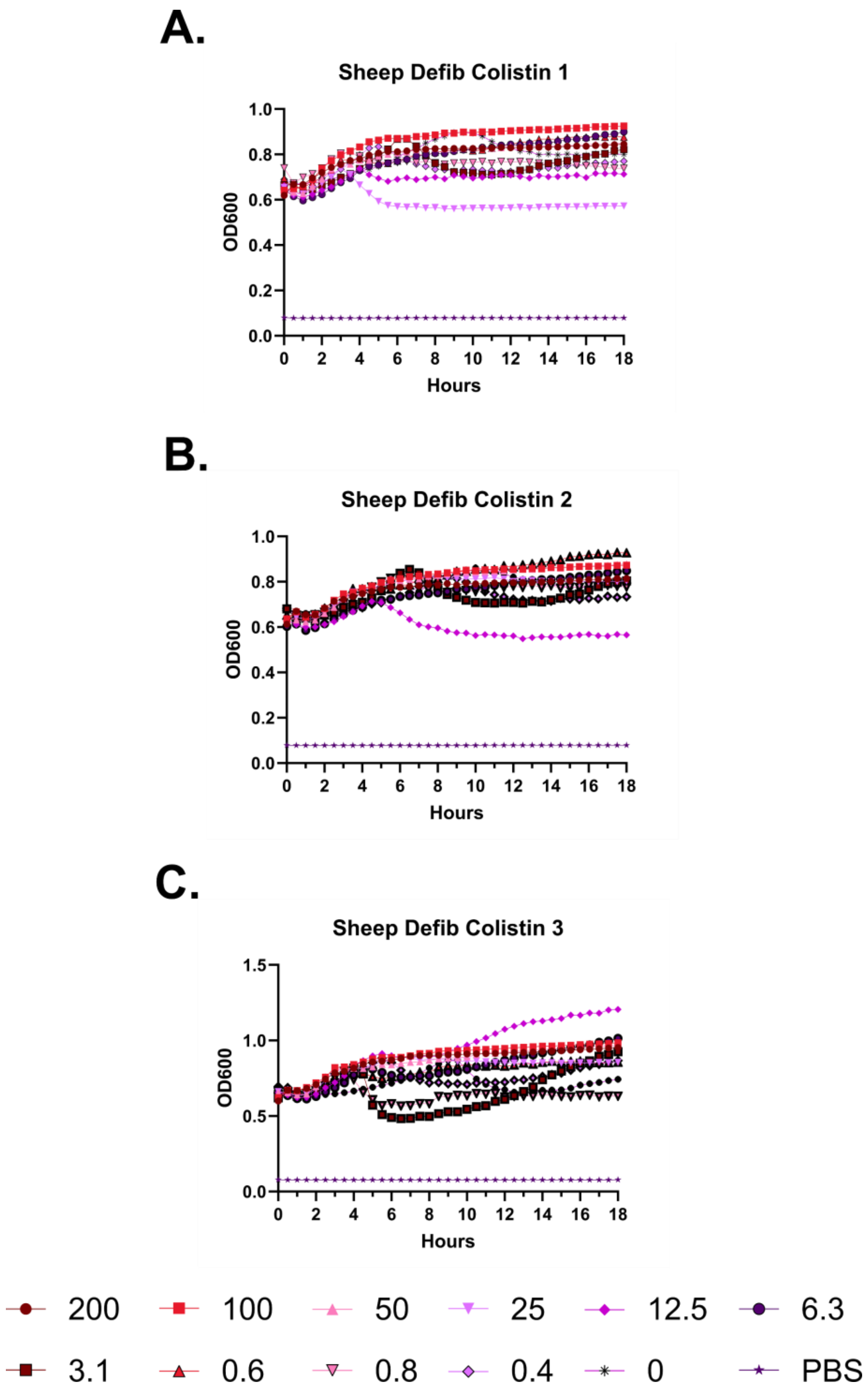
A.
Sheep Defib Colistin 1
OD600
Hours
B.
Sheep Defib Colistin 2
OD600
Hours
C.
Sheep Defib Colistin 3
OD600
Hours
200
100
50
25
12.5
6.3
3.1
0.6
0.8
0.4
0
PBS

**Supplementary Figure 51 – Kinetic hemolysis assays for colistin in defibrinated sheep blood. A-C.** Hemolysis was monitored by OD600 over 18 hours at 37 °C in the presence of the indicated peptide concentrations. Decreasing OD600 indicates increasing hemolysis. Peptide concentrations are given in μM. Each curve indicates an independent experiment.

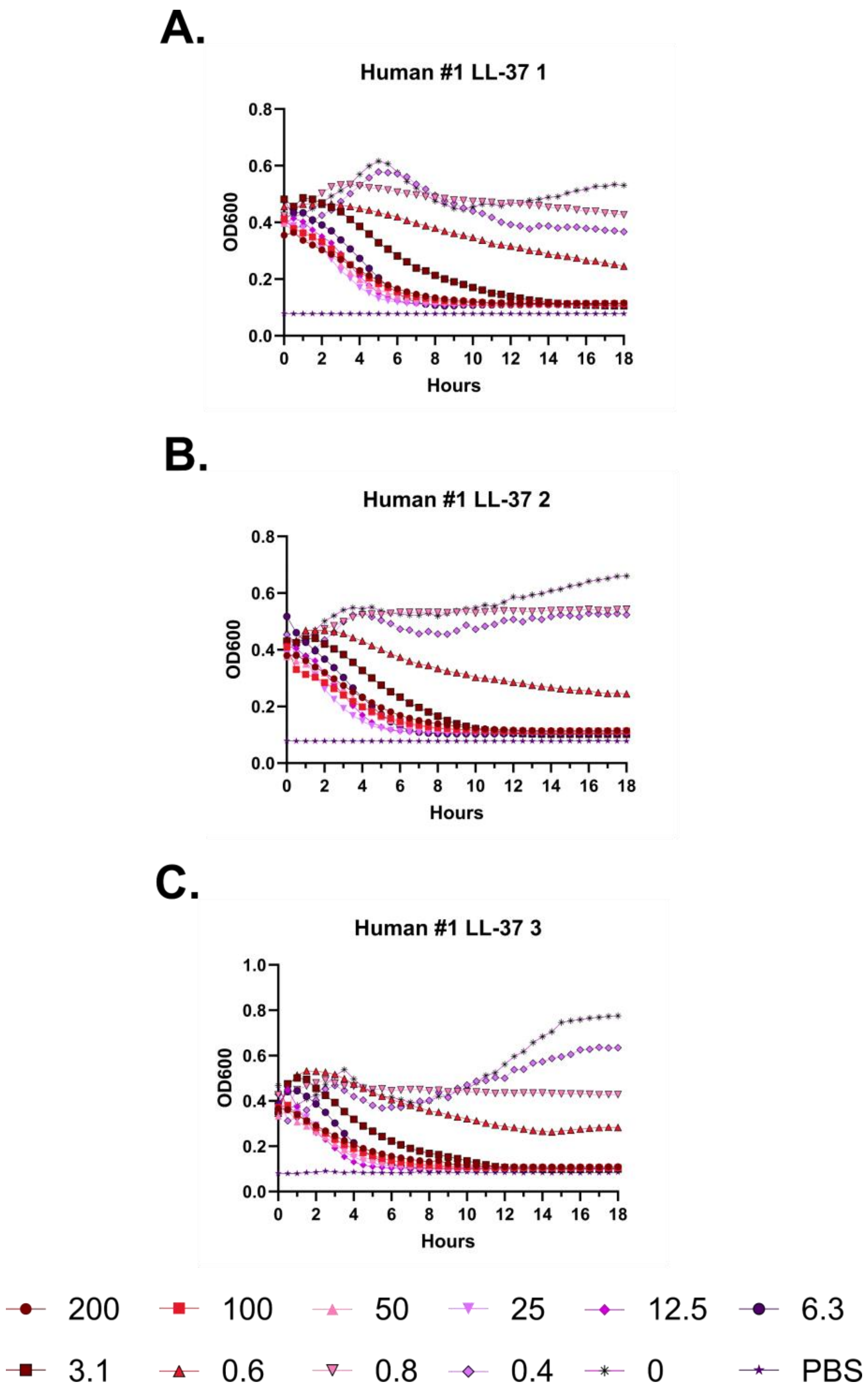
A.
Human #1 LL-37 1
0.8
0.6
0.4
0.2
0.0
OD600
0
2
4
6
8
10
12
14
16
18
Hours
B.
Human #1 LL-37 2
0.8
0.6
0.4
0.2
0.0
OD600
0
2
4
6
8
10
12
14
16
18
Hours
C.
Human #1 LL-37 3
1.0
0.8
0.6
0.4
0.2
0.0
OD600
0
2
4
6
8
10
12
14
16
18
Hours
200
100
50
25
12.5
6.3
3.1
0.6
0.8
0.4
0
PBS

**Supplementary Figure 52 – Kinetic hemolysis assays for LL-37 in human whole blood donor #1. A-C.** Hemolysis was monitored by OD600 over 18 hours at 37 °C in the presence of the indicated peptide concentrations. Decreasing OD600 indicates increasing hemolysis. Peptide concentrations are given in μM. Each curve indicates an independent experiment.

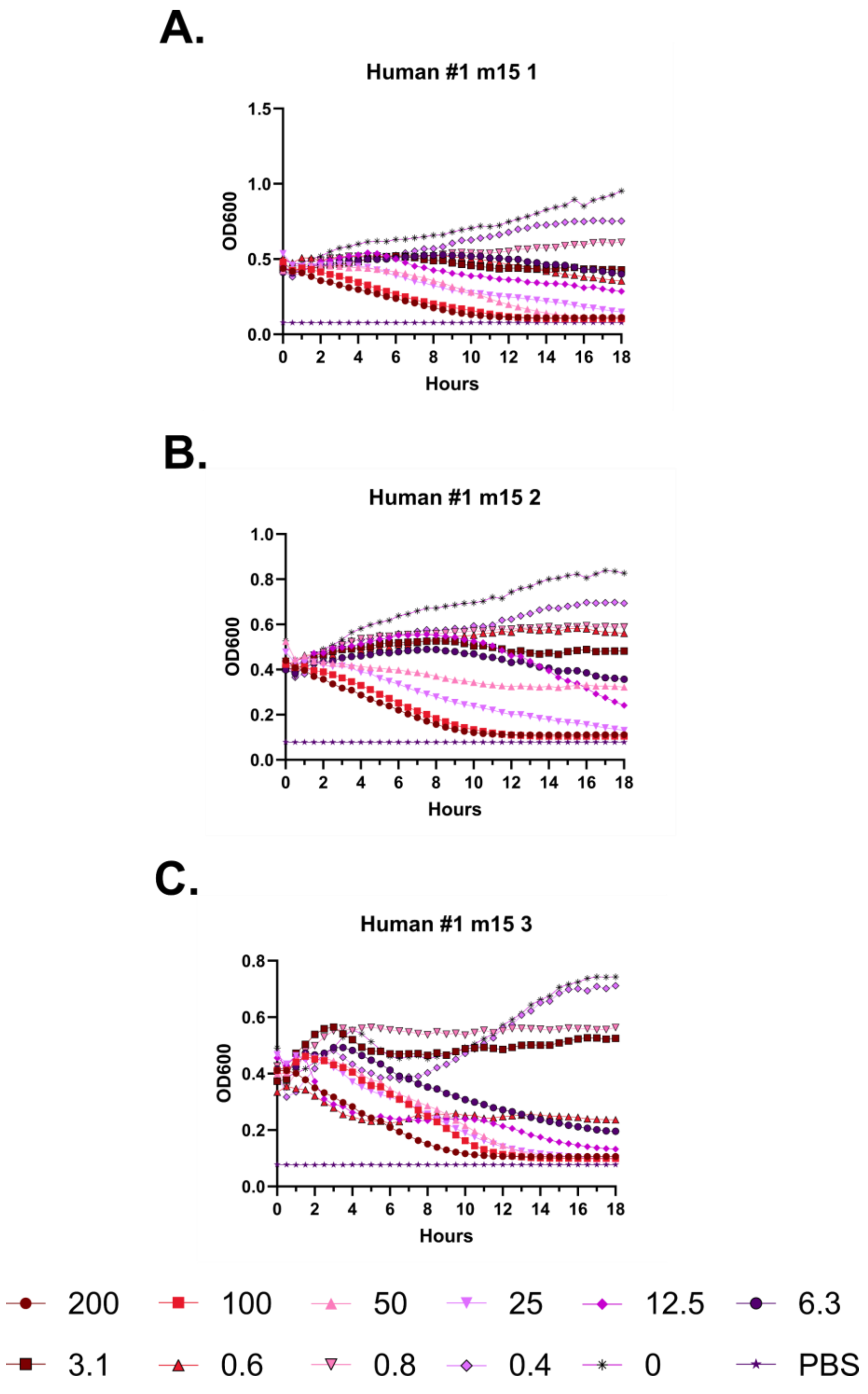
A.
Human #1 m15 1
OD600
Hours
B.
Human #1 m15 2
OD600
Hours
C.
Human #1 m15 3
OD600
Hours
200
100
50
25
12.5
6.3
3.1
0.6
0.8
0.4
0
PBS

**Supplementary Figure 53 – Kinetic hemolysis assays for m15 in human whole blood donor #1.** **A-C.** Hemolysis was monitored by OD600 over 18 hours at 37 °C in the presence of the indicated peptide concentrations. Decreasing OD600 indicates increasing hemolysis. Peptide concentrations are given in μM. Each curve indicates an independent experiment.

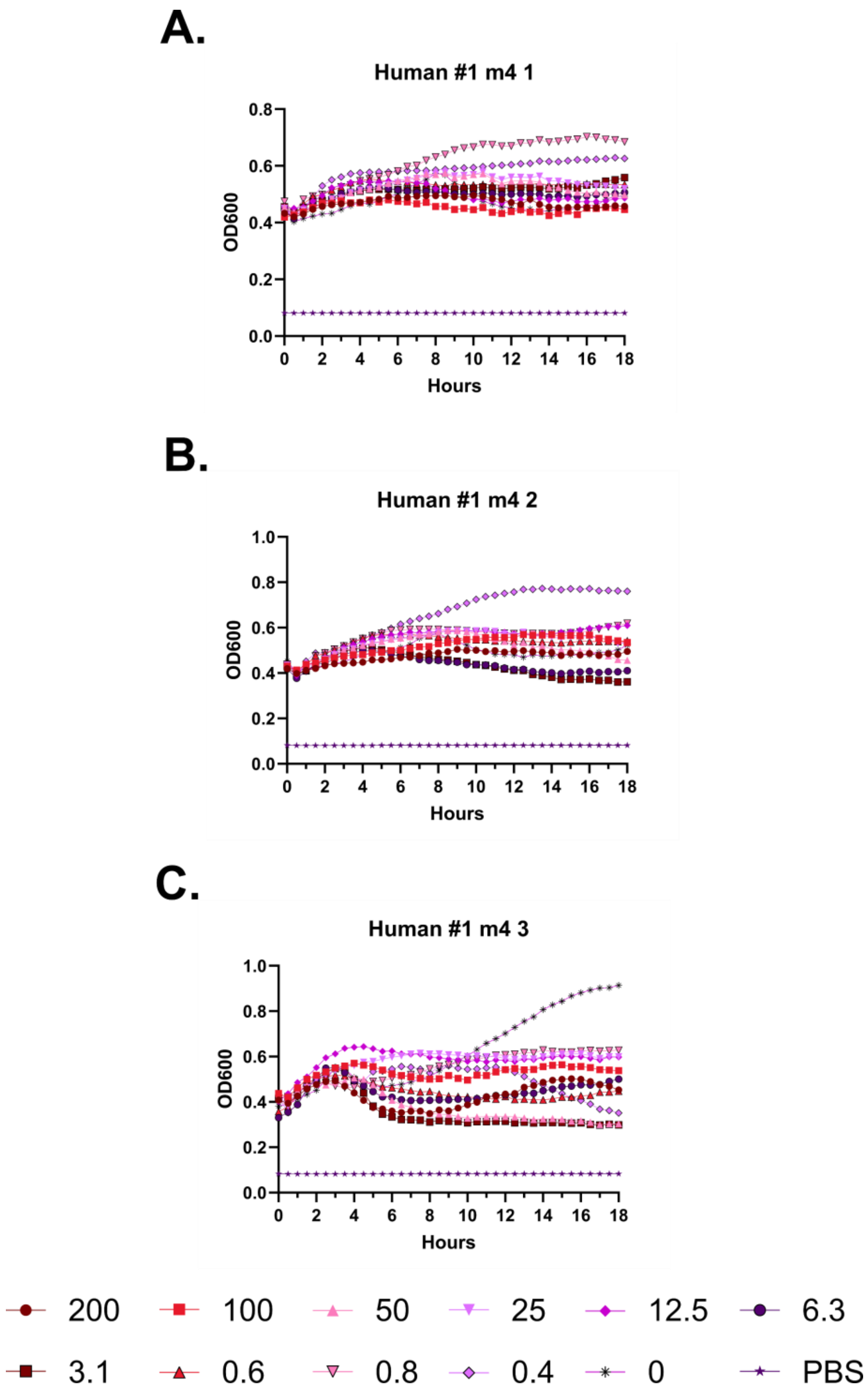
A.
Human #1 m4 1
OD600
Hours
B.
Human #1 m4 2
OD600
Hours
C.
Human #1 m4 3
OD600
Hours
200
100
50
25
12.5
6.3
3.1
0.6
0.8
0.4
0
PBS

**Supplementary Figure 54 – Kinetic hemolysis assays for m4 in human whole blood donor #1. A-C.** Hemolysis was monitored by OD600 over 18 hours at 37 °C in the presence of the indicated peptide concentrations. Decreasing OD600 indicates increasing hemolysis. Peptide concentrations are given in μM. Each curve indicates an independent experiment.

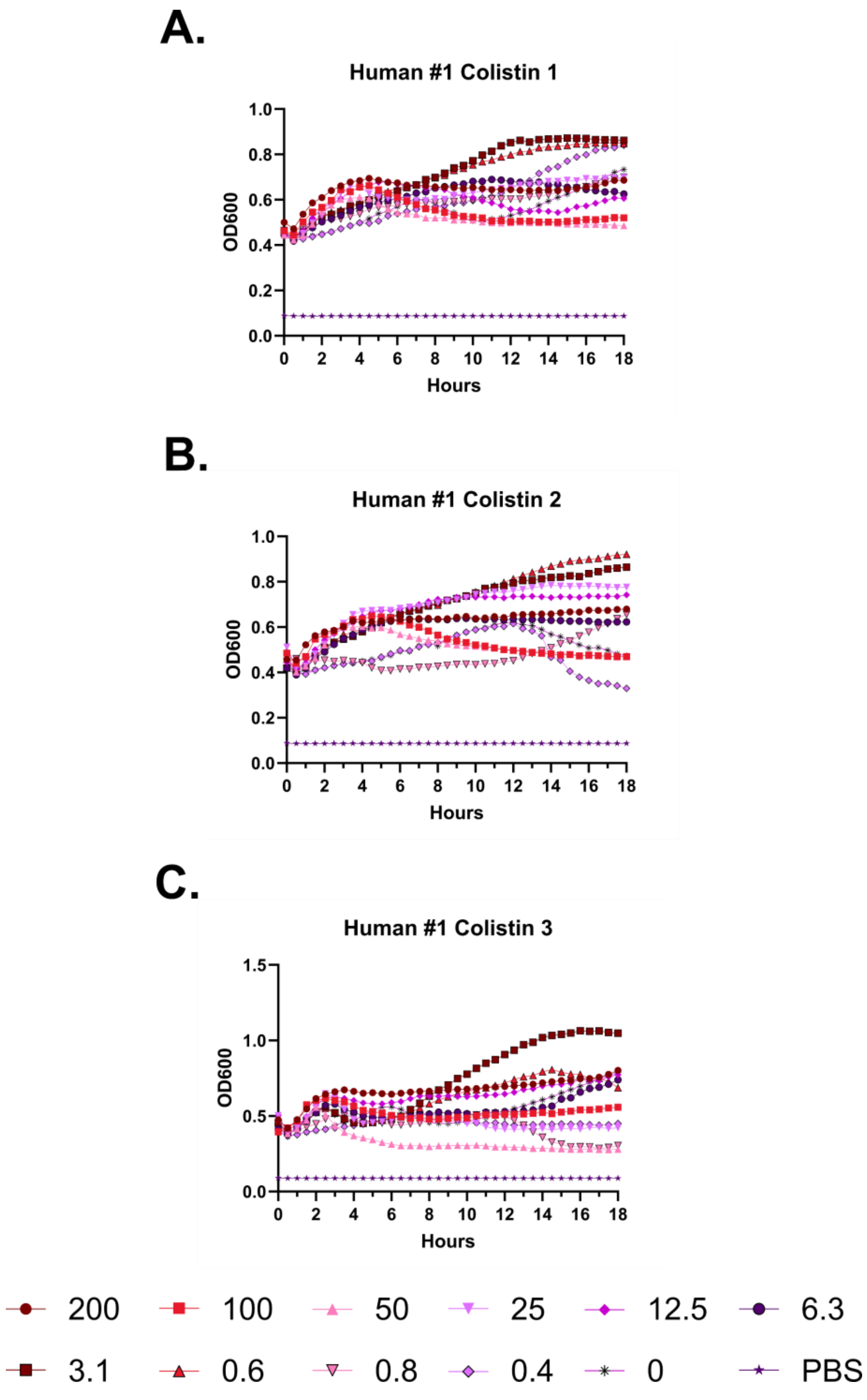
A.
Human #1 Colistin 1
OD600
Hours
B.
Human #1 Colistin 2
OD600
Hours
C.
Human #1 Colistin 3
OD600
Hours
200
100
50
25
12.5
6.3
3.1
0.6
0.8
0.4
0
PBS

**Supplementary Figure 55 – Kinetic hemolysis assays for colistin in human whole blood donor #1. A-C.** Hemolysis was monitored by OD600 over 18 hours at 37 °C in the presence of the indicated peptide concentrations. Decreasing OD600 indicates increasing hemolysis. Peptide concentrations are given in μM. Each curve indicates an independent experiment.

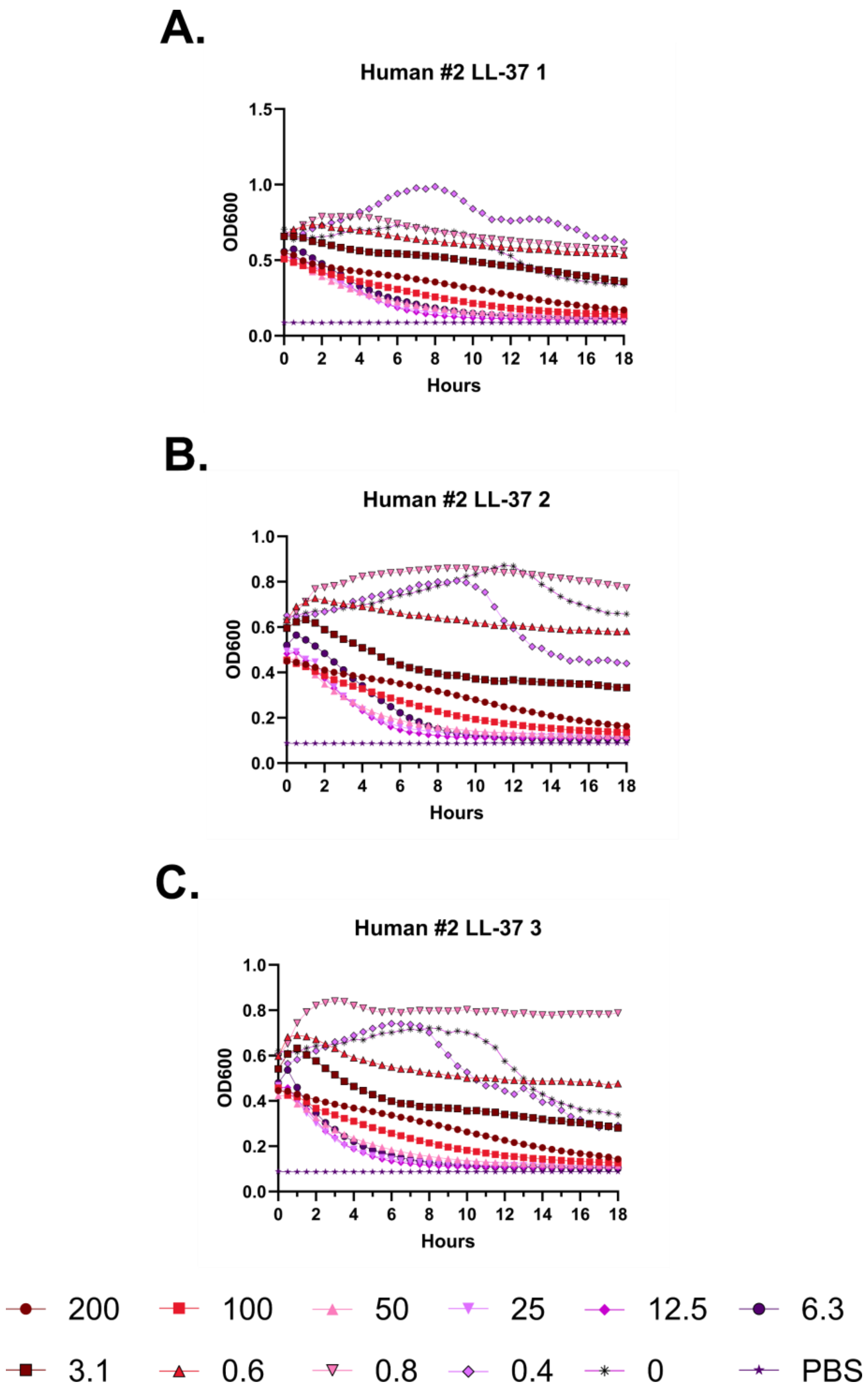
A.
Human #2 LL-37 1
OD600
Hours
B.
Human #2 LL-37 2
OD600
Hours
C.
Human #2 LL-37 3
OD600
Hours
200
100
50
25
12.5
6.3
3.1
0.6
0.8
0.4
0
PBS

**Supplementary Figure 56 – Kinetic hemolysis assays for LL-37 in human whole blood donor #2. A-C.** Hemolysis was monitored by OD600 over 18 hours at 37 °C in the presence of the indicated peptide concentrations. Decreasing OD600 indicates increasing hemolysis. Peptide concentrations are given in μM. Each curve indicates an independent experiment.

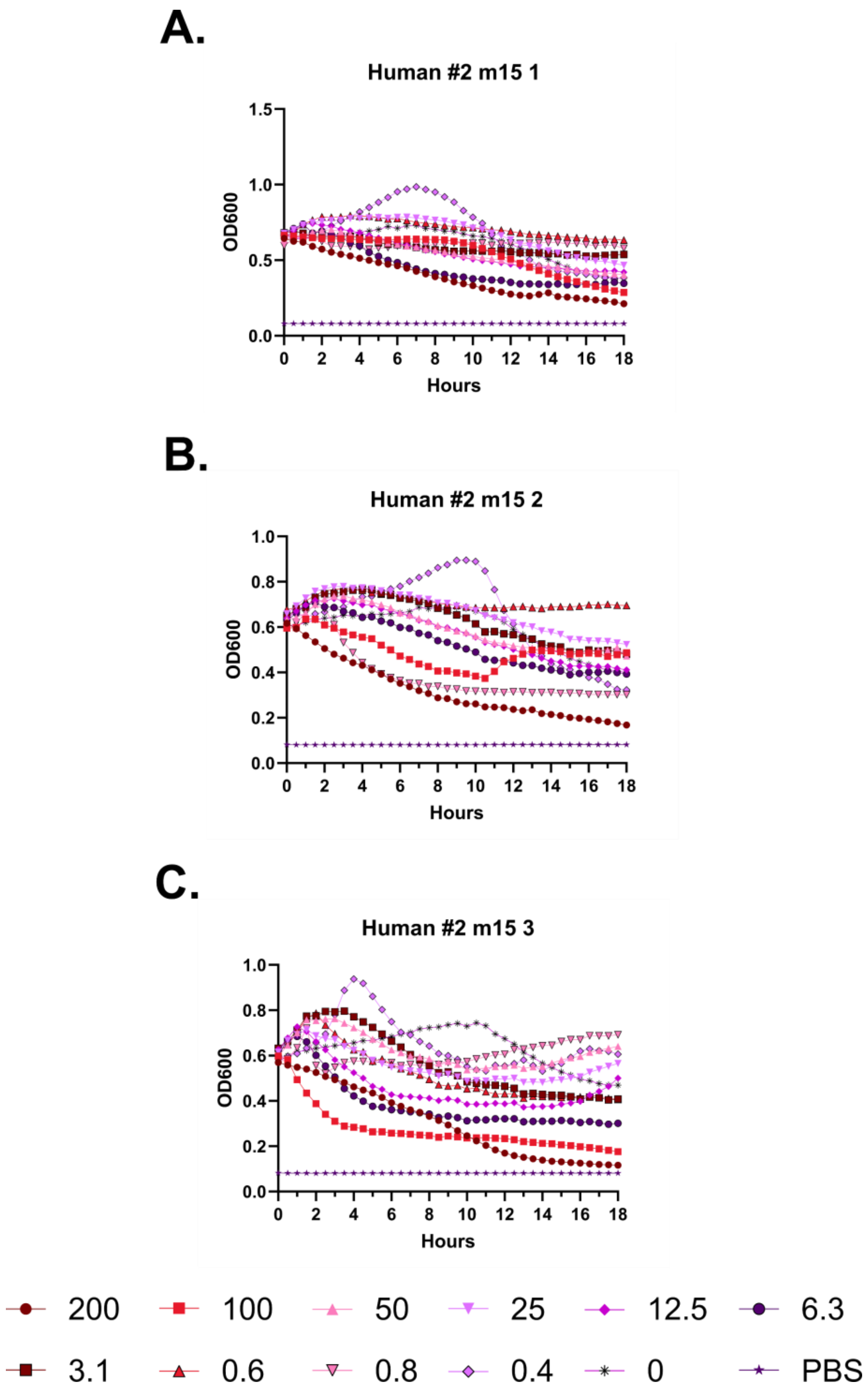
A.
Human #2 m15 1
OD600
Hours
B.
Human #2 m15 2
OD600
Hours
C.
Human #2 m15 3
OD600
Hours
200
100
50
25
12.5
6.3
3.1
0.6
0.8
0.4
0
PBS

**Supplementary Figure 57 – Kinetic hemolysis assays for m15 in human whole blood donor #2. A-C.** Hemolysis was monitored by OD600 over 18 hours at 37 °C in the presence of the indicated peptide concentrations. Decreasing OD600 indicates increasing hemolysis. Peptide concentrations are given in μM. Each curve indicates an independent experiment.

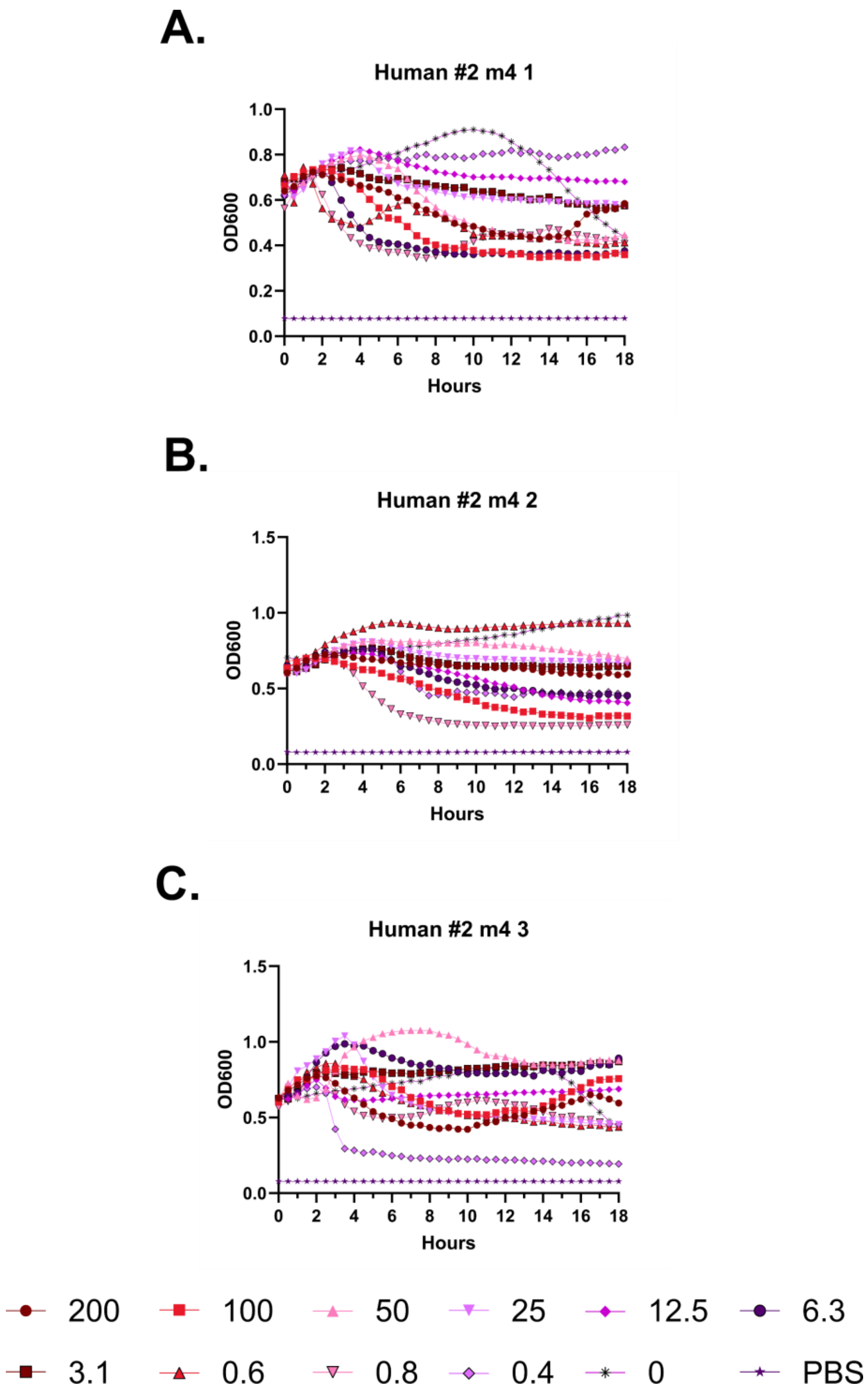
A.
Human #2 m4 1
OD600
Hours
B.
Human #2 m4 2
OD600
Hours
C.
Human #2 m4 3
OD600
Hours
200
100
50
25
12.5
6.3
3.1
0.6
0.8
0.4
0
PBS

**Supplementary Figure 58 – Kinetic hemolysis assays for m4 in human whole blood donor #2. A-C.** Hemolysis was monitored by OD600 over 18 hours at 37 °C in the presence of the indicated peptide concentrations. Decreasing OD600 indicates increasing hemolysis. Peptide concentrations are given in μM. Each curve indicates an independent experiment.

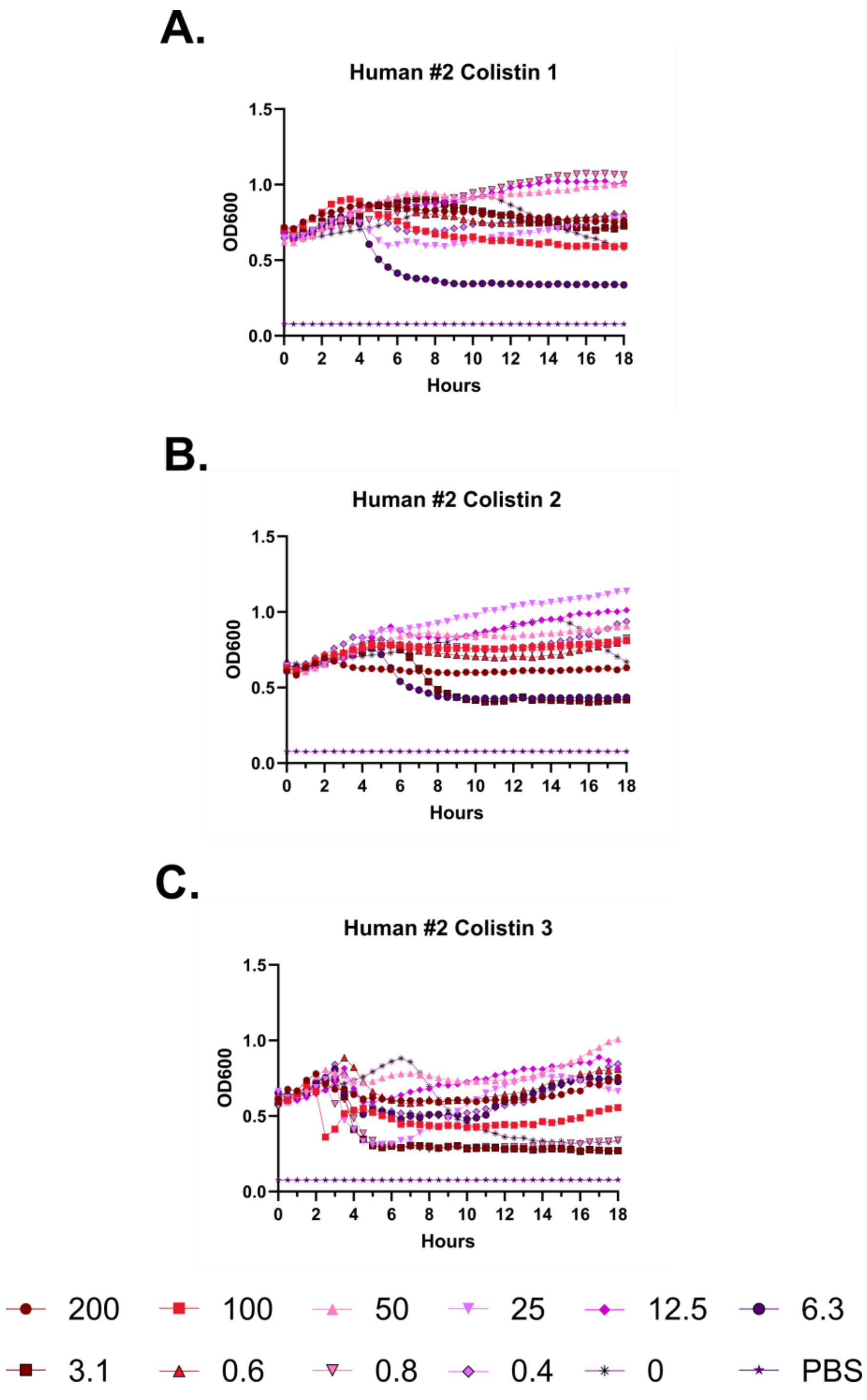
A.
Human #2 Colistin 1
OD600
Hours
B.
Human #2 Colistin 2
OD600
Hours
C.
Human #2 Colistin 3
OD600
Hours
200
100
50
25
12.5
6.3
3.1
0.6
0.8
0.4
0
PBS

**Supplementary Figure 59 – Kinetic hemolysis assays for colistin in human whole blood donor #2. A-C.** Hemolysis was monitored by OD600 over 18 hours at 37 °C in the presence of the indicated peptide concentrations. Decreasing OD600 indicates increasing hemolysis. Peptide concentrations are given in μM. Each curve indicates an independent experiment.

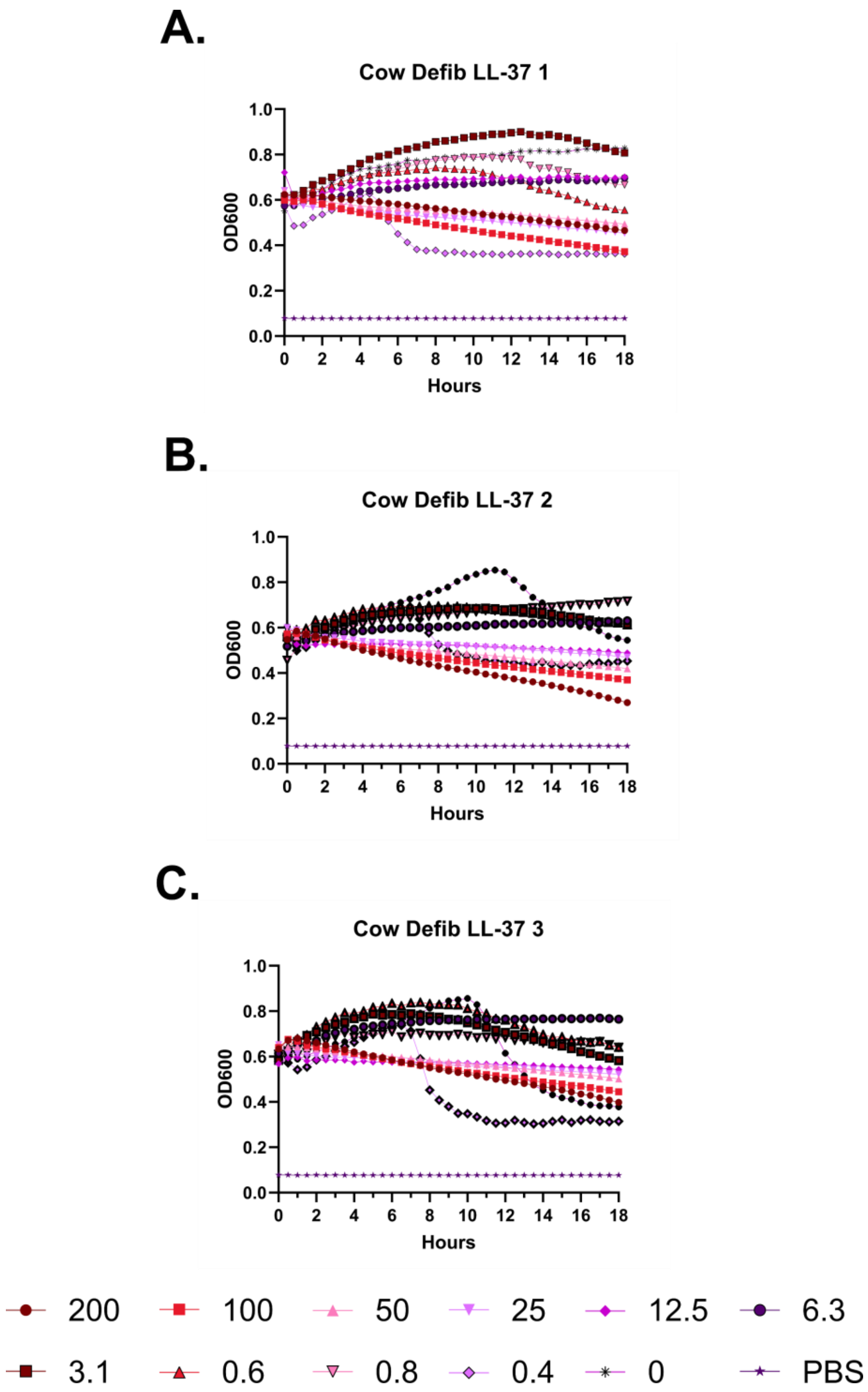
A.
Cow Defib LL-37 1
OD600
Hours
B.
Cow Defib LL-37 2
OD600
Hours
C.
Cow Defib LL-37 3
OD600
Hours
200
100
50
25
12.5
6.3
3.1
0.6
0.8
0.4
0
PBS

**Supplementary Figure 60 – Kinetic hemolysis assays for LL-37 in defibrinated cow blood. A-C.** Hemolysis was monitored by OD600 over 18 hours at 37 °C in the presence of the indicated peptide concentrations. Decreasing OD600 indicates increasing hemolysis. Peptide concentrations are given in μM. Each curve indicates an independent experiment.

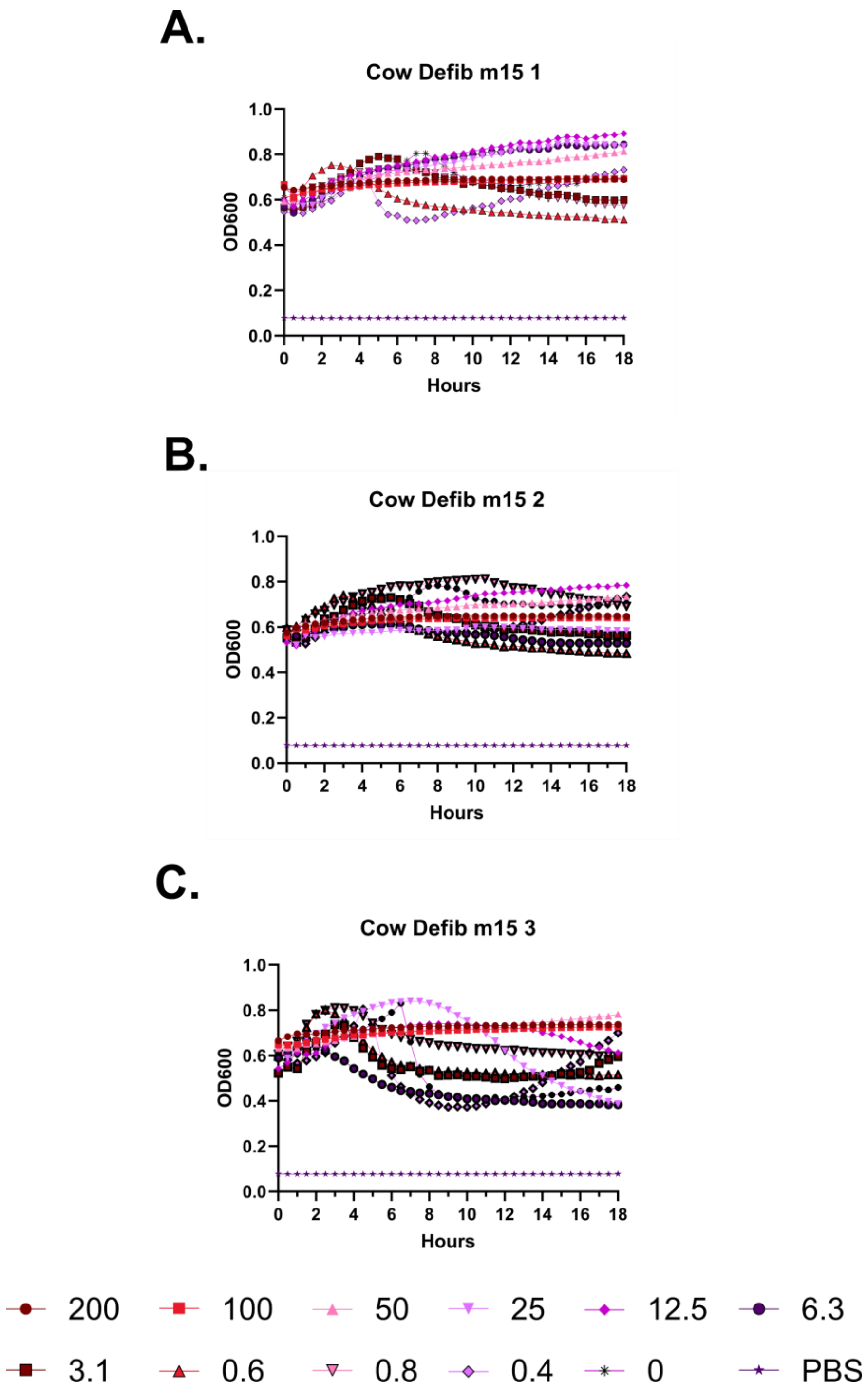
A.
Cow Defib m15 1
B.
Cow Defib m15 2
C.
Cow Defib m15 3
OD600
Hours
0.0
0.2
0.4
0.6
0.8
1.0
0
2
4
6
8
10
12
14
16
18
200
100
50
25
12.5
6.3
3.1
0.6
0.8
0.4
0
PBS

**Supplementary Figure 61 – Kinetic hemolysis assays for m15 in defibrinated cow blood. A-C.** Hemolysis was monitored by OD600 over 18 hours at 37 °C in the presence of the indicated peptide concentrations. Decreasing OD600 indicates increasing hemolysis. Peptide concentrations are given in μM. Each curve indicates an independent experiment.

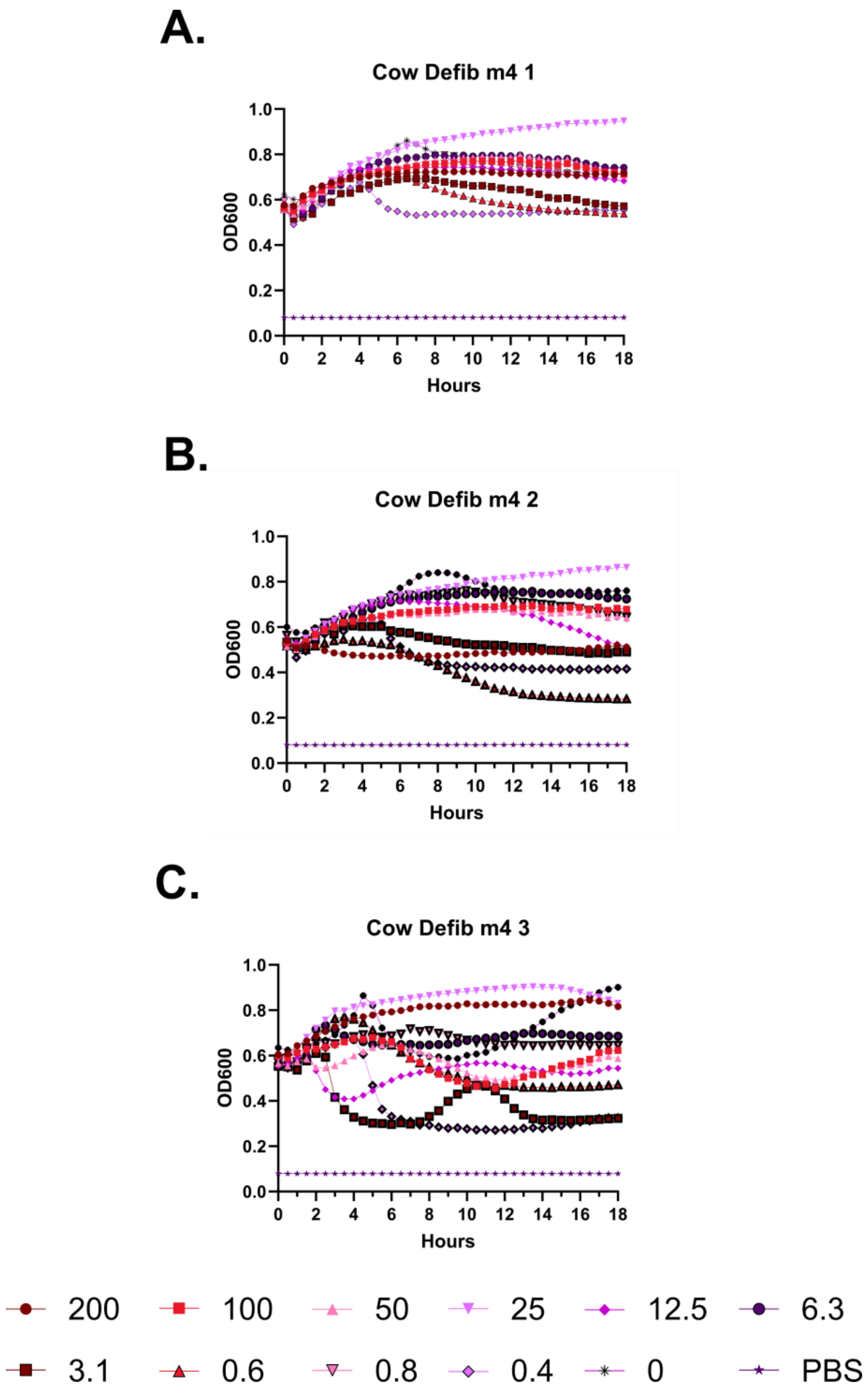
A.
Cow Defib m4 1
OD600
Hours
B.
Cow Defib m4 2
OD600
Hours
C.
Cow Defib m4 3
OD600
Hours
200
100
50
25
12.5
6.3
3.1
0.6
0.8
0.4
0
PBS

**Supplementary Figure 62 – Kinetic hemolysis assays for m4 in defibrinated cow blood. A-C.** Hemolysis was monitored by OD600 over 18 hours at 37 °C in the presence of the indicated peptide concentrations. Decreasing OD600 indicates increasing hemolysis. Peptide concentrations are given in μM. Each curve indicates an independent experiment.

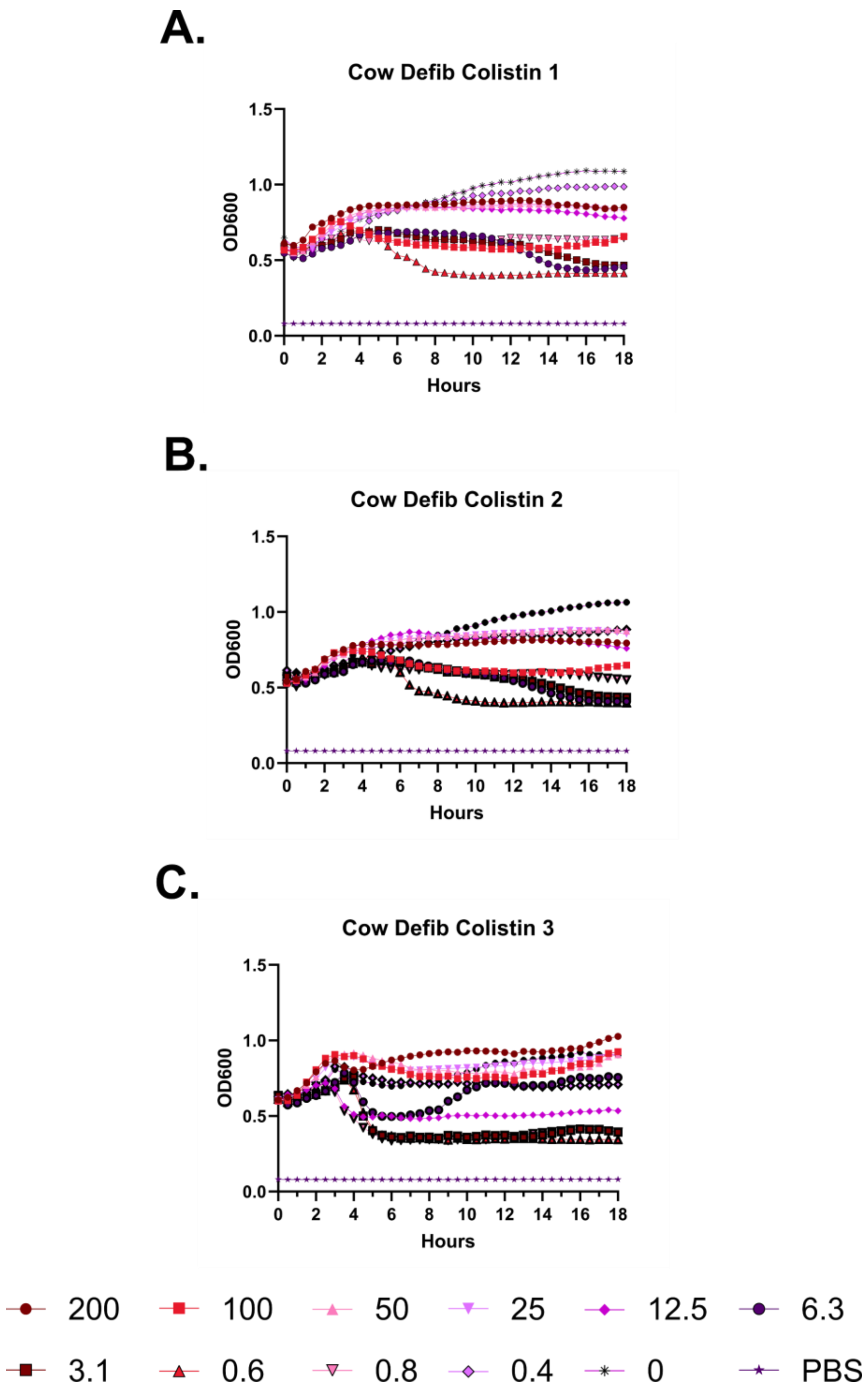
A.
Cow Defib Colistin 1
OD600
Hours
B.
Cow Defib Colistin 2
OD600
Hours
C.
Cow Defib Colistin 3
OD600
Hours
200
100
50
25
12.5
6.3
3.1
0.6
0.8
0.4
0
PBS

**Supplementary Figure 63 – Kinetic hemolysis assays for colistin in defibrinated cow blood. A-C.** Hemolysis was monitored by OD600 over 18 hours at 37 °C in the presence of the indicated peptide concentrations. Decreasing OD600 indicates increasing hemolysis. Peptide concentrations are given in μM. Each curve indicates an independent experiment.

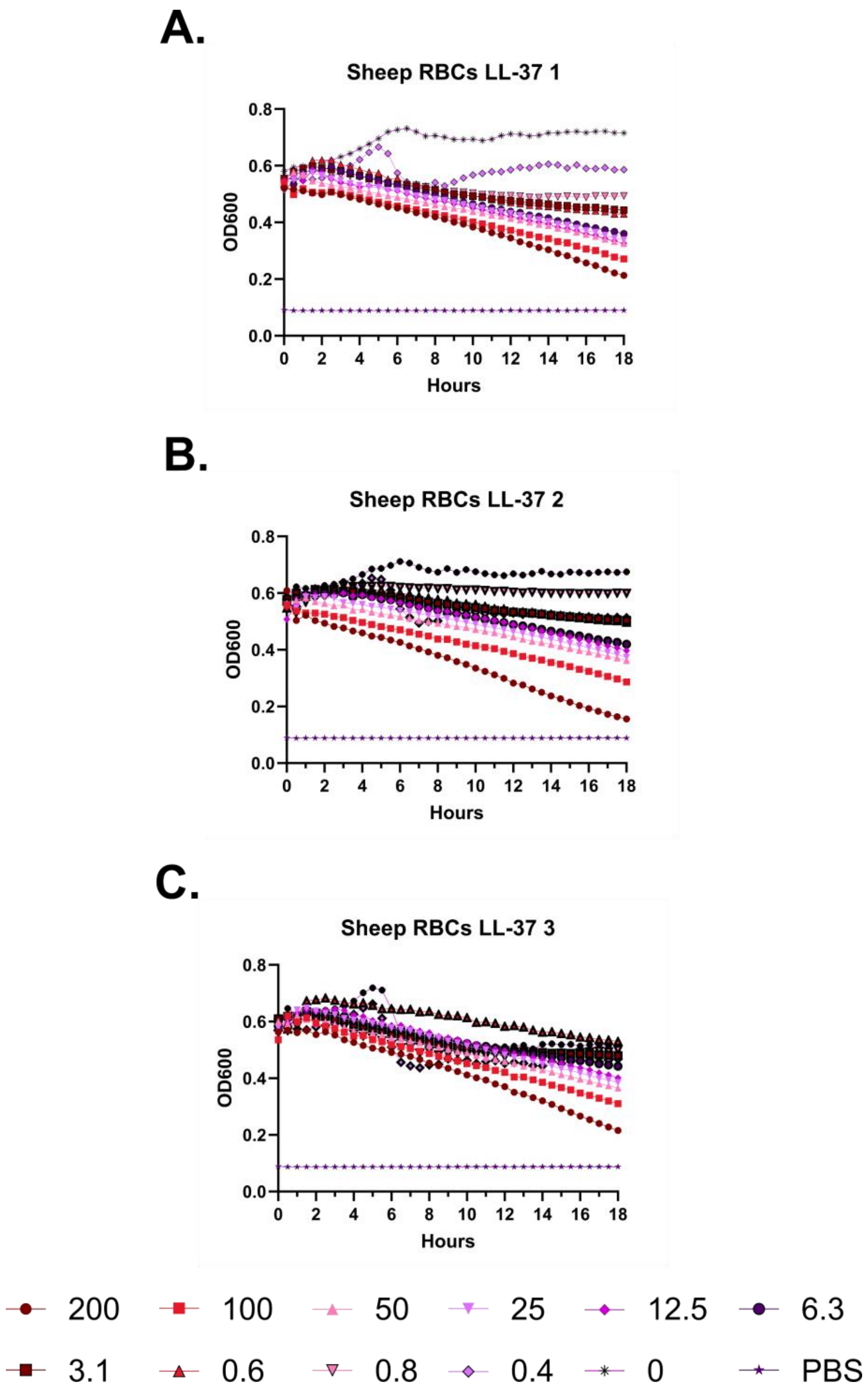
A.
Sheep RBCs LL-37 1
OD600
0.0
0.2
0.4
0.6
0.8
0 2 4 6 8 10 12 14 16 18
Hours
B.
Sheep RBCs LL-37 2
OD600
0.0
0.2
0.4
0.6
0.8
0 2 4 6 8 10 12 14 16 18
Hours
C.
Sheep RBCs LL-37 3
OD600
0.0
0.2
0.4
0.6
0.8
0 2 4 6 8 10 12 14 16 18
Hours
200
100
50
25
12.5
6.3
3.1
0.6
0.8
0.4
0
PBS

**Supplementary Figure 64 – Kinetic hemolysis assays for LL-37 in sheep red blood cells. A-C.** Hemolysis was monitored by OD600 over 18 hours at 37 °C in the presence of the indicated peptide concentrations. Decreasing OD600 indicates increasing hemolysis. Peptide concentrations are given in μM. Each curve indicates an independent experiment.

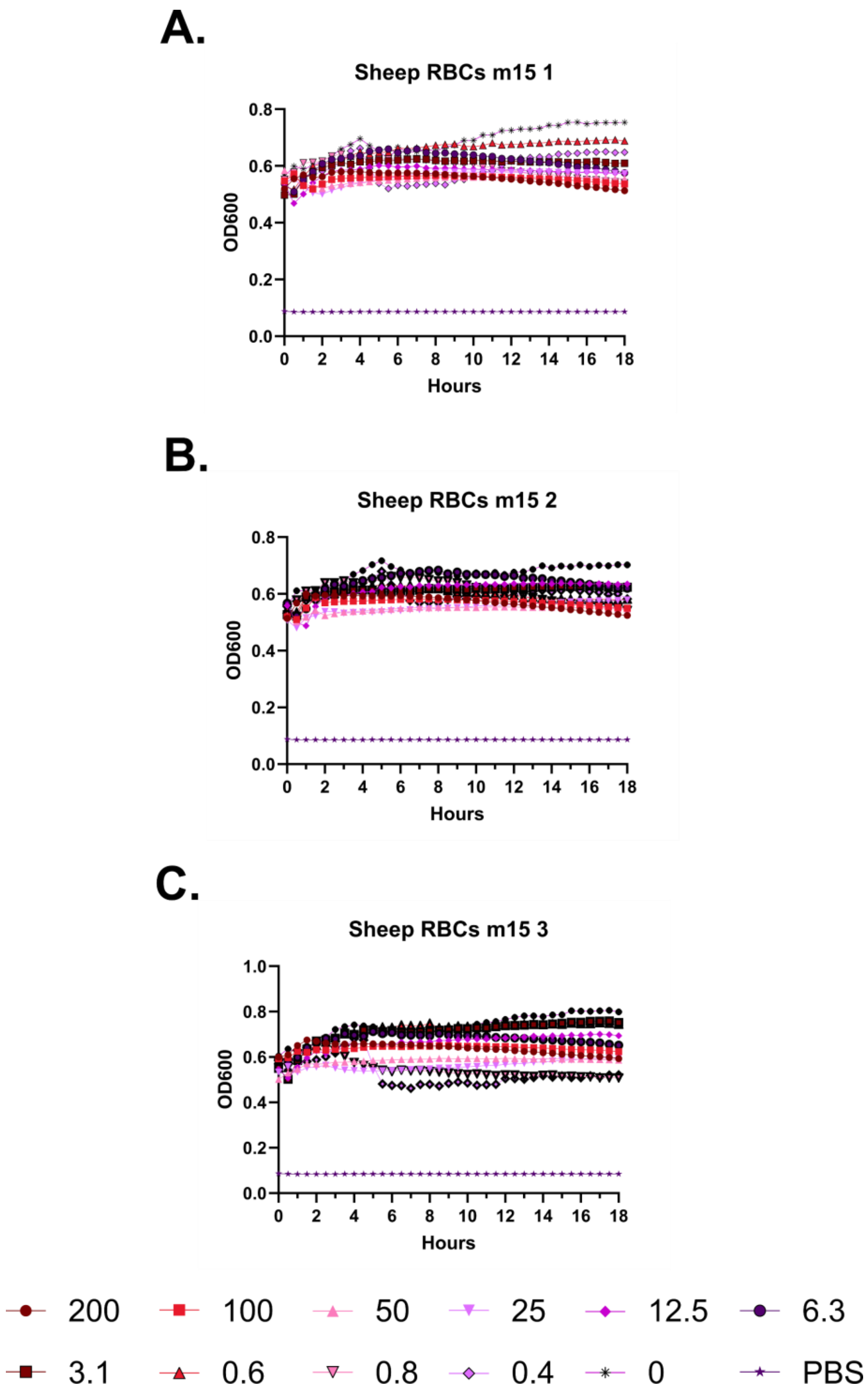
A.
Sheep RBCs m15 1
OD600
Hours
B.
Sheep RBCs m15 2
OD600
Hours
C.
Sheep RBCs m15 3
OD600
Hours
200
100
50
25
12.5
6.3
3.1
0.6
0.8
0.4
0
PBS

**Supplementary Figure 65 – Kinetic hemolysis assays for m15 in sheep red blood cells. A-C.** Hemolysis was monitored by OD600 over 18 hours at 37 °C in the presence of the indicated peptide concentrations. Decreasing OD600 indicates increasing hemolysis. Peptide concentrations are given in μM. Each curve indicates an independent experiment.

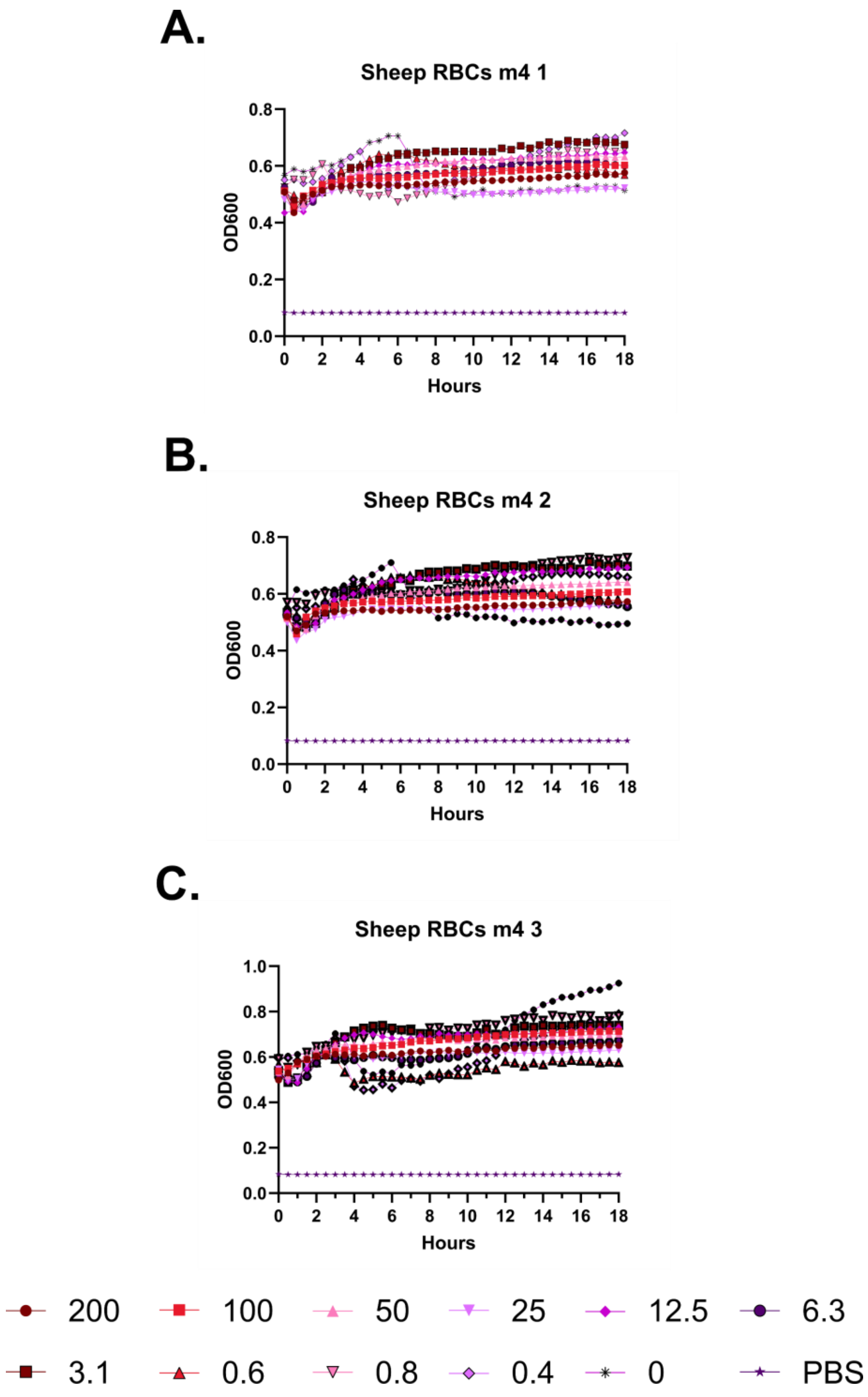
A.
Sheep RBCs m4 1
OD600
Hours
B.
Sheep RBCs m4 2
OD600
Hours
C.
Sheep RBCs m4 3
OD600
Hours
200
100
50
25
12.5
6.3
3.1
0.6
0.8
0.4
0
PBS

**Supplementary Figure 66 – Kinetic hemolysis assays for m4 in sheep red blood cells. A-C.** Hemolysis was monitored by OD600 over 18 hours at 37 °C in the presence of the indicated peptide concentrations. Decreasing OD600 indicates increasing hemolysis. Peptide concentrations are given in μM. Each curve indicates an independent experiment.

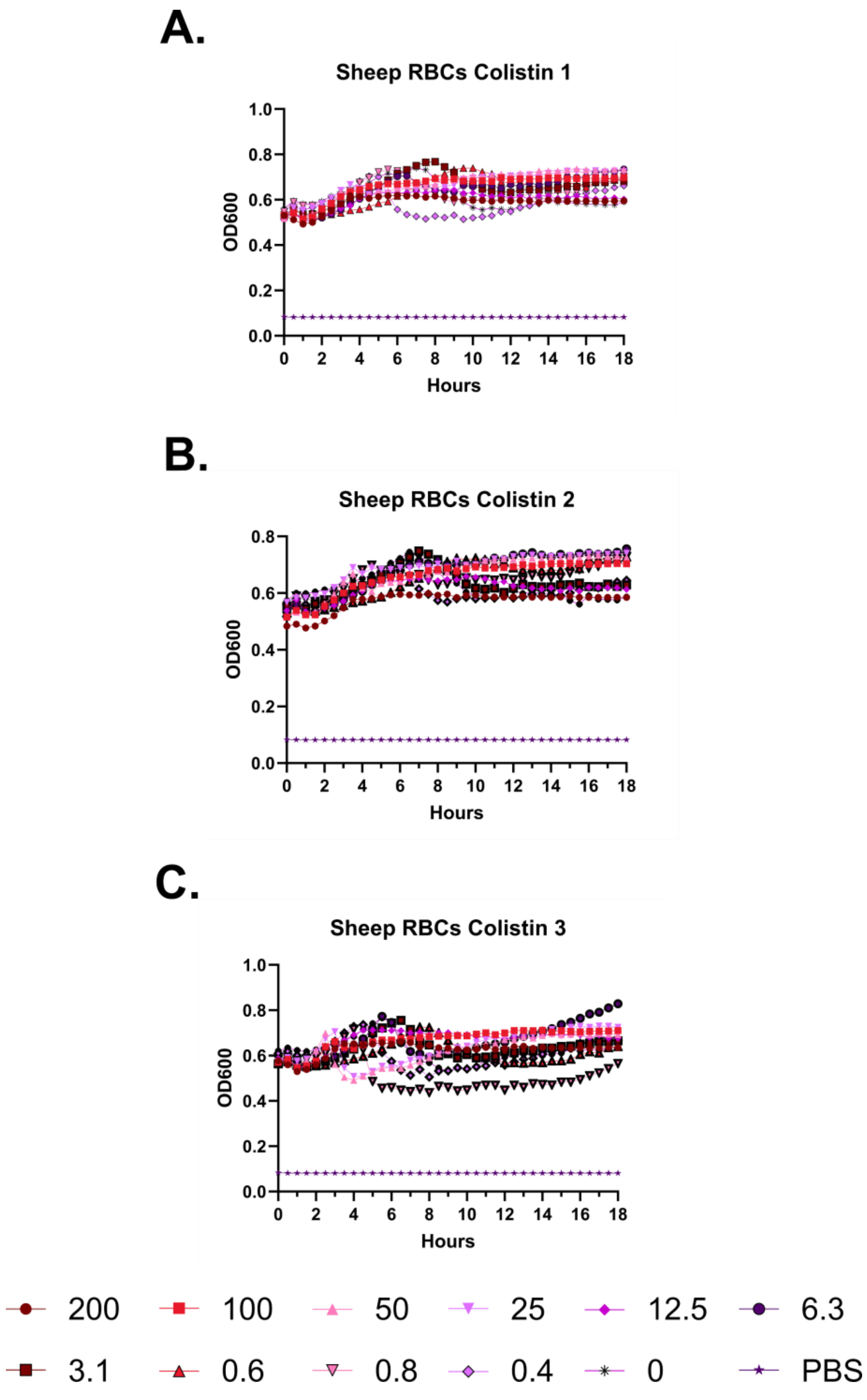
A.
Sheep RBCs Colistin 1
OD600
Hours
B.
Sheep RBCs Colistin 2
OD600
Hours
C.
Sheep RBCs Colistin 3
OD600
Hours
200
100
50
25
12.5
6.3
3.1
0.6
0.8
0.4
0
PBS

**Supplementary Figure 67 – Kinetic hemolysis assays for colistin in sheep red blood cells. A-C.** Hemolysis was monitored by OD600 over 18 hours at 37 °C in the presence of the indicated peptide concentrations. Decreasing OD600 indicates increasing hemolysis. Peptide concentrations are given in μM. Each curve indicates an independent experiment.

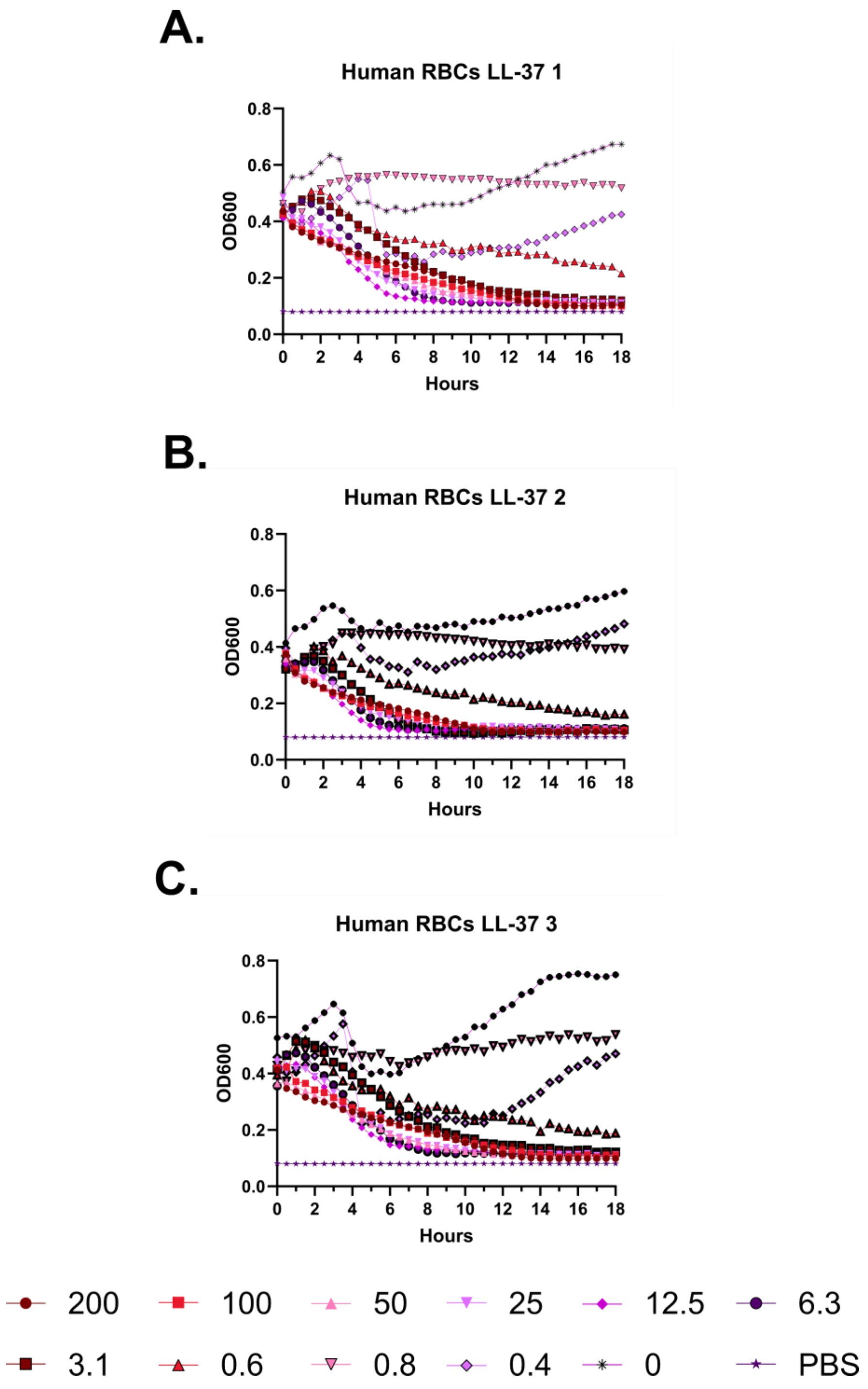
A.
Human RBCs LL-37 1
OD600
0.0
0.2
0.4
0.6
0.8
0 2 4 6 8 10 12 14 16 18
Hours
B.
Human RBCs LL-37 2
OD600
0.0
0.2
0.4
0.6
0.8
0 2 4 6 8 10 12 14 16 18
Hours
C.
Human RBCs LL-37 3
OD600
0.0
0.2
0.4
0.6
0.8
0 2 4 6 8 10 12 14 16 18
Hours
200
100
50
25
12.5
6.3
3.1
0.6
0.8
0.4
0
PBS

**Supplementary Figure 68 – Kinetic hemolysis assays for LL-37 in human red blood cells. A-C.** Hemolysis was monitored by OD600 over 18 hours at 37 °C in the presence of the indicated peptide concentrations. Decreasing OD600 indicates increasing hemolysis. Peptide concentrations are given in μM. Each curve indicates an independent experiment.

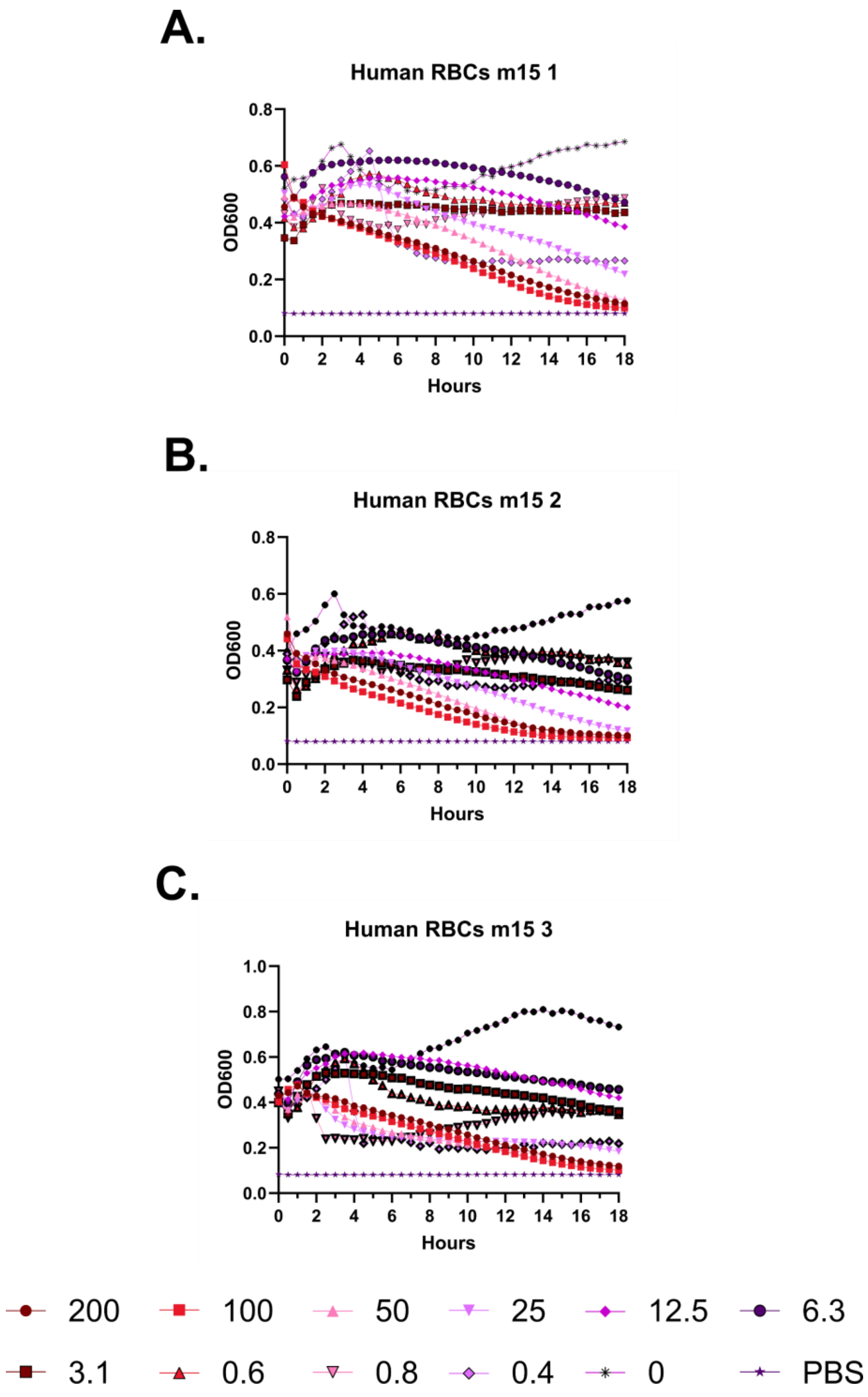
A.
Human RBCs m15 1
OD600
Hours
0.0
0.2
0.4
0.6
0.8
0 2 4 6 8 10 12 14 16 18
B.
Human RBCs m15 2
OD600
Hours
0.0
0.2
0.4
0.6
0.8
0 2 4 6 8 10 12 14 16 18
C.
Human RBCs m15 3
OD600
Hours
0.0
0.2
0.4
0.6
0.8
1.0
0 2 4 6 8 10 12 14 16 18
200
100
50
25
12.5
6.3
3.1
0.6
0.8
0.4
0
PBS

**Supplementary Figure 69 – Kinetic hemolysis assays for m15 in human red blood cells. A-C.** Hemolysis was monitored by OD600 over 18 hours at 37 °C in the presence of the indicated peptide concentrations. Decreasing OD600 indicates increasing hemolysis. Peptide concentrations are given in μM. Each curve indicates an independent experiment.

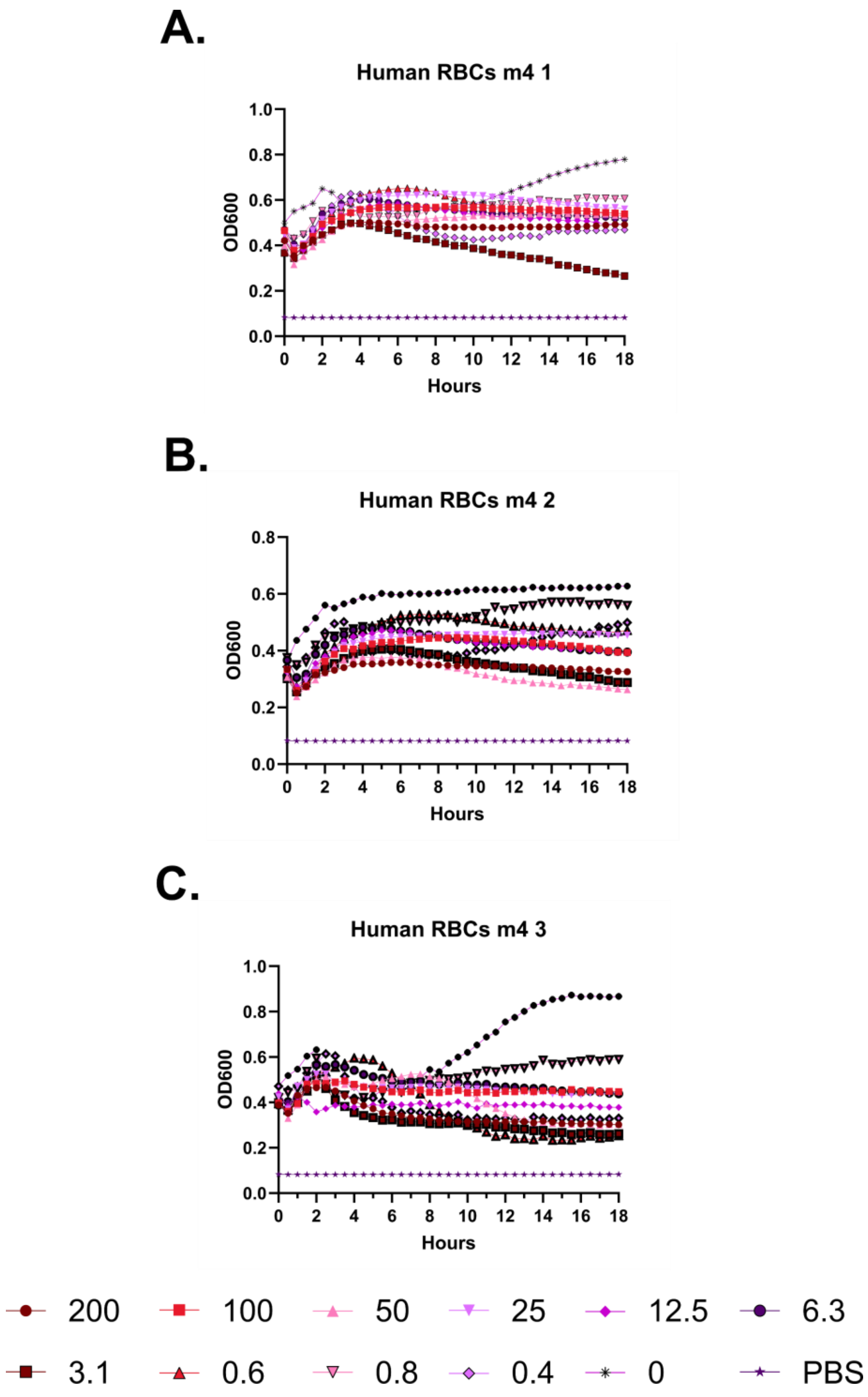
A.
Human RBCs m4 1
OD600
Hours
B.
Human RBCs m4 2
OD600
Hours
C.
Human RBCs m4 3
OD600
Hours
200
100
50
25
12.5
6.3
3.1
0.6
0.8
0.4
0
PBS

**Supplementary Figure 70 – Kinetic hemolysis assays for m4 in human red blood cells. A-C.** Hemolysis was monitored by OD600 over 18 hours at 37 °C in the presence of the indicated peptide concentrations. Decreasing OD600 indicates increasing hemolysis. Peptide concentrations are given in μM. Each curve indicates an independent experiment.

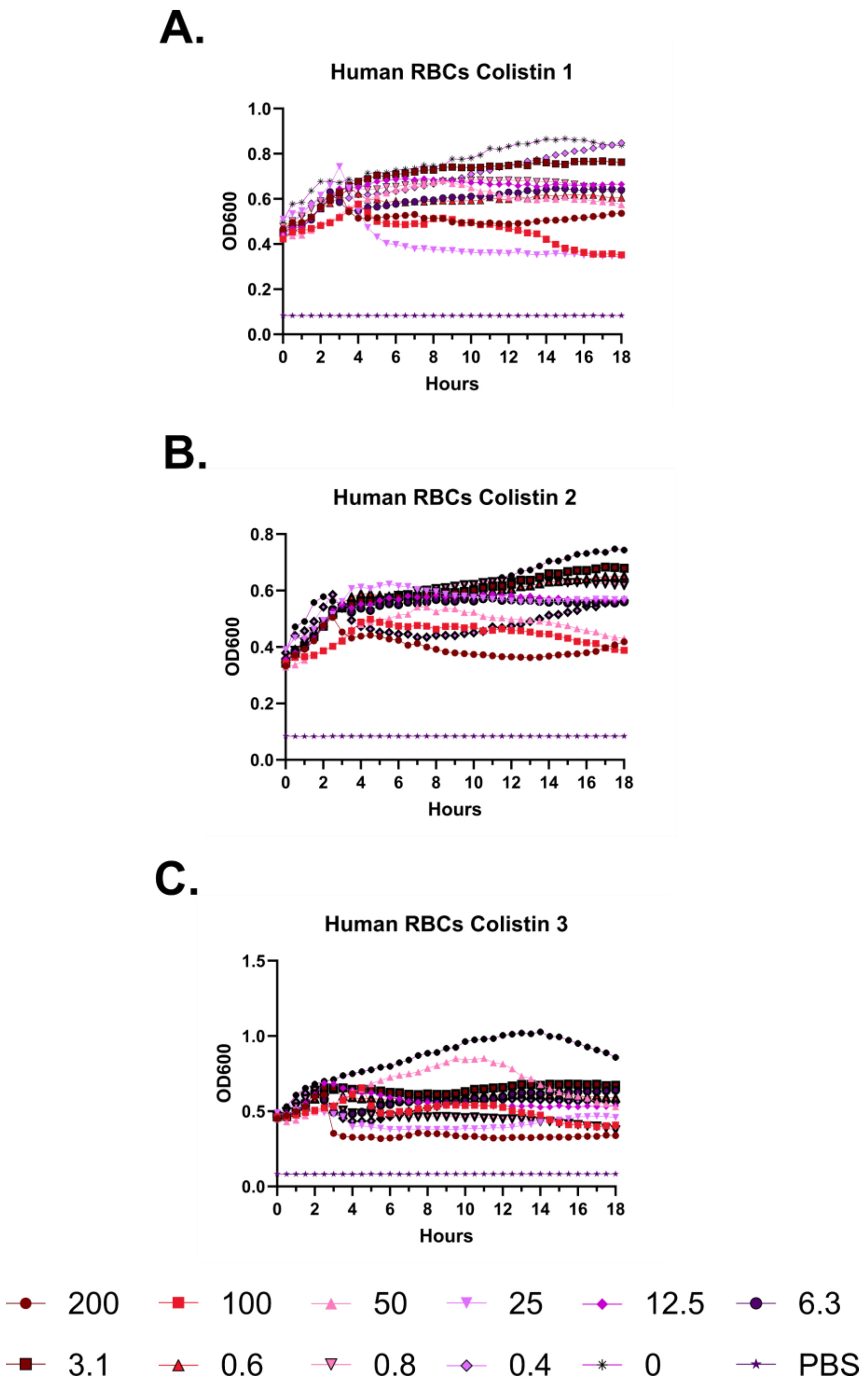
A.
Human RBCs Colistin 1
OD600
Hours
B.
Human RBCs Colistin 2
OD600
Hours
C.
Human RBCs Colistin 3
OD600
Hours
200
100
50
25
12.5
6.3
3.1
0.6
0.8
0.4
0
PBS

**Supplementary Figure 71 – Kinetic hemolysis assays for colistin in human red blood cells. A-C.** Hemolysis was monitored by OD600 over 18 hours at 37 °C in the presence of the indicated peptide concentrations. Decreasing OD600 indicates increasing hemolysis. Peptide concentrations are given in μM. Each curve indicates an independent experiment.

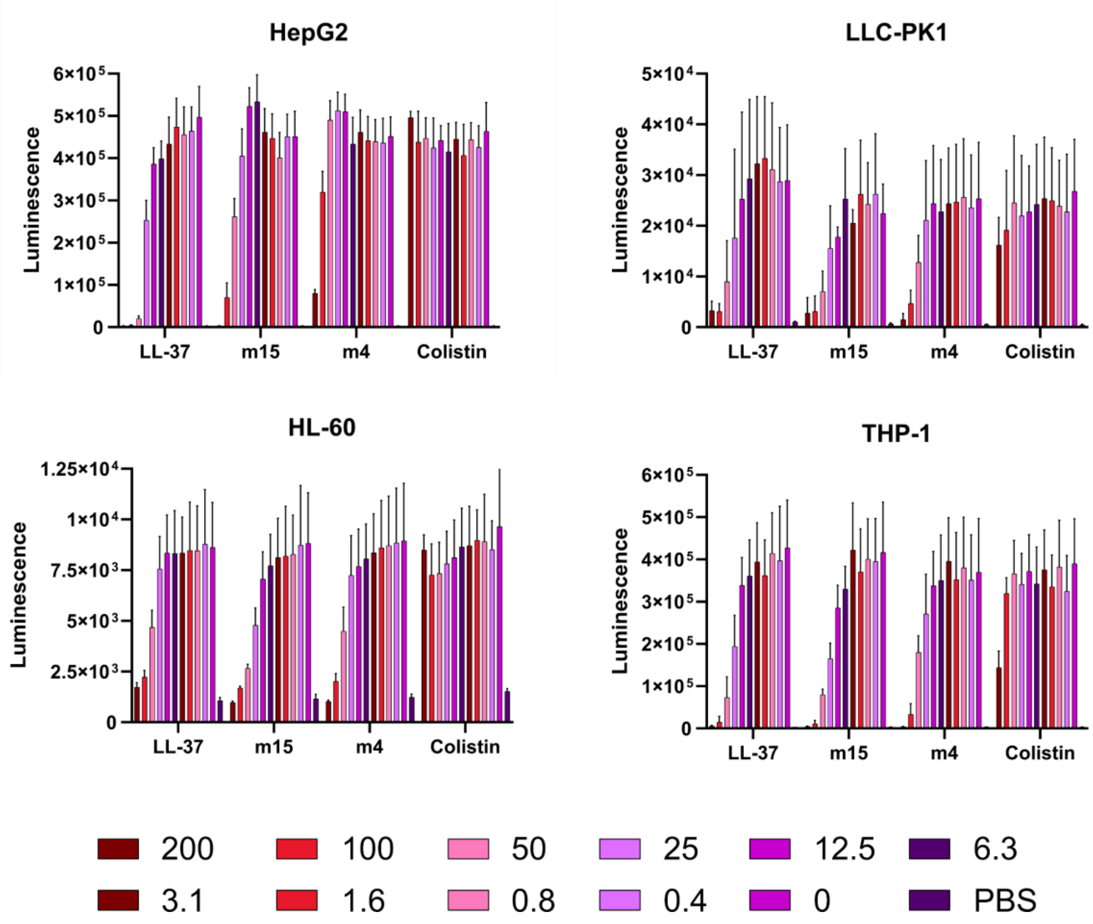
HepG2
LLC-PK1
HL-60
THP-1
Luminescence
LL-37
m15
m4
Colistin
6×10⁵
5×10⁵
4×10⁵
3×10⁵
2×10⁵
1×10⁵
0
5×10⁴
4×10⁴
3×10⁴
2×10⁴
1×10⁴
1.25×10⁴
1×10⁴
7.5×10³
5×10³
2.5×10³
200
100
50
25
12.5
6.3
3.1
1.6
0.8
0.4
0
PBS

**Supplementary Figure 72 – Histogram depiction of Figure 4B data. A.** Endpoint of RealTime Glo luciferase toxicity assay using the indicated cell types. Data are the mean and standard deviation from the raw luminescence measures of three independent experiments.

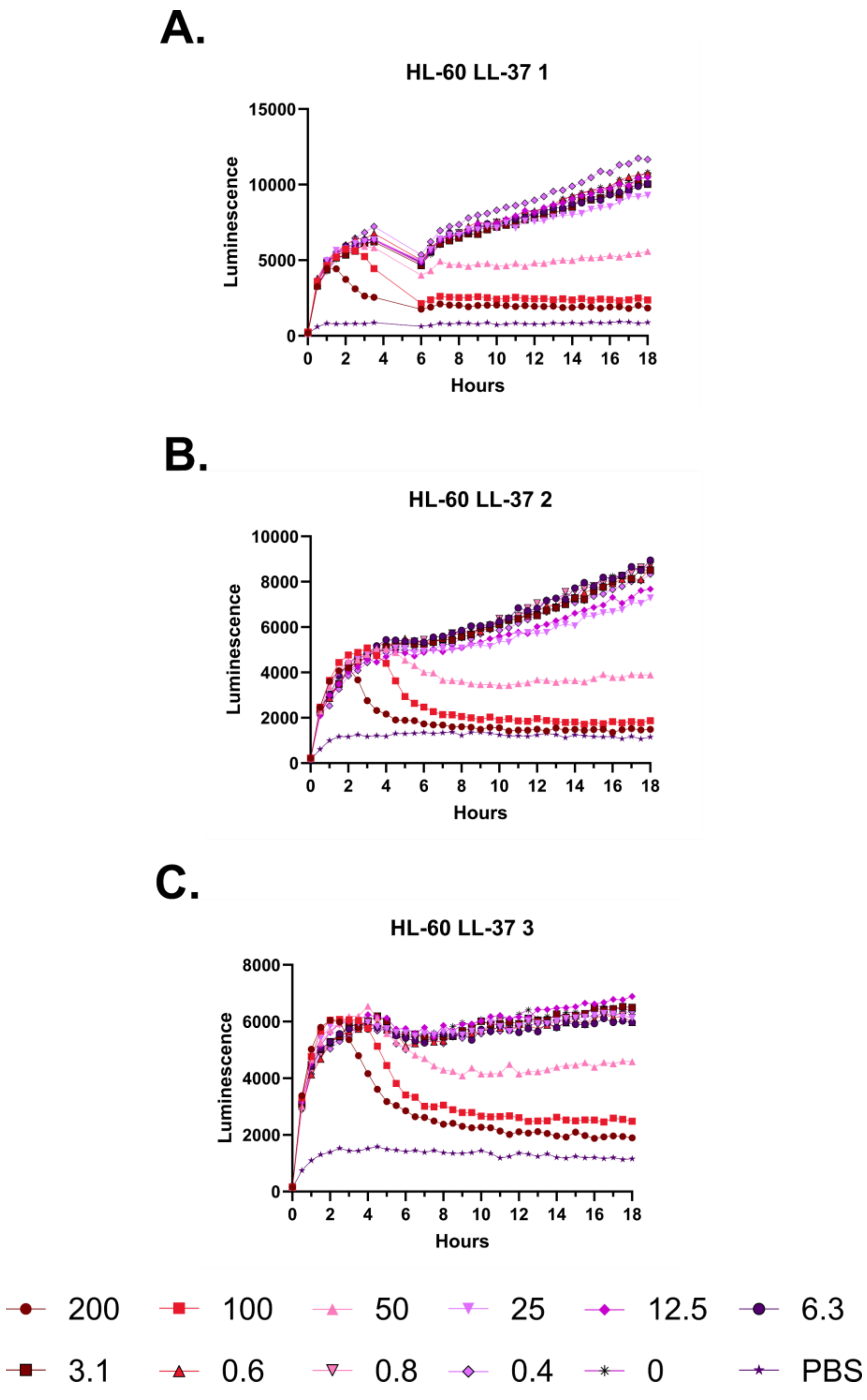
A.
HL-60 LL-37 1
15000
10000
5000
0
Luminescence
0 2 4 6 8 10 12 14 16 18
Hours
B.
HL-60 LL-37 2
10000
8000
6000
4000
2000
0
Luminescence
0 2 4 6 8 10 12 14 16 18
Hours
C.
HL-60 LL-37 3
8000
6000
4000
2000
0
Luminescence
0 2 4 6 8 10 12 14 16 18
Hours
200
100
50
25
12.5
6.3
3.1
0.6
0.8
0.4
0
PBS

**Supplementary Figure 73 – Kinetic luminescence toxicity assay for HL-60 treated with LL-37.** **A-C.** Luminescence was monitored over 18 hours at 37 °C with 5% $CO_2$ in a Small Humidity Cassette in the presence of the indicated peptide concentrations. Toxicity is indicated by curves that fall off of a steady upward trajectory. An instrument error in Experiment 1 resulted in no reads being taken during the gap in the curve. Peptide concentrations are given in μM. Each curve indicates an independent experiment.

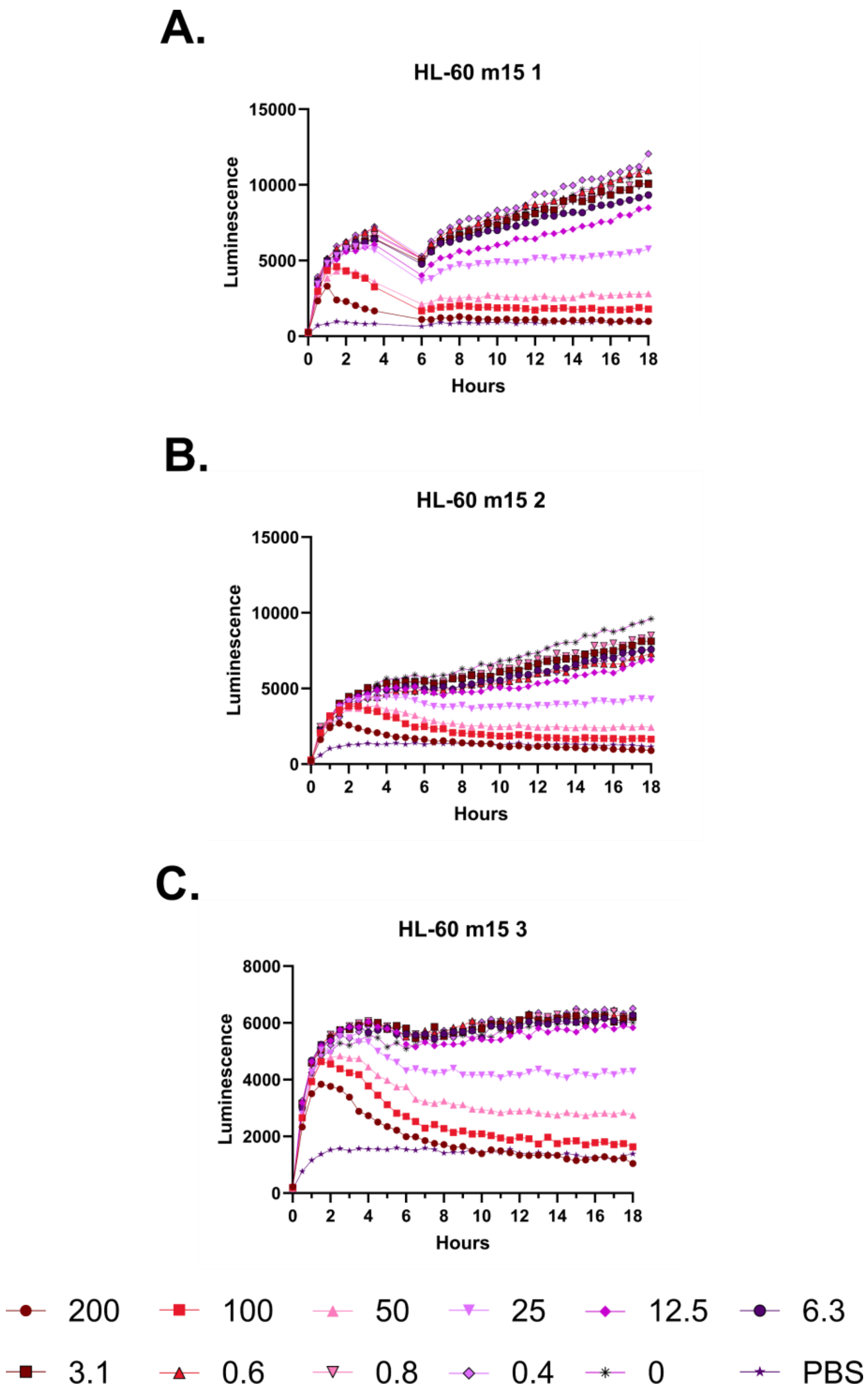
A.
HL-60 m15 1
15000
10000
5000
0
Luminescence
0 2 4 6 8 10 12 14 16 18
Hours
B.
HL-60 m15 2
15000
10000
5000
0
Luminescence
0 2 4 6 8 10 12 14 16 18
Hours
C.
HL-60 m15 3
8000
6000
4000
2000
0
Luminescence
0 2 4 6 8 10 12 14 16 18
Hours
200
100
50
25
12.5
6.3
3.1
0.6
0.8
0.4
0
PBS

**Supplementary Figure 74 – Kinetic luminescence toxicity assay for HL-60 treated with m15. A-C.** Luminescence was monitored over 18 hours at 37 °C with 5% $CO_2$ in a Small Humidity Cassette in the presence of the indicated peptide concentrations. Toxicity is indicated by curves that fall off of a steady upward trajectory. An instrument error in Experiment 1 resulted in no reads being taken during the gap in the curve. Peptide concentrations are given in μM. Each curve indicates an independent experiment.

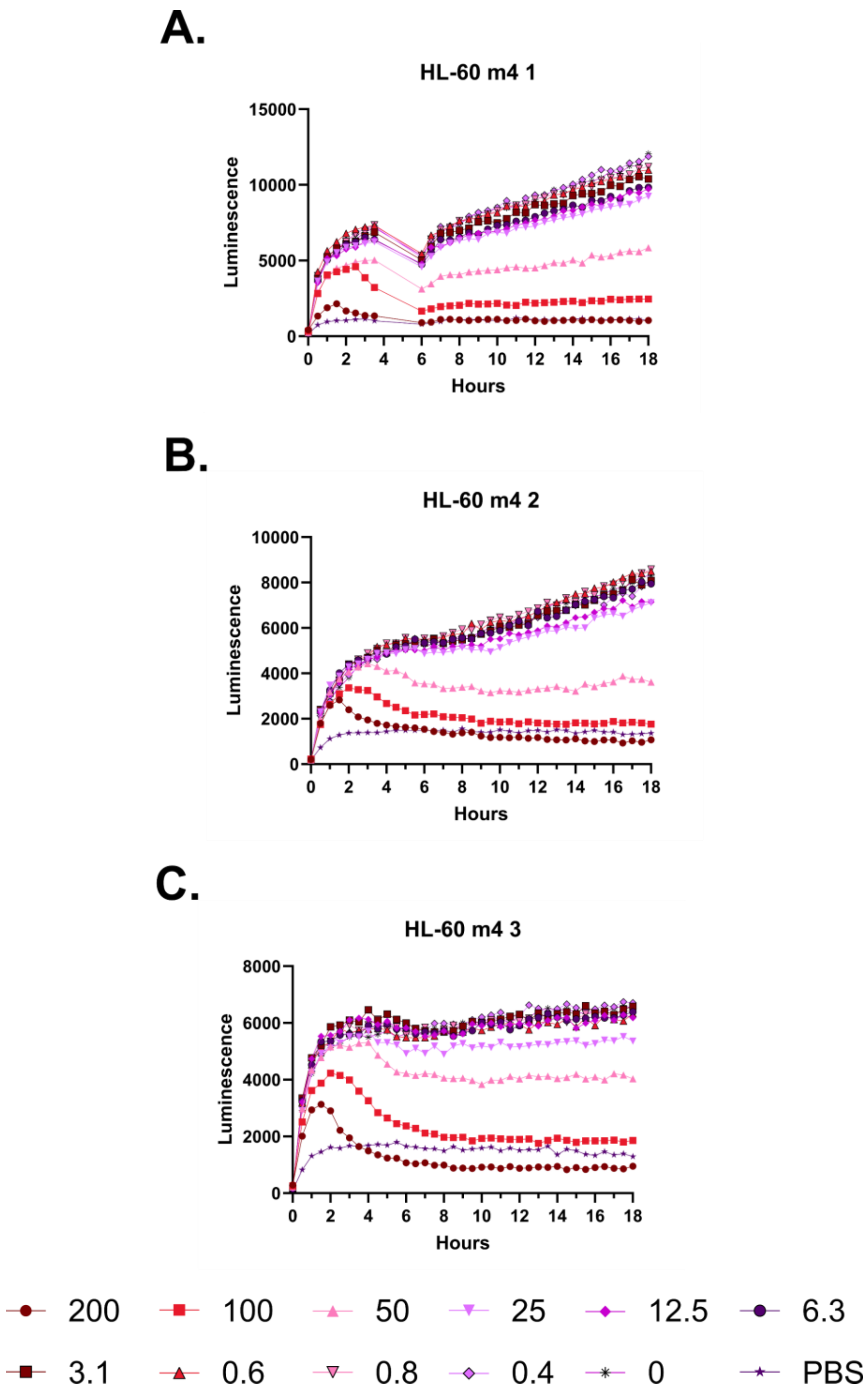
A.
HL-60 m4 1
Luminescence
Hours
B.
HL-60 m4 2
Luminescence
Hours
C.
HL-60 m4 3
Luminescence
Hours
200
100
50
25
12.5
6.3
3.1
0.6
0.8
0.4
0
PBS

**Supplementary Figure 75 – Kinetic luminescence toxicity assay for HL-60 treated with m4. A-C.** Luminescence was monitored over 18 hours at 37 °C with 5% $CO_2$ in a Small Humidity Cassette in the presence of the indicated peptide concentrations. Toxicity is indicated by curves that fall off of a steady upward trajectory. An instrument error in Experiment 1 resulted in no reads being taken during the gap in the curve. Peptide concentrations are given in μM. Each curve indicates an independent experiment.

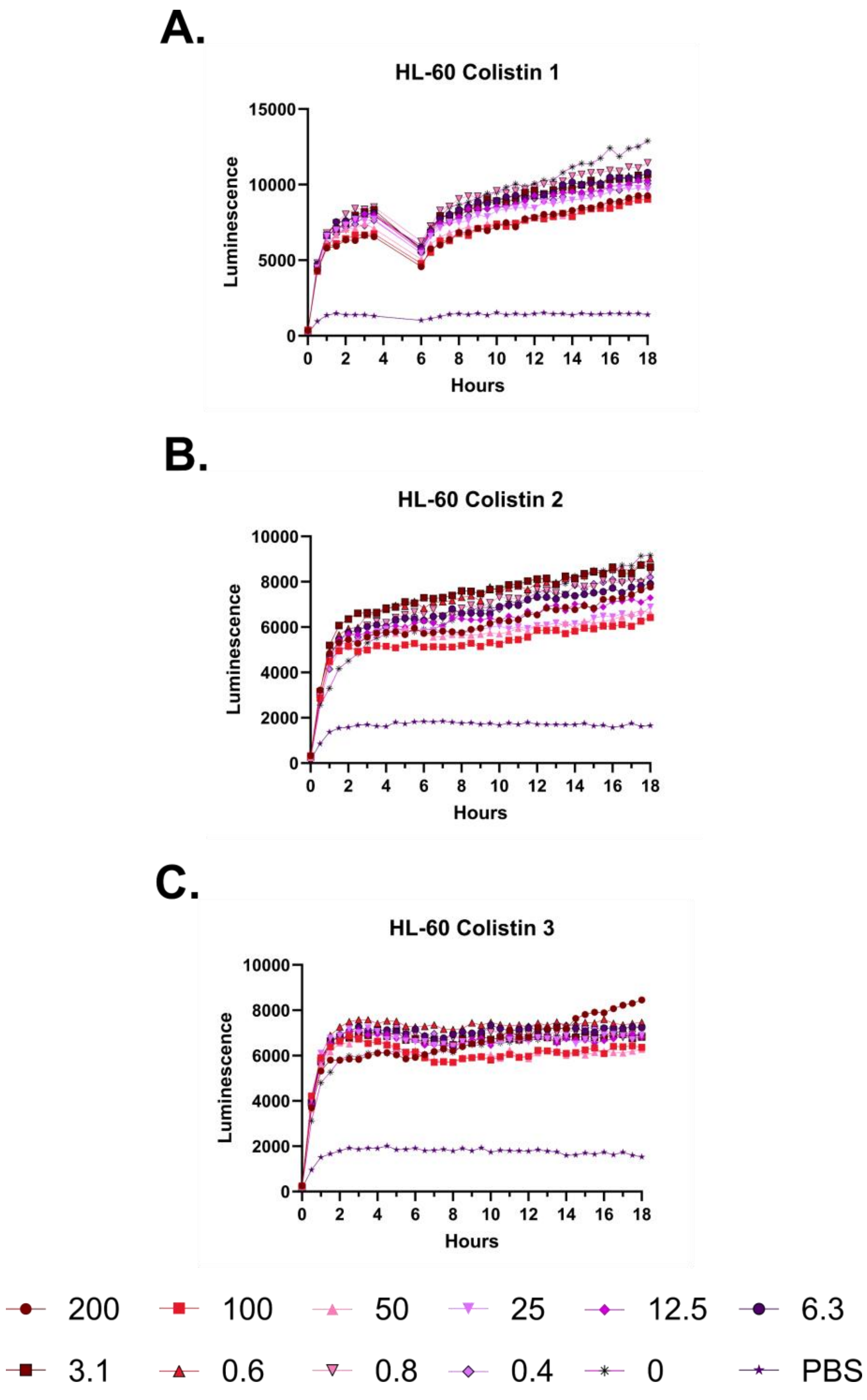
A.
HL-60 Colistin 1
Luminescence
Hours
B.
HL-60 Colistin 2
Luminescence
Hours
C.
HL-60 Colistin 3
Luminescence
Hours
200
100
50
25
12.5
6.3
3.1
0.6
0.8
0.4
0
PBS

**Supplementary Figure 76 – Kinetic luminescence toxicity assay for HL-60 treated with colistin. A-C.** Luminescence was monitored over 18 hours at 37 °C with 5% $CO_2$ in a Small Humidity Cassette in the presence of the indicated peptide concentrations. Toxicity is indicated by curves that fall off of a steady upward trajectory. An instrument error in Experiment 1 resulted in no reads being taken during the gap in the curve. Peptide concentrations are given in µM. Each curve indicates an independent experiment.

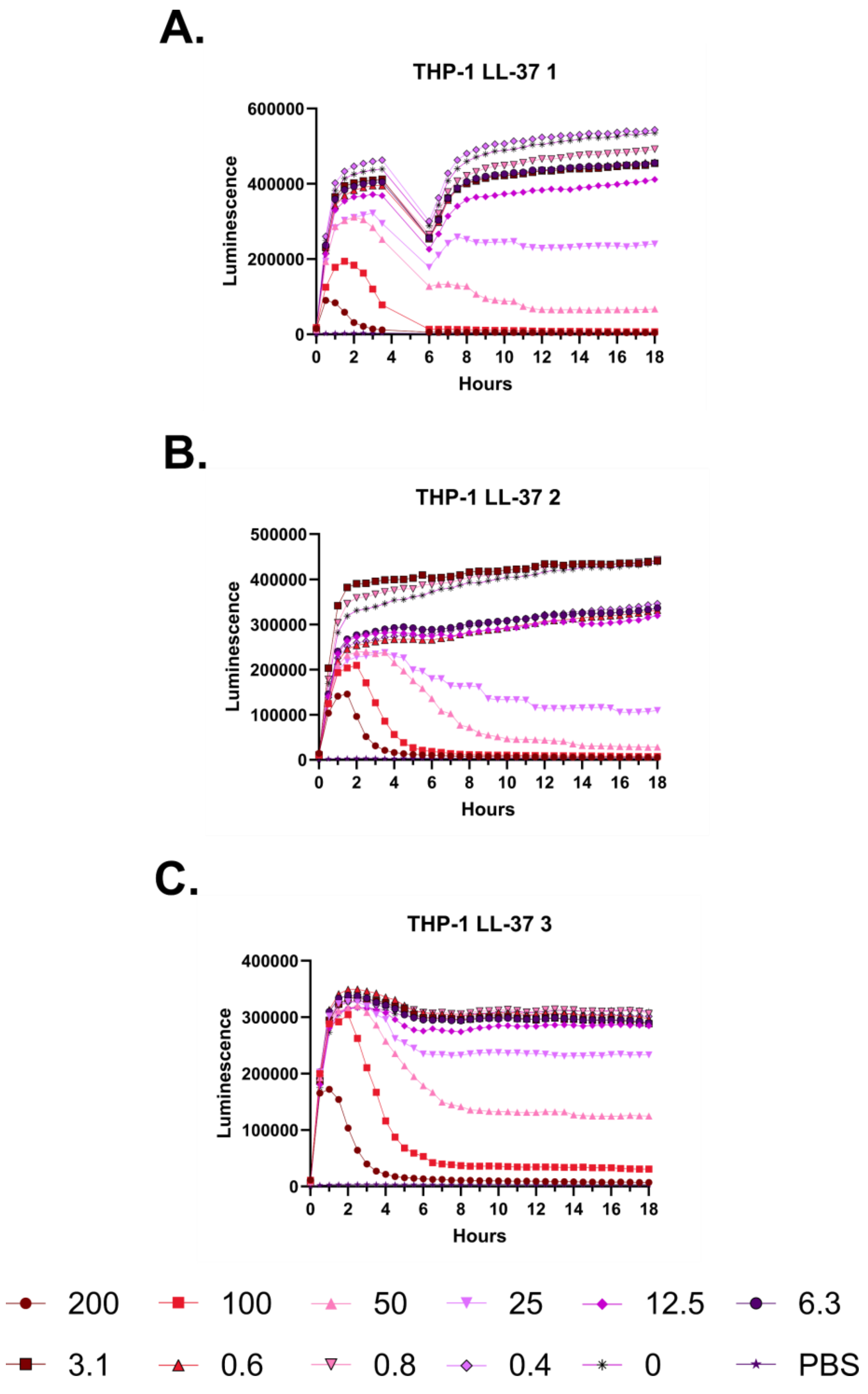
A.
THP-1 LL-37 1
600000
400000
200000
0
Luminescence
0 2 4 6 8 10 12 14 16 18
Hours
B.
THP-1 LL-37 2
500000
400000
300000
200000
100000
0
Luminescence
0 2 4 6 8 10 12 14 16 18
Hours
C.
THP-1 LL-37 3
400000
300000
200000
100000
0
Luminescence
0 2 4 6 8 10 12 14 16 18
Hours
200
100
50
25
12.5
6.3
3.1
0.6
0.8
0.4
0
PBS

**Supplementary Figure 77 – Kinetic luminescence toxicity assay for THP-1 treated with LL-37. A-C.** Luminescence was monitored over 18 hours at 37 °C with 5% $CO_2$ in a Small Humidity Cassette in the presence of the indicated peptide concentrations. Toxicity is indicated by curves that fall off of a steady upward trajectory. An instrument error in Experiment 1 resulted in no reads being taken during the gap in the curve. Peptide concentrations are given in µM. Each curve indicates an independent experiment.

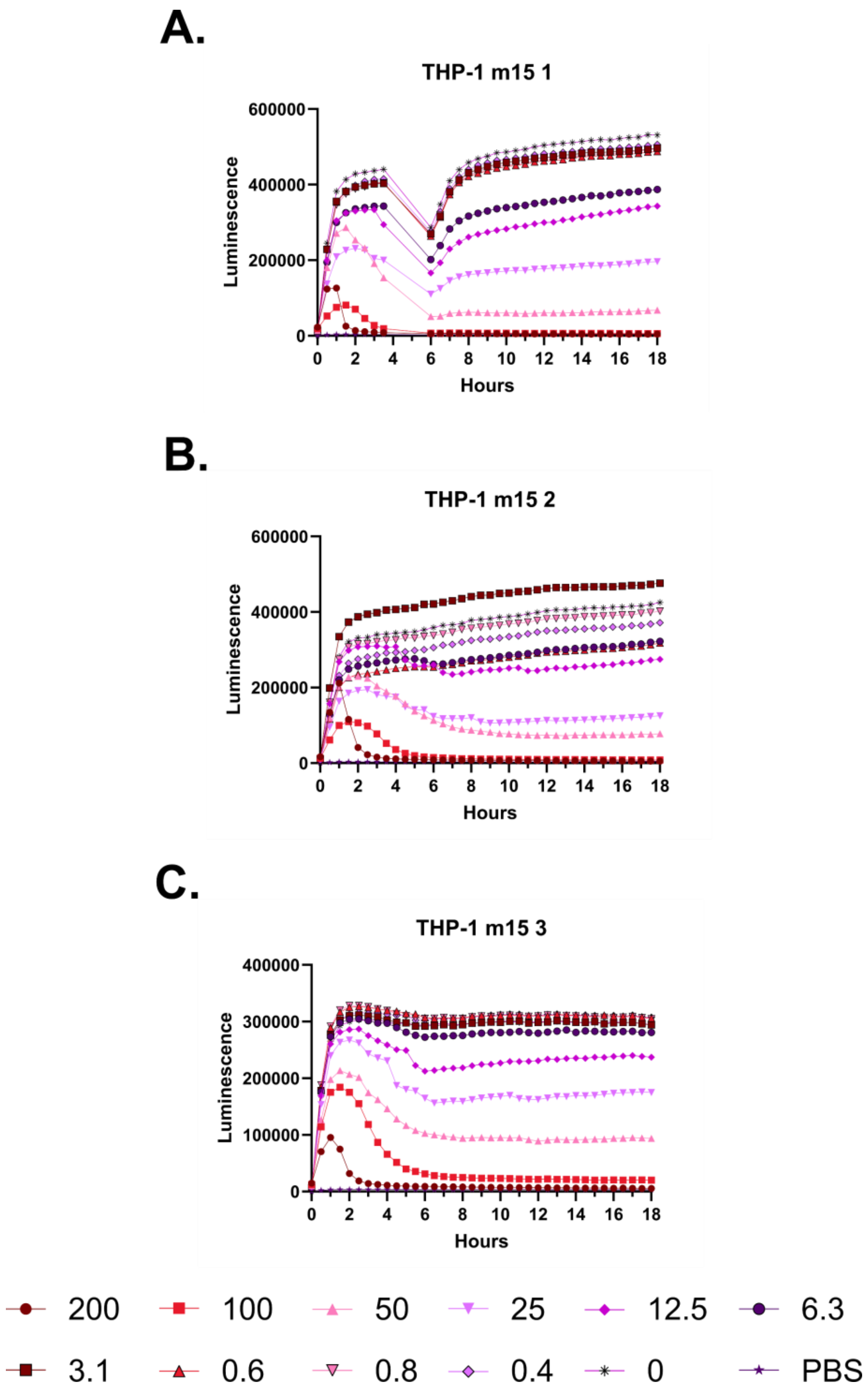
A.
THP-1 m15 1
600000
400000
200000
0
Luminescence
0 2 4 6 8 10 12 14 16 18
Hours
B.
THP-1 m15 2
600000
400000
200000
0
Luminescence
0 2 4 6 8 10 12 14 16 18
Hours
C.
THP-1 m15 3
400000
300000
200000
100000
0
Luminescence
0 2 4 6 8 10 12 14 16 18
Hours
200
100
50
25
12.5
6.3
3.1
0.6
0.8
0.4
0
PBS

**Supplementary Figure 78 – Kinetic luminescence toxicity assay for THP-1 treated with m15. A-C.** Luminescence was monitored over 18 hours at 37 °C with 5% $CO_2$ in a Small Humidity Cassette in the presence of the indicated peptide concentrations. Toxicity is indicated by curves that fall off of a steady upward trajectory. An instrument error in Experiment 1 resulted in no reads being taken during the gap in the curve. Peptide concentrations are given in µM. Each curve indicates an independent experiment.

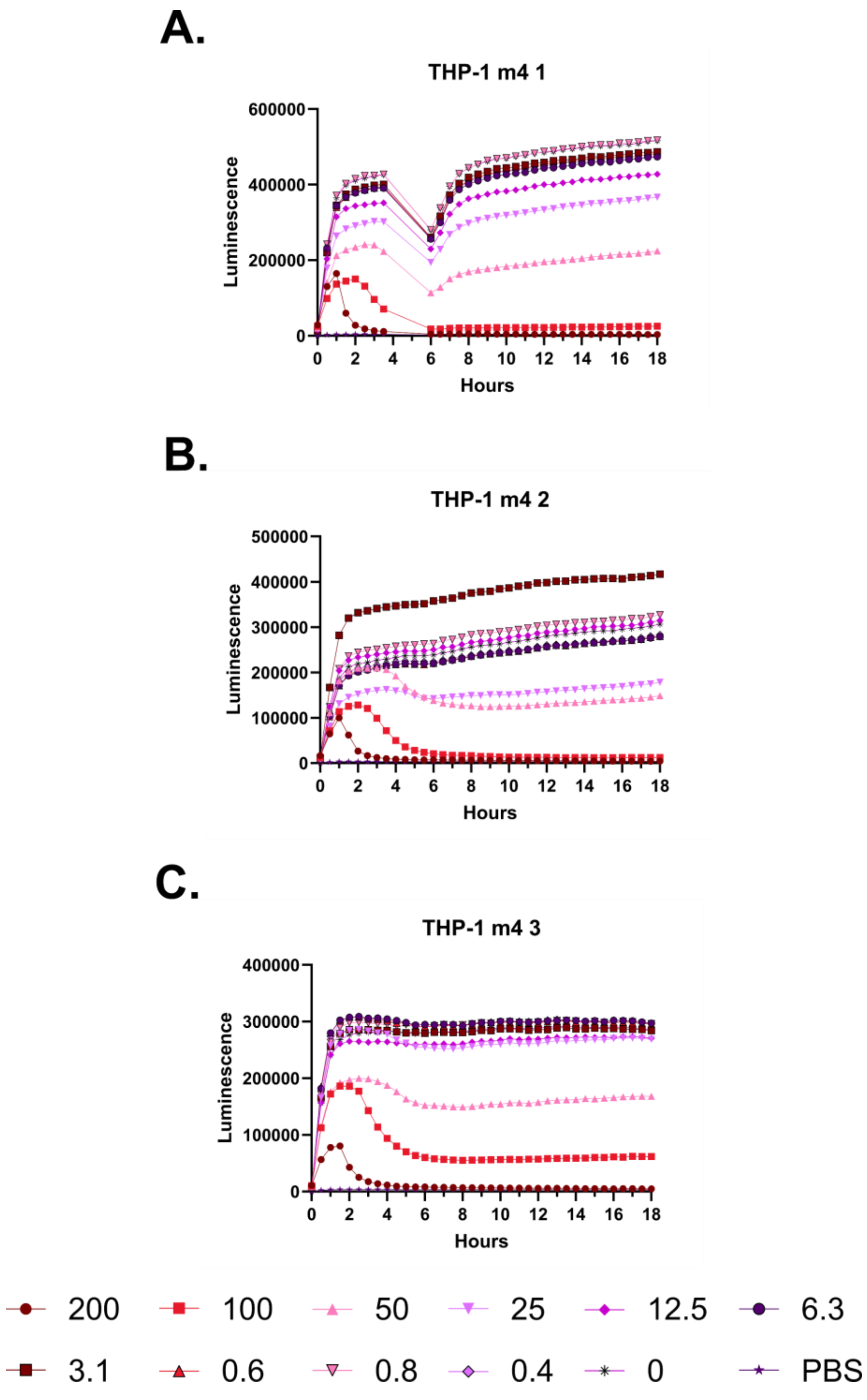
A.
THP-1 m4 1
Luminescence
Hours
B.
THP-1 m4 2
Luminescence
Hours
C.
THP-1 m4 3
Luminescence
Hours
200
100
50
25
12.5
6.3
3.1
0.6
0.8
0.4
0
PBS

**Supplementary Figure 79 – Kinetic luminescence toxicity assay for THP-1 treated with m4. A-C.** Luminescence was monitored over 18 hours at 37 °C with 5% $CO_2$ in a Small Humidity Cassette in the presence of the indicated peptide concentrations. Toxicity is indicated by curves that fall off of a steady upward trajectory. An instrument error in Experiment 1 resulted in no reads being taken during the gap in the curve. Peptide concentrations are given in μM. Each curve indicates an independent experiment.

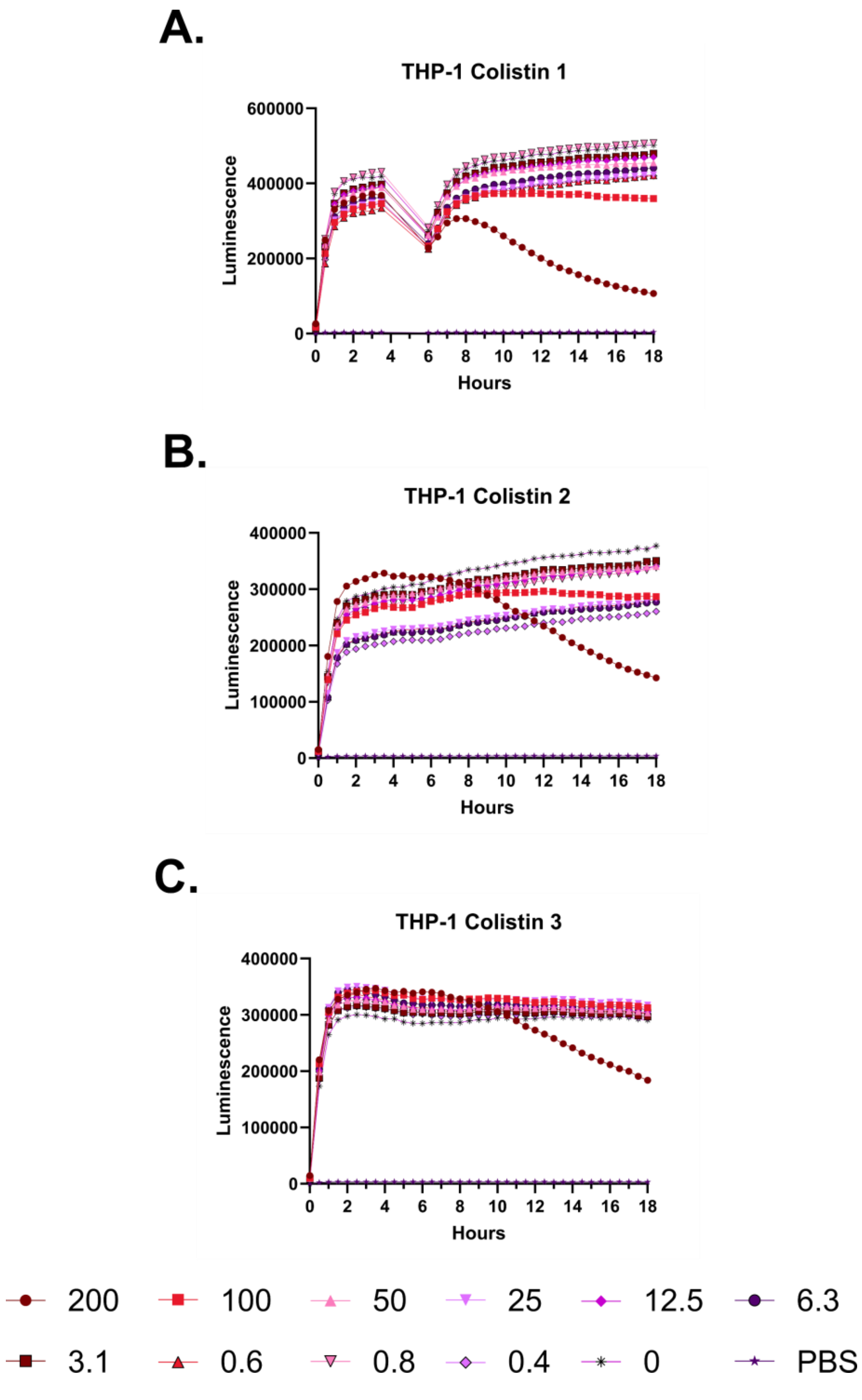
A.
THP-1 Colistin 1
Luminescence
600000
400000
200000
0
0 2 4 6 8 10 12 14 16 18
Hours
B.
THP-1 Colistin 2
Luminescence
400000
300000
200000
100000
0
0 2 4 6 8 10 12 14 16 18
Hours
C.
THP-1 Colistin 3
Luminescence
400000
300000
200000
100000
0
0 2 4 6 8 10 12 14 16 18
Hours
200
100
50
25
12.5
6.3
3.1
0.6
0.8
0.4
0
PBS

**Supplementary Figure 80 – Kinetic luminescence toxicity assay for THP-1 treated with colistin. A-C.** Luminescence was monitored over 18 hours at 37 °C with 5% $CO_2$ in a Small Humidity Cassette in the presence of the indicated peptide concentrations. Toxicity is indicated by curves that fall off of a steady upward trajectory. An instrument error in Experiment 1 resulted in no reads being taken during the gap in the curve. Peptide concentrations are given in µM. Each curve indicates an independent experiment.

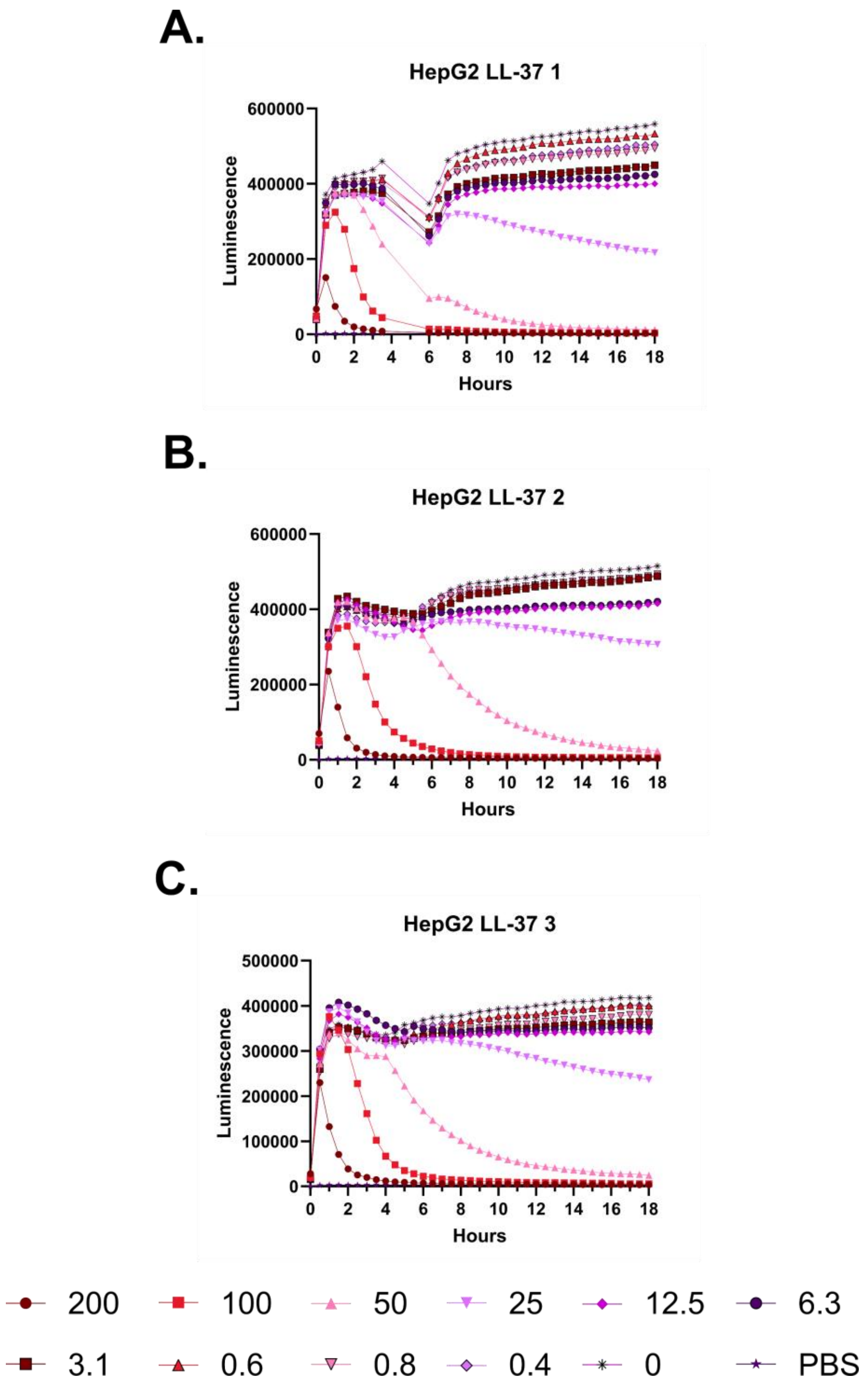
A.
HepG2 LL-37 1
Luminescence
Hours
B.
HepG2 LL-37 2
Luminescence
Hours
C.
HepG2 LL-37 3
Luminescence
Hours
200
100
50
25
12.5
6.3
3.1
0.6
0.8
0.4
0
PBS

**Supplementary Figure 81 – Kinetic luminescence toxicity assay for HepG2 treated with LL-37. A-C.** Luminescence was monitored over 18 hours at 37 °C with 5% $CO_2$ in a Small Humidity Cassette in the presence of the indicated peptide concentrations. Toxicity is indicated by curves that fall off of a steady upward trajectory. An instrument error in Experiment 1 resulted in no reads being taken during the gap in the curve. Peptide concentrations are given in µM. Each curve indicates an independent experiment.

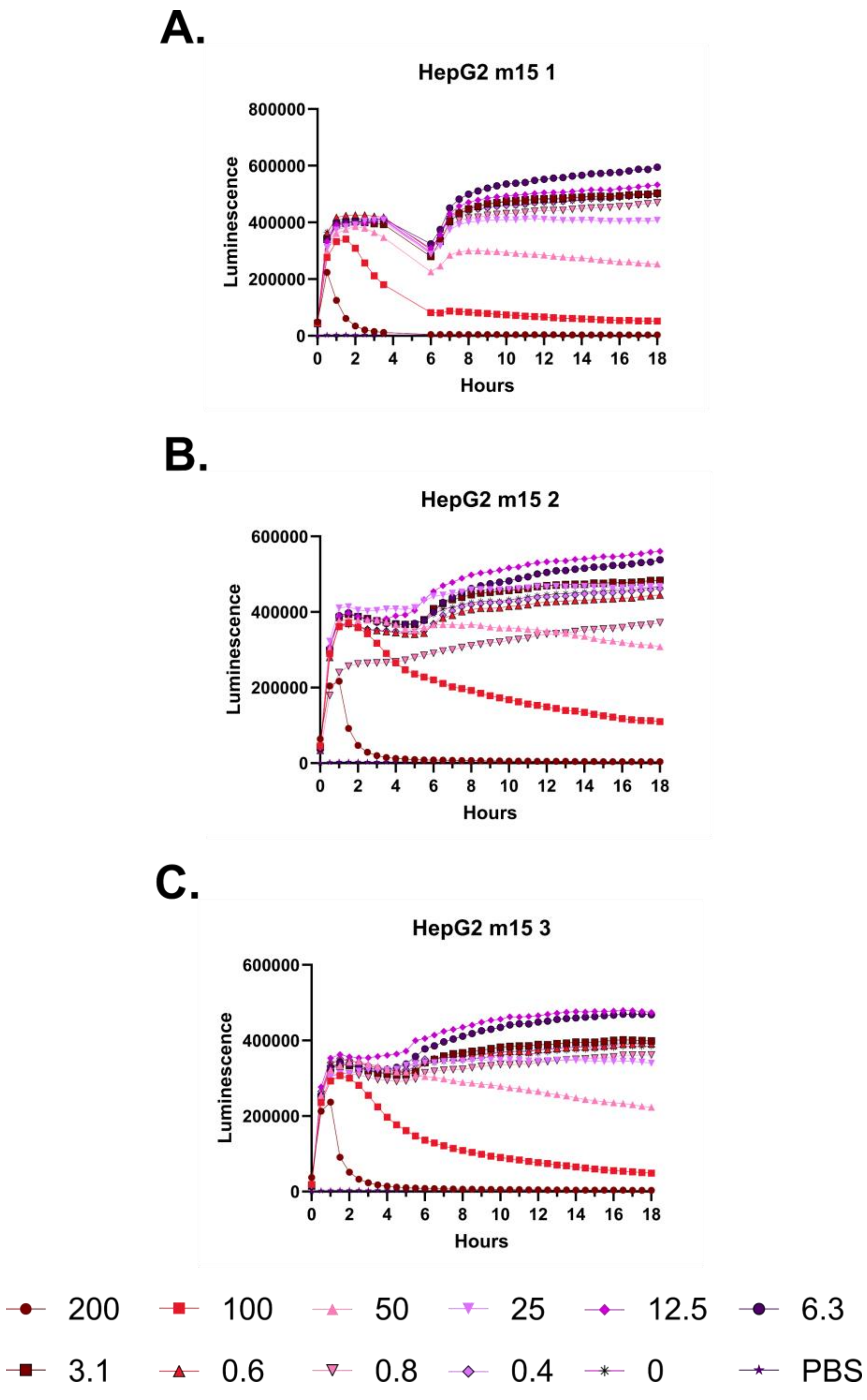
A.
HepG2 m15 1
Luminescence
Hours
B.
HepG2 m15 2
Luminescence
Hours
C.
HepG2 m15 3
Luminescence
Hours
200
100
50
25
12.5
6.3
3.1
0.6
0.8
0.4
0
PBS

**Supplementary Figure 82 – Kinetic luminescence toxicity assay for HepG2 treated with m15. A-C.** Luminescence was monitored over 18 hours at 37 °C with 5% $CO_2$ in a Small Humidity Cassette in the presence of the indicated peptide concentrations. Toxicity is indicated by curves that fall off of a steady upward trajectory. An instrument error in Experiment 1 resulted in no reads being taken during the gap in the curve. Peptide concentrations are given in μM. Each curve indicates an independent experiment.

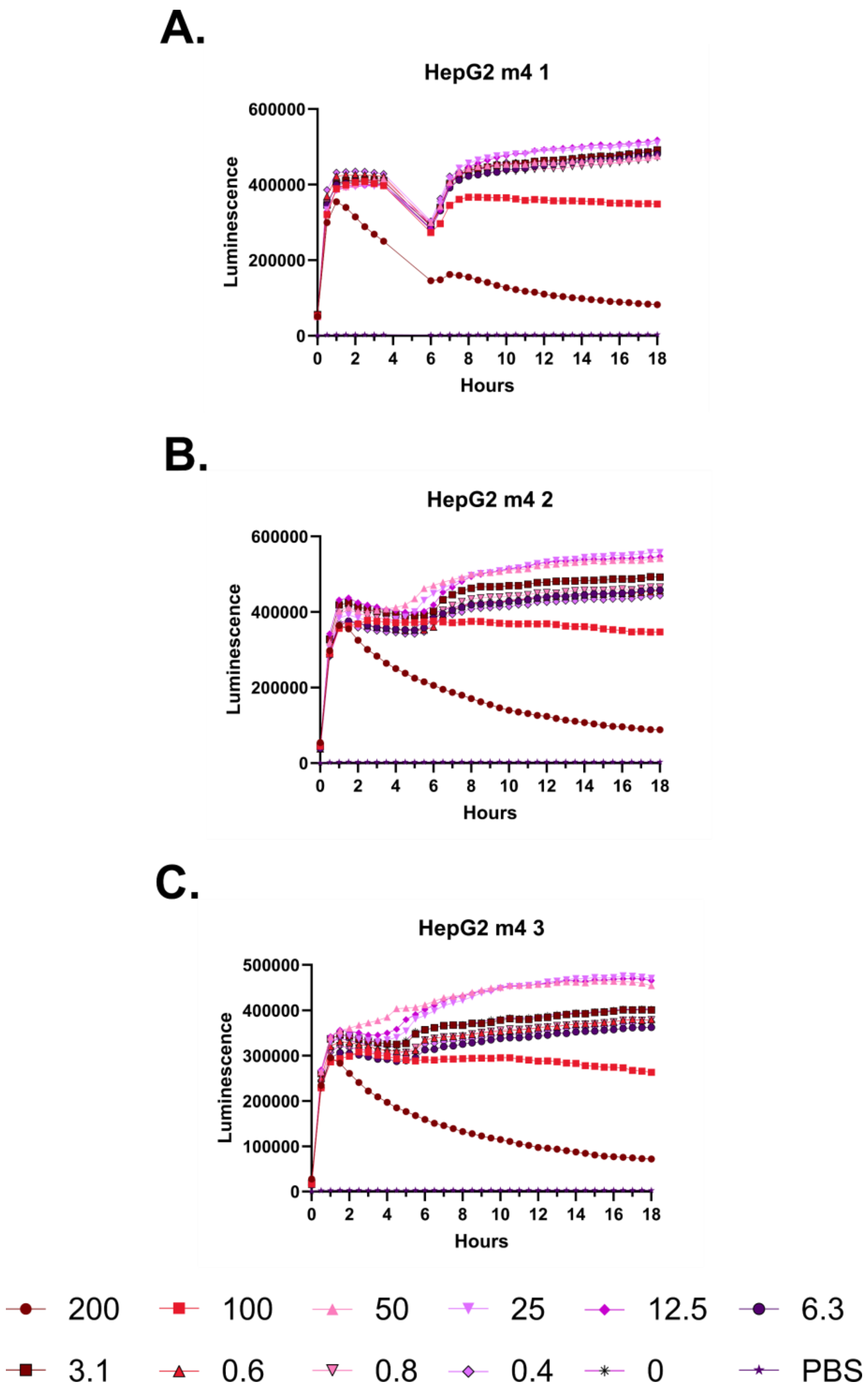
A.
HepG2 m4 1
Luminescence
Hours
B.
HepG2 m4 2
Luminescence
Hours
C.
HepG2 m4 3
Luminescence
Hours
200
100
50
25
12.5
6.3
3.1
0.6
0.8
0.4
0
PBS

**Supplementary Figure 83 – Kinetic luminescence toxicity assay for HepG2 treated with m4. A-C.** Luminescence was monitored over 18 hours at 37 °C with 5% $CO_2$ in a Small Humidity Cassette in the presence of the indicated peptide concentrations. Toxicity is indicated by curves that fall off of a steady upward trajectory. An instrument error in Experiment 1 resulted in no reads being taken during the gap in the curve. Peptide concentrations are given in μM. Each curve indicates an independent experiment.

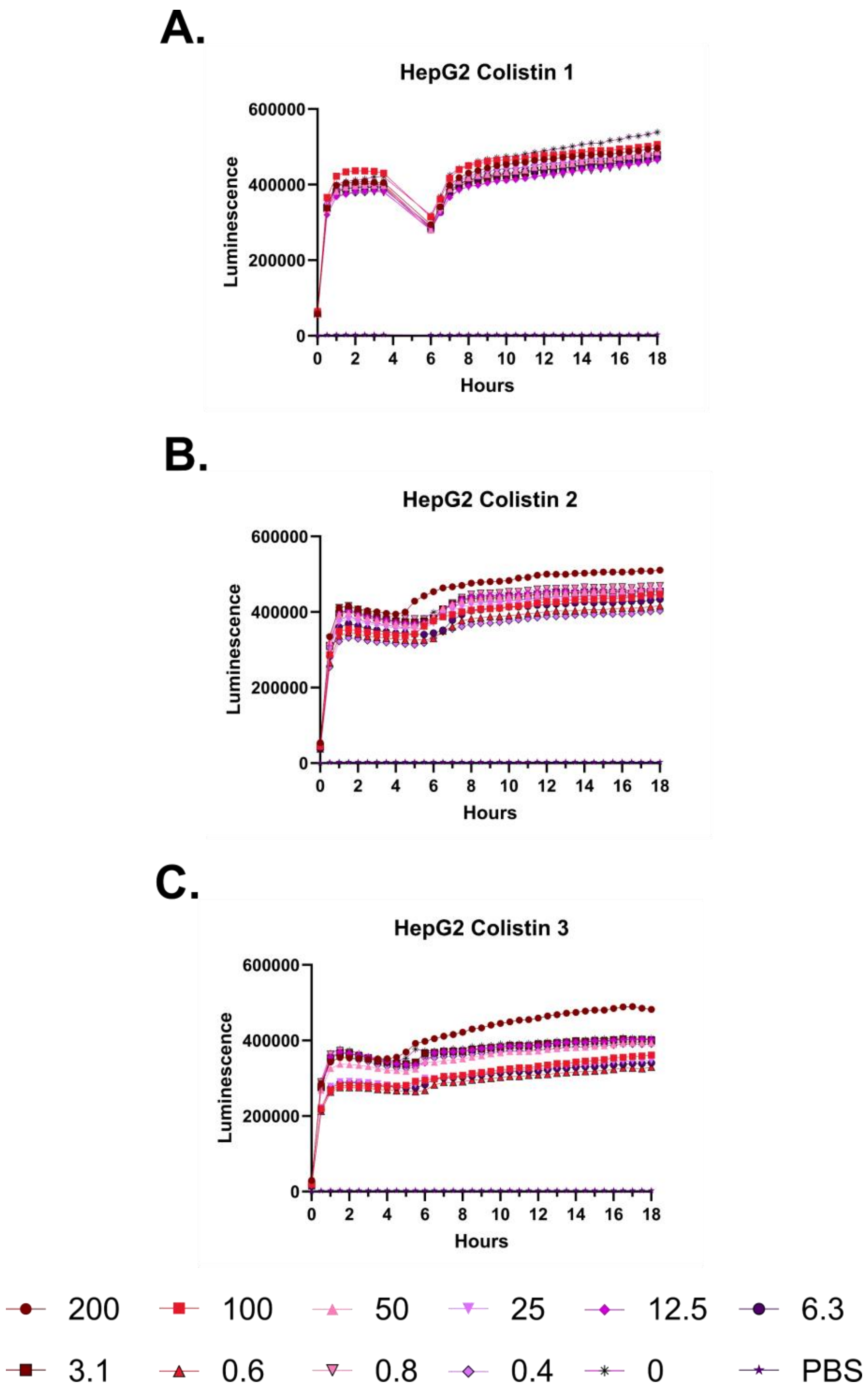
A.
HepG2 Colistin 1
600000
400000
200000
0
Luminescence
0 2 4 6 8 10 12 14 16 18
Hours
B.
HepG2 Colistin 2
600000
400000
200000
0
Luminescence
0 2 4 6 8 10 12 14 16 18
Hours
C.
HepG2 Colistin 3
600000
400000
200000
0
Luminescence
0 2 4 6 8 10 12 14 16 18
Hours
200
100
50
25
12.5
6.3
3.1
0.6
0.8
0.4
0
PBS

**Supplementary Figure 84 – Kinetic luminescence toxicity assay for HepG2 treated with colistin. A-C.** Luminescence was monitored over 18 hours at 37 °C with 5% $CO_2$ in a Small Humidity Cassette in the presence of the indicated peptide concentrations. Toxicity is indicated by curves that fall off of a steady upward trajectory. An instrument error in Experiment 1 resulted in no reads being taken during the gap in the curve. Peptide concentrations are given in µM. Each curve indicates an independent experiment.

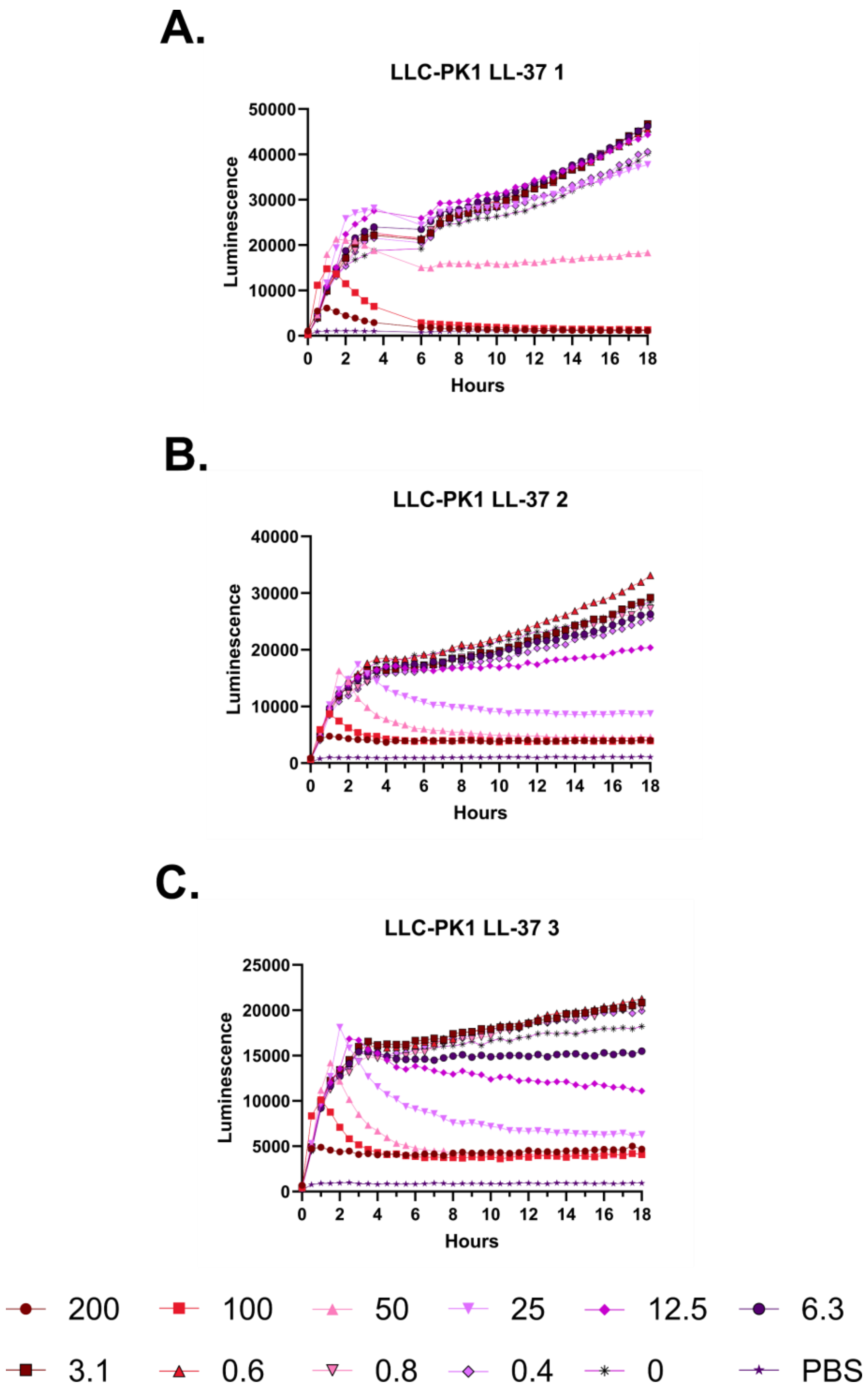
A.
LLC-PK1 LL-37 1
Luminescence
Hours
B.
LLC-PK1 LL-37 2
Luminescence
Hours
C.
LLC-PK1 LL-37 3
Luminescence
Hours
200
100
50
25
12.5
6.3
3.1
0.6
0.8
0.4
0
PBS

**Supplementary Figure 85 – Kinetic luminescence toxicity assay for LLC-PK1 treated with LL-37. A-C.** Luminescence was monitored over 18 hours at 37 °C with 5% $CO_2$ in a Small Humidity Cassette in the presence of the indicated peptide concentrations. Toxicity is indicated by curves that fall off of a steady upward trajectory. An instrument error in Experiment 1 resulted in no reads being taken during the gap in the curve. Peptide concentrations are given in µM. Each curve indicates an independent experiment.

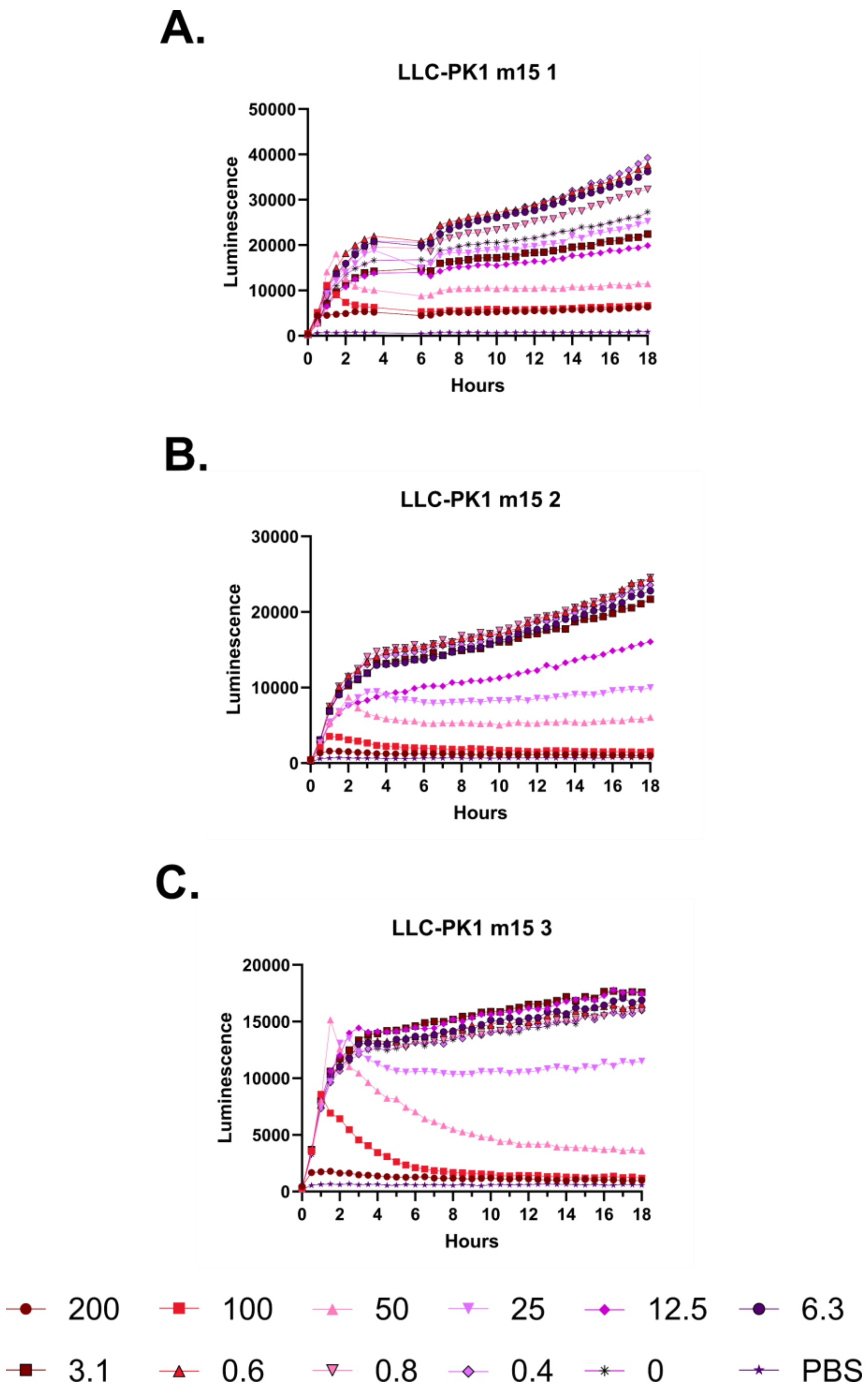
A.
LLC-PK1 m15 1
Luminescence
50000
40000
30000
20000
10000
0
0 2 4 6 8 10 12 14 16 18
Hours
B.
LLC-PK1 m15 2
Luminescence
30000
20000
10000
0
0 2 4 6 8 10 12 14 16 18
Hours
C.
LLC-PK1 m15 3
Luminescence
20000
15000
10000
5000
0
0 2 4 6 8 10 12 14 16 18
Hours
200
100
50
25
12.5
6.3
3.1
0.6
0.8
0.4
0
PBS

**Supplementary Figure 86 – Kinetic luminescence toxicity assay for LLC-PK1 treated with m15. A-C.** Luminescence was monitored over 18 hours at 37 °C with 5% $CO_2$ in a Small Humidity Cassette in the presence of the indicated peptide concentrations. Toxicity is indicated by curves that fall off of a steady upward trajectory. An instrument error in Experiment 1 resulted in no reads being taken during the gap in the curve. Peptide concentrations are given in μM. Each curve indicates an independent experiment.

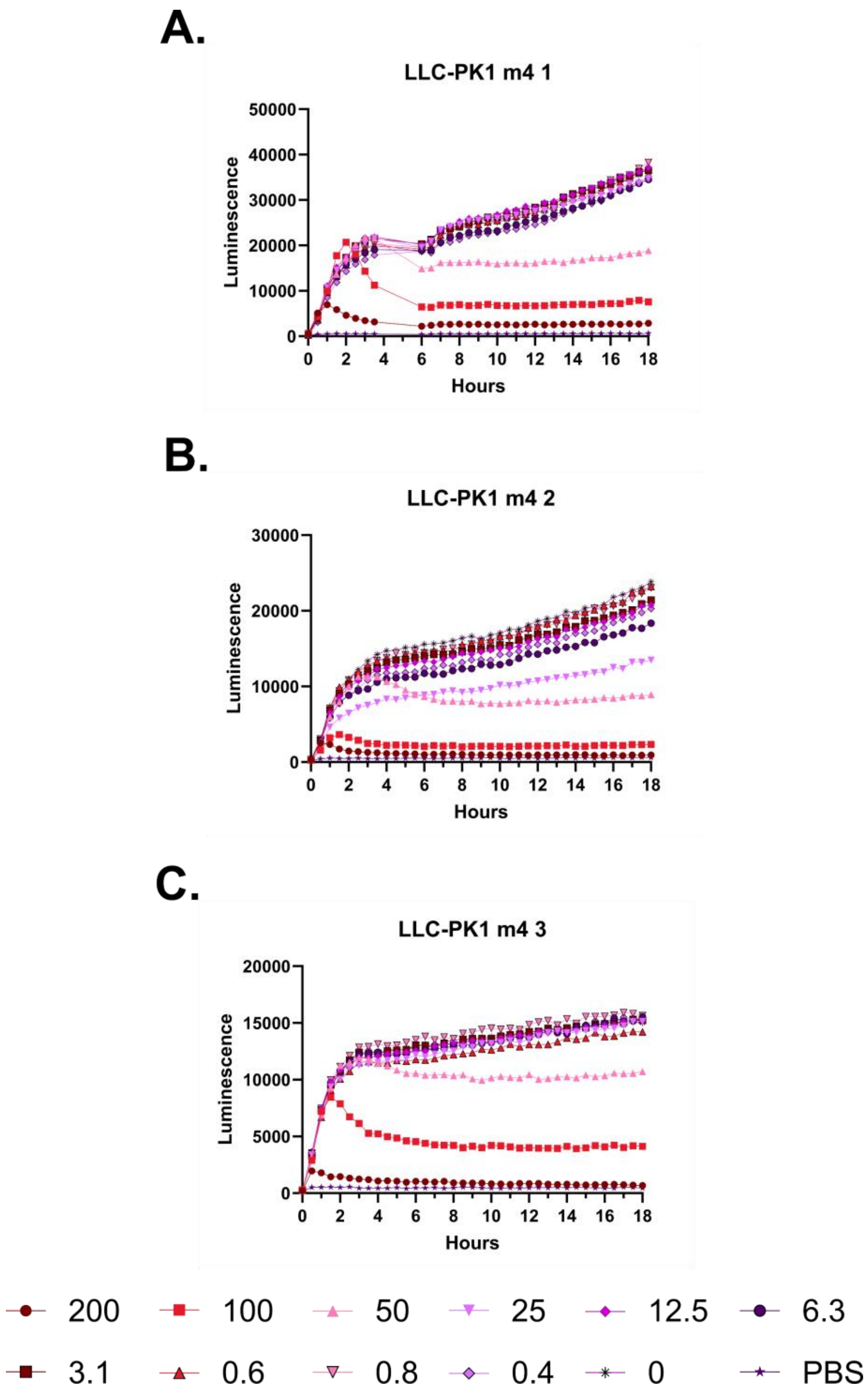
A.
LLC-PK1 m4 1
Luminescence
Hours
0 10000 20000 30000 40000 50000
0 2 4 6 8 10 12 14 16 18
B.
LLC-PK1 m4 2
Luminescence
Hours
0 10000 20000 30000
0 2 4 6 8 10 12 14 16 18
C.
LLC-PK1 m4 3
Luminescence
Hours
0 5000 10000 15000 20000
0 2 4 6 8 10 12 14 16 18
200
100
50
25
12.5
6.3
3.1
0.6
0.8
0.4
0
PBS

**Supplementary Figure 87 – Kinetic luminescence toxicity assay for LLC-PK1 treated with m4. A-C.** Luminescence was monitored over 18 hours at 37 °C with 5% $CO_2$ in a Small Humidity Cassette in the presence of the indicated peptide concentrations. Toxicity is indicated by curves that fall off of a steady upward trajectory. An instrument error in Experiment 1 resulted in no reads being taken during the gap in the curve. Peptide concentrations are given in µM. Each curve indicates an independent experiment.

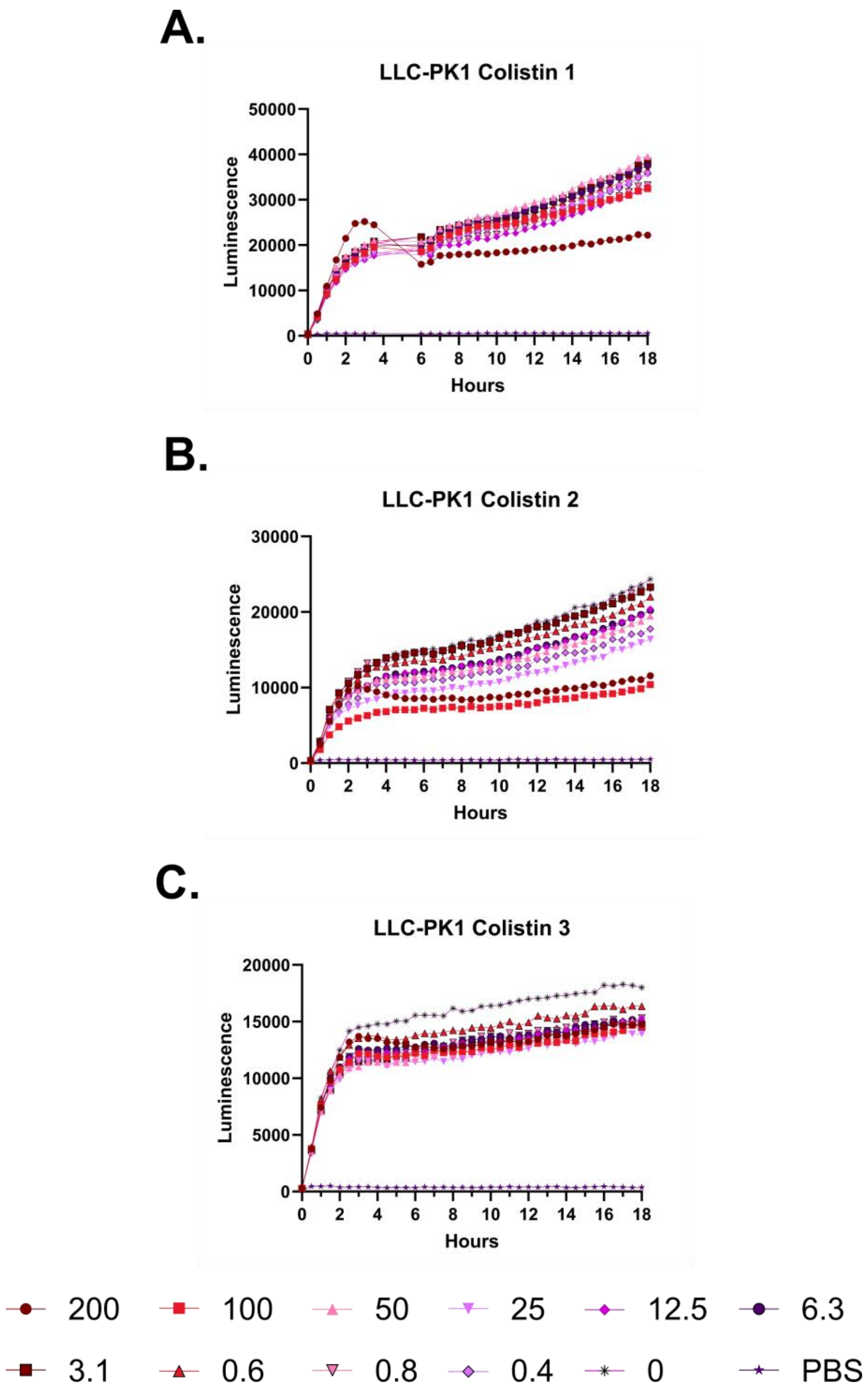
A.
LLC-PK1 Colistin 1
Luminescence
0
10000
20000
30000
40000
50000
0 2 4 6 8 10 12 14 16 18
Hours
B.
LLC-PK1 Colistin 2
Luminescence
0
10000
20000
30000
0 2 4 6 8 10 12 14 16 18
Hours
C.
LLC-PK1 Colistin 3
Luminescence
0
5000
10000
15000
20000
0 2 4 6 8 10 12 14 16 18
Hours
200
100
50
25
12.5
6.3
3.1
0.6
0.8
0.4
0
PBS

**Supplementary Figure 88 – Kinetic luminescence toxicity assay for LLC-PK1 treated with colistin. A-C.** Luminescence was monitored over 18 hours at 37 °C with 5% $CO_2$ in a Small Humidity Cassette in the presence of the indicated peptide concentrations. Toxicity is indicated by curves that fall off of a steady upward trajectory. An instrument error in Experiment 1 resulted in no reads being taken during the gap in the curve. Peptide concentrations are given in μM. Each curve indicates an independent experiment.

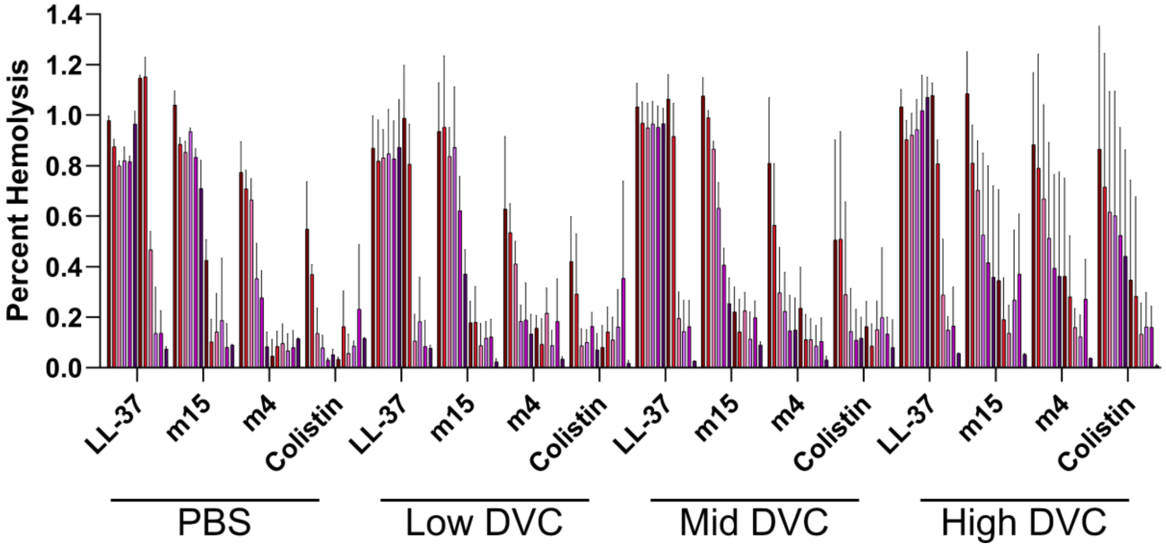
Percent Hemolysis
1.4
1.2
1.0
0.8
0.6
0.4
0.2
0.0
LL-37
m15
m4
Colistin
PBS
Low DVC
Mid DVC
High DVC

**Supplementary Figure 89 – Histogram depiction of Figure 5 data.** Endpoint hemolytic activity of each peptide against defibrinated sheep blood relative to a 100% hemolysis thrice freeze-thawed control after 18 hours at 37 °C in the presence of PBS or PBS supplemented with Low, Mid, or High concentrations of $Ca^{2+}$ / $Mg^{2+}$. Raw data were obtained by absorbance at 414 nm in well supernatants for free hemoglobin. Peptide concentrations are given in μM.

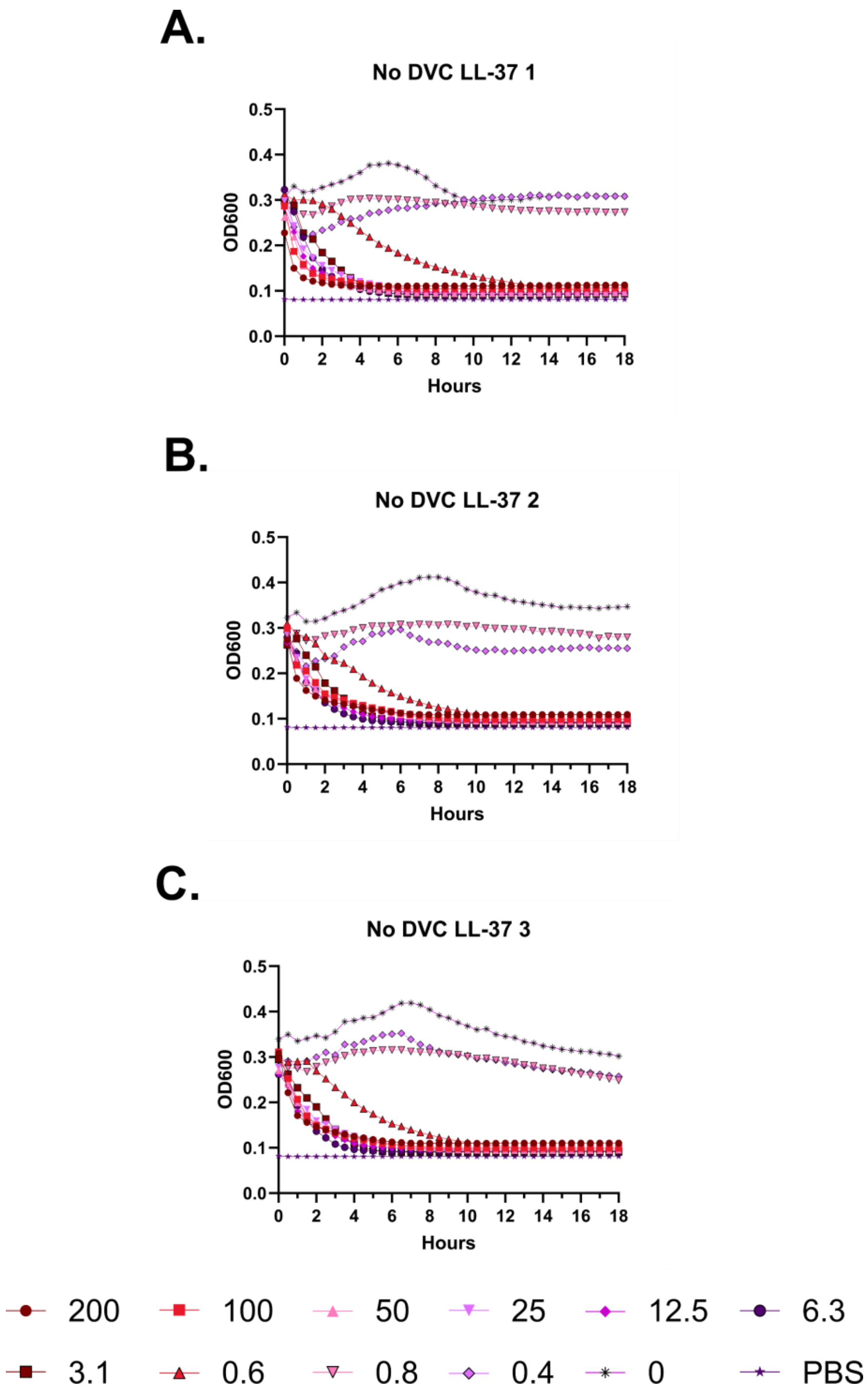
A.
No DVC LL-37 1
OD600
Hours
B.
No DVC LL-37 2
OD600
Hours
C.
No DVC LL-37 3
OD600
Hours
200
100
50
25
12.5
6.3
3.1
0.6
0.8
0.4
0
PBS

**Supplementary Figure 90 – Kinetic hemolysis assays for LL-37 in the presence of PBS.** Hemolysis was monitored by OD600 over 18 hours at 37 °C in the presence of the indicated peptide concentrations. Decreasing OD600 indicates increasing hemolysis. Peptide concentrations are given in μM. Each curve indicates an independent experiment.

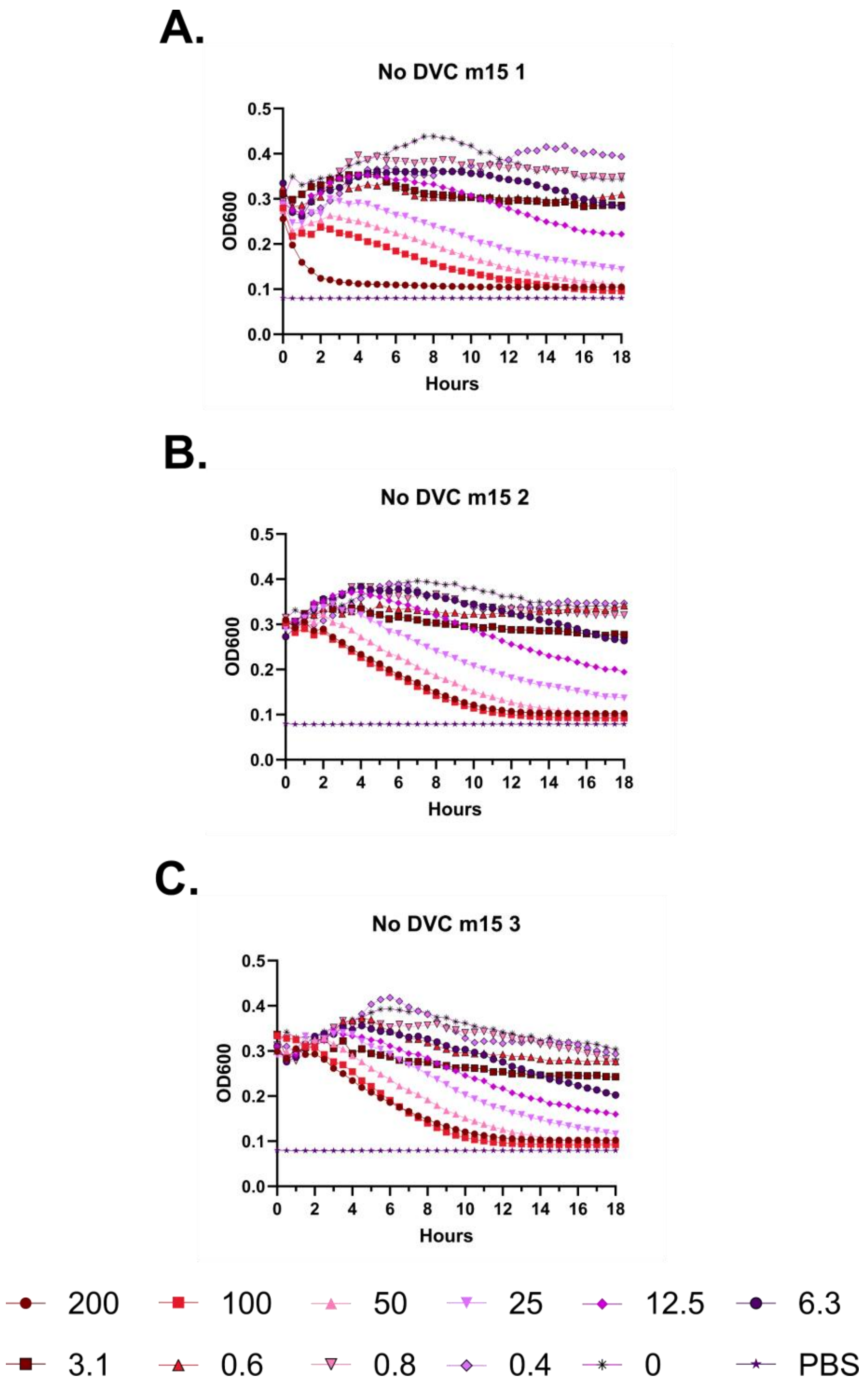
A.
No DVC m15 1
OD600
Hours
B.
No DVC m15 2
OD600
Hours
C.
No DVC m15 3
OD600
Hours
200
100
50
25
12.5
6.3
3.1
0.6
0.8
0.4
0
PBS

**Supplementary Figure 91 – Kinetic hemolysis assays for m15 in the presence of PBS.** Hemolysis was monitored by OD600 over 18 hours at 37 °C in the presence of the indicated peptide concentrations. Decreasing OD600 indicates increasing hemolysis. Peptide concentrations are given in μM. Each curve indicates an independent experiment.

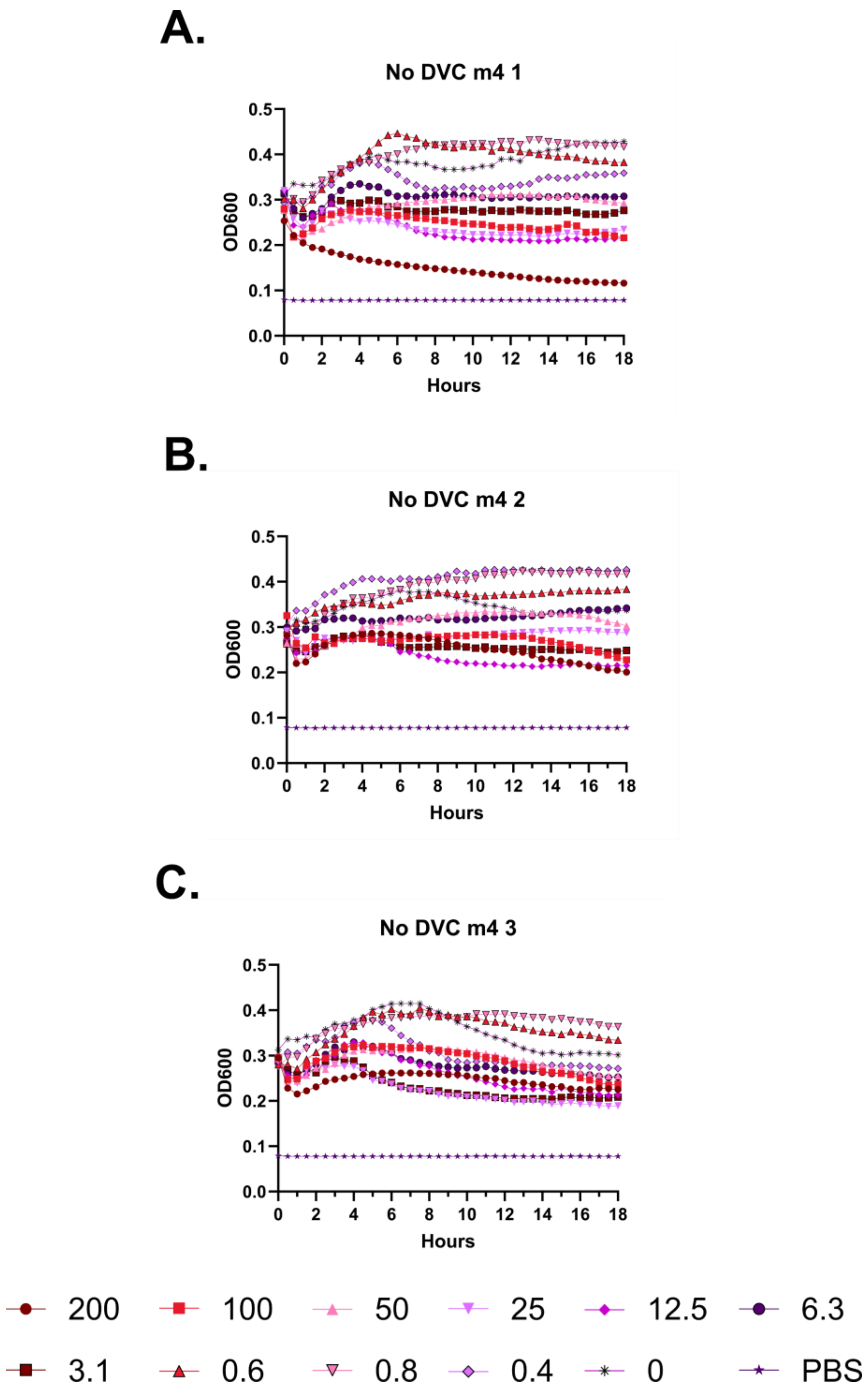
A.
No DVC m4 1
OD600
Hours
B.
No DVC m4 2
OD600
Hours
C.
No DVC m4 3
OD600
Hours
200
100
50
25
12.5
6.3
3.1
0.6
0.8
0.4
0
PBS

**Supplementary Figure 92 – Kinetic hemolysis assays for m4 in the presence of PBS.** Hemolysis was monitored by OD600 over 18 hours at 37 °C in the presence of the indicated peptide concentrations. Decreasing OD600 indicates increasing hemolysis. Peptide concentrations are given in μM. Each curve indicates an independent experiment.

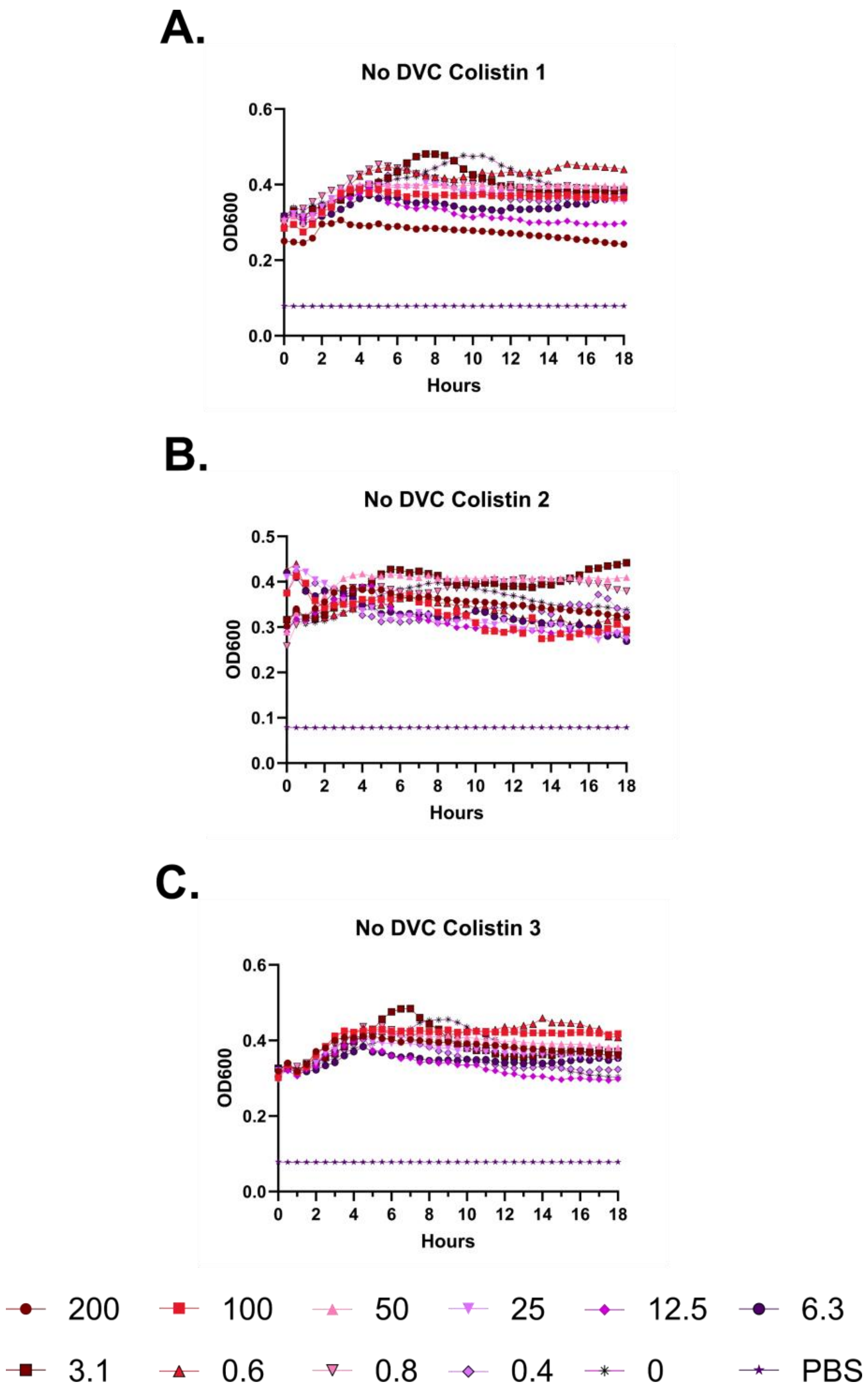
A.
No DVC Colistin 1
OD600
Hours
B.
No DVC Colistin 2
OD600
Hours
C.
No DVC Colistin 3
OD600
Hours
200
100
50
25
12.5
6.3
3.1
0.6
0.8
0.4
0
PBS

**Supplementary Figure 93 – Kinetic hemolysis assays for colistin in the presence of PBS.** Hemolysis was monitored by OD600 over 18 hours at 37 °C in the presence of the indicated peptide concentrations. Decreasing OD600 indicates increasing hemolysis. Peptide concentrations are given in μM. Each curve indicates an independent experiment.

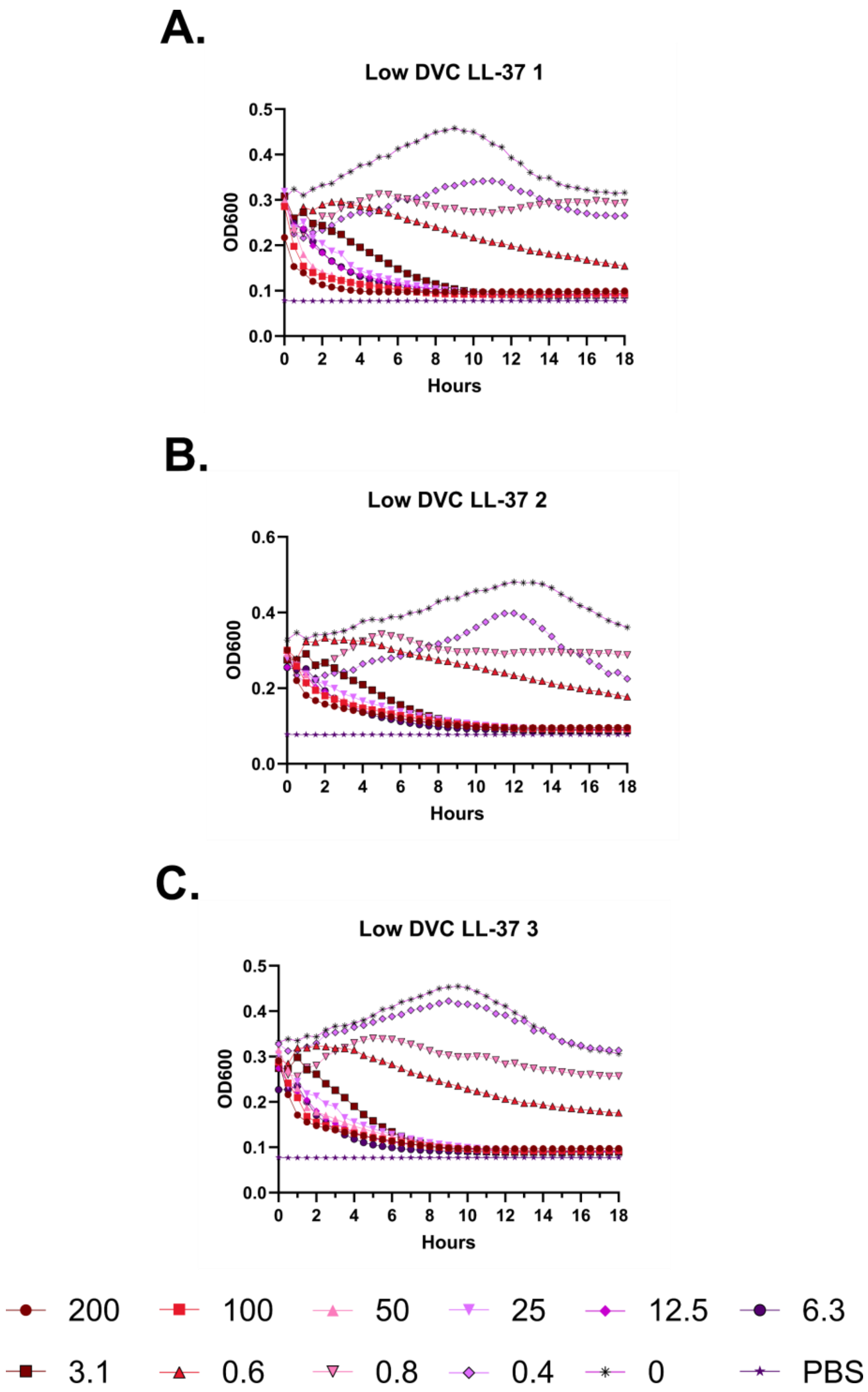
A.
Low DVC LL-37 1
0.5
0.4
0.3
0.2
0.1
0.0
OD600
0
2
4
6
8
10
12
14
16
18
Hours
B.
Low DVC LL-37 2
0.6
0.4
0.2
0.0
OD600
0
2
4
6
8
10
12
14
16
18
Hours
C.
Low DVC LL-37 3
0.5
0.4
0.3
0.2
0.1
0.0
OD600
0
2
4
6
8
10
12
14
16
18
Hours
200
100
50
25
12.5
6.3
3.1
0.6
0.8
0.4
0
PBS

**Supplementary Figure 94 – Kinetic hemolysis assays for LL-37 in the presence of PBS and Low Divalent Cations.** Hemolysis was monitored by OD600 over 18 hours at 37 °C in the presence of the indicated peptide concentrations. Decreasing OD600 indicates increasing hemolysis. Peptide concentrations are given in μM. Each curve indicates an independent experiment.

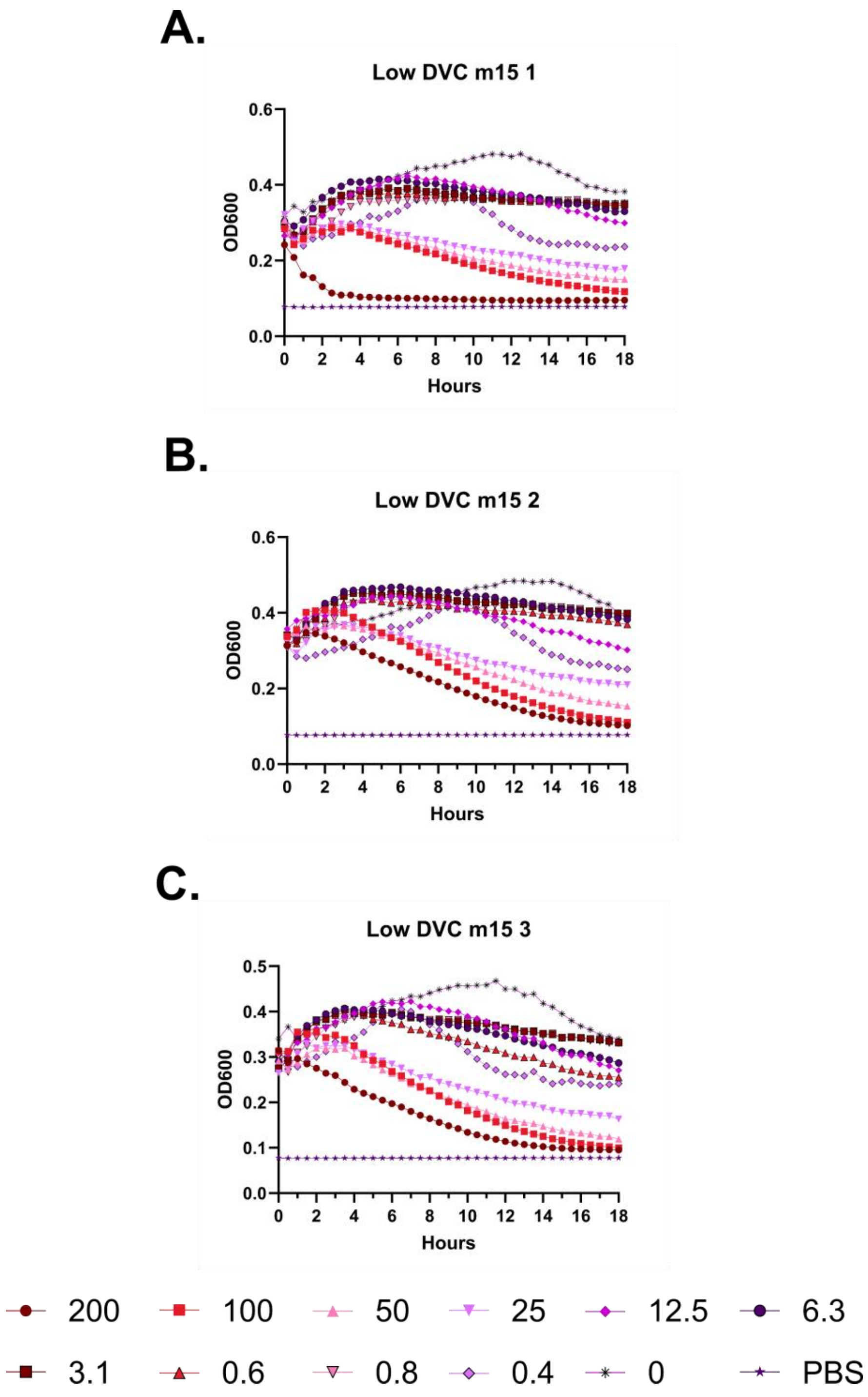
A.
Low DVC m15 1
0.6
0.4
0.2
0.0
OD600
0 2 4 6 8 10 12 14 16 18
Hours
B.
Low DVC m15 2
0.6
0.4
0.2
0.0
OD600
0 2 4 6 8 10 12 14 16 18
Hours
C.
Low DVC m15 3
0.5
0.4
0.3
0.2
0.1
0.0
OD600
0 2 4 6 8 10 12 14 16 18
Hours
200
100
50
25
12.5
6.3
3.1
0.6
0.8
0.4
0
PBS

**Supplementary Figure 95 – Kinetic hemolysis assays for m15 in the presence of PBS and Low Divalent Cations.** Hemolysis was monitored by OD600 over 18 hours at 37 °C in the presence of the indicated peptide concentrations. Decreasing OD600 indicates increasing hemolysis. Peptide concentrations are given in μM. Each curve indicates an independent experiment.

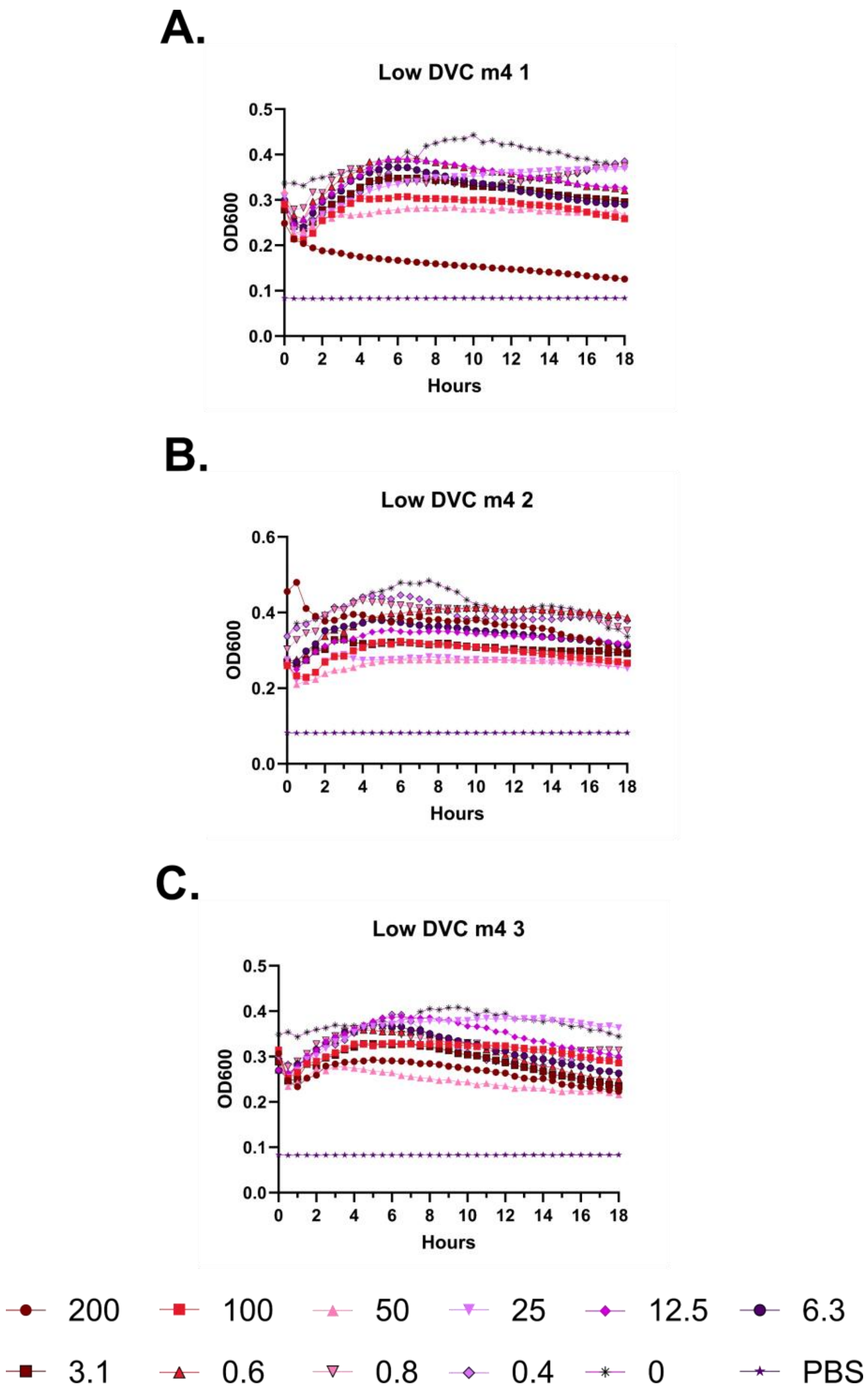
A.
Low DVC m4 1
OD600
Hours
B.
Low DVC m4 2
OD600
Hours
C.
Low DVC m4 3
OD600
Hours
200
100
50
25
12.5
6.3
3.1
0.6
0.8
0.4
0
PBS

**Supplementary Figure 96 – Kinetic hemolysis assays for m4 in the presence of PBS and Low Divalent Cations.** Hemolysis was monitored by OD600 over 18 hours at 37 °C in the presence of the indicated peptide concentrations. Decreasing OD600 indicates increasing hemolysis. Peptide concentrations are given in μM. Each curve indicates an independent experiment.

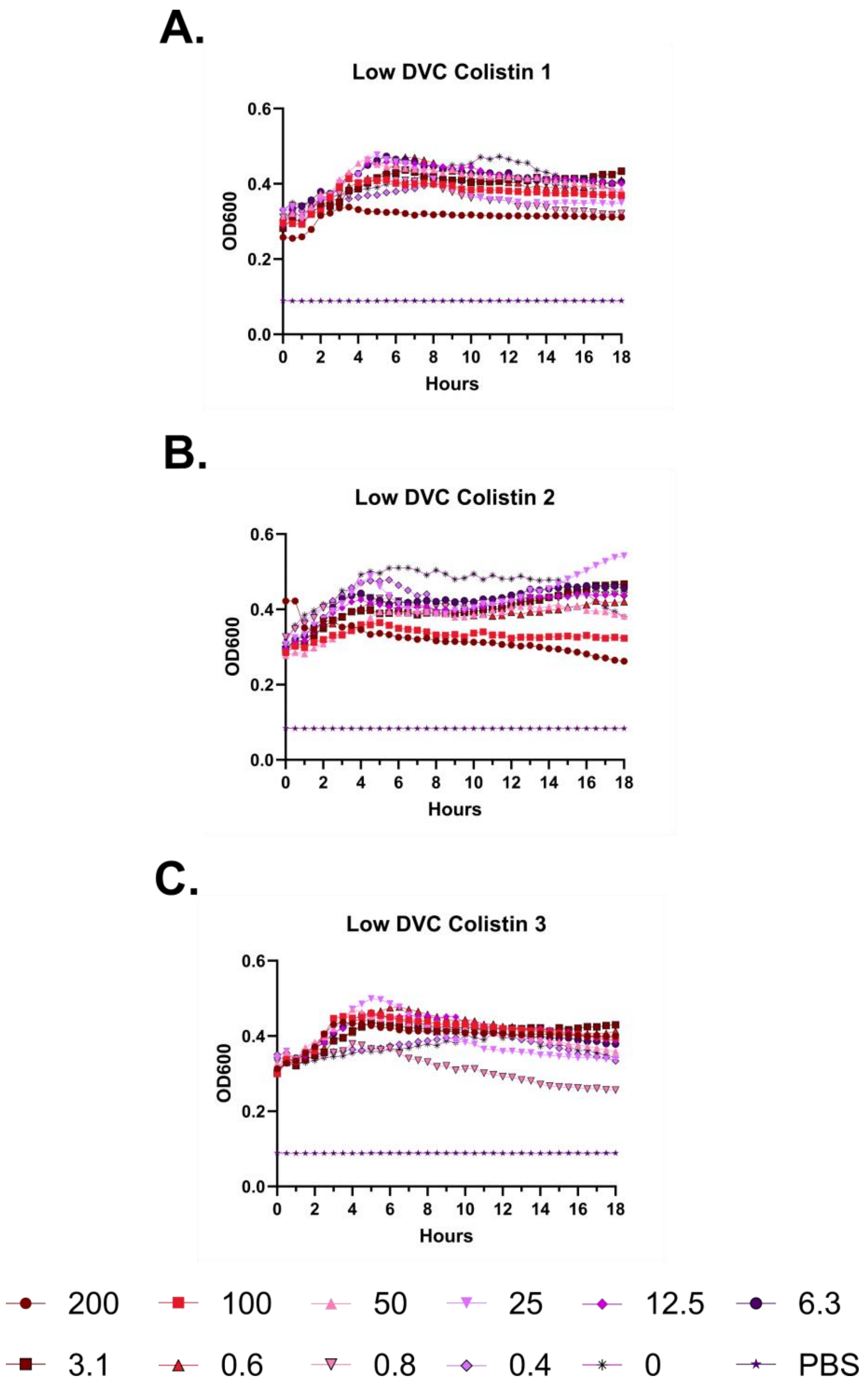
A.
Low DVC Colistin 1
OD600
Hours
0.0
0.2
0.4
0.6
0
2
4
6
8
10
12
14
16
18
B.
Low DVC Colistin 2
OD600
Hours
C.
Low DVC Colistin 3
OD600
Hours
200
100
50
25
12.5
6.3
3.1
0.6
0.8
0.4
0
PBS

**Supplementary Figure 97 – Kinetic hemolysis assays for colistin in the presence of PBS and Low Divalent Cations.** Hemolysis was monitored by OD600 over 18 hours at 37 °C in the presence of the indicated peptide concentrations. Decreasing OD600 indicates increasing hemolysis. Peptide concentrations are given in μM. Each curve indicates an independent experiment.

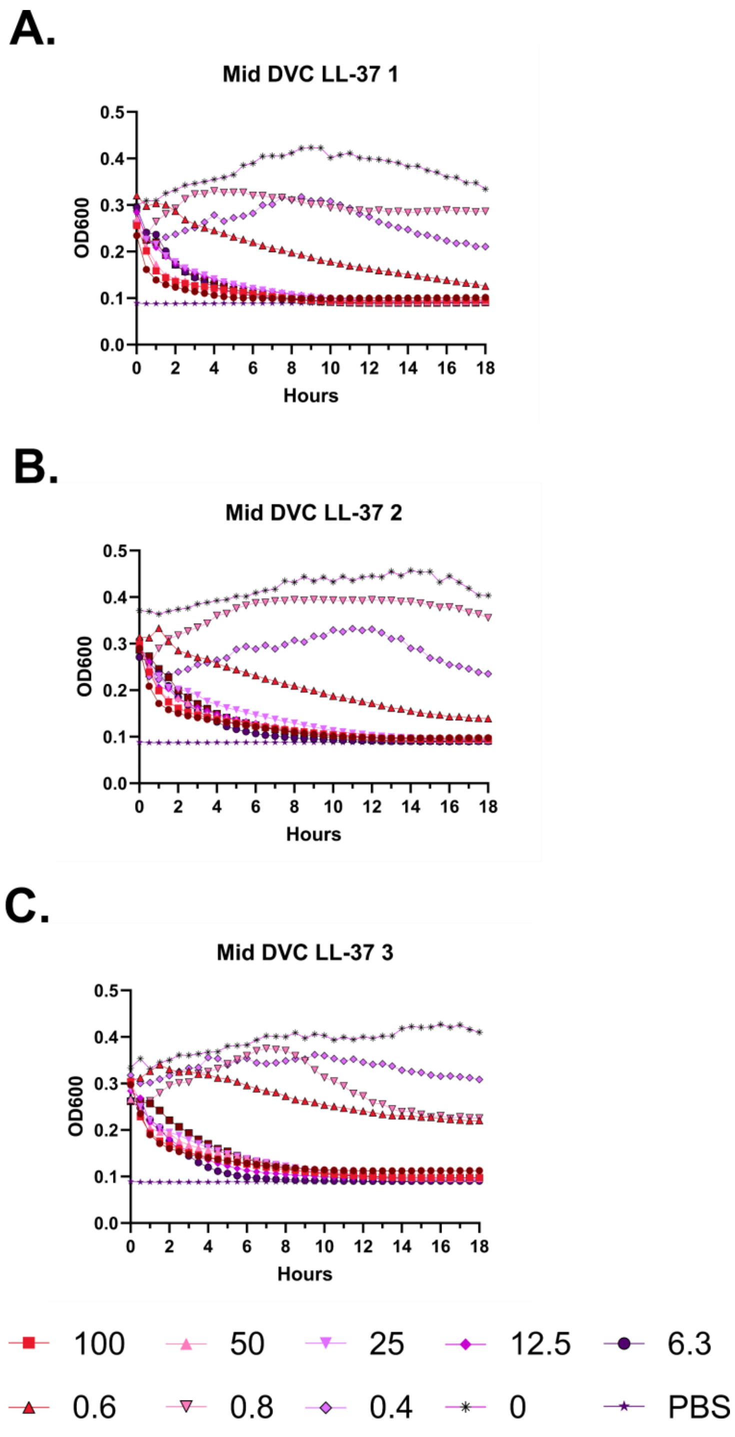
A.
Mid DVC LL-37 1
OD600
Hours
B.
Mid DVC LL-37 2
OD600
Hours
C.
Mid DVC LL-37 3
OD600
Hours
200
100
50
25
12.5
6.3
3.1
0.6
0.8
0.4
0
PBS

**Supplementary Figure 98 – Kinetic hemolysis assays for LL-37 in the presence of PBS and Mid Divalent Cations.** Hemolysis was monitored by OD600 over 18 hours at 37 °C in the presence of the indicated peptide concentrations. Decreasing OD600 indicates increasing hemolysis. Peptide concentrations are given in μM. Each curve indicates an independent experiment.

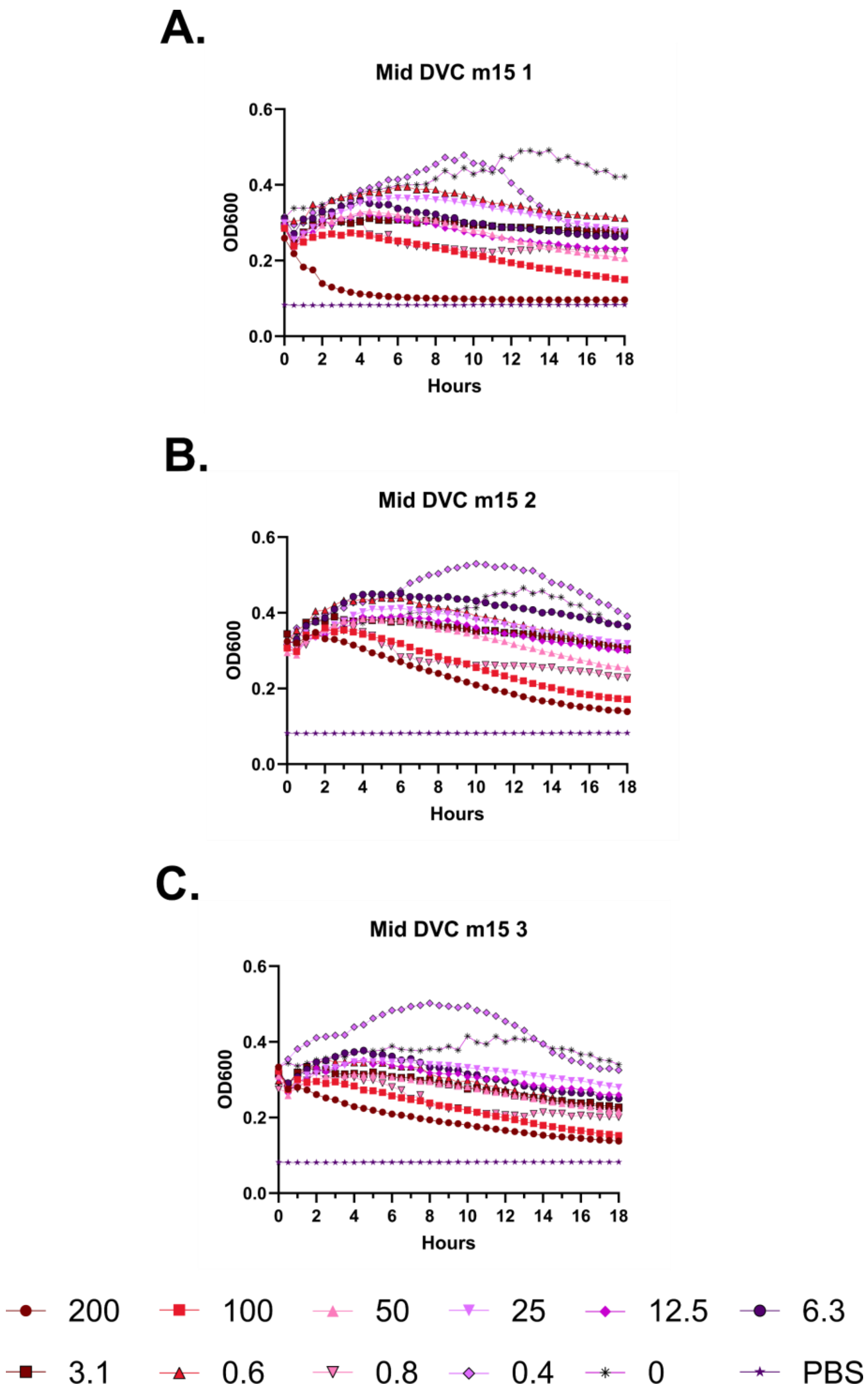
A.
Mid DVC m15 1
OD600
Hours
B.
Mid DVC m15 2
OD600
Hours
C.
Mid DVC m15 3
OD600
Hours
200
100
50
25
12.5
6.3
3.1
0.6
0.8
0.4
0
PBS

**Supplementary Figure 99 – Kinetic hemolysis assays for m15 in the presence of PBS and Mid Divalent Cations.** Hemolysis was monitored by OD600 over 18 hours at 37 °C in the presence of the indicated peptide concentrations. Decreasing OD600 indicates increasing hemolysis. Peptide concentrations are given in μM. Each curve indicates an independent experiment.

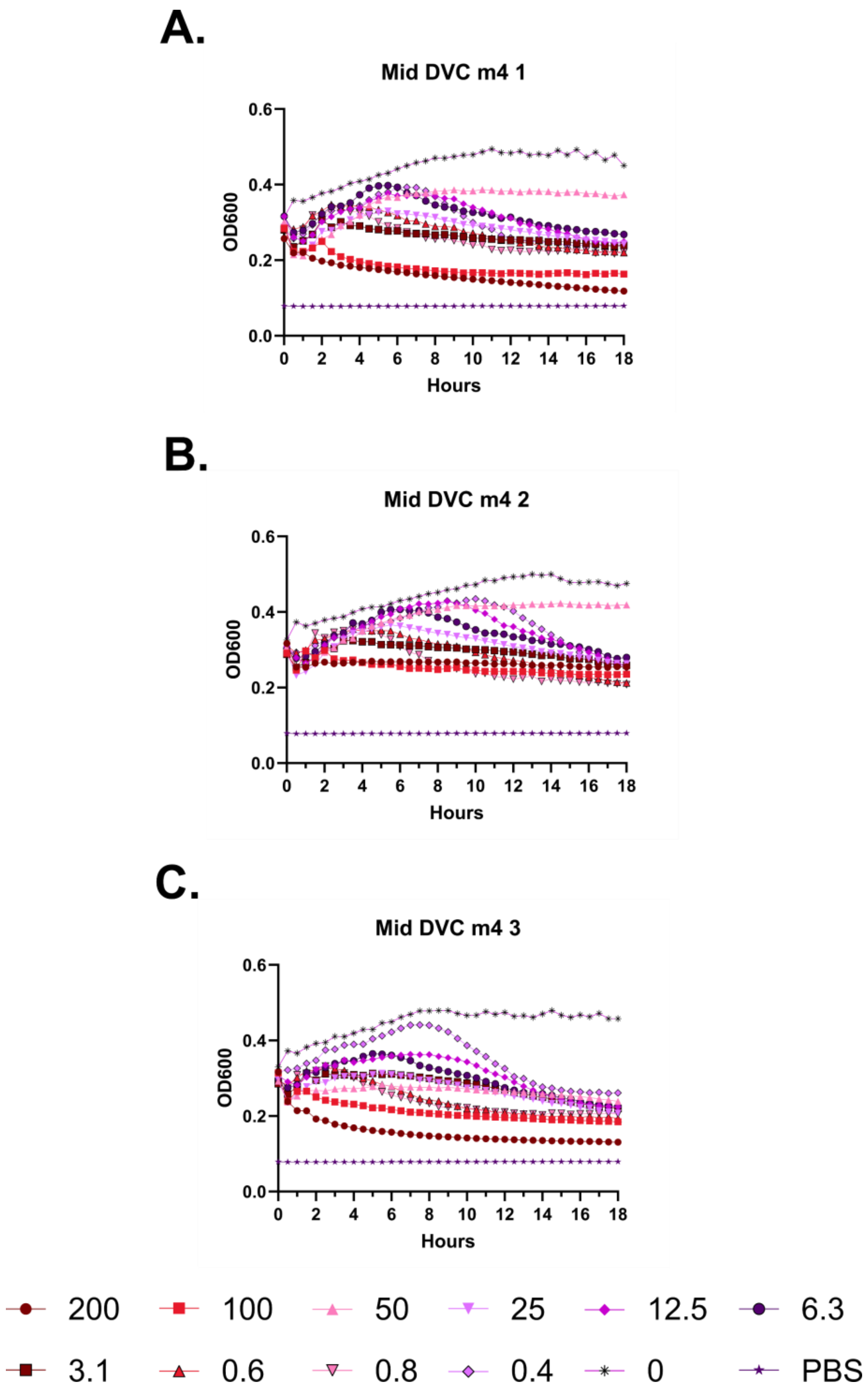
A.
Mid DVC m4 1
OD600
Hours
B.
Mid DVC m4 2
OD600
Hours
C.
Mid DVC m4 3
OD600
Hours
200
100
50
25
12.5
6.3
3.1
0.6
0.8
0.4
0
PBS

**Supplementary Figure 100 – Kinetic hemolysis assays for m4 in the presence of PBS and Mid Divalent Cations.** Hemolysis was monitored by OD600 over 18 hours at 37 °C in the presence of the indicated peptide concentrations. Decreasing OD600 indicates increasing hemolysis. Peptide concentrations are given in μM. Each curve indicates an independent experiment.

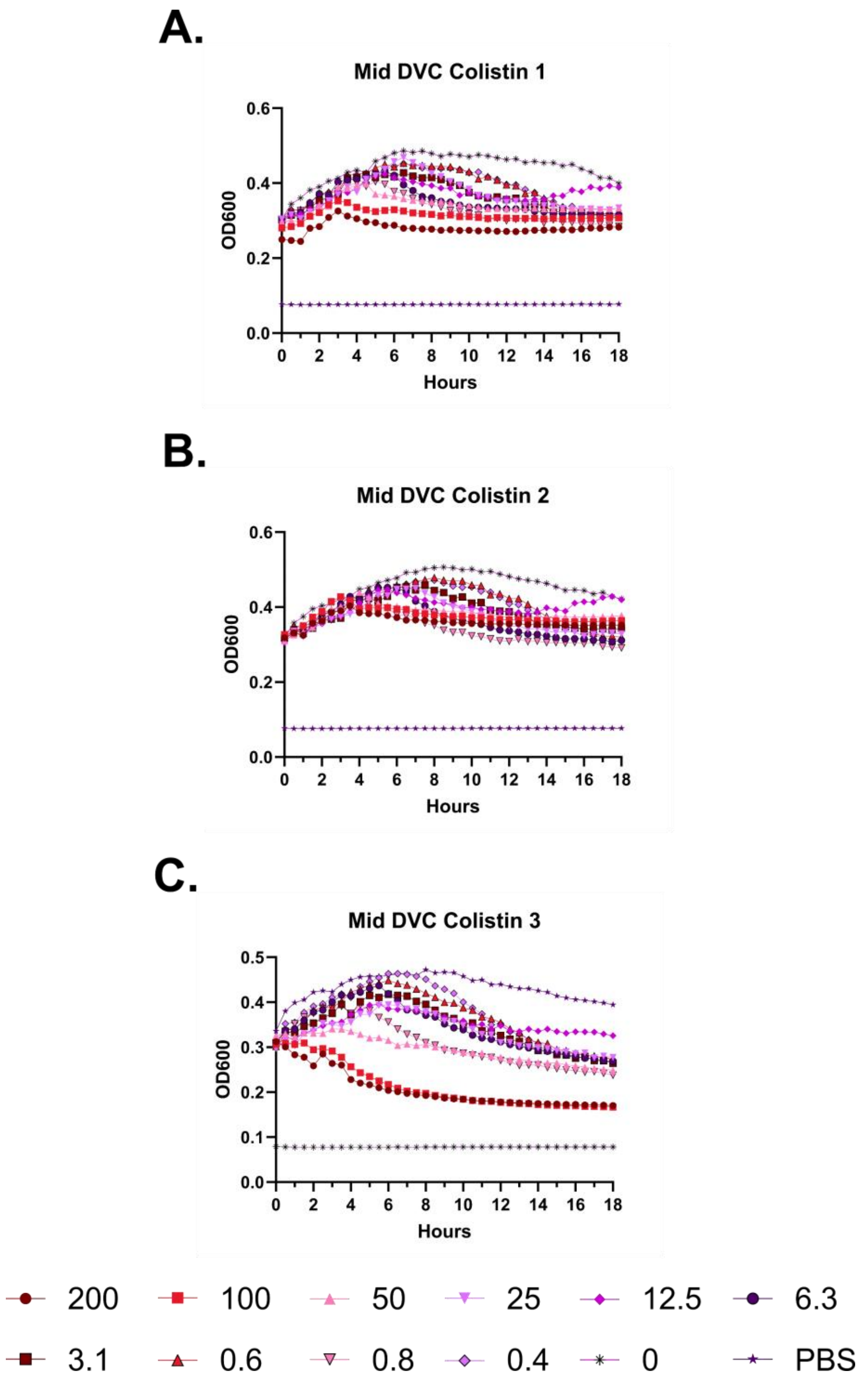
A.
Mid DVC Colistin 1
OD600
Hours
B.
Mid DVC Colistin 2
OD600
Hours
C.
Mid DVC Colistin 3
OD600
Hours
200
100
50
25
12.5
6.3
3.1
0.6
0.8
0.4
0
PBS

**Supplementary Figure 101 – Kinetic hemolysis assays for colistin in the presence of PBS and Mid Divalent Cations.** Hemolysis was monitored by OD600 over 18 hours at 37 °C in the presence of the indicated peptide concentrations. Decreasing OD600 indicates increasing hemolysis. Peptide concentrations are given in μM. Each curve indicates an independent experiment.

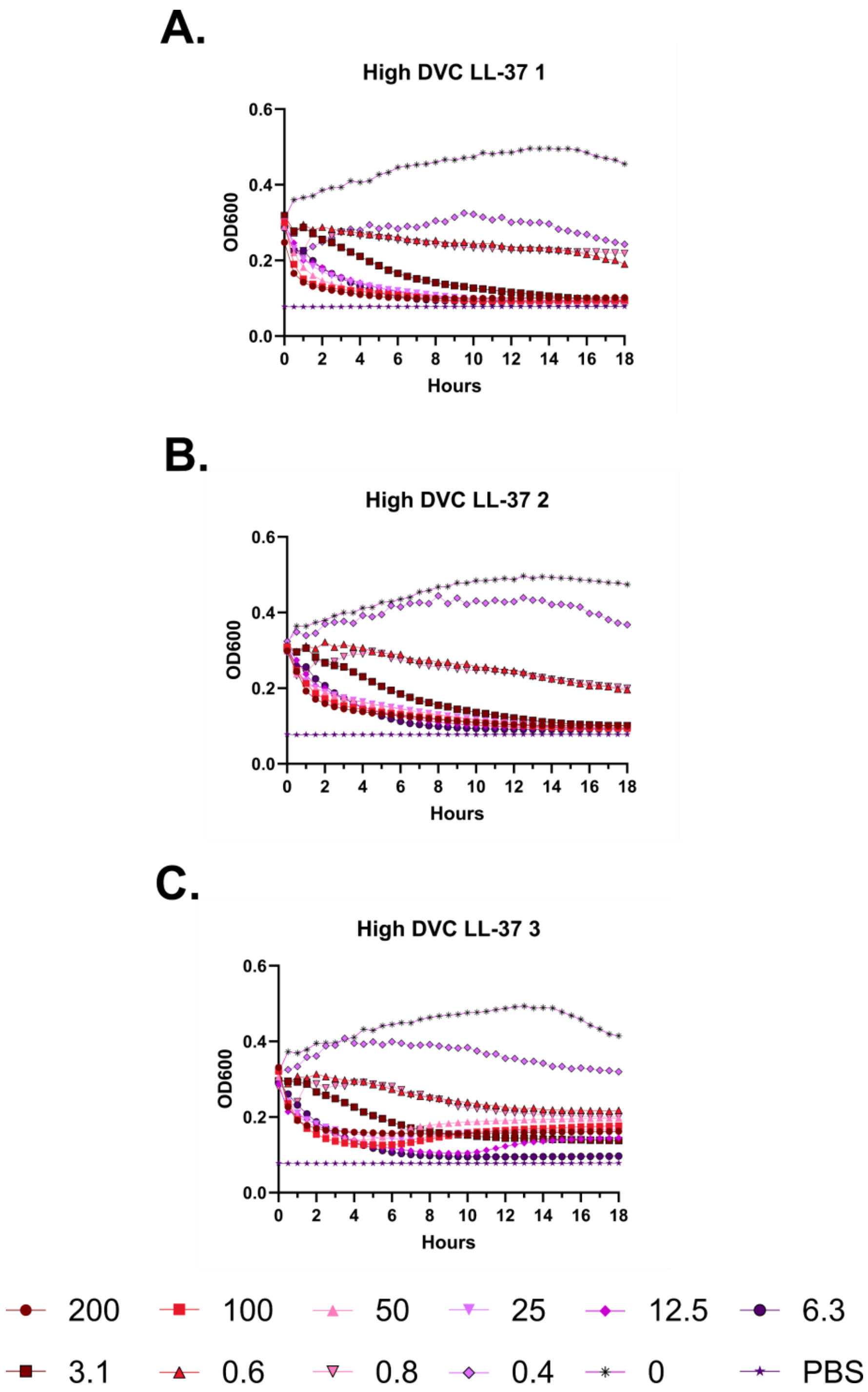
A.
High DVC LL-37 1
B.
High DVC LL-37 2
C.
High DVC LL-37 3
OD600
0.0
0.2
0.4
0.6
Hours
0 2 4 6 8 10 12 14 16 18
200
100
50
25
12.5
6.3
3.1
0.6
0.8
0.4
0
PBS

**Supplementary Figure 102 – Kinetic hemolysis assays for LL-37 in the presence of PBS and High Divalent Cations.** Hemolysis was monitored by OD600 over 18 hours at 37 °C in the presence of the indicated peptide concentrations. Decreasing OD600 indicates increasing hemolysis. Peptide concentrations are given in μM. Each curve indicates an independent experiment.

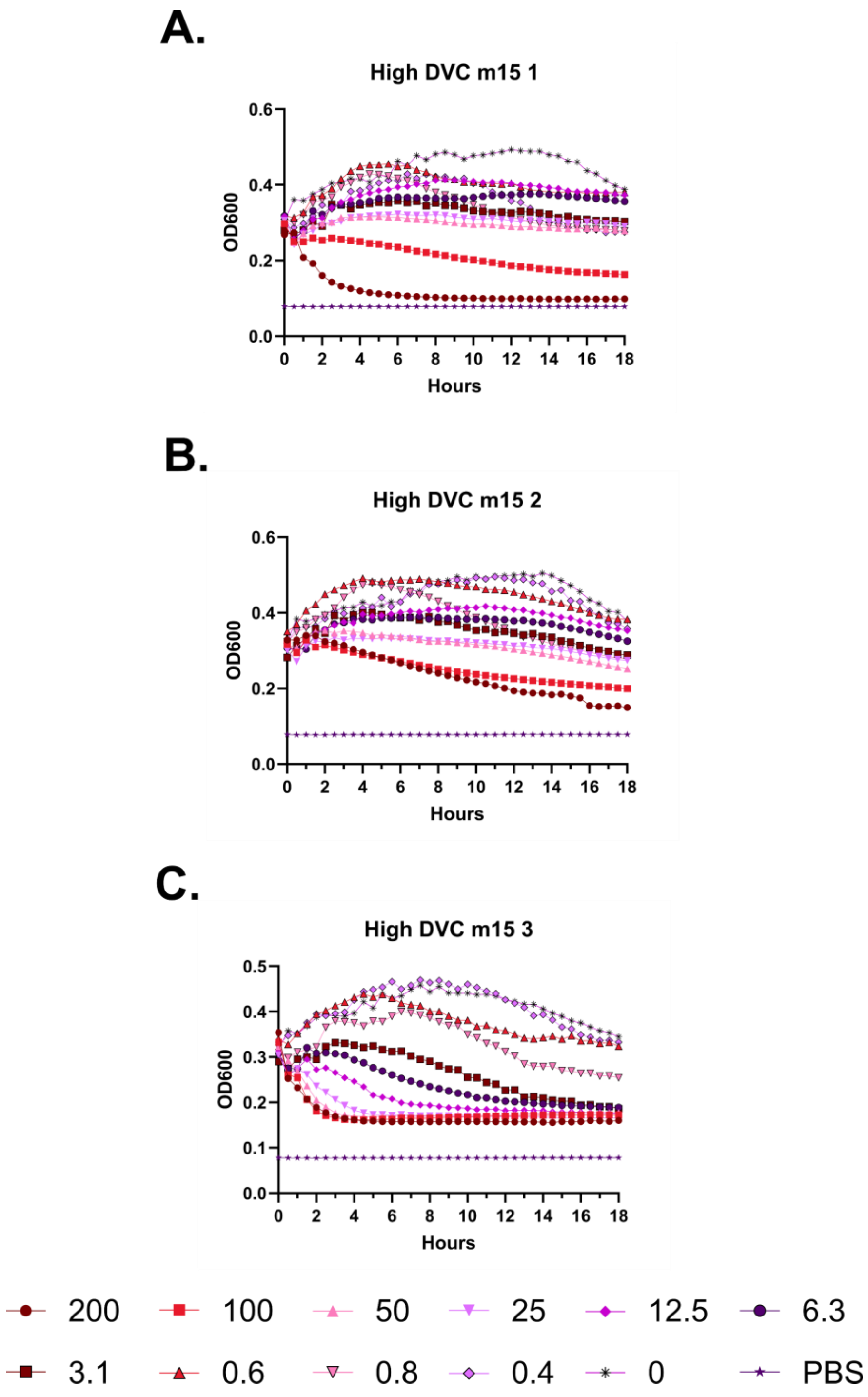
A.
High DVC m15 1
OD600
Hours
B.
High DVC m15 2
OD600
Hours
C.
High DVC m15 3
OD600
Hours
200
100
50
25
12.5
6.3
3.1
0.6
0.8
0.4
0
PBS

**Supplementary Figure 103 – Kinetic hemolysis assays for m15 in the presence of PBS and High Divalent Cations.** Hemolysis was monitored by OD600 over 18 hours at 37 °C in the presence of the indicated peptide concentrations. Decreasing OD600 indicates increasing hemolysis. Peptide concentrations are given in μM. Each curve indicates an independent experiment.

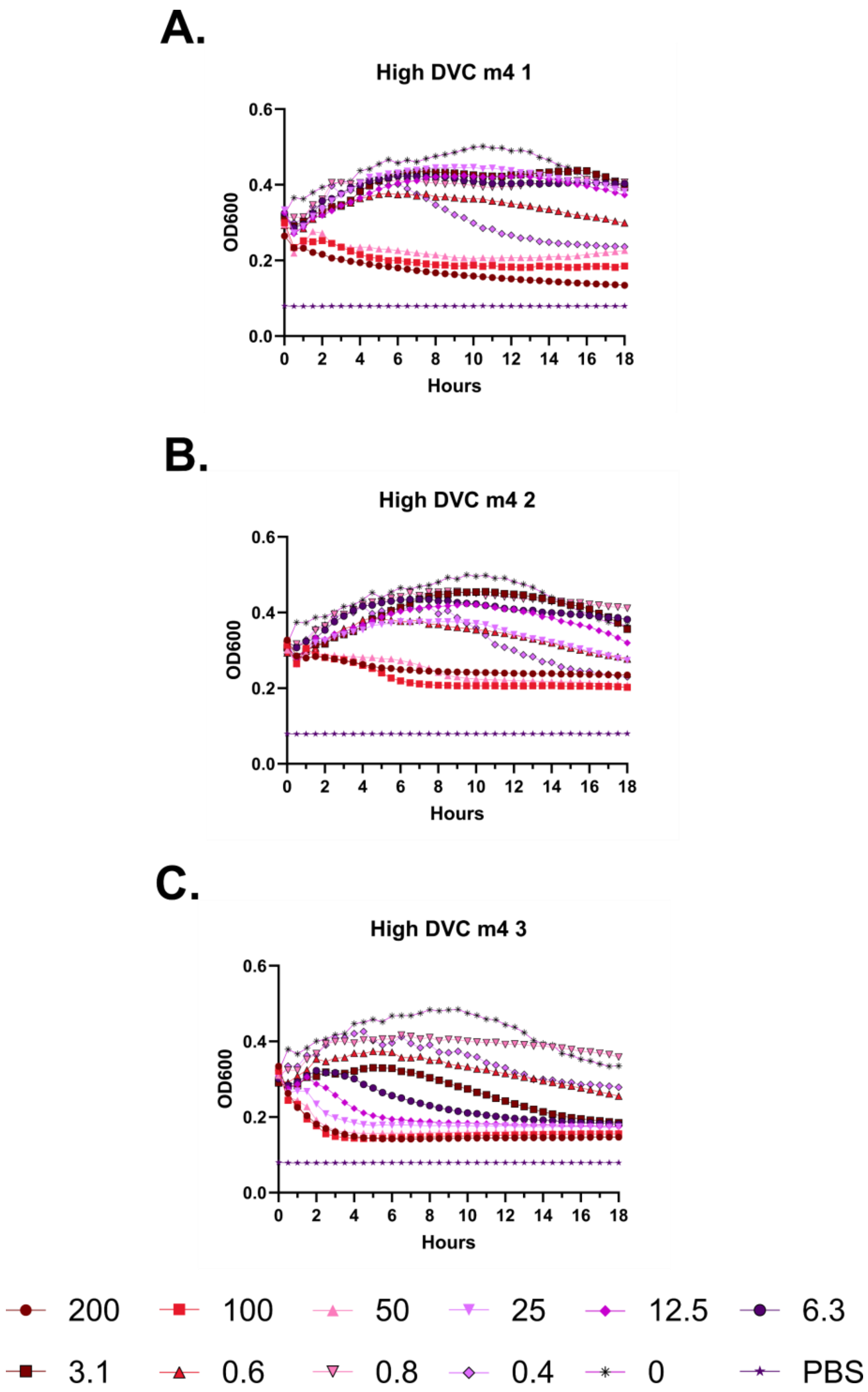
A.
High DVC m4 1
OD600
Hours
0.0
0.2
0.4
0.6
0 2 4 6 8 10 12 14 16 18
B.
High DVC m4 2
OD600
Hours
0.0
0.2
0.4
0.6
0 2 4 6 8 10 12 14 16 18
C.
High DVC m4 3
OD600
Hours
0.0
0.2
0.4
0.6
0 2 4 6 8 10 12 14 16 18
200
100
50
25
12.5
6.3
3.1
0.6
0.8
0.4
0
PBS

**Supplementary Figure 104 – Kinetic hemolysis assays for m4 in the presence of PBS and High Divalent Cations.** Hemolysis was monitored by OD600 over 18 hours at 37 °C in the presence of the indicated peptide concentrations. Decreasing OD600 indicates increasing hemolysis. Peptide concentrations are given in μM. Each curve indicates an independent experiment.

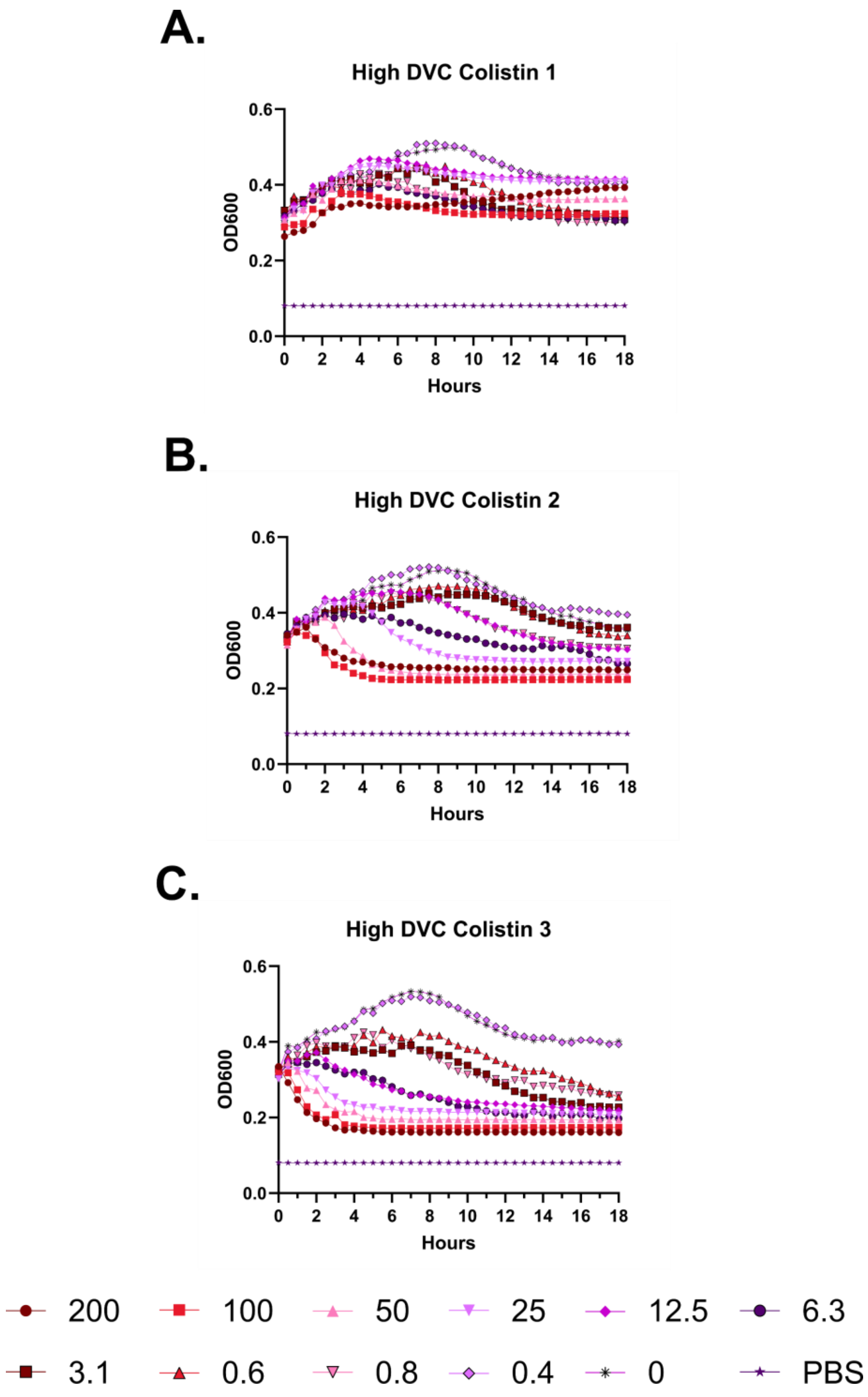
A.
High DVC Colistin 1
OD600
0.0
0.2
0.4
0.6
0 2 4 6 8 10 12 14 16 18
Hours
B.
High DVC Colistin 2
OD600
0.0
0.2
0.4
0.6
0 2 4 6 8 10 12 14 16 18
Hours
C.
High DVC Colistin 3
OD600
0.0
0.2
0.4
0.6
0 2 4 6 8 10 12 14 16 18
Hours
200
100
50
25
12.5
6.3
3.1
0.6
0.8
0.4
0
PBS

**Supplementary Figure 105 – Kinetic hemolysis assays for colistin in the presence of PBS and High Divalent Cations.** Hemolysis was monitored by OD600 over 18 hours at 37 °C in the presence of the indicated peptide concentrations. Decreasing OD600 indicates increasing hemolysis. Peptide concentrations are given in μM. Each curve indicates an independent experiment.

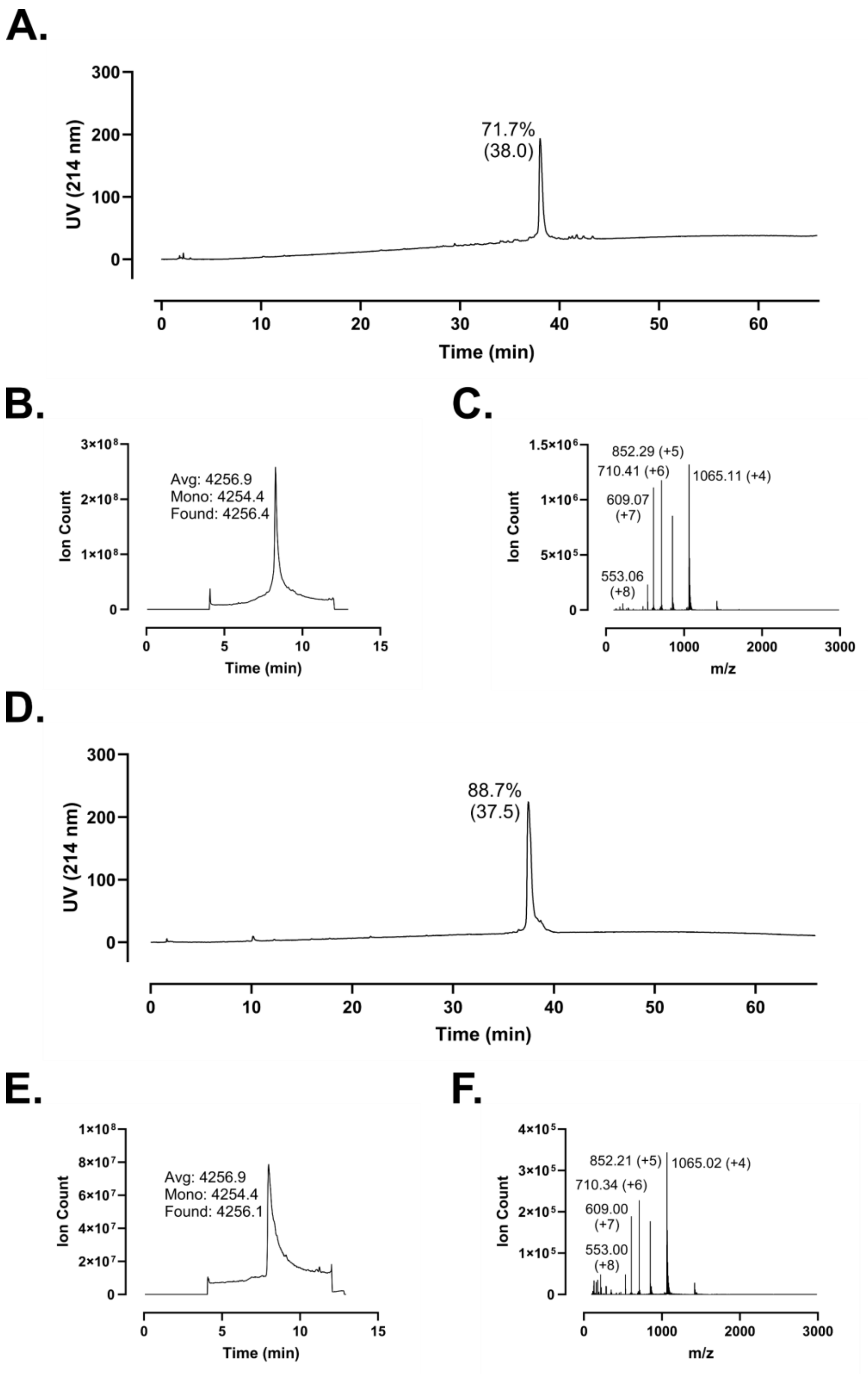
A.
UV (214 nm)
Time (min)
71.7%
(38.0)
B.
Ion Count
Time (min)
Avg: 4256.9
Mono: 4254.4
Found: 4256.4
C.
Ion Count
m/z
852.29 (+5)
710.41 (+6)
1065.11 (+4)
609.07
(+7)
553.06
(+8)
D.
UV (214 nm)
Time (min)
88.7%
(37.5)
E.
Ion Count
Time (min)
Avg: 4256.9
Mono: 4254.4
Found: 4256.1
F.
Ion Count
m/z
852.21 (+5)
1065.02 (+4)
710.34 (+6)
609.00
(+7)
553.00
(+8)

**Supplementary Figure 106 – Analytical data associated with LL-37 m1 synthesis. A.** HPLC of crude peptide. Purity by trace integration is shown along with retention time in parentheses. **B.** Total ion chromatogram of crude peptide, including the average, monoisotopic, and observed masses. **C.** Mass spectrum at peak of **B.** with selected *m/z* labelled. **D.** HPLC of peptide following a single round of purification. Peak labeling as in **A. E.** Total ion chromatogram for purified peptide. **F.** Mass spectrum at the peak of **E.**

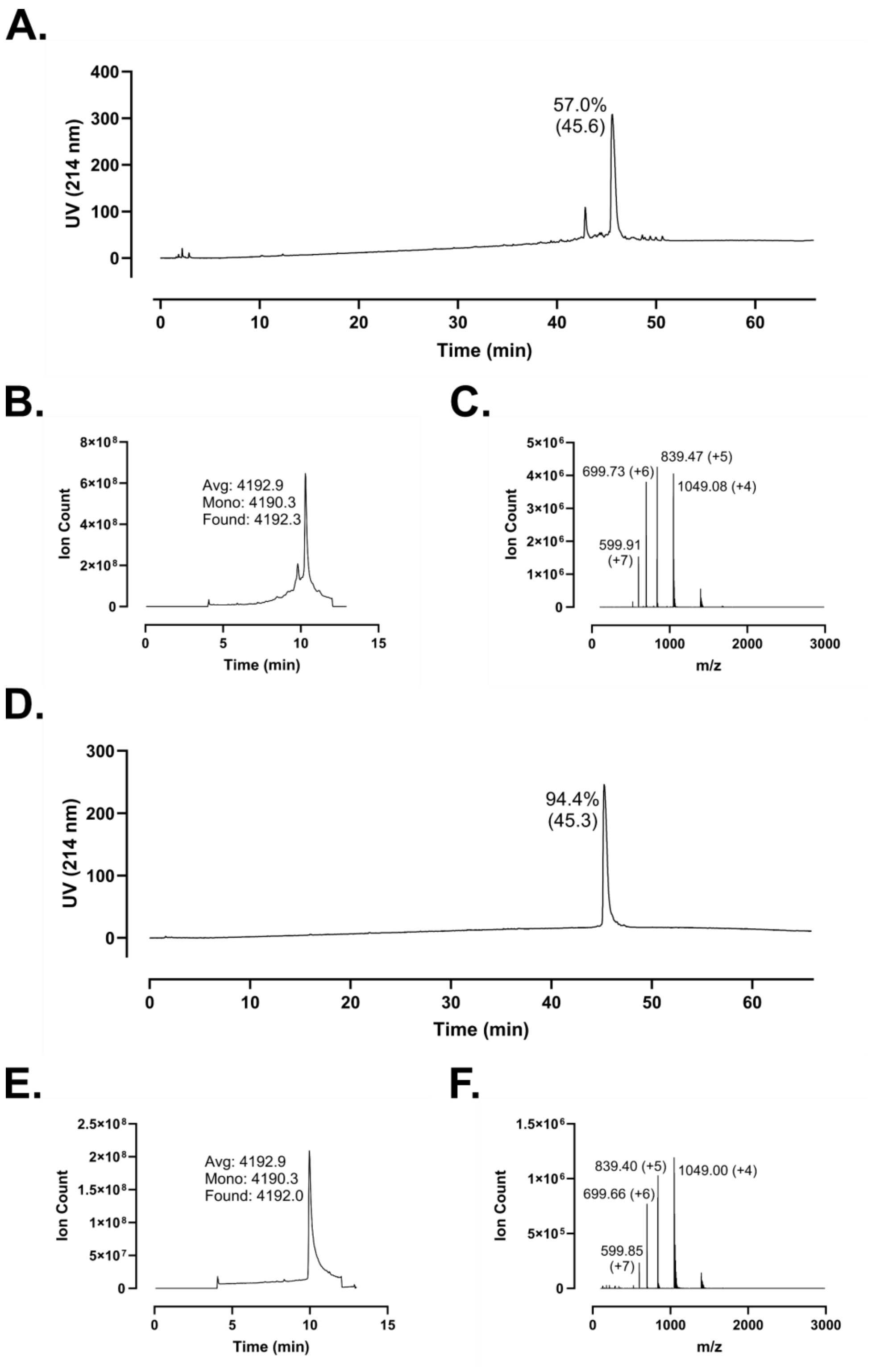
A.
UV (214 nm)
Time (min)
57.0%
(45.6)
B.
Ion Count
Time (min)
Avg: 4192.9
Mono: 4190.3
Found: 4192.3
C.
Ion Count
m/z
699.73 (+6)
839.47 (+5)
1049.08 (+4)
599.91
(+7)
D.
UV (214 nm)
Time (min)
94.4%
(45.3)
E.
Ion Count
Time (min)
Avg: 4192.9
Mono: 4190.3
Found: 4192.0
F.
Ion Count
m/z
839.40 (+5)
1049.00 (+4)
699.66 (+6)
599.85
(+7)

**Supplementary Figure 107 – Analytical data associated with LL-37 m2 synthesis. A.** HPLC of crude peptide. Purity by trace integration is shown along with retention time in parentheses. **B.** Total ion chromatogram of crude peptide, including the average, monoisotopic, and observed masses. **C.** Mass spectrum at peak of **B.** with selected *m/z* labelled. **D.** HPLC of peptide following a single round of purification. Peak labeling as in **A. E.** Total ion chromatogram for purified peptide. **F.** Mass spectrum at the peak of **E.**

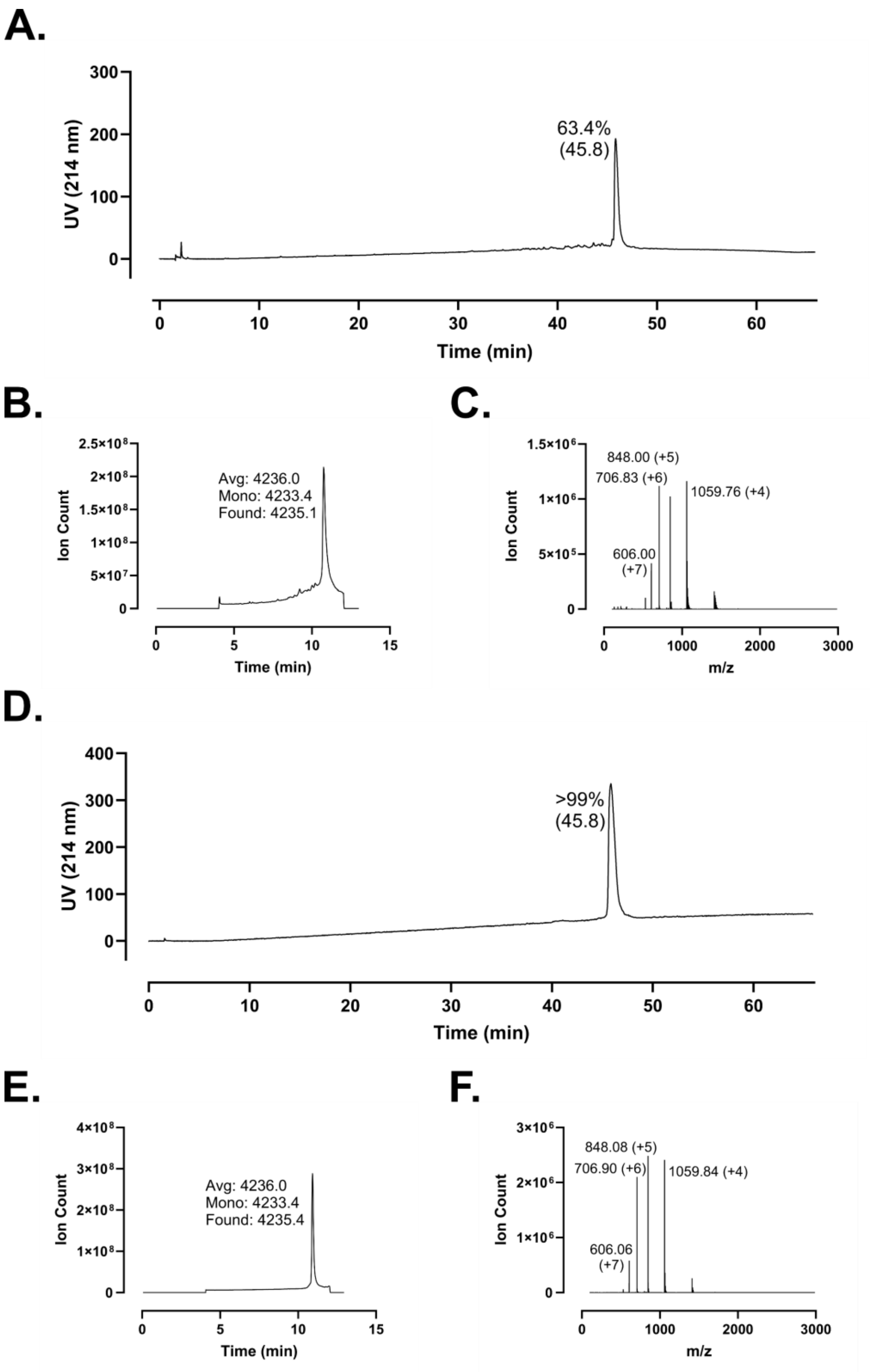
A.
UV (214 nm)
Time (min)
63.4%
(45.8)
B.
Ion Count
Time (min)
Avg: 4236.0
Mono: 4233.4
Found: 4235.1
C.
Ion Count
m/z
848.00 (+5)
706.83 (+6)
1059.76 (+4)
606.00
(+7)
D.
UV (214 nm)
Time (min)
>99%
(45.8)
E.
Ion Count
Time (min)
Avg: 4236.0
Mono: 4233.4
Found: 4235.4
F.
Ion Count
m/z
848.08 (+5)
706.90 (+6)
1059.84 (+4)
606.06
(+7)

**Supplementary Figure 108 – Analytical data associated with LL-37 m3 synthesis. A.** HPLC of crude peptide. Purity by trace integration is shown along with retention time in parentheses. **B.** Total ion chromatogram of crude peptide, including the average, monoisotopic, and observed masses. **C.** Mass spectrum at peak of **B.** with selected *m/z* labelled. **D.** HPLC of peptide following a single round of purification. Peak labeling as in **A. E.** Total ion chromatogram for purified peptide. **F.** Mass spectrum at the peak of **E.**

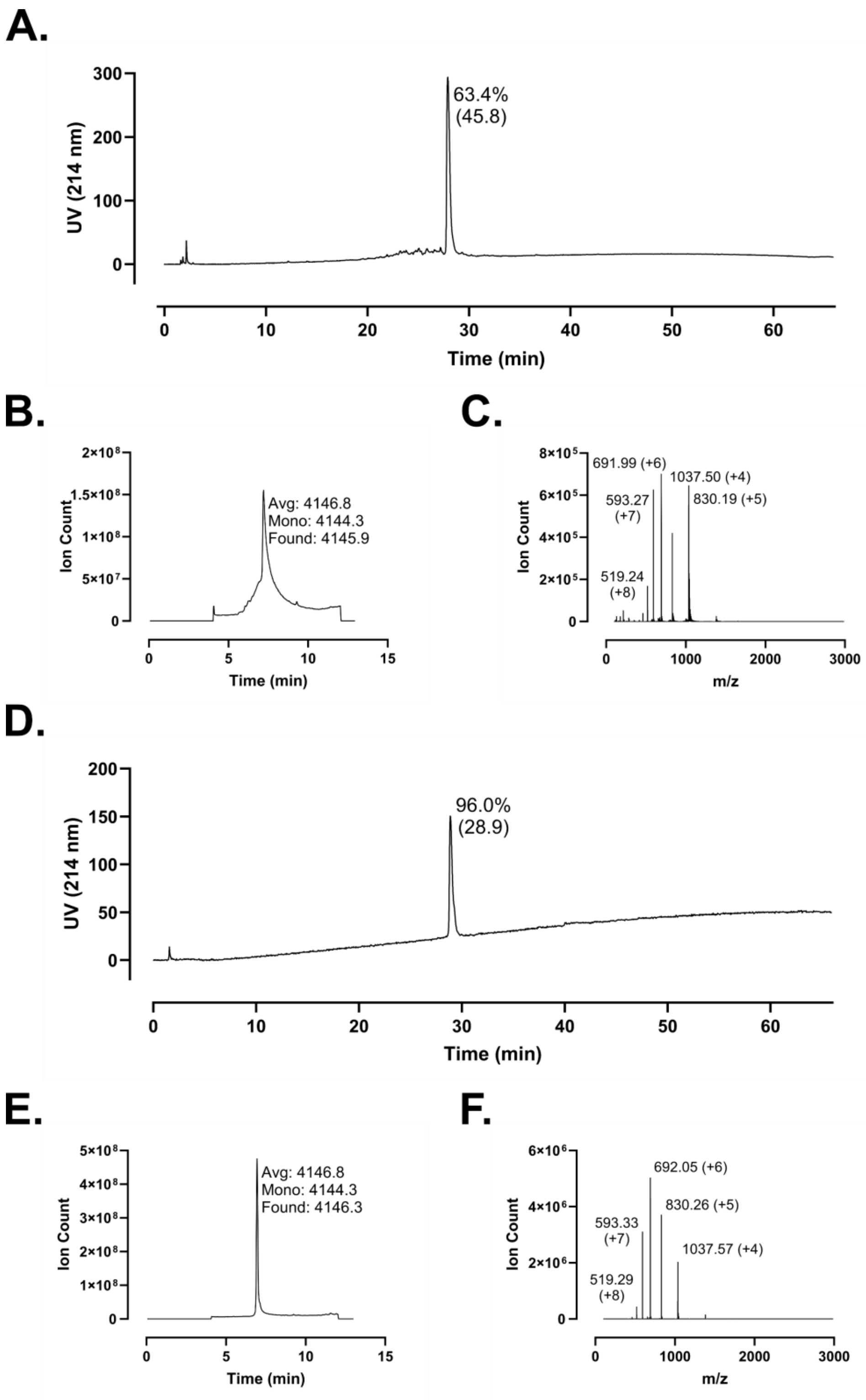
A.
UV (214 nm)
300
200
100
0
63.4%
(45.8)
Time (min)
0
10
20
30
40
50
60
B.
Ion Count
2×10⁸
1.5×10⁸
1×10⁸
5×10⁷
0
Avg: 4146.8
Mono: 4144.3
Found: 4145.9
Time (min)
0
5
10
15
C.
Ion Count
8×10⁵
6×10⁵
4×10⁵
2×10⁵
0
691.99 (+6)
1037.50 (+4)
593.27
(+7)
830.19 (+5)
519.24
(+8)
m/z
0
1000
2000
3000
D.
UV (214 nm)
200
150
100
50
0
96.0%
(28.9)
Time (min)
0
10
20
30
40
50
60
E.
Ion Count
5×10⁸
4×10⁸
3×10⁸
2×10⁸
1×10⁸
0
Avg: 4146.8
Mono: 4144.3
Found: 4146.3
Time (min)
0
5
10
15
F.
Ion Count
6×10⁶
4×10⁶
2×10⁶
0
692.05 (+6)
830.26 (+5)
593.33
(+7)
1037.57 (+4)
519.29
(+8)
m/z
0
1000
2000
3000

**Supplementary Figure 109 – Analytical data associated with LL-37 m4 synthesis. A.** HPLC of crude peptide. Purity by trace integration is shown along with retention time in parentheses. **B.** Total ion chromatogram of crude peptide, including the average, monoisotopic, and observed masses. **C.** Mass spectrum at peak of **B.** with selected *m/z* labelled. **D.** HPLC of peptide following a single round of purification. Peak labeling as in **A. E.** Total ion chromatogram for purified peptide. **F.** Mass spectrum at the peak of **E.** The synthesis shown here was used in multiple projects. The data presented here are thus identical to those shown in Reference 31 S154.

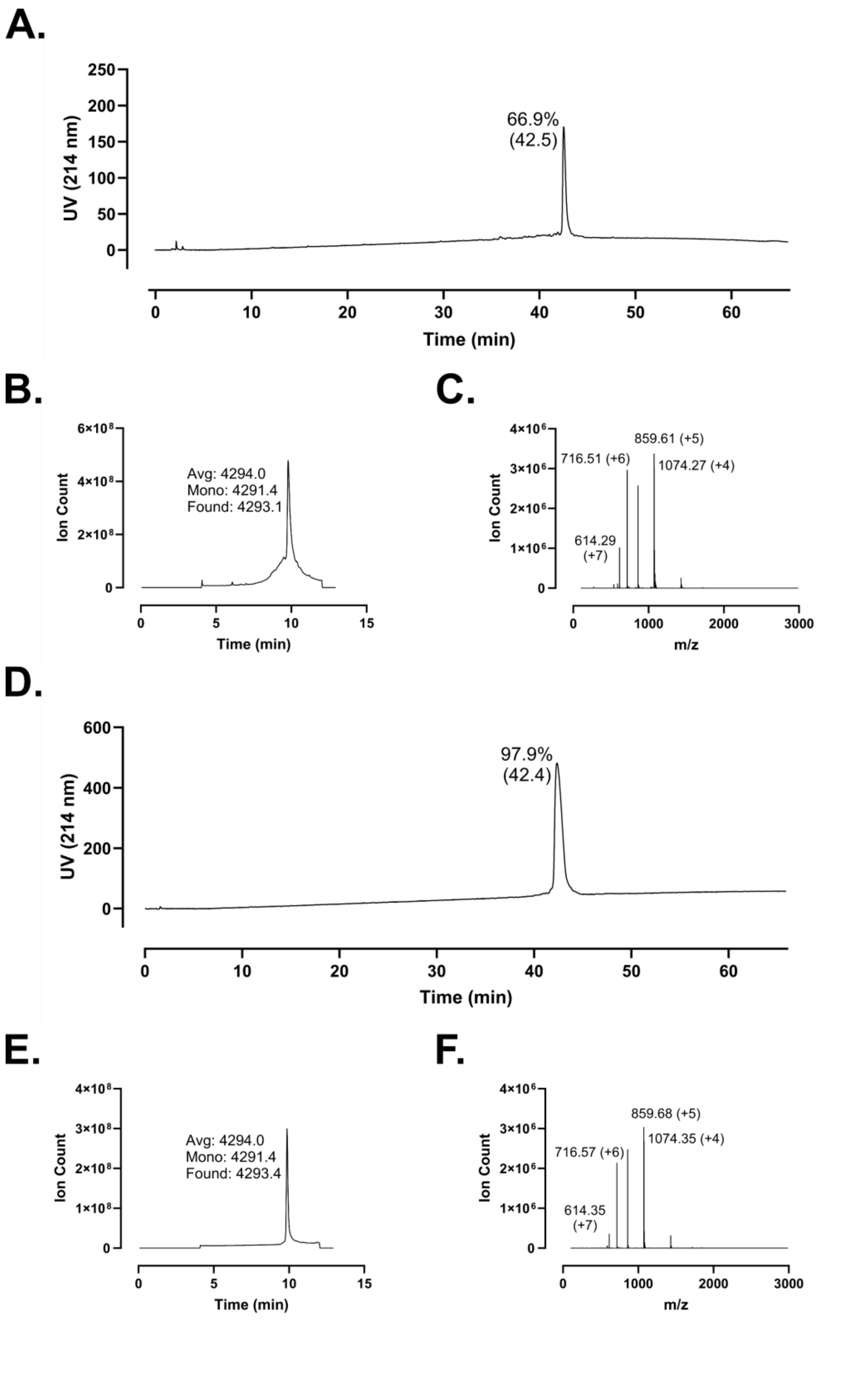
A.
UV (214 nm)
66.9%
(42.5)
Time (min)
B.
Ion Count
Avg: 4294.0
Mono: 4291.4
Found: 4293.1
Time (min)
C.
Ion Count
859.61 (+5)
716.51 (+6)
1074.27 (+4)
614.29
(+7)
m/z
D.
UV (214 nm)
97.9%
(42.4)
Time (min)
E.
Ion Count
Avg: 4294.0
Mono: 4291.4
Found: 4293.4
Time (min)
F.
Ion Count
859.68 (+5)
716.57 (+6)
1074.35 (+4)
614.35
(+7)
m/z

**Supplementary Figure 110 – Analytical data associated with LL-37 m5 synthesis. A.** HPLC of crude peptide. Purity by trace integration is shown along with retention time in parentheses. **B.** Total ion chromatogram of crude peptide, including the average, monoisotopic, and observed masses. **C.** Mass spectrum at peak of **B.** with selected *m/z* labelled. **D.** HPLC of peptide following a single round of purification. Peak labeling as in **A. E.** Total ion chromatogram for purified peptide. **F.** Mass spectrum at the peak of **E.** The synthesis shown here was used in multiple projects. The data presented here are thus identical to those shown in Reference 31 S155.

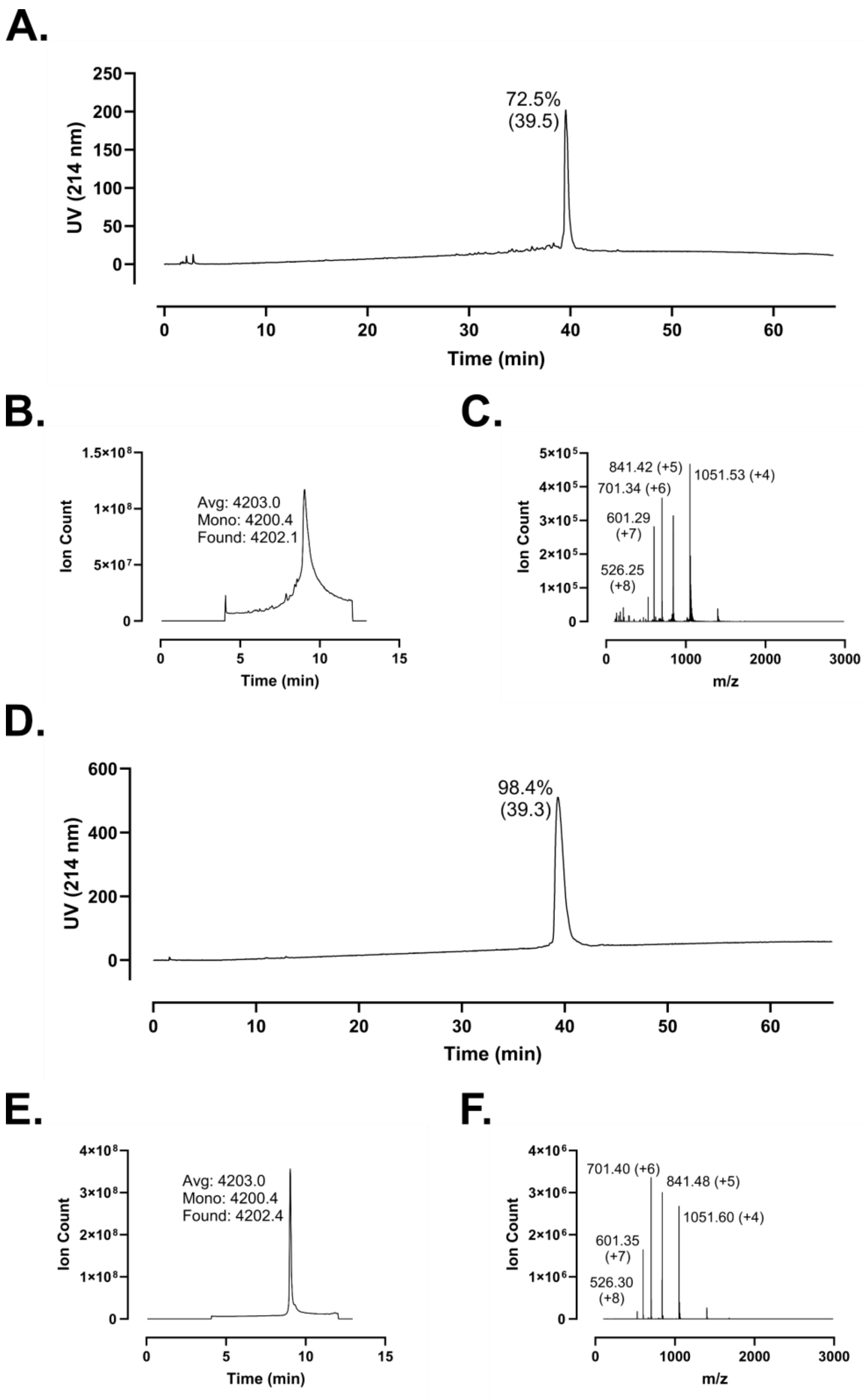
A.
250
200
150
100
50
0
UV (214 nm)
72.5%
(39.5)
0
10
20
30
40
50
60
Time (min)
B.
1.5×10⁸
1×10⁸
5×10⁷
0
Ion Count
Avg: 4203.0
Mono: 4200.4
Found: 4202.1
0
5
10
15
Time (min)
C.
5×10⁵
4×10⁵
3×10⁵
2×10⁵
1×10⁵
0
Ion Count
841.42 (+5)
1051.53 (+4)
701.34 (+6)
601.29
(+7)
526.25
(+8)
0
1000
2000
3000
m/z
D.
600
400
200
0
UV (214 nm)
98.4%
(39.3)
0
10
20
30
40
50
60
Time (min)
E.
4×10⁸
3×10⁸
2×10⁸
1×10⁸
0
Ion Count
Avg: 4203.0
Mono: 4200.4
Found: 4202.4
0
5
10
15
Time (min)
F.
4×10⁶
3×10⁶
2×10⁶
1×10⁶
0
Ion Count
701.40 (+6)
841.48 (+5)
1051.60 (+4)
601.35
(+7)
526.30
(+8)
0
1000
2000
3000
m/z

**Supplementary Figure 111 – Analytical data associated with LL-37 m6 synthesis. A.** HPLC of crude peptide. Purity by trace integration is shown along with retention time in parentheses. **B.** Total ion chromatogram of crude peptide, including the average, monoisotopic, and observed masses. **C.** Mass spectrum at peak of **B.** with selected *m/z* labelled. **D.** HPLC of peptide following a single round of purification. Peak labeling as in **A. E.** Total ion chromatogram for purified peptide. **F.** Mass spectrum at the peak of **E.** The synthesis shown here was used in multiple projects. The data presented here are thus identical to those shown in Reference 31 S156.

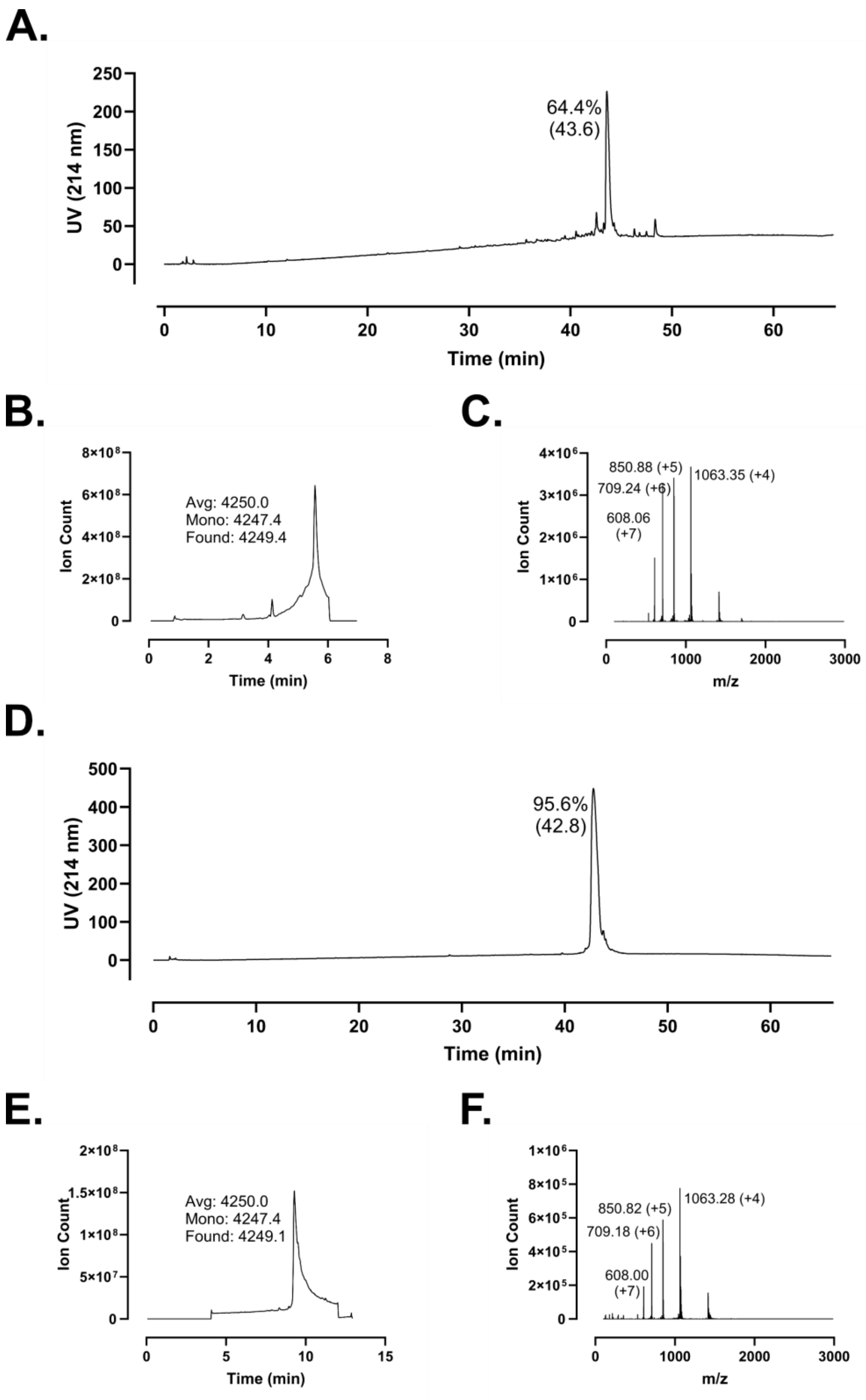
A.
64.4%
(43.6)
UV (214 nm)
Time (min)
B.
Avg: 4250.0
Mono: 4247.4
Found: 4249.4
Ion Count
Time (min)
C.
850.88 (+5)
709.24 (+6)
1063.35 (+4)
608.06
(+7)
Ion Count
m/z
D.
95.6%
(42.8)
UV (214 nm)
Time (min)
E.
Avg: 4250.0
Mono: 4247.4
Found: 4249.1
Ion Count
Time (min)
F.
850.82 (+5)
709.18 (+6)
1063.28 (+4)
608.00
(+7)
Ion Count
m/z

**Supplementary Figure 112 – Analytical data associated with LL-37 m7 synthesis. A.** HPLC of crude peptide. Purity by trace integration is shown along with retention time in parentheses. **B.** Total ion chromatogram of crude peptide, including the average, monoisotopic, and observed masses. **C.** Mass spectrum at peak of **B.** with selected *m/z* labelled. **D.** HPLC of peptide following a single round of purification. Peak labeling as in **A. E.** Total ion chromatogram for purified peptide. **F.** Mass spectrum at the peak of **E.**

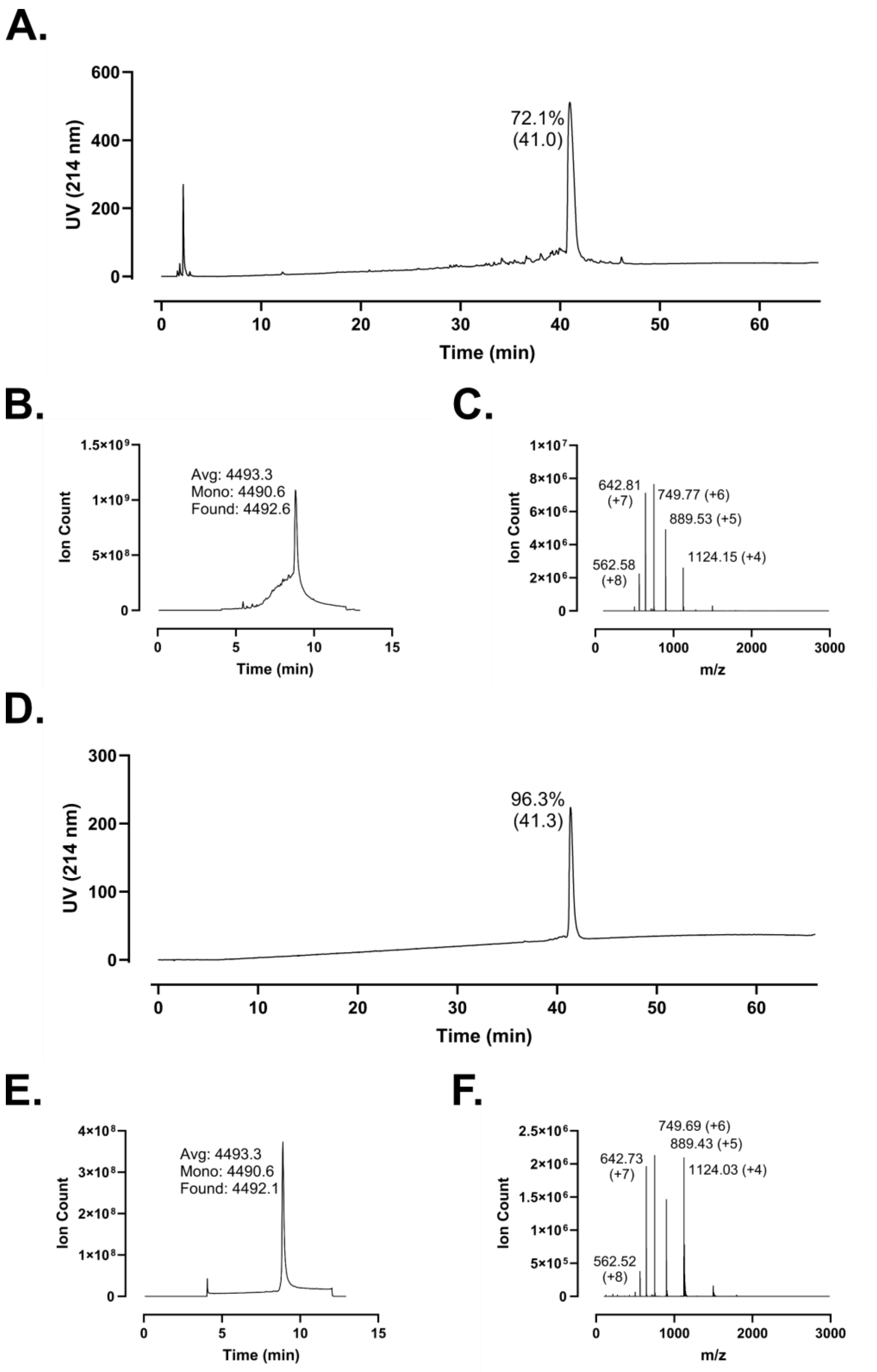
A.
600
400
200
0
UV (214 nm)
72.1%
(41.0)
0
10
20
30
40
50
60
Time (min)
B.
1.5×10⁹
1×10⁹
5×10⁸
0
Ion Count
Avg: 4493.3
Mono: 4490.6
Found: 4492.6
0
5
10
15
Time (min)
C.
1×10⁷
8×10⁶
6×10⁶
4×10⁶
2×10⁶
0
Ion Count
642.81
(+7)
749.77 (+6)
889.53 (+5)
562.58
(+8)
1124.15 (+4)
0
1000
2000
3000
m/z
D.
300
200
100
0
UV (214 nm)
96.3%
(41.3)
0
10
20
30
40
50
60
Time (min)
E.
4×10⁸
3×10⁸
2×10⁸
1×10⁸
0
Ion Count
Avg: 4493.3
Mono: 4490.6
Found: 4492.1
0
5
10
15
Time (min)
F.
2.5×10⁶
2×10⁶
1.5×10⁶
1×10⁶
5×10⁵
0
Ion Count
749.69 (+6)
889.43 (+5)
642.73
(+7)
1124.03 (+4)
562.52
(+8)
0
1000
2000
3000
m/z

**Supplementary Figure 113 – Analytical data associated with LL-37 synthesis. A.** HPLC of crude peptide. Purity by trace integration is shown along with retention time in parentheses. **B.** Total ion chromatogram of crude peptide, including the average, monoisotopic, and observed masses. **C.** Mass spectrum at peak of **B.** with selected *m/z* labelled. **D.** HPLC of peptide following a single round of purification. Peak labeling as in **A. E.** Total ion chromatogram for purified peptide. **F.** Mass spectrum at the peak of **E.** The synthesis shown here was used in multiple projects. The data presented here are thus identical to the same characterization shown in References 30 S151 and 31 S157.

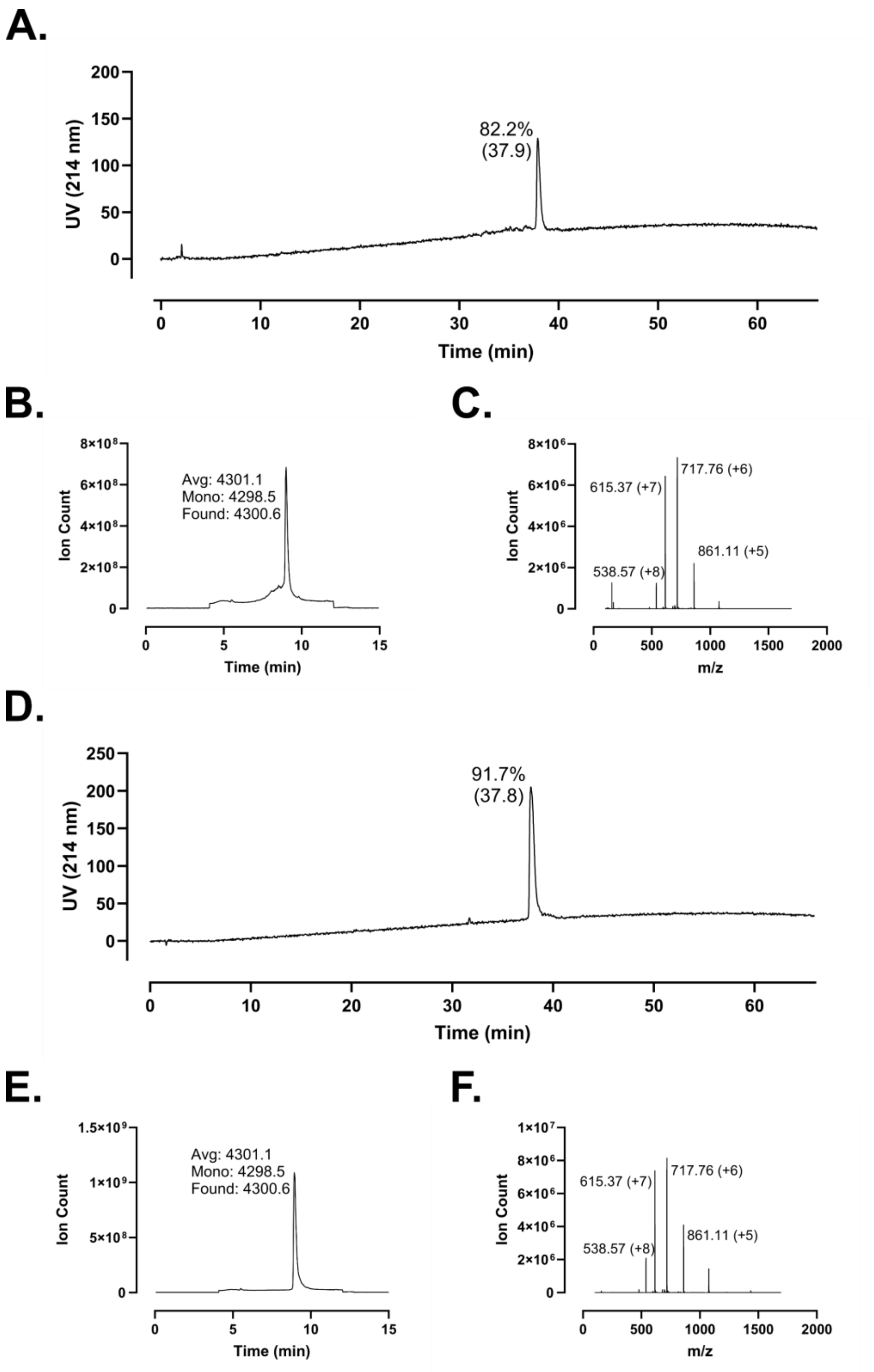
A.
UV (214 nm)
Time (min)
82.2%
(37.9)
B.
Ion Count
Time (min)
Avg: 4301.1
Mono: 4298.5
Found: 4300.6
C.
Ion Count
m/z
717.76 (+6)
615.37 (+7)
861.11 (+5)
538.57 (+8)
D.
UV (214 nm)
Time (min)
91.7%
(37.8)
E.
Ion Count
Time (min)
Avg: 4301.1
Mono: 4298.5
Found: 4300.6
F.
Ion Count
m/z
717.76 (+6)
615.37 (+7)
861.11 (+5)
538.57 (+8)

**Supplementary Figure 114 – Analytical data associated with LL-37 m8 synthesis. A.** HPLC of crude peptide. Purity by trace integration is shown along with retention time in parentheses. **B.** Total ion chromatogram of crude peptide, including the average, monoisotopic, and observed masses. **C.** Mass spectrum at peak of **B.** with selected *m/z* labelled. **D.** HPLC of peptide following a single round of purification. Peak labeling as in **A. E.** Total ion chromatogram for purified peptide. **F.** Mass spectrum at the peak of **E.**

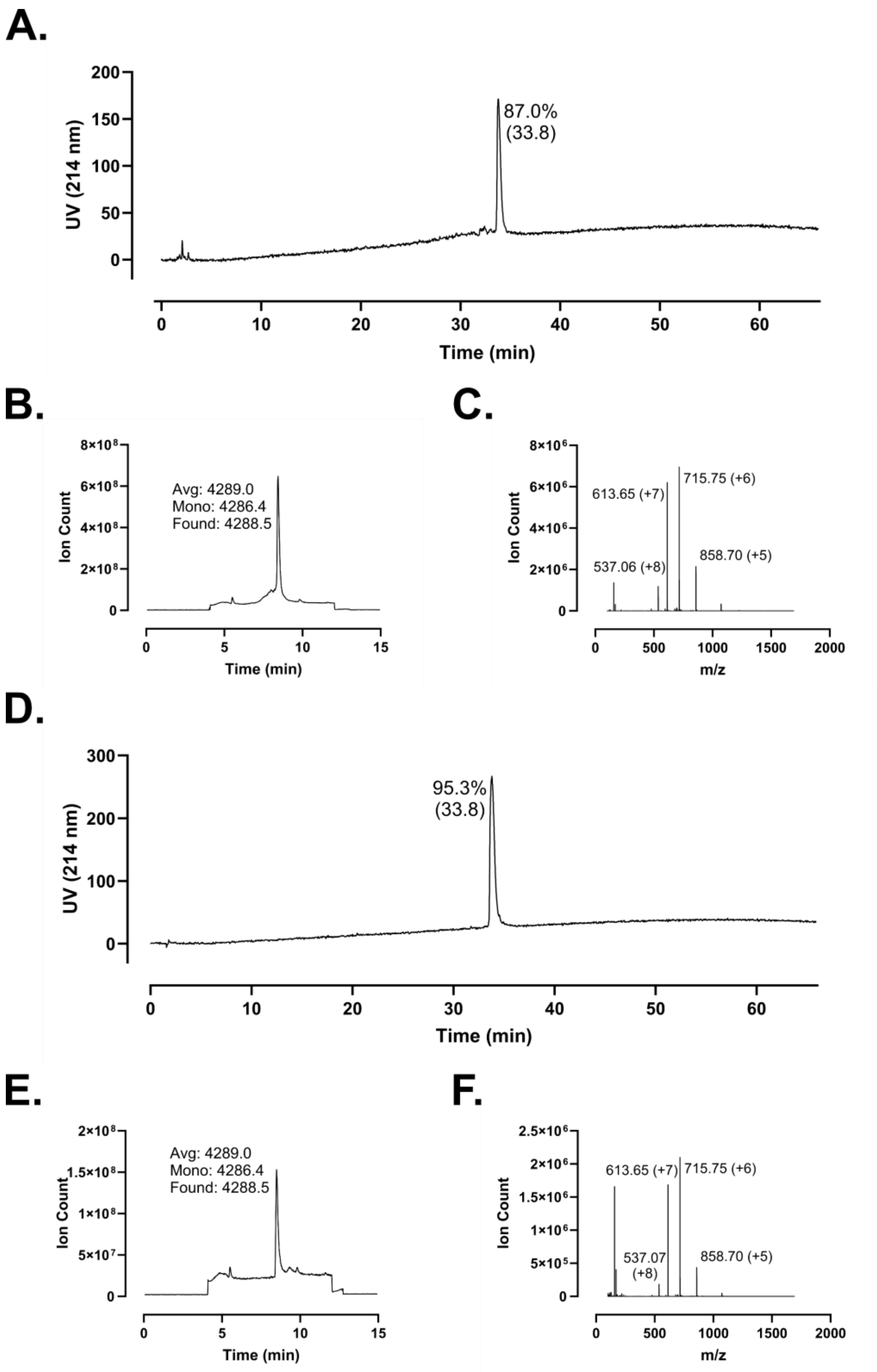
A.
UV (214 nm)
Time (min)
87.0%
(33.8)
B.
Ion Count
Time (min)
Avg: 4289.0
Mono: 4286.4
Found: 4288.5
C.
Ion Count
m/z
613.65 (+7)
715.75 (+6)
537.06 (+8)
858.70 (+5)
D.
UV (214 nm)
Time (min)
95.3%
(33.8)
E.
Ion Count
Time (min)
Avg: 4289.0
Mono: 4286.4
Found: 4288.5
F.
Ion Count
m/z
613.65 (+7)
715.75 (+6)
537.07
(+8)
858.70 (+5)

**Supplementary Figure 115 – Analytical data associated with LL-37 m9 synthesis. A.** HPLC of crude peptide. Purity by trace integration is shown along with retention time in parentheses. **B.** Total ion chromatogram of crude peptide, including the average, monoisotopic, and observed masses. **C.** Mass spectrum at peak of **B.** with selected *m/z* labelled. **D.** HPLC of peptide following a single round of purification. Peak labeling as in **A. E.** Total ion chromatogram for purified peptide. **F.** Mass spectrum at the peak of **E.**

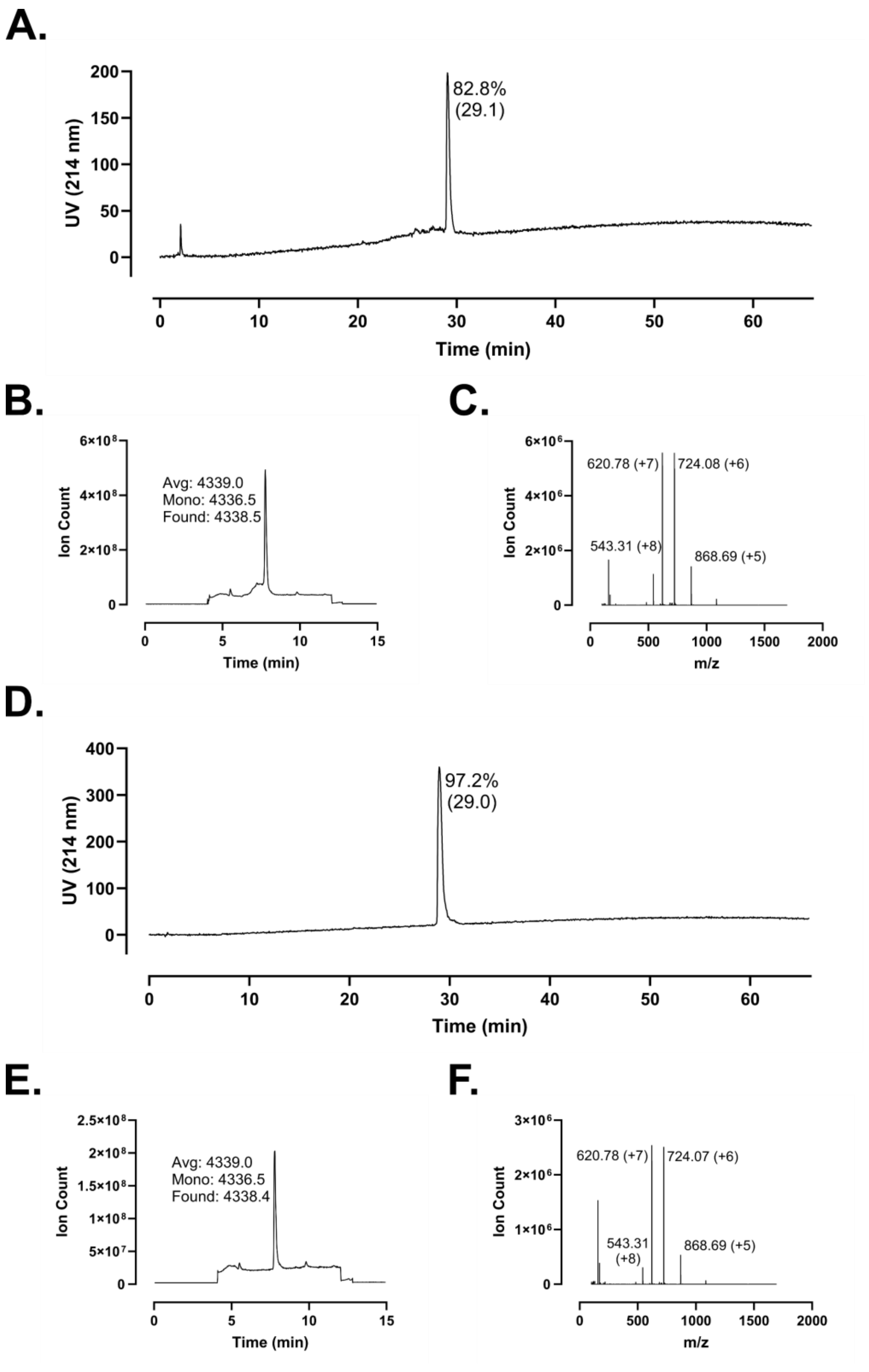

A.
200
150
100
50
0
UV (214 nm)
82.8%
(29.1)
0
10
20
30
40
50
60
Time (min)
B.
6×10⁸
4×10⁸
2×10⁸
0
Ion Count
Avg: 4339.0
Mono: 4336.5
Found: 4338.5
0
5
10
15
Time (min)
C.
6×10⁶
4×10⁶
2×10⁶
0
Ion Count
620.78 (+7)
724.08 (+6)
543.31 (+8)
868.69 (+5)
0
500
1000
1500
2000
m/z
D.
400
300
200
100
0
UV (214 nm)
97.2%
(29.0)
0
10
20
30
40
50
60
Time (min)
E.
2.5×10⁸
2×10⁸
1.5×10⁸
1×10⁸
5×10⁷
0
Ion Count
Avg: 4339.0
Mono: 4336.5
Found: 4338.4
0
5
10
15
Time (min)
F.
3×10⁶
2×10⁶
1×10⁶
0
Ion Count
620.78 (+7)
724.07 (+6)
543.31
(+8)
868.69 (+5)
0
500
1000
1500
2000
m/z

**Supplementary Figure 116 – Analytical data associated with LL-37 m10 synthesis. A.** HPLC of crude peptide. Purity by trace integration is shown along with retention time in parentheses. **B.** Total ion chromatogram of crude peptide, including the average, monoisotopic, and observed masses. **C.** Mass spectrum at peak of **B.** with selected *m/z* labelled. **D.** HPLC of peptide following a single round of purification. Peak labeling as in **A. E.** Total ion chromatogram for purified peptide. **F.** Mass spectrum at the peak of **E.**

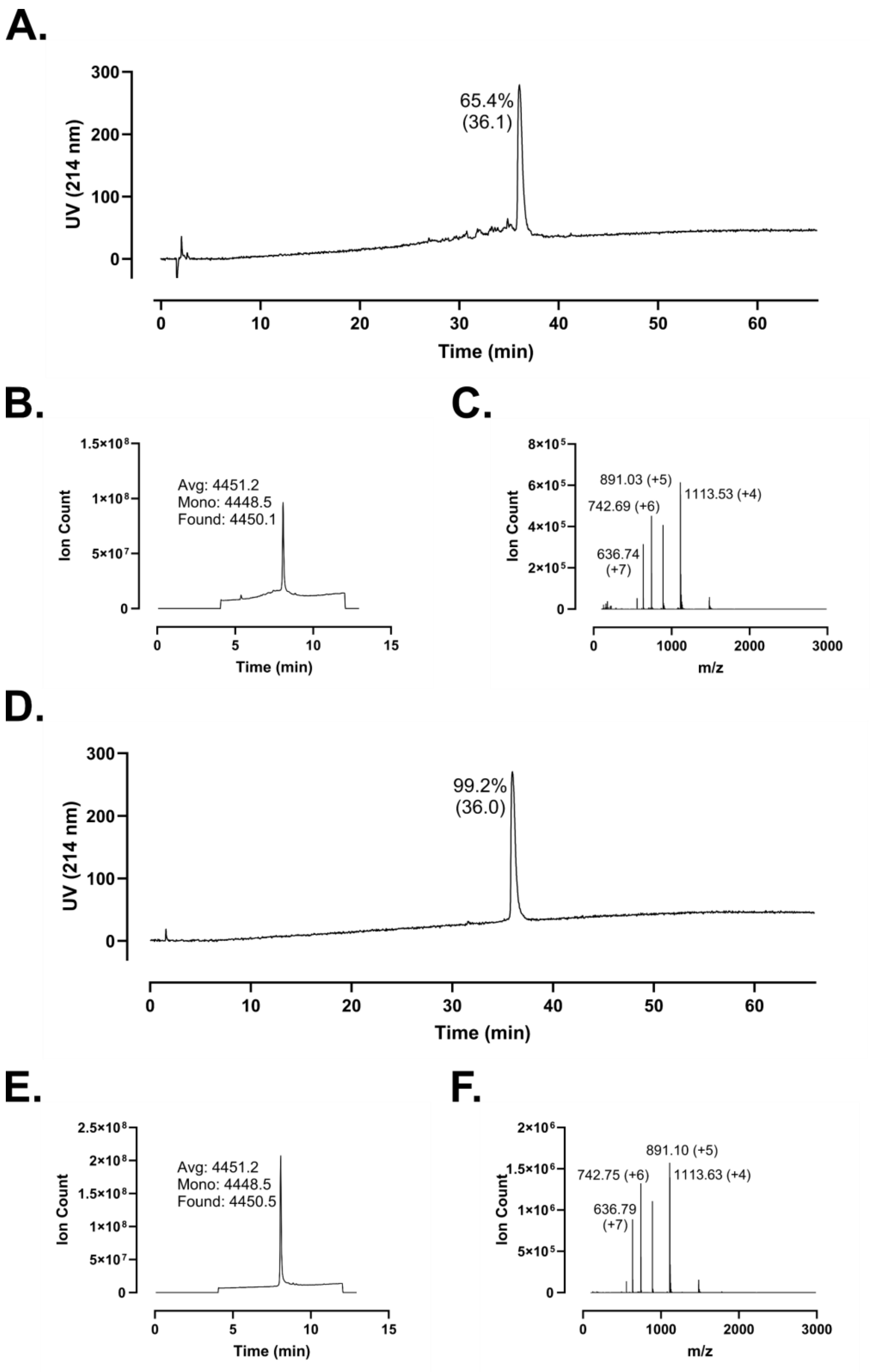

A.
300
200
100
0
UV (214 nm)
65.4%
(36.1)
0
10
20
30
40
50
60
Time (min)
B.
1.5×10⁸
1×10⁸
5×10⁷
0
Ion Count
Avg: 4451.2
Mono: 4448.5
Found: 4450.1
0
5
10
15
Time (min)
C.
8×10⁵
6×10⁵
4×10⁵
2×10⁵
0
Ion Count
891.03 (+5)
742.69 (+6)
1113.53 (+4)
636.74
(+7)
0
1000
2000
3000
m/z
D.
300
200
100
0
UV (214 nm)
99.2%
(36.0)
0
10
20
30
40
50
60
Time (min)
E.
2.5×10⁸
2×10⁸
1.5×10⁸
1×10⁸
5×10⁷
0
Ion Count
Avg: 4451.2
Mono: 4448.5
Found: 4450.5
0
5
10
15
Time (min)
F.
2×10⁶
1.5×10⁶
1×10⁶
5×10⁵
0
Ion Count
891.10 (+5)
742.75 (+6)
1113.63 (+4)
636.79
(+7)
0
1000
2000
3000
m/z

**Supplementary Figure 117 – Analytical data associated with LL-37 m11 synthesis. A.** HPLC of crude peptide. Purity by trace integration is shown along with retention time in parentheses. **B.** Total ion chromatogram of crude peptide, including the average, monoisotopic, and observed masses. **C.** Mass spectrum at peak of **B.** with selected *m/z* labelled. **D.** HPLC of peptide following a single round of purification. Peak labeling as in **A. E.** Total ion chromatogram for purified peptide. **F.** Mass spectrum at the peak of **E.**

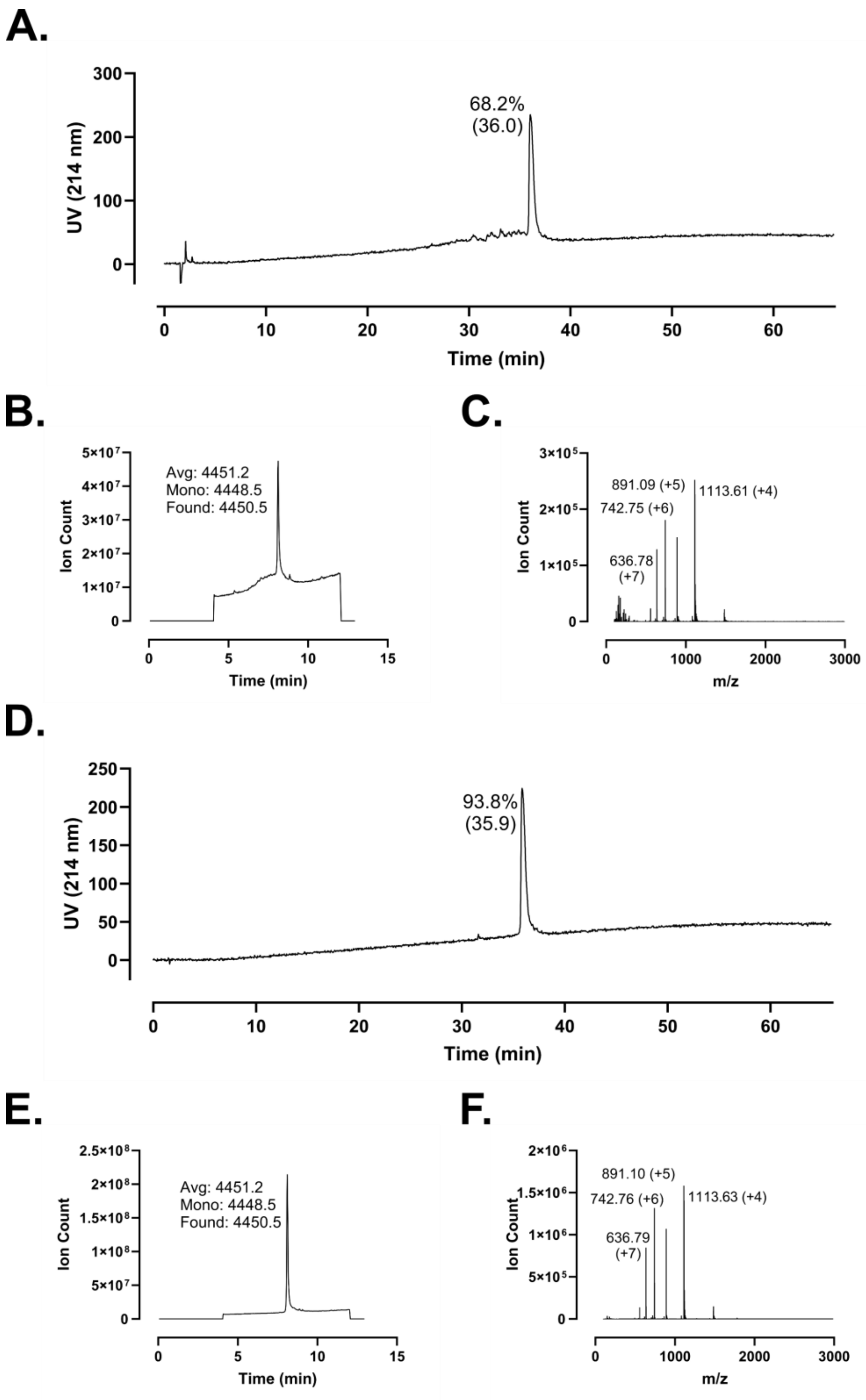
A.
68.2%
(36.0)
UV (214 nm)
Time (min)
B.
Avg: 4451.2
Mono: 4448.5
Found: 4450.5
Ion Count
Time (min)
C.
891.09 (+5)
1113.61 (+4)
742.75 (+6)
636.78
(+7)
Ion Count
m/z
D.
93.8%
(35.9)
UV (214 nm)
Time (min)
E.
Avg: 4451.2
Mono: 4448.5
Found: 4450.5
Ion Count
Time (min)
F.
891.10 (+5)
742.76 (+6)
1113.63 (+4)
636.79
(+7)
Ion Count
m/z

**Supplementary Figure 118 – Analytical data associated with LL-37 m12 synthesis. A.** HPLC of crude peptide. Purity by trace integration is shown along with retention time in parentheses. **B.** Total ion chromatogram of crude peptide, including the average, monoisotopic, and observed masses. **C.** Mass spectrum at peak of **B.** with selected *m/z* labelled. **D.** HPLC of peptide following a single round of purification. Peak labeling as in **A. E.** Total ion chromatogram for purified peptide. **F.** Mass spectrum at the peak of **E.**

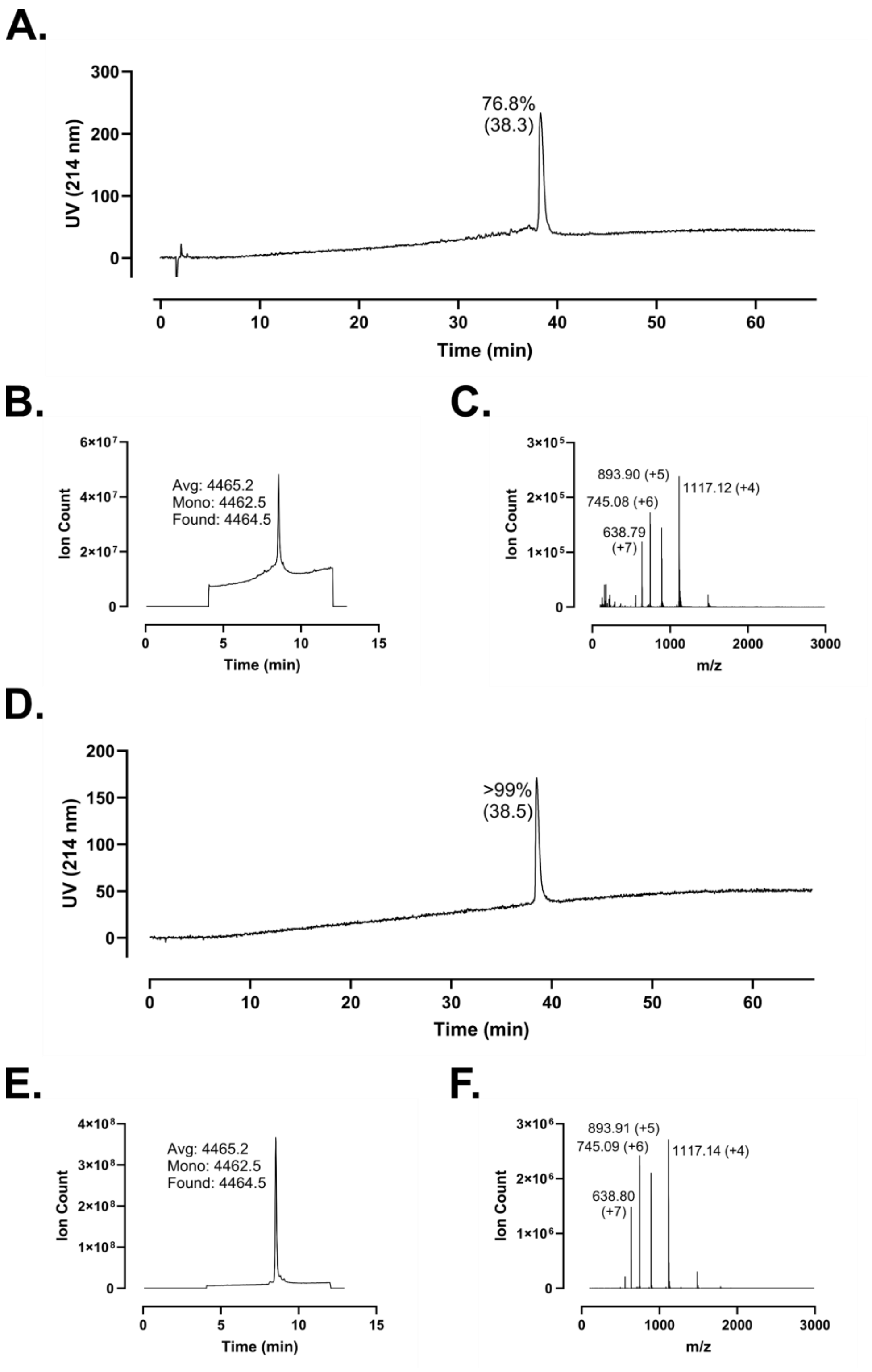
A.
76.8%
(38.3)
UV (214 nm)
Time (min)
B.
Avg: 4465.2
Mono: 4462.5
Found: 4464.5
Ion Count
Time (min)
C.
893.90 (+5)
1117.12 (+4)
745.08 (+6)
638.79
(+7)
Ion Count
m/z
D.
>99%
(38.5)
UV (214 nm)
Time (min)
E.
Avg: 4465.2
Mono: 4462.5
Found: 4464.5
Ion Count
Time (min)
F.
893.91 (+5)
745.09 (+6)
1117.14 (+4)
638.80
(+7)
Ion Count
m/z

**Supplementary Figure 119 – Analytical data associated with LL-37 m13 synthesis. A.** HPLC of crude peptide. Purity by trace integration is shown along with retention time in parentheses. **B.** Total ion chromatogram of crude peptide, including the average, monoisotopic, and observed masses. **C.** Mass spectrum at peak of **B.** with selected *m/z* labelled. **D.** HPLC of peptide following a single round of purification. Peak labeling as in **A. E.** Total ion chromatogram for purified peptide. **F.** Mass spectrum at the peak of **E.**

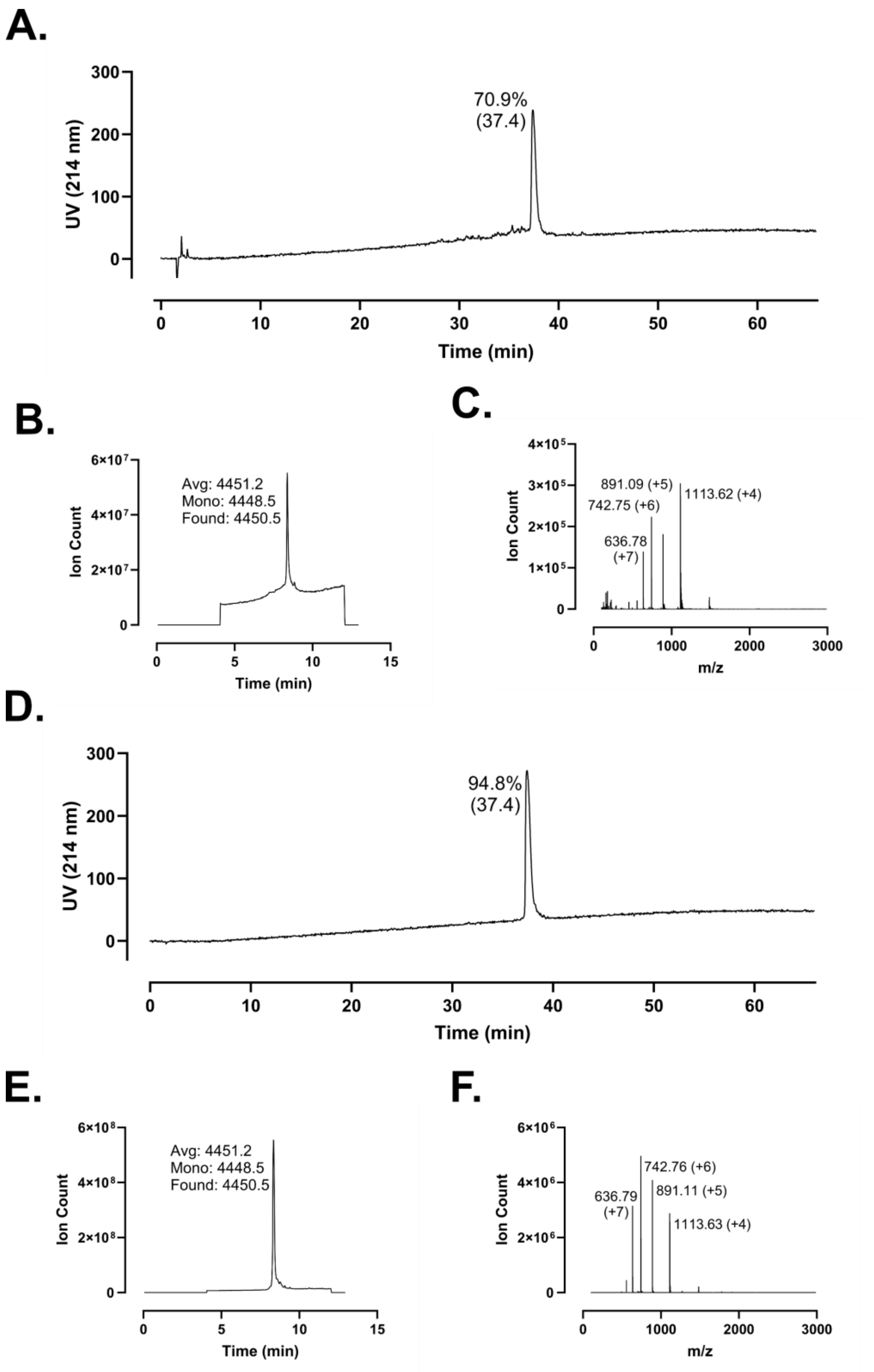
A.
UV (214 nm)
300
200
100
0
70.9%
(37.4)
0
10
20
30
40
50
60
Time (min)
B.
Ion Count
6×10^7
4×10^7
2×10^7
0
Avg: 4451.2
Mono: 4448.5
Found: 4450.5
0
5
10
15
Time (min)
C.
Ion Count
4×10^5
3×10^5
2×10^5
1×10^5
0
891.09 (+5)
742.75 (+6)
1113.62 (+4)
636.78
(+7)
0
1000
2000
3000
m/z
D.
UV (214 nm)
300
200
100
0
94.8%
(37.4)
0
10
20
30
40
50
60
Time (min)
E.
Ion Count
6×10^8
4×10^8
2×10^8
0
Avg: 4451.2
Mono: 4448.5
Found: 4450.5
0
5
10
15
Time (min)
F.
Ion Count
6×10^6
4×10^6
2×10^6
0
742.76 (+6)
891.11 (+5)
636.79
(+7)
1113.63 (+4)
0
1000
2000
3000
m/z

**Supplementary Figure 120 – Analytical data associated with LL-37 m14 synthesis. A.** HPLC of crude peptide. Purity by trace integration is shown along with retention time in parentheses. **B.** Total ion chromatogram of crude peptide, including the average, monoisotopic, and observed masses. **C.** Mass spectrum at peak of **B.** with selected *m/z* labelled. **D.** HPLC of peptide following a single round of purification. Peak labeling as in **A. E.** Total ion chromatogram for purified peptide. **F.** Mass spectrum at the peak of **E.**

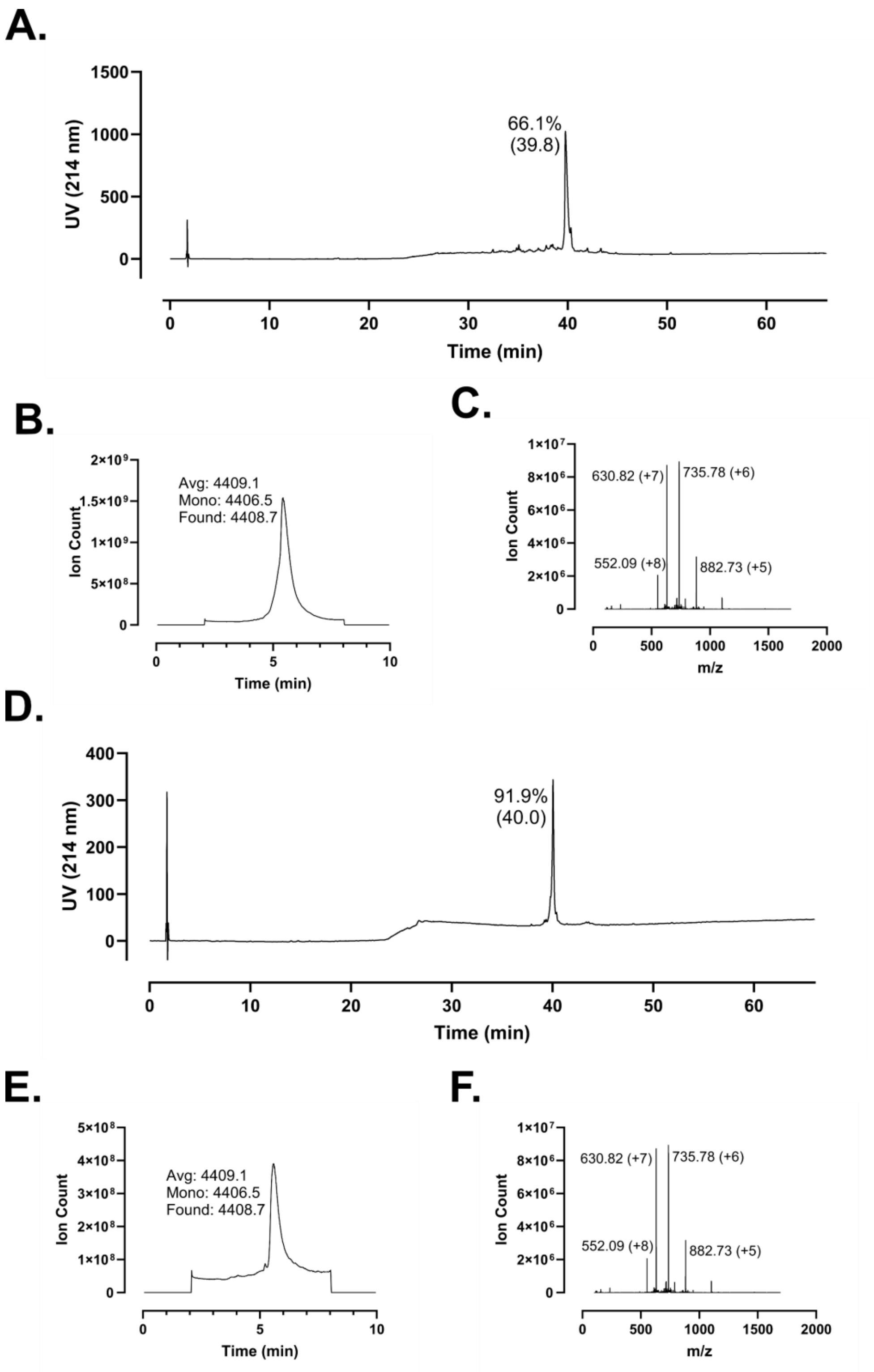
A.
66.1%
(39.8)
UV (214 nm)
Time (min)
B.
Avg: 4409.1
Mono: 4406.5
Found: 4408.7
Ion Count
Time (min)
C.
630.82 (+7)
735.78 (+6)
552.09 (+8)
882.73 (+5)
Ion Count
m/z
D.
91.9%
(40.0)
UV (214 nm)
Time (min)
E.
Avg: 4409.1
Mono: 4406.5
Found: 4408.7
Ion Count
Time (min)
F.
630.82 (+7)
735.78 (+6)
552.09 (+8)
882.73 (+5)
Ion Count
m/z

**Supplementary Figure 121 – Analytical data associated with LL-37 m15 synthesis. A.** HPLC of crude peptide. Purity by trace integration is shown along with retention time in parentheses. **B.** Total ion chromatogram of crude peptide, including the average, monoisotopic, and observed masses. **C.** Mass spectrum at peak of **B.** with selected *m/z* labelled. **D.** HPLC of peptide following a single round of purification. Peak labeling as in **A.** **E.** Total ion chromatogram for purified peptide. **F.** Mass spectrum at the peak of **E.**

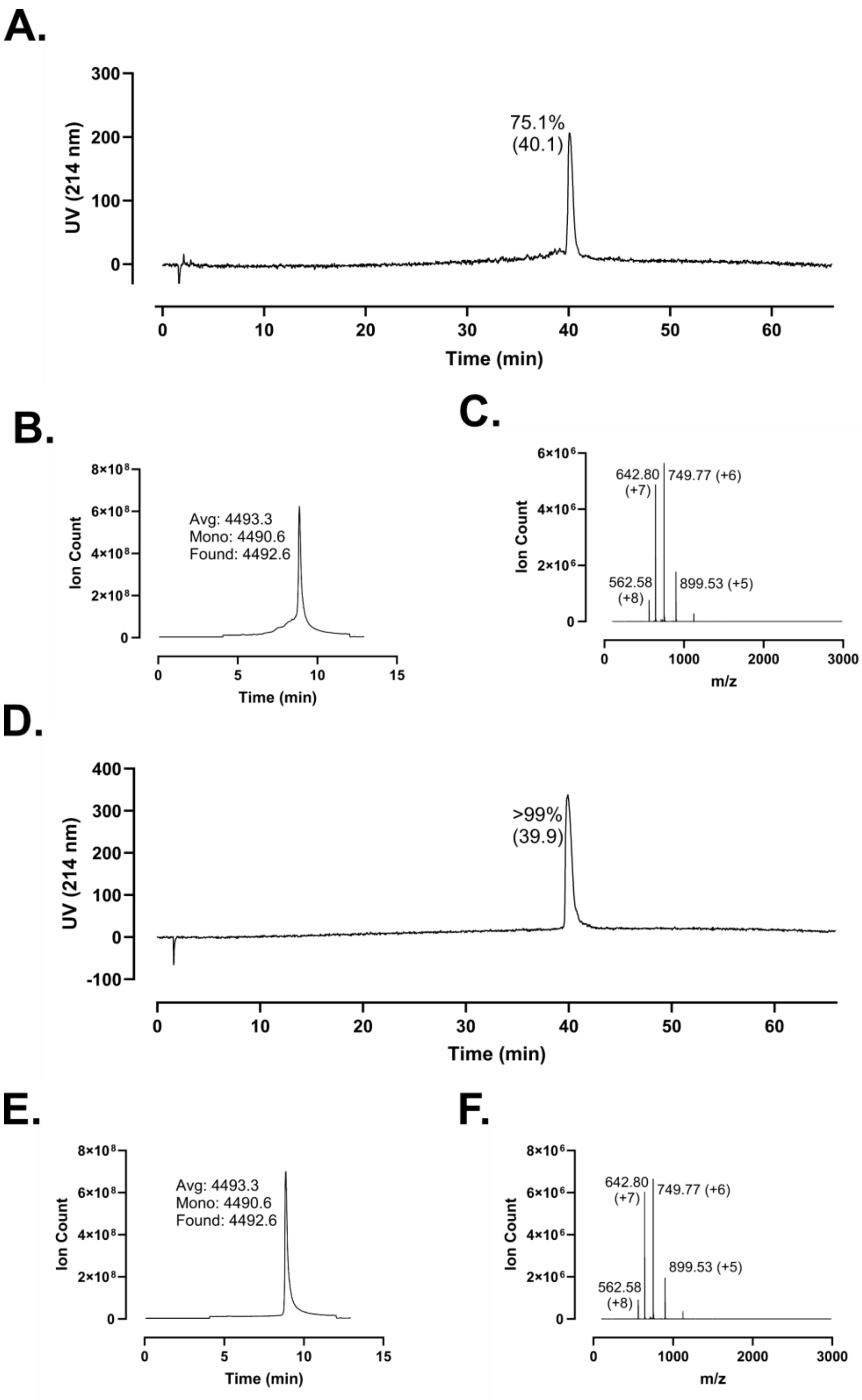
A.
UV (214 nm)
75.1%
(40.1)
Time (min)
B.
Ion Count
Avg: 4493.3
Mono: 4490.6
Found: 4492.6
Time (min)
C.
Ion Count
642.80
(+7)
749.77 (+6)
562.58
(+8)
899.53 (+5)
m/z
D.
UV (214 nm)
>99%
(39.9)
Time (min)
E.
Ion Count
Avg: 4493.3
Mono: 4490.6
Found: 4492.6
Time (min)
F.
Ion Count
642.80
(+7)
749.77 (+6)
562.58
(+8)
899.53 (+5)
m/z

**Supplementary Figure 122 – Analytical data associated with LL-37 synthesis. A.** HPLC of crude peptide. Purity by trace integration is shown along with retention time in parentheses. **B.** Total ion chromatogram of crude peptide, including the average, monoisotopic, and observed masses. **C.** Mass spectrum at peak of **B.** with selected *m/z* labelled. **D.** HPLC of peptide following a single round of purification. Peak labeling as in **A. E.** Total ion chromatogram for purified peptide. **F.** Mass spectrum at the peak of **E.** The synthesis here is distinct from that shown in **Supplementary Figure 113**.

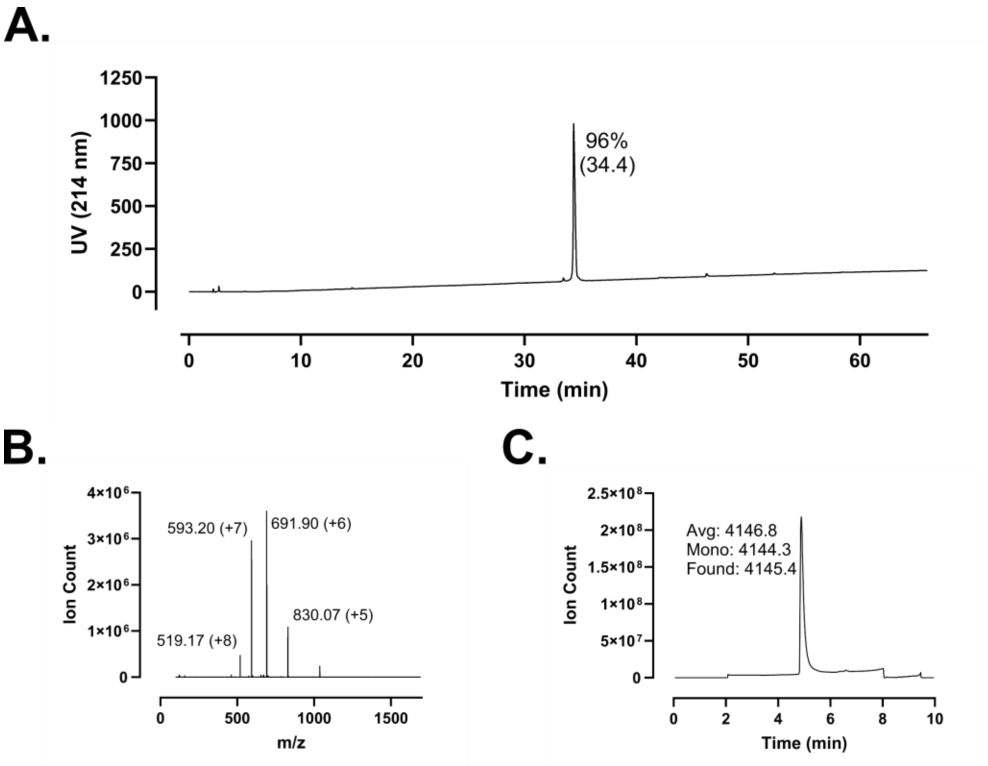
A.
UV (214 nm)
1250
1000
750
500
250
0
96%
(34.4)
0
10
20
30
40
50
60
Time (min)
B.
Ion Count
4×10⁶
3×10⁶
2×10⁶
1×10⁶
0
593.20 (+7)
691.90 (+6)
830.07 (+5)
519.17 (+8)
0
500
1000
1500
m/z
C.
Ion Count
2.5×10⁸
2×10⁸
1.5×10⁸
1×10⁸
5×10⁷
0
Avg: 4146.8
Mono: 4144.3
Found: 4145.4
0
2
4
6
8
10
Time (min)

**Supplementary Figure 123 – Analytical data associated with LL-37 m4 synthesis. A.** HPLC of peptide following a single round of purification. Purity by trace integration is shown along with retention time in parentheses. **B.** Total ion chromatogram of peptide as in **A**, including the average, monoisotopic, and observed masses. **C.** Mass spectrum at peak of **B.** with selected *m/z* labelled. The synthesis shown here is distinct from that shown in **Supplementary Figure 109**.

A.

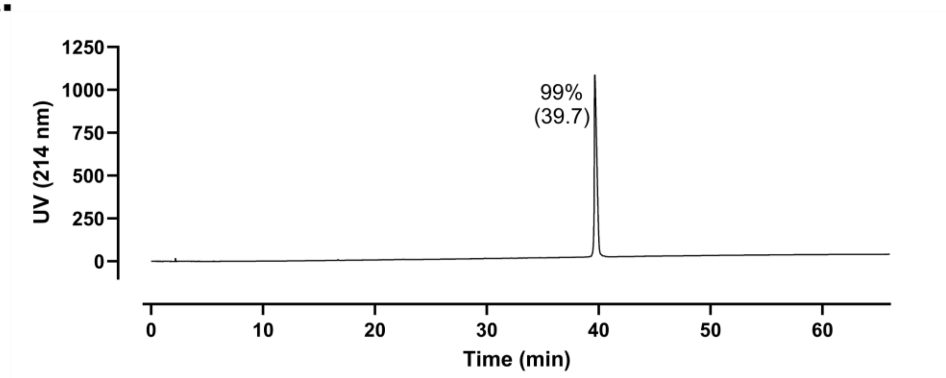

UV (214 nm)
1250
1000
750
500
250
0
99%
(39.7)
0
10
20
30
40
50
60
Time (min)


B.

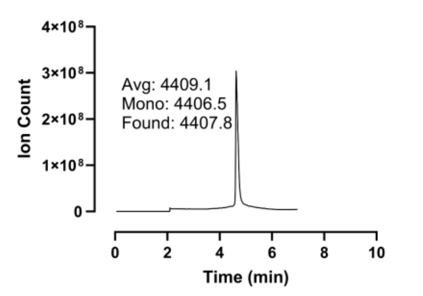

Ion Count
4×10⁸
3×10⁸
2×10⁸
1×10⁸
0
Avg: 4409.1
Mono: 4406.5
Found: 4407.8
0
2
4
6
8
10
Time (min)


C.

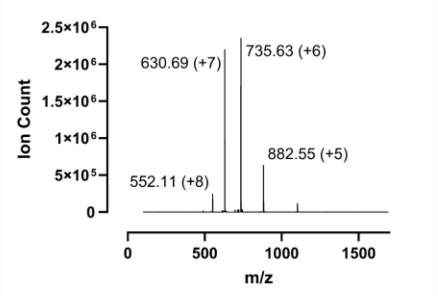

Ion Count
2.5×10⁶
2×10⁶
1.5×10⁶
1×10⁶
5×10⁵
0
630.69 (+7)
735.63 (+6)
882.55 (+5)
552.11 (+8)
0
500
1000
1500
m/z

**Supplementary Figure 124 – Analytical data associated with LL-37 m15 synthesis. A.** HPLC of peptide following a single round of purification. Purity by trace integration is shown along with retention time in parentheses. **B.** Total ion chromatogram of peptide as in **A**, including the average, monoisotopic, and observed masses. **C.** Mass spectrum at peak of **B.** with selected *m/z* labelled. The synthesis shown here is distinct from that shown in **Supplementary Figure 121**.

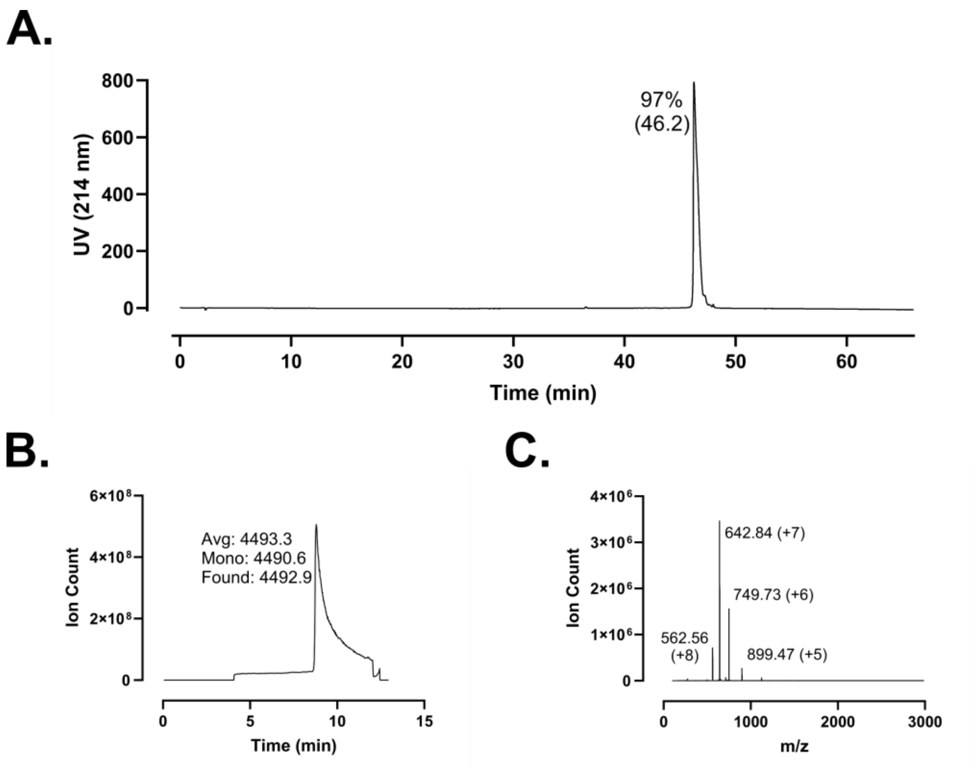
A.
97%
(46.2)
UV (214 nm)
Time (min)
B.
Avg: 4493.3
Mono: 4490.6
Found: 4492.9
Ion Count
Time (min)
C.
642.84 (+7)
749.73 (+6)
562.56
(+8)
899.47 (+5)
Ion Count
m/z

**Supplementary Figure 125 – Analytical data associated with LL-37 synthesis. A.** HPLC of peptide following a single round of purification. Purity by trace integration is shown along with retention time in parentheses. **B.** Total ion chromatogram of peptide as in **A**, including the average, monoisotopic, and observed masses. **C.** Mass spectrum at peak of **B.** with selected *m/z* labelled. The synthesis shown here is distinct from that shown in **Supplementary Figures 113** and **122**. The synthesis shown here was used in multiple projects. The data presented here are thus identical to the same characterization shown in Reference 30 S205.

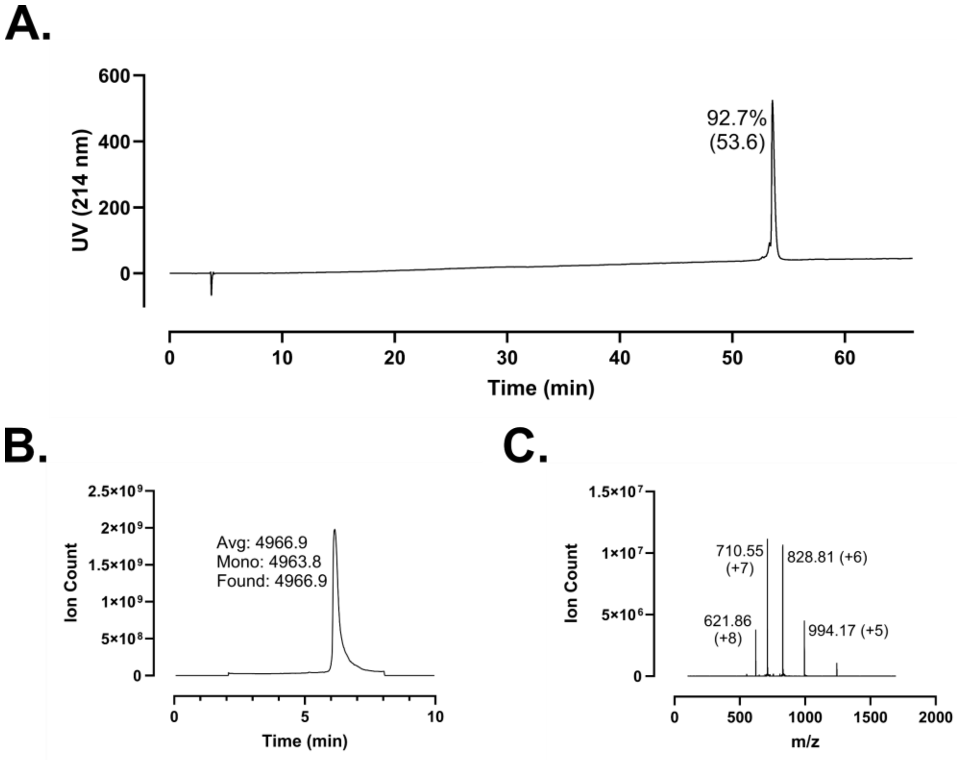
A.
92.7%
(53.6)
UV (214 nm)
Time (min)
B.
Avg: 4966.9
Mono: 4963.8
Found: 4966.9
Ion Count
Time (min)
C.
710.55
(+7)
828.81 (+6)
621.86
(+8)
994.17 (+5)
Ion Count
m/z

**Supplementary Figure 126 – Analytical data associated with PEG4-LL-37 synthesis. A.** HPLC of peptide following a single round of purification. Purity by trace integration is shown along with retention time in parentheses. **B.** Total ion chromatogram of peptide as in **A**, including the average, monoisotopic, and observed masses. **C.** Mass spectrum at peak of **B.** with selected *m/z* labelled.
